# Fast, accurate and scalable normalization for RNA sequencing data with the RUVprps software package

**DOI:** 10.64898/2026.01.28.702384

**Authors:** Marie Trussart, Momeneh Foroutan, Michael Milton, Himisha Beltran, Terence P Speed, Ramyar Molania

**Author notes:** These authors contributed equally (Marie Trussart, Momeneh Foroutan, Michael Milton). Corresponding author, Ramyar Molania.

## Abstract

Unwanted variation refers to any source of variability in the data that can compromise down-stream analysis. Effective removal of such variation from gene expression data is essential to derive accurate and meaningful biological results. We refer to this process as normalization. Data may come from a single study or from multiple studies with different sources of unwanted variation. We have previously developed the RUV-III method for normalizing omics data with a strong focus on transcriptomics. Initially, we introduced RUV-III for the normalization of Nanostring nCounter gene expression data, utilizing genuine technical replicates and pseudo-replicates as control samples. Subsequently, we proposed RUV-III with pseudo-replicates of pseudo-samples (PRPS), and which demonstrated its potential in mitigating the effects of different sources of unwanted variation in large and complex RNA-seq studies. To enhance accessibility and performance of this method, we present a new comprehensive R package named RUVprps. The package offers over 100 functions including ones for assessing variation in both biological and unwanted variables, an automated RUV-III normalization process, and metrics for evaluating the effectiveness of the resulting normalizations. Further, it introduces several new features such as ways of identifying unknown sources of unwanted variation, strategies to identify suitable negative control genes, and methods for generating PRPS when information on the biological and unwanted variation is unavailable. The package also implements a faster approach to RUV-III normalization, streamlining its application to large RNA-seq datasets. Our freely available R package and normalization assessment pipeline can help find effective data normalization methods for new data and help benchmark new methods.

## Introduction

RNA-seq data is almost always affected by sources of unwanted variation, irrespective of sample size [1–4]. RNA-seq data can be significantly influenced by variations in library size and tissue composition such as tumor purity in cancer studies, as well as batch effects when the data comes from multiple batches [3]. It is crucial to note that distinct sources of unwanted variation may affect distinct subsets of genes in varying ways [3, 5, 6]. Effective identification and removal of the impact of this variation is essential to derive accurate and meaningful biological conclusions from the data.

Previously, we introduced a normalization method called removing unwanted variation III (RUV-III), which is designed to remove different sources of unwanted variation in omics data, with a particular emphasis on transcriptomics [3, 5, 7–10]. RUV-III leverages both control samples and control genes, termed negative control genes (NCG), to effectively estimate both known and unknown sources of unwanted variation and remove them from the data. It uses gene-wise expression differences across control samples, along with variation in NCG, to perform effective normalization. Initially, control samples were technical replicates, which are the same sample profiled across anticipated sources of unwanted variation in a study. Any gene-wise expression differences between sets of technical replicates are purely unwanted variation. However, there are situations where technical replicates are unavailable or where they are available, but in insufficient numbers or in sub-optimal locations in relation to the sources of unwanted variation. More importantly, some sources of unwanted variation, such as tumor purity, cannot be removed by technical replicates [3]. To overcome these limitations, we previously proposed a novel approach called PRPS, where pseudo-replicates (PR) of pseudo-samples (PS) play the role of technical replicates in the original RUV-III method [3, 5]. Pseudo-samples are *in silico* samples derived from small groups of samples that are roughly homogeneous with respect to unwanted variation and biology. Two or more pseudo-samples with the same biology coming from (“spanning”) two or more batches will be regarded as a pseudo-replicate set. The gene expression differences between such pseudo-samples will largely be unwanted variation. We have previously demonstrated that RUV-III is highly robust to poorly chosen samples to create PRPS data [3].

Here, we present a user-friendly R package and a comprehensive pipeline designed to streamline all the essential steps involved in the normalization of RNA-seq data. This includes evaluating both biological and unwanted variation, applying RUV-III normalization, and assessing the effectiveness of the resulting normalizations. Moreover, RUVprps presents four novel features that address various limitations encountered in the utilization of RUV-III with PRPS [3]. First, it enables estimation of unknown sources of variation within RNA-seq data, which is particularly helpful in scenarios where some or all sources of unwanted variation are unknown. An example is when we use publicly available data where no information on unwanted variation is available. As a second feature, the package facilitates the identification of NCG and the generation of PRPS in situations where biological variables are entirely unknown. We have previously shown the satisfactory performance of RUV-III-PRPS normalization when biological variables are only partially known [3]. However, there might be situations where none at all are known. We refer to these steps as the “unsupervised” implementation of the method. Third, the package implements a faster approach to RUV-III normalization, enabling easy execution of the method with different parameters, NCG and PRPS sets. Finally, the package summarizes all the evaluation assessments and plots into a single score for each individual normalization. This enables users to rank normalizations based on their overall performance and select a suitable one for downstream analysis. It should be noted that the package can also accommodate technical replicates as control samples for RUV-III normalization if they are available.

We use several publicly available RNA-seq datasets and design different real-world scenarios to demonstrate the effectiveness of both the supervised and the unsupervised applications of RUV-III with PRPS. Additionally, we show how our variation and normalization evaluations can aid in achieving a satisfactory normalization and determining the most effective method.

## Results

### Application of RUVprps

The RUVprps R package is a general-purpose computing tool designed for the end-to-end normalization of high-dimensional gene expression data using the RUV-III method (Methods and Fig. 1). RUVprps provides a comprehensive set of gene-level and global statistical and computational tools for identifying and assessing sources of unwanted variation (Methods), as well as evaluating the performance of normalization methods (Supplementary File, Fig. 1b). RUVprps can be applied to data from single or multiple studies, regardless of sample size, and is compatible with both whole-transcriptome and targeted RNA-seq experiments that profile a specific panel of genes. We have previously shown excellent performance of RUV-III on Nanostring nCounter gene expression data with a 250 gene panel that were highly variable with respect to the biological conditions of interest [5]. RUV-III requires both control samples and NCG to estimate and remove known and unknown sources of unwanted variation from the data. The control samples may include genuine technical replicates (TR), pseudo-replicates (PR) [5], pseudo-replicates of pseudo-samples (PRPS)[3] or a combination of these (Methods). NCG can be a pre-selected set of genes such as housekeeping genes. However, if those are not affected by unwanted variation in the data, NCG can be obtained using different data-driven approaches provided by RUVprps.

**Figure 1:**
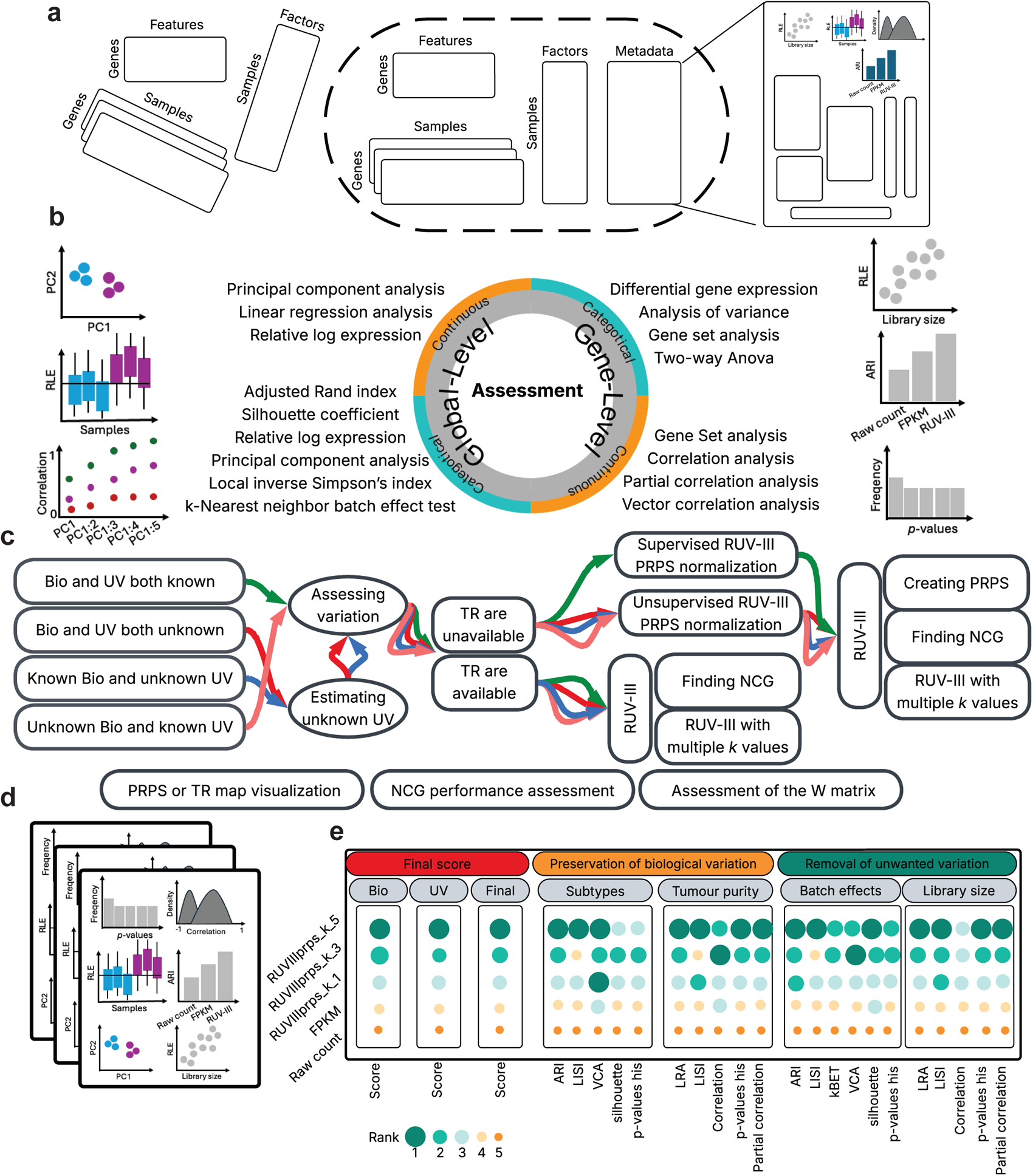
Overview of the RUVprps R package. a) Widely available tabular matrices used in RNA-seq data analysis, including gene expression data (assay), gene annotation, and sample annotation are shown. The `prepareSeObj()` function in RUVprps can construct a *SummarizedExperiment* object from these matrices. All outputs generated by RUVprps functions can be stored in the metadata of the*SummarizedExperiment* object. b) Overview of the gene-level and global statistical and computational tools used during the variation and normalization-assessment steps. RUVprps provides two classes of tools designed to evaluate variation associated with either categorical or continuous variables. c) Schematic representation of the major processing steps in RUV-III normalization. RUV-III can be applied across scenarios regardless of whether the biological or unwanted sources of variation are known or unknown. d) RUVprps produces a large number of graphical outputs created by the RUVprps variation and normalization performance-assessment modules for user investigation. e) RUVprps numerically summarizes all performance-assessment metrics to facilitate selection of the most suitable normalized dataset for downstream analysis.

Figure 1c shows that the RUV-III normalization framework across different scenarios, ranging from fully supervised to fully unsupervised applications of the RUV-III method. Fully supervised refers to situations where sources of both biological and unwanted variation are known. RUVprps introduces novel features that enable application of the method in settings where both biological and unwanted variation are completely unknown, which we refer to as fully unsupervised. Finally, RUVprps generates a comprehensive set of diagnostic plots and numerical summaries to assist users in identifying the most suitable normalized data for downstream analyses (Fig. 1d and e).

The RUV-III method has also been applied to other omics data, including single-cell RNA-seq [9, 11], single-cell CyTOF [8], and bulk proteomics [7] and metabolomics analyses. Currently, RUV-III generates log-normalized data as output. This data can be utilized for various downstream analyses, such as unsupervised clustering, prediction, gene set enrichment analysis and more. The estimated unwanted variation terms from RUV-III can serve as covariates in models within edgeR [12] and DESeq2 [13] designed for raw count data.

### Overview of the RUVprps R package

The underlying data structure in RUVprps is based on the *SummarizedExperiment* object (Fig. 1a). The object integrates experimental data termed an “assay” (such as gene expression values from RNA-seq), sample and gene feature information, along with metadata, into a single container [14]. Further, this enables the numerical and graphical outputs of individual RUVprps functions to be stored within the metadata of the object, allowing them to be readily accessed and utilized in subsequent analysis steps. RUVprps leverages the features of the *SummarizedExperiment* object to streamline the normalization process and minimize potential user-specific programming errors. RUVprps facilitates the creation of a *SummarizedExperiment* object from tabular data through the `prepareSeObj()` function (Methods). The Supplementary File provides guidance on accessing the various data types in a *SummarizedExperiment* object. We will subsequently detail the structure of the metadata type and show how and where various outputs of the RUVprps functions are stored.

The RUVprps package includes over 100 functions designed to address the full spectrum of tasks involved in end-to-end RNA-seq normalization (Fig. 2). Individual functions can be used separately; however, these functions are integrated in various ways to form a few main functions that streamline the core steps of the analysis (Fig. 1). The first step involves a comprehensive assessment of variation in both categorical and continuous variables (Fig. 2c). The second step is the application of RUV-III normalization, which includes: (1) selecting and evaluating the performance of NCG, (2) generating PR or PRPS sets, unless suitable TR are already available, (3) applying RUV-III with varying parameters, particularly different values of *k*, and performing a normalization assessment pipeline that evaluates the performance of RNA-seq normalization methods. All numerical and graphical outputs of individual steps are by default stored in the *SummarizedExperiment* object, which will be used in subsequent steps.

**Figure 2:**
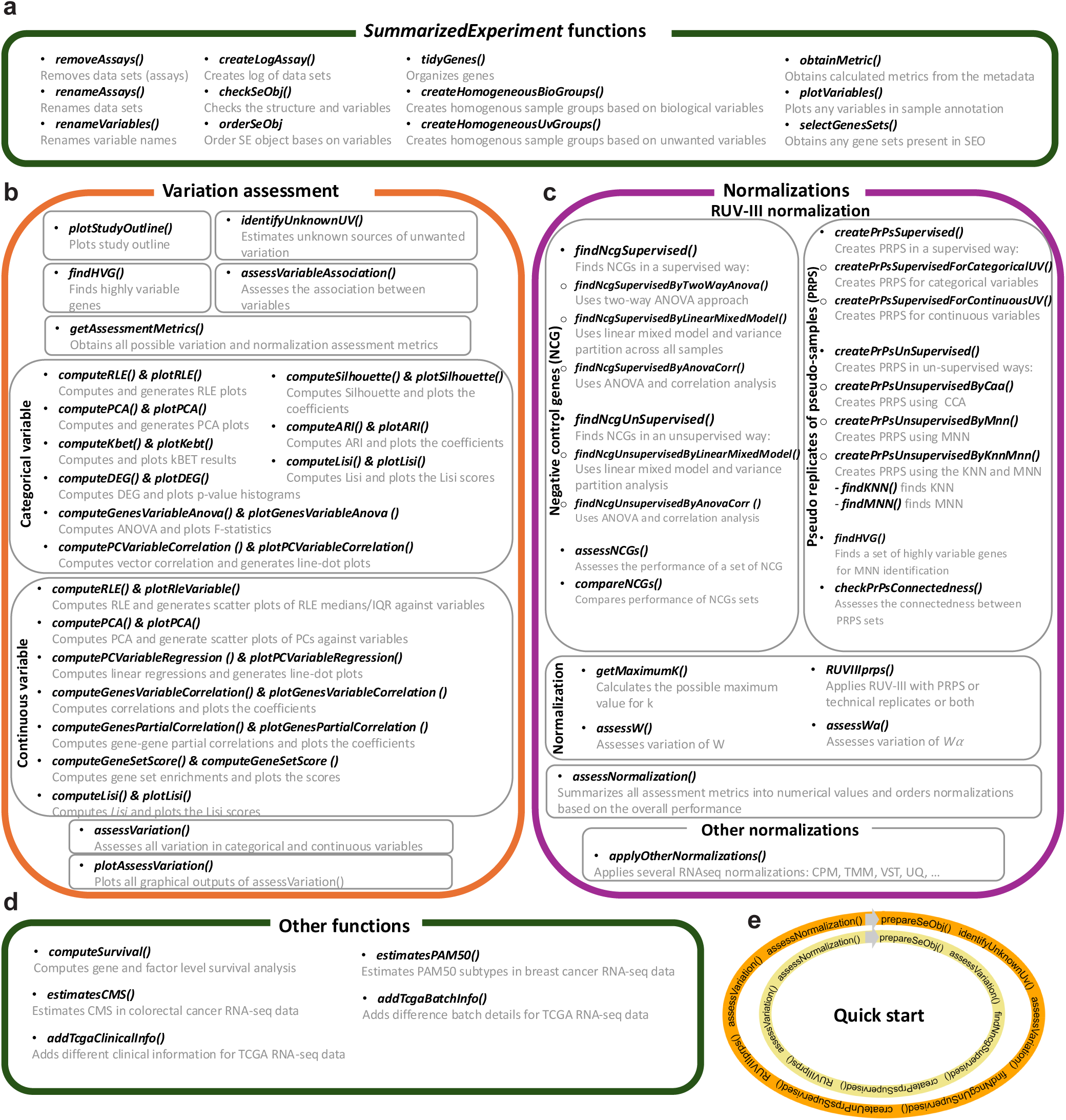
Overview of the functions and their applications in RUVprps. a)Displays functions that facilitate working with data and variables in a *SummarizedExperiment* object. b)Exhibits all the functions that are involved in the variation assessment step. Two different class of functions and approaches are proposed to assess categorical or continuous variables. All these functions can be run simultaneously just by using the `assessVariation()` function. c)Displays different functions that are involved in different steps of the RUV-III normalization. The steps are the identification and assessment of NCG, PRPS construction, and the application of RUV-III. The first two major steps can be performed in supervised or unsupervised manners. Further, the `assessNormalization()` function numerically summarizes all the results of the functions involved in the variation assessment step and orders normalizations based on their performance. RUVprps offers the `applyOtherNormalization(`) to apply widely-used normalizations for RNA-seq data. d) Shows functions that facilitate working with different types of data, with a strong focus on the TCGA RNA-seq data. e) Shows only the required functions for fully supervised (yellow ellipse) and fully unsupervised applications (orange ellipse) of RUV-III for RNA-seq normalization.

RUVprps provides a set of general-purpose functions (Fig. 2a and d) designed to streamline working with *SummarizedExperiment* objects and to facilitate the processing of the Cancer Genome Atlas (TCGA) RNA-seq data. Detailed descriptions of all individual functions are provided in the Supplementary File.

RUVprps leverages data vectorization and parallelization to maximize speed and improve the scalability of each function. Further, running the entire function each time parameters are adjusted can be computationally expensive. To address this, the major functions include an option that allows users to re-run only the necessary steps affected by the parameter changes. Outputs that need to be generated only once are stored in intermediate files and made accessible through the *SummarizedExperiment* object. When this option is enabled, the function retrieves and reuses these stored results, significantly improving efficiency during iterative tuning.

### Assessment of variation in the data

A thorough assessment of both biological and unwanted variation at the gene-level and globally, conducted before and after normalization, is essential for robust RNA-seq data analysis [3, 5]. Our analysis of the Recount3 database (Methods) including 220,000 samples from 2,284 RNA-seq studies, demonstrates the importance of assessing variation at both levels to reveal and quantify individual sources of unwanted variation (Supplementary File). Across this large collection of RNA-seq datasets, the results show that the commonly used principal component analysis (PCA) and interpretation of the first few principal components (PCs) do not fully indicate how effectively library size effects are removed, even after counts-per-million (CPM) normalization (Supplementary Fig. 1a). In contrast, gene-level correlations between library size and individual gene expression provide a strong and complementary approach to PCA-based assessments. Our analysis shows that a reasonable proportion of genes are significantly correlated with library size, whereas the first three PCs are not highly associated with library size. Similar analyses were performed to evaluate the performance of both the gene-level and global assessments for batch effects (Supplementary Fig. 1b). Similar to library size, gene-level ANOVA between individual gene expression and batch effects reveals a strong impact of batch effects, whereas the first three PC show little or no association with batch effects in a large proportion of datasets. Further, we will show the importance of gene-level and global assessments when normalizing different RNA-seq datasets.

RUVprps offers eight functions for evaluating categorical variables and seven for continuous variables. These are all integrated into a major function, `assessVariation()`, which streamlines the workflow while still allowing each function to be used independently for more targeted analyses (Fig. 2b). For individual categorical variables e.g. batches, the function calculates relative log expression (RLE), PCA, vector correlation analysis (VCA), adjusted Rand index (ARI), silhouette coefficient, local inverse Simpson’s index (LISI), k-nearest neighbor batch effect test (kBET), gene level ANOVA, and differential gene expression analysis. For each continuous variable e.g. library size, the function computes gene-level correlation, gene-gene partial correlation, correlation with RLE medians and IQRs, correlation with principal components, local inverse Simpson’s index (LISI) and and linear regression analysis (LRA). We refer to the Supplementary File for details of individual functions and Methods to show how to interpret individual metrics.

### Identification of unknown sources of unwanted variation

Accurate identification of sources of unwanted variation is essential for evaluating normalization performance. Further, this step is critical for constructing PRPS data and for assessing and identifying NCGs for RUV-III with PRPS normalization. There may be instances in which certain sources of unwanted variation remain unrecorded, or where full details on experimental conditions and sample processing is unavailable, for example when analyzing publicly available data [3, 5]. We refer to these as “unknown” sources of unwanted variation.

RUVprps provides a function called `identifyUnknownUV()` that can help estimate possible sources of unknown unwanted variation in RNA-seq data (Fig. 3a). This function offers three different approaches including `rle`, `pca` and `sample.scoring` with various strategies (Methods).

**Figure 3:**
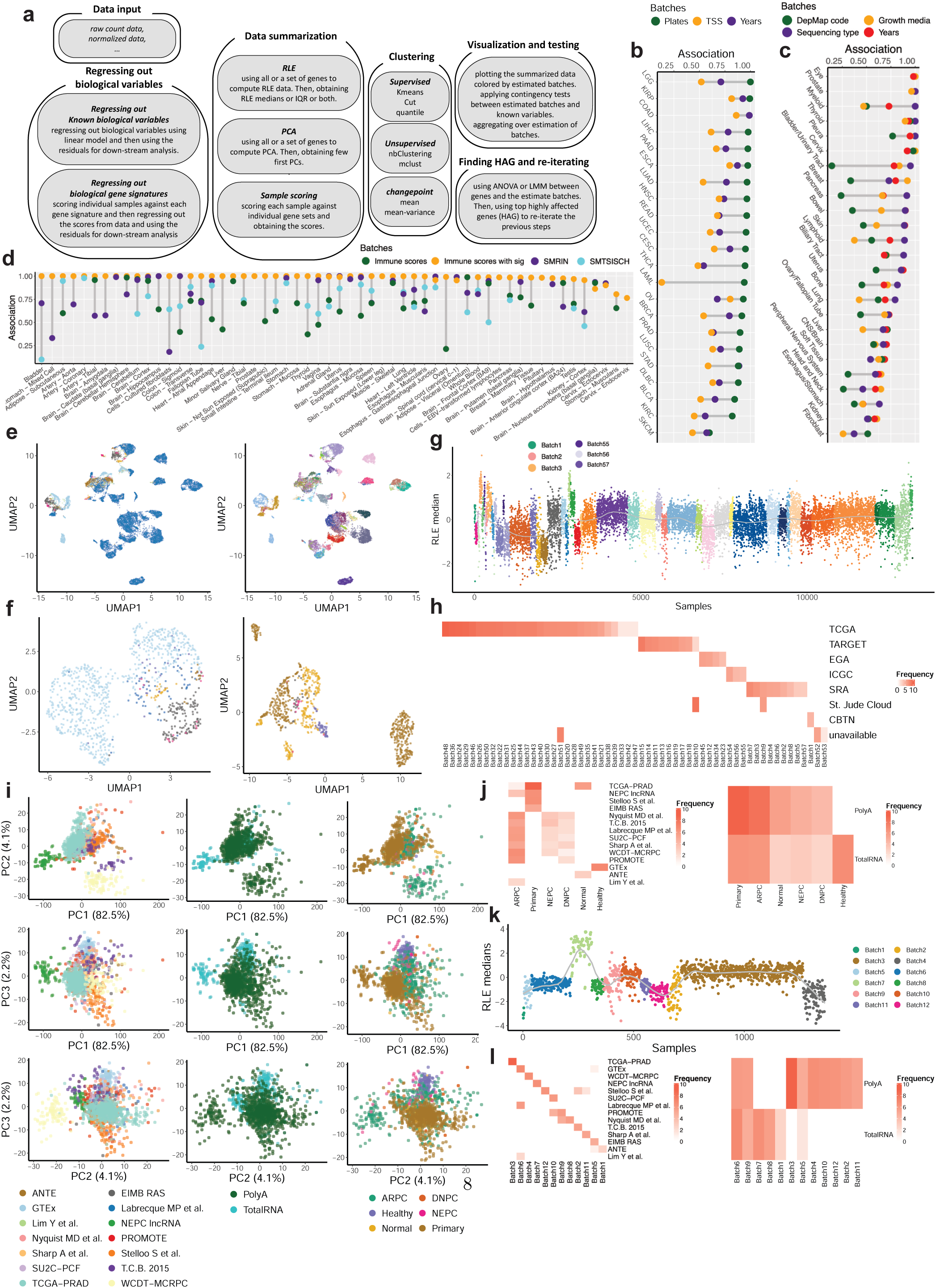
Identification of unknown sources of unwanted variation in RNA-seq data by RUVprps. a) Schematic illustrating identification of unknown sources of unwanted variation using the `identifyUnknownUV()` function (Methods). Briefly, gene expression data are summarized using relative log expression (RLE), principal component analysis (PCA), or sample-scoring approaches, followed by clustering of the summarized data to estimate sources of unwanted variation. b–d) Plots showing associations between unwanted variation estimated by RUVprps and known sources of unwanted variation in TCGA pan-cancer (b), CCLE (c), and GTEx (d) RNA-seq datasets, respectively. e) Scatter plot of the first two UMAP dimensions of the Treehouse raw count (*log*_2_ + 1) data, colored by study(8 studies) (left) and cancer type (102 types) (right). f) Scatter plot of the first two UMAP dimensions for glioma and acute lymphoblastic leukemia samples, colored by studies. Legends for panels e and f are omitted due to the large number of factors. g) RLE medians of the Treehouse raw count (*log*_2_ + 1) RNA-seq data, colored by estimated batches identified by RUVprps. h) Confusion matrix comparing estimated batches with projects or studies in the Treehouse cohort. i) Scatter plots of the first three principal components of the PCCAT raw count (*log*_2_ + 1) data, colored by major molecular subtype (left), library preparation protocol (middle), and study (right).j) Confusion matrices between major molecular subtype and study (left) and library preparation protocol (right). k) RLE medians of the raw count (*log*_2_ + 1) PCCAT RNA-seq data, colored by estimated batches identified by RUVprps. l) Confusion matrices comparing RUVprps-estimated batches with study (left) and library preparation protocol (right).

Each approach has distinct advantages and disadvantages, depending on the impact of unwanted variation on the data. It should be noted that effective identification of all unknown sources of unwanted variation may require the use of multiple approaches with iterations and exploratory data analysis, specifically, in situations where all biological variation are unknown.

We used several large-scale RNA-seq datasets, including the TCGA pan-cancer cohort, the Cancer Cell Line Encyclopedia (CCLE), the Genotype-Tissue Expression (GTEx)[15], Treehouse [16], and the Prostate Cancer Cell Atlas (PCCAT)[17] (Methods) to evaluate the performance of RUVprps in identifying sources of unwanted variation. In these analyses, sources of unwanted variation were estimated using RUVprps, and their associations with known sources of unwanted variation were subsequently examined. We refer to the Supplementary File for details on how RUVprps was applied to estimate unwanted variation within individual studies.

Figure 3b, c and d demonstrate the strong performance of RUVprps in estimating unwanted variation within the TCGA, CCLE and GTEx RNA-seq data. In this analysis, sources of unwanted variation were estimated within each study or tissue, and their associations with known sources of unwanted variation were evaluated. It should be noted that for certain studies or tissues, where the associations between estimated and known sources of unwanted variation are not strong, additional uncharacterized sources of variation may exist that are not publicly available.

The Treehouse cohort comprises a unified collection of cancer RNA-seq data assembled from three large-scale projects, with sequencing reads from all samples processed using a uniform pipeline. UMAP embeddings of both the full dataset and tissue-specific subsets reveal pronounced batch effects between studies (Fig. 3e and f). Estimates of unwanted variation inferred by RUVprps demonstrate that the method effectively captures batch effects both within and between studies in the Treehouse RNA-seq dataset (Fig. 3g and h).

Identifying unwanted variation in the PCCAT RNA-seq dataset is particularly challenging because of strong confounding between sources of unwanted variation (e.g., study) and major biological subtypes (Fig. 3i and j). However, the RLE-based approach robustly identifies sources of unwanted variation both within and across studies in the PCCAT RNA-seq dataset(Fig. 3k and l). We will show that a combined `rle` and `pca` approaches further reveal more unwanted variation e.g., the years effect in the TCGA data, within studies in the PCCAT dataset. Finally, we applied RUVprps across the entire Recount3 database and the results show that all studies exhibited at least two distinct groups, most likely reflecting batch effects, in the distribution of RLE medians of the CPM-normalized data (Supplementary Fig. 2).

### The procedure of RUV-III normalization

Figure 2c exhibits functions that are designed to apply RUV-III in either supervised or unsupervised manner. To implement RUV-III effectively, it is essential to use true NCG and control samples that capture variation from individual sources of unwanted variation. The NCG may be pre-selected; however, if they fail to capture all sources of unwanted variation, they should be identified from the data. Control samples can be TR; however, if TRs are not well distributed across batches or are unavailable, PR or PRPS should be created. It should be emphasized that RUVprps can accommodate different types of control samples for RUV-III normalization. It provides both supervised and unsupervised approaches to find true NCG and creating PR or PRPS. In the supervised approach, sources of both biological and unwanted variation must be known, while unsupervised methods do not require such prior information. RUVprps implements a novel and faster approach to RUV-III normalization, enabling efficient assessment of various method parameters, particularly for determining the optimal value of *k* in the normalization process. For each *k* value specified in the optimization step, RUVprps will store the corresponding RUVIII adjusted data and estimated unwanted variation *W* matrices as different assays within the same *SummarizedExperiment* object. The different steps involved in the RUV-III normalization procedure will be explained in the subsequent section.

### Supervised and unsupervised selection of NCG

RUVprps offers both supervised and unsupervised approaches for deriving suitable sets of NCG from the data. There are five major steps involved in selecting NCG in both approaches (Methods). RUVprps provides three functions, each offering a distinct approach for finding NCG in a supervised manner (Methods). These functions are combined into a major function called `findNcgSupervised()` to streamline the analysis. Our general recommendation is to apply all the three functions and compare their performance using the `compareNCGs()` function.

In the unsupervised selection of NCG, both biological and unwanted sources of variation may be unknown. RUVprps provides two approaches and combines them into a major function called `findNcgUnSupervised()` to identify NCG in an unsupervised manner (Methods).

We initially used the TCGA (cancer samples) and GTEx (healthy samples) RNA-seq datasets to assess the performance of unsupervised NCG selection. In this analysis, sources of unwanted variation were estimated in an unsupervised manner, and a set of NCGs was identified for each dataset (Supplementary File). For each dataset, PCA was performed using only the selected NCGs, and the association of the first few PCs with the estimated batches was examined. This analysis was repeated for three sets of well-known housekeeping genes. As shown in Figure 4a-c, NCGs selected in an unsupervised manner by RUVprps perform similarly to, or slightly better than, the housekeeping gene sets in both TCGA and GTEx RNA-seq datasets. For the TCGA RNA-seq studies, we conducted further analyses to verify that RUVprps selected NCGs do not include genes that are highly variable with respect to biological variation. Highly variable genes (HVGs) were first identified (Methods), and the overlap between selected NCGs and HVGs was assessed. Figure 4b shows no significant overlap between the two sets. We also evaluated the overlap between the NCGs and pan-cancer stable genes from the singscore R package, and observed substantial concordance across TCGA studies. We refer to the Supplementary File for more details. We will demonstrate the performance of unsupervised NCG selection in RUV-III normalization across several RNA-seq datasets in subsequent sections.

**Figure 4:**
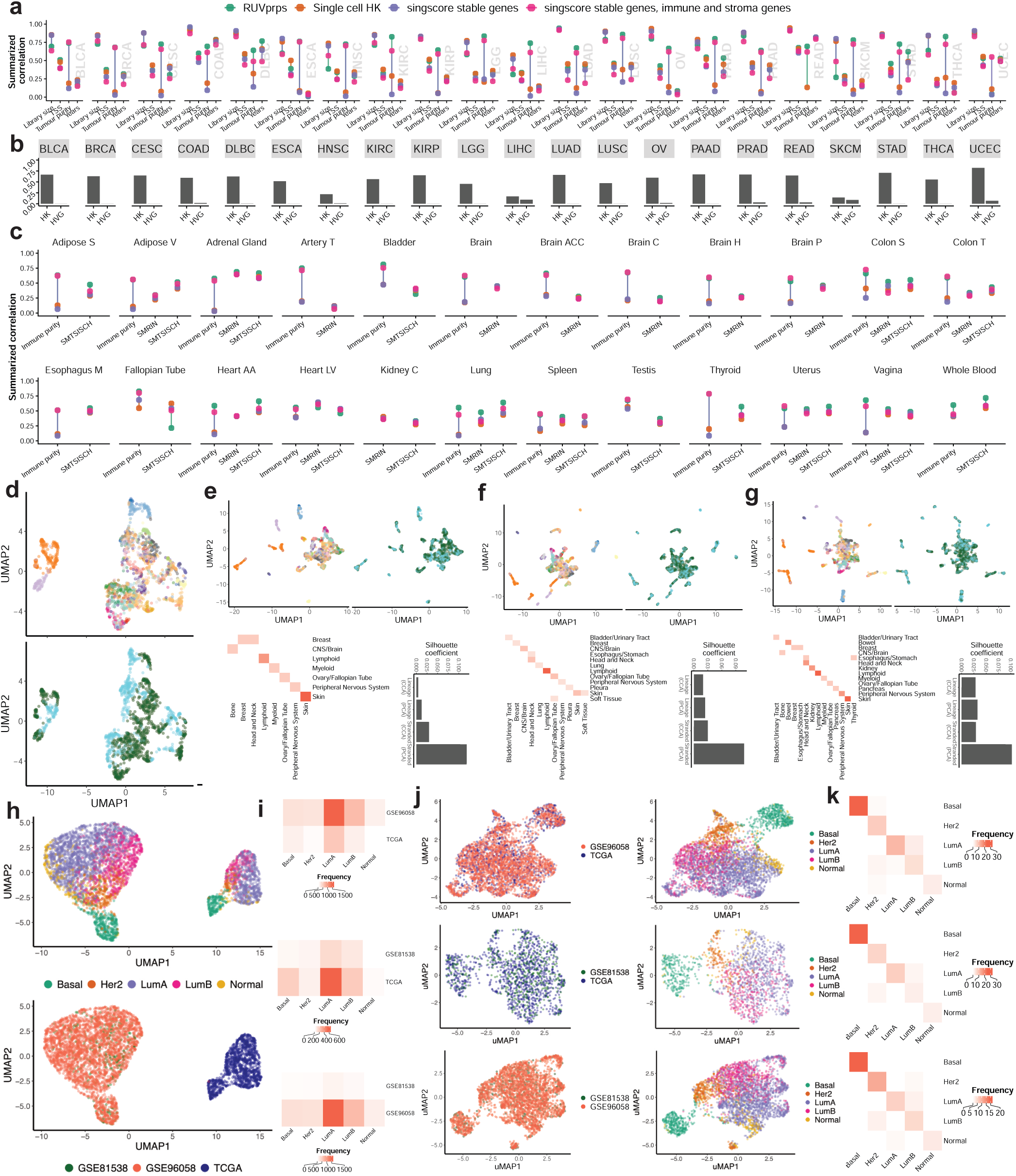
Performance of RUVprps for unsupervised NCG selection and PRPS construction. a) Performance of NCGs selected by RUVprps compared with three sets of housekeeping genes in capturing different sources of unwanted variation in TCGA RNA-seq studies. b) Bar plots showing the proportion of overlap between NCGs selected by RUVprps and housekeeping genes or highly variable genes for individual TCGA studies. c) Same analysis as in a for the GTEx RNA-seq datasets. d) UMAP projections of CCLE RNA-seq raw count data (*log*_2_ + 1), colored by cell line identity (top) and RNA-seq strandedness (bottom). e) UMAP projections of the first 20 canonical correlation analysis (CCA) components, colored by cell line identity (left) and strandedness (right). The confusion matrix below the UMAP projections show the number of samples matched across the stranded and unstranded batches by CCA. Silhouette coefficients quantify cluster separation by cell line and strandedness in principal components and CCA components. f,g) The same as e, using the first 30 (f) and 40 (g) CCA components, respectively. h) UMAP projections of FPKM-normalized expression data (*log*_2_ + 1) from three large breast cancer RNA-seq studies, colored by PAM50 subtype (left) and study (right). i) Distribution of PAM50 subtypes across each pair of studies. j) For each study pair, UMAP projections of the first 30 CCA components, colored by study (left) and PAM50 subtype (right). k) Confusion matrix showing PAM50 subtype concordance across studies identified by CCA.

### Supervised and unsupervised selection of PRPS

The PRPS can play the role of TR in the RUV-III method [3, 5] when there are no, too few or only poorly placed genuine TR. RUVprps provides both supervised and unsupervised approaches to create PRPS data (Methods). It is important to highlight that all sources of unwanted variation must be categorical for PRPS construction. If any are continuous, they will be grouped into categorical bins as specified by users. For example, library size can be stratified into three groups corresponding to low, intermediate and high library sizes, and PRPS can be generated either across all groups or just between the low and high groups. All RUVprps functions designed for generating PRPS include options to assess PRPS contamination and connectivity (Methods).

In the unsupervised case, identifying biologically homogeneous samples across different batches for PRPS construction is substantially challenging. This difficulty arises because the data are influenced by multiple sources of unwanted variation, making it difficult to distinguish truly biologically homogeneous samples across batches. To overcome this challenge, RUVprps employs canonical correlation analysis (CCA), *k* and mutual nearest-neighbor approaches with different strategies to create PRPS in an unsupervised way. It provides three distinct functions, which are integrated into a main function, `createUnSupervisedPrPs()` (Methods). In general, each function does the following: (1) identifies samples with similar gene expression profiles across known or estimated sources of unwanted variation; (2) generates pseudo-samples (PS) by averaging *n* samples (at least *n* ≥ 3 by default) within batches; and (3) matches PS samples to construct distinct PR sets.

We first assessed the performance of the CCA approach for identifying homogeneous samples across different batches. These samples were used to construct PRPS. As shown in Figure 4d, the CCLE RNA-seq data are strongly affected by RNA-seq library strandedness, forming distinct stranded and unstranded cohorts. To address this, CCA was applied to identify homogeneous samples, ideally corresponding to the same cell lines, across the stranded and unstranded cohorts. Figure 4e-g shows UMAP embeddings based on the first 20, 30, and 40 canonical components, demonstrating that the dominant effects of cell line are no longer visible, while the underlying biological differences are largely preserved. The resulting canonical components were then used to identify MNNs across batches for PRPS construction. Confusion matrices of matched samples across batches demonstrate the strong performance of the CCA-based approach in identifying homogeneous samples when varying the number of canonical components used as input (Fig. 4e).

We next applied the CCA-based approach to three large breast cancer RNA-seq studies (Methods). As shown in Figure 4h, substantial batch effects are evident between TCGA and non-TCGA datasets. CCA was performed between all pairwise combinations of studies. UMAP projections of the first 30 canonical components show that study-specific effects are effectively removed, while the expected clustering by the PAM50 subtype is clearly preserved (Fig. 4h). Application of the MNN using the first 30 canonical components successfully identified homogeneous PAM50 samples across all pairwise study comparisons. We will show the the performance of CCA-based PRPS construction in RUV-III normalization of several RNA-seq data.

### RUV-III normalization

RUVprps provides a core function, `RUVIIIprps()`, for implementing the RUV-III normalization method. This function supports all types of control samples, including TR, PR, PRPS, or any combination thereof. The construction of the mapping matrix for different control samples (the M matrix) is described in the Methods and Supplementary File. The PRPS sets can be a combination of both supervised and unsupervised approaches. Similarly, the NCG can be identified using supervised, unsupervised, or pre-selected gene sets, or any combination of these. We refer to Supplementary File for details on the `RUVIIIprps()` function.

To apply RUV-III with the chosen control samples and NCG set, one needs to specify the number *k* of (linear) dimensions of unwanted variation to estimate and remove from the data (Methods). As with the choice of control samples and NCG, there are no clear rules to determine the optimal value of *k*. Our general approach is to repeat the analysis with different values of *k*, and then evaluate the quality of each analysis using various statistical metrics and prior biological knowledge, as explained in the above. Further, RUVprps provides the function `assessW()` to evaluate the association of the W matrix with both biological and unwanted variation, which can help identify an appropriate value for *K* (Methods).

### Data types required for RUVprps

There are specific data type requirements for various steps of the RUVprps package. In general, we highly recommend using raw count data without any prior transformation as input, if available. By default, the `RUVprps` functions apply a log_2_-based transformation with a pseudo-count. We refer to Supplementary File for more details. While RUVprps can accommodate transformed or initially normalized data types such as FPKM or CPM, it is important to note that certain transformations may remove biological signals from the data [3]. If such signals are lost during preprocessing, RUVprps may not be able to recover them during normalization.

### Other RNA-seq normalizations

RUVprps provides a wrapper function, `applyOtherNormalizations()`, which implements several widely used RNA-seq normalization methods, including counts per million (CPM) [12], trimmed mean of M-values (TMM) [18], variance-stabilizing transformation (VST) [13], upper-quartile normalization, full quantile normalization, and median-scaling normalization [19]. Each normalized dataset is stored as a separate assay within a *SummarizedExperiment* object, enabling systematic evaluation of normalization performance. More generally, any RNA-seq normalization method can be incorporated, with normalized data added as additional assays to the *SummarizedExperiment* for normalizations assessment provided by RUVprps.

### Normalization performance assessments

It is essential to assess the performance of a normalization as a remover of unwanted variation and a preserver of biological variation in the data. An ideal normalization should preserve and/or enhance all known sources of biological variation, and remove or reduce the impact of all known or estimated (unknown) sources of unwanted variation. RUVprps offers the `assessNormalization()` function to address those aspects on both the biological and unwanted variation. This function first uses the `assessVariation()` function to compute all the corresponding metrics of assessment and then uses different approaches to numerically summarize them. Finally, all evaluation metrics are aggregated into an overall score to rank the methods according to their overall performance. Individual scores range from 0 to 1, and are subsequently transformed so that higher scores consistently correspond to better performance, regardless of whether they evaluate biological or unwanted variation. We refer to the Supplementary File for details on the numerical summarization of the different assessment metrics.

### Examples of RUVprps normalization performance

To demonstrate the ability of RUVprps to normalize RNA-seq data, we showcase its performance on diverse RNA-seq datasets with various challenges. For individual examples, full details of the RUV-III normalization process are provided in the Supplementary File.

### The TCGA BRCA RNA-seq data

In this example, we evaluated the performance of RUVprps in removing library size effects, tumor purity variation, and batch effects from TCGA breast cancer invasive carcinoma (BRCA) RNA-seq data. We defined four scenarios spanning fully supervised to fully unsupervised applications of RUVprps. Here, we present results from the fully unsupervised application and refer to the Supplementary File for details of the other scenarios. The Supplementary File also provides detailed descriptions of data preprocessing, estimation of PAM50 subtypes, unsupervised estimation of unwanted variation, RUV-III normalization, and assessment of sources of variation.

Fig. 5a and b show that NCG selection using a linear mixed model performs slightly better than the other unsupervised approach implemented in RUVprps. Further, the selected NCG exhibit substantial overlap with a pan-cancer stable genes obtained from the singscore R package as well as immune and stromal gene sets from [20], while showing limited overlap with highly variable genes (HVGs), due to biological variation, identified in the data (Fig. 5c). Different sets of PRPS were generated to capture variation in library size, tumor purity, and estimated batch effects (Fig. 5d). Library size and tumor purity were each divided into four groups, and PRPS sets were constructed between the low and high groups. Fig. 5e demonstrates the strong performance of the unsupervised CCA-based PRPS approach in identifying biologically similar samples across batches. RUV-III was then applied using different values of *K*. The association between the RUV-III W matrix and the specified biological and unwanted variables indicated that the unsupervised approach effectively captures all sources of unwanted variation, while it did not strongly capture major biological variation. The numerical normalization performance assessment showed that RUV-III-PRPS with *k* = 20 effectively removed unwanted variation while preserving variation in known biological factors (Fig. 5g). Scatter plots of the first three PC demonstrated clear mixing of the flow cell chemistry batches, while preserving the biologically expected separation of the PAM50 subtypes in the RUV-III–normalized data (Fig. 5h). Kaplan–Meier survival analysis revealed that PAM50 subtypes estimated from RUV-III–normalized data were significantly associated with patient-level survival data (Fig. 5i).

**Figure 5:**
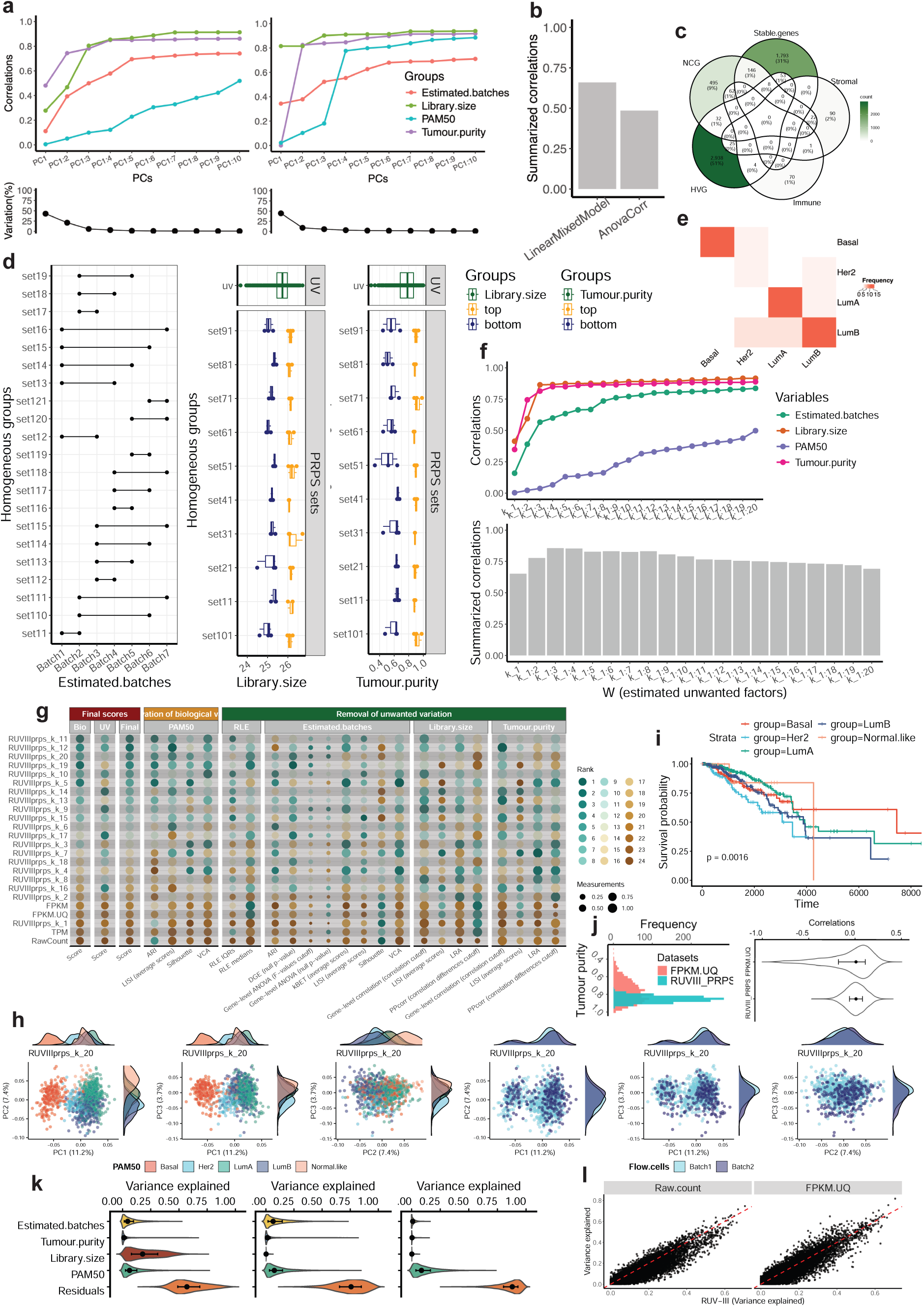
Unsupervised RUV-III-PRPS normalization of TCGA BRCA RNA-seq data. a) Line–dot plots show the performance of selected NCGs using different unsupervised approaches, including `LinearMixedModel` (left) and `AnovaCorr` (right). b) Barplot summarizing the performance of the selected NCG sets; higher correlations indicate better performance. c) Overlap of the selected NCGs with a pan-cancer stable gene set obtained from singscore R package, stromal and immune gene sets from [20], and a set of highly variable genes (HVGs) identified by the `findHVGs` function of RUVprps in the data. d) PRPS maps for estimated batch effects (left), library size (middle), and tumor purity (right). The y-axis displays biologically homogeneous sample groups, and the x-axes show the distributions of samples across estimated batches, library size, and tumor purity, respectively. In the batch PRPS map, each horizontal line connecting two points defines a PRPS set across batches. In the library size PRPS map, each horizontal line corresponds to six samples representing the highest and lowest library sizes within a homogeneous group; every three samples are averaged to create a pseudo-sample (PS). In the tumor purity PRPS map, each horizontal line corresponds to six samples representing the highest and lowest tumor purity values within a homogeneous group; every three samples are averaged to create a PS. e) Confusion matrix shows the PAM50 subtypes selected by the unsupervised CCA-based PRPS approach across all possible pairs of batches. f) Line–dot plot shows associations between columns of the estimated W matrix and both biological and unwanted variables, with the accompanying barplot summarizing these correlations. g) Numerical assessments of all metrics for biological and unwanted variables. Each point represents the performance of a normalization method, with higher values indicating better performance. Normalized datasets are ranked according to their effectiveness in removing unwanted variation while preserving biological signal. h) Scatter plots of the first three PCs, colored by PAM50 subtype and the flow cell chemistries, for RUV-III with *K* = 20. i) Kaplan–Meier survival analysis of PAM50 subtypes estimated from RUV-III–normalized data. j The histogram shows the tumor purity estimates from the TCGA FPKM.UQ data and RUV-III–PRPS normalization with *K* = 20. The violin plots displays gene-level correlation with tumor purity for the FPKM.UQ and RUV-III normalized data. k) Results of gene-level variance decomposition using a linear mixed model for TCGA BRCA raw counts (left), FPKM-UQ (middle), and RUV-III–PRPS–normalized data (right). The violin plots show the total variance of individual genes across specified variables. l)Shows the results of a linear mixed model analysis of gene-level variation associated with PAM50 subtypes across different normalization methods.

Fig. 5j illustrates that gene expression variation introduced as a result of the different values of tumor purity was substantially reduced by RUV-III-PRPS normalization. Finally, linear mixed model analyses performed on raw counts (log_2_ +1), FPKM-UQ, and RUV-III-PRPS–normalized data show that RUV-III-PRPS effectively removed multiple sources of unwanted variation while preserving variation associated with known biological factors (Fig. 5k and l).

### The TCGA READ RNA-seq data

We used the TCGA rectal adenocarcinoma (READ) RNA-seq data to demonstrate the removal of substantial batch and library size effects while preserving variation associated with tumor purity and other known biological factors. Analogous to the TCGA BRCA example, we considered four scenarios ranging from fully supervised to fully unsupervised applications of RUVprps. Here, we present results from the fully unsupervised scenario and refer to the Supplementary File for details of the remaining scenarios.

The first three PCs of the TCGA READ raw data (log_2_ +1) clearly show the impact of major time interval effects. In contrast, RUV-III-PRPS normalization effectively removed this unwanted variation and preserved the separation of the consensus molecular subtypes (CMS) in the data (Fig. 6a,b). These results were further supported by multiple complementary assessment metrics (Fig. 6c–h). Both gene-level and global analyses indicated that variation associated with library size was successfully removed, while biologically meaningful variation related to tumor purity was preserved following RUV-III-PRPS normalization (Fig. 6i–m). Moreover, a broad range of downstream assessments demonstrated that variation associated with CMS was retained in the RUV-III-PRPS–normalized data (Fig. 6n–q). Consistent with this observation, Kaplan–Meier survival analyses based on CMS revealed that RUV-III-PRPS normalization improved the expected association between CMS and patient survival (Fig. 6r). Detailed results for all variation assessment metrics across datasets are provided in the Supplementary File.

**Figure 6:**
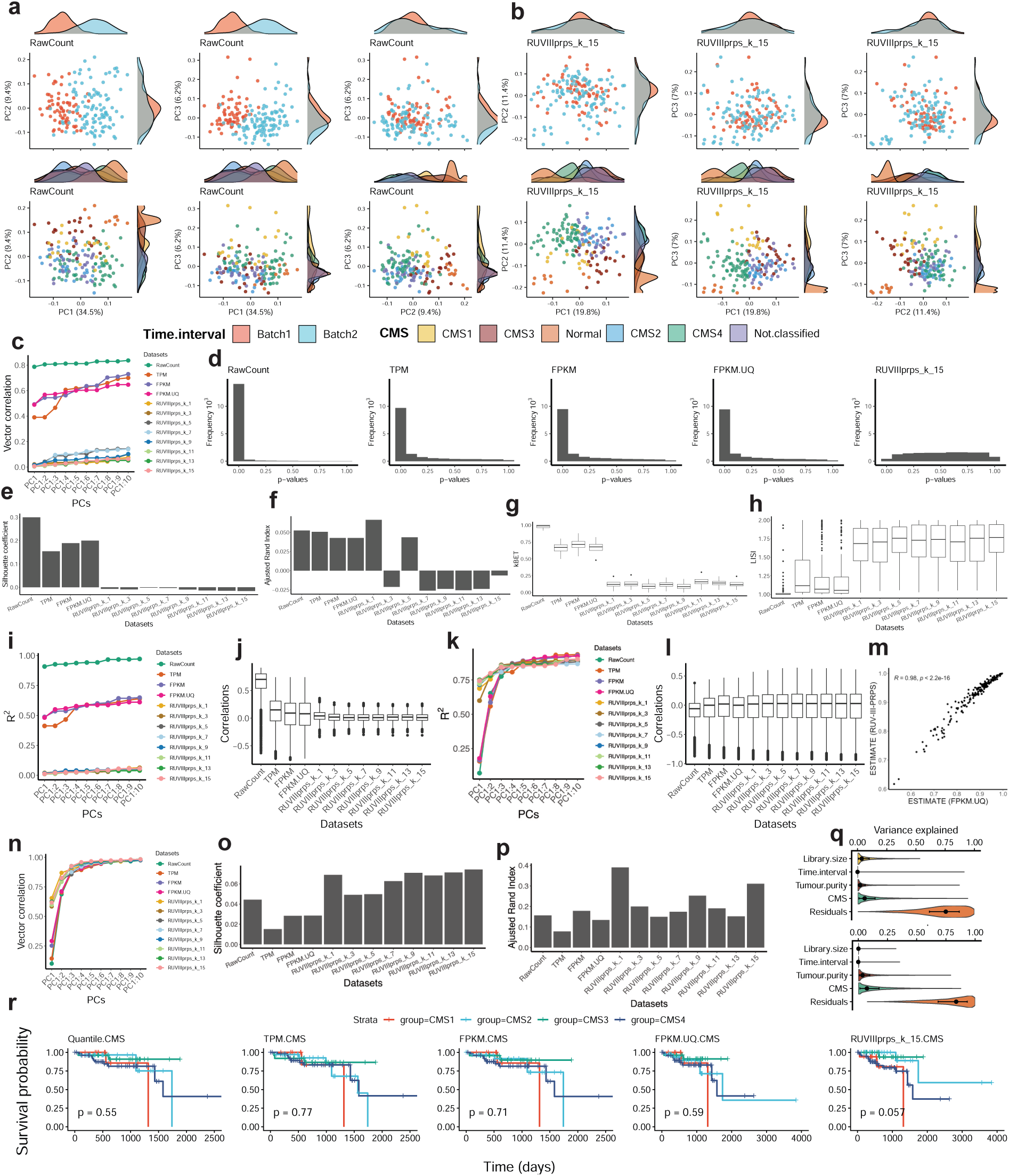
Supervised RUV-III normalization of TCGA READ RNA-seq data. a) Scatter plots of the first three PCs of raw count (log_2_ +1) TCGA READ RNA-seq data, colored by major time interval (top row) and consensus molecular subtype (CMS; bottom row). b) As in a, but for data normalized using RUV-III-PRPS. The PCA was performed using 500 genes proposed by the CMScaller R package for CMS identification. c) Vector correlations between the first ten PCs (cumulatively) and major time interval across datasets. d) Histograms of p-values from differential gene expression analyses comparing major time intervals across datasets. e–h) Silhouette coefficient (e), adjusted Rand index (f), kBET acceptance rate (g), and LISI scores (h), respectively, evaluating sample mixing by major time interval for different datasets. i) Linear regression analysis assessing the association between the first ten PCs (cumulatively) and library size across multiple datasets. j) Boxplots showing correlation coefficients between individual gene expression levels and library size for each dataset. k,l) As in i,j, respectively, but assessing associations with tumor purity. m) Correlation between estimated tumor purity from TCGA FPKM-UQ and RUV-III-PRPS–normalized data. n) Vector correlations between the first ten PCs (cumulatively) and CMS. o and p) Silhouette coefficient and adjusted Rand index assessing separation of CMS subtypes. q) Results of linear mixed-effects modeling for TCGA FPKM-UQ (top) and RUV-III-PRPS–normalized data (bottom). r) Kaplan–Meier survival analyses stratified by CMS across datasets

### Three large breast cancer RNA-seq data

Here, our goal is to show the performance of fully supervised and unsupervised RUVprps to normalize multiple RNA-seq studies. We used three large breast cancer RNA-seq studies from TCGA and two studies by Brueffer et al. [21, 22]. Our aim is to remove unwanted variation within and between studies, while preserving known biological variation, including the PAM50 subtypes and tumor purity. We refer to the Supplementary File for full details on data preprocessing and RUVprps normalization procedures. We defined two scenarios, one is fully supervised and the other one is fully unsupervised.

The first three PCs reveal substantial study effects between the TCGA and non-TCGA cohorts in the FPKM data (Fig. 7a). This observation is supported by the RLE plot of the FPKM data (Fig. 7b). Figure 7c shows that variation in tumor purity and PAM50 subtypes is not strongly associated with the studies. For the unsupervised application of RUVprps, sources of unwanted variation were estimated using the *rle* approach. Figure 7a and b illustrate that RUVprps identifies both within-study and between-study batch effects.

**Figure 7:**
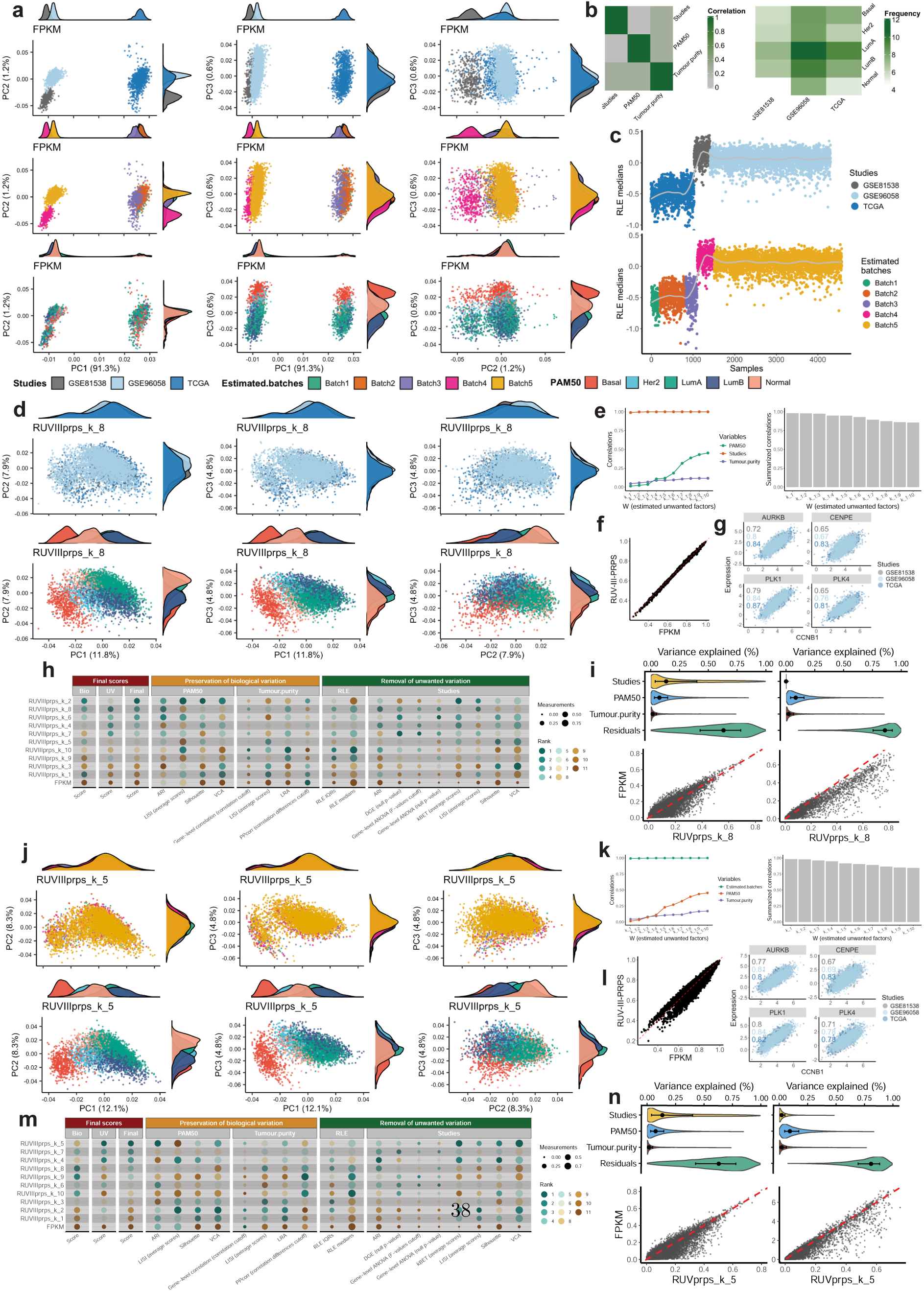
Supervised and unsupervised RUV-III normalization of three large breast cancer RNA-seq datasets. a) Scatter plots of the first three PCs of the FPKM data, colored by the studies (top row), estimated unwanted variation inferred by RUVprps (middle row), and the PAM50 molecular subtypes (bottom row). b) Left: heatmap showing pairwise correlations between PAM50 subtype, tumor purity, and studies. Right: distribution of PAM50 subtypes across studies. c) Relative log expression (RLE) plots showing sample-wise medians of the FPKM data, colored by studies (top) and by estimated unwanted variation from RUVprps (bottom). (d–i), Assessment of RUV-III-PRPS applied in a fully supervised setting (d); Scatter plots of the first three PCs of the RUV-III-PRPS–normalized data, colored by study (top row) and PAM50 subtype (bottom row) (e); Association of the RUV-III unwanted factor matrix (W) with biological and technical variables (f); Comparison of tumor purity estimates between FPKM data and RUV-III-PRPS–normalized data (g); Co-expression patterns of selected well-characterized breast cancer marker genes (h); Summary matrix of normalization performance metrics for RUVprps (i); Linear mixed-effects model analysis comparing variance components in the FPKM and RUV-III-PRPS–normalized data. (j–n) As in (d–i) but showing corresponding assessment results for RUV-III-PRPS applied in a fully unsupervised manner.

For fully supervised RUVprps normalization, the PAM50 subtypes and tumor purity were considered as known sources of biological variation, whereas studies were treated as known source of unwanted variation. RUV-III was applied using different sets of PRPS, selected NCGs, and values of *k* (Supplementary File). Fig. 7d clearly shows that RUV-III normalization effectively removes study-related variation while preserving the expected biological clustering of PAM50 subtypes. Figure 7e demonstrates that RUV-III strongly captures differences between studies while remaining weakly associated with known biological variables. The strong correlation between tumor purity estimates derived from RUV-III–normalized data and those from FPKM data indicates that RUV-III preserves biological variation. Consistent with this, well-established gene–gene correlation patterns are maintained following RUV-III normalization. Finally, linear mixed model analyses applied to FPKM and RUV-III-PRPS–normalized data show that RUV-III-PRPS effectively removes study effects while preserving variation associated with known biological factors.

Fig. 7j–n show analogous analyses for RUV-III-PRPS normalization performed in a fully unsupervised manner. In this setting, the estimated sources of unwanted variation were treated as technical variation, while all biological variation was assumed to be unknown. Similar to the supervised application, the unsupervised approach resulted in satisfactory normalization of the three large RNA-seq datasets.

### GTEx RNA-seq data

Fig. 3 shows that GTEx RNA-seq datasets are influenced by multiple sources of unwanted variation, including total ischemic time, defined as the interval between actual or presumed death (or cross-clamp application) and final tissue stabilization. We selected GTEx adipose tissue RNA-seq data comprising two subtypes, subcutaneous and visceral—to assess the ability of RUVprps to remove this source of unwanted variation. PCA of the raw count data shows that the first PC primarily captures variation explained by the ischemic time (Fig. 8a). This result is further supported by different gene-level and global assessments, RLE plots, differential gene expression analyses between the groups of the ischemic time, and correlation analyses between individual gene expression levels and total ischemic time (Fig. 8b,c). A linear mixed model followed by gene-level variance analysis further indicates that a substantial number of genes are affected by ischemic time (Fig. 8d). Further, Fig. 8f gives two representative examples showing how gene expression is affected by ischemic time. These results highlight the fact that genes are influenced by unwanted variation in heterogeneous, gene-specific ways.

**Figure 8:**
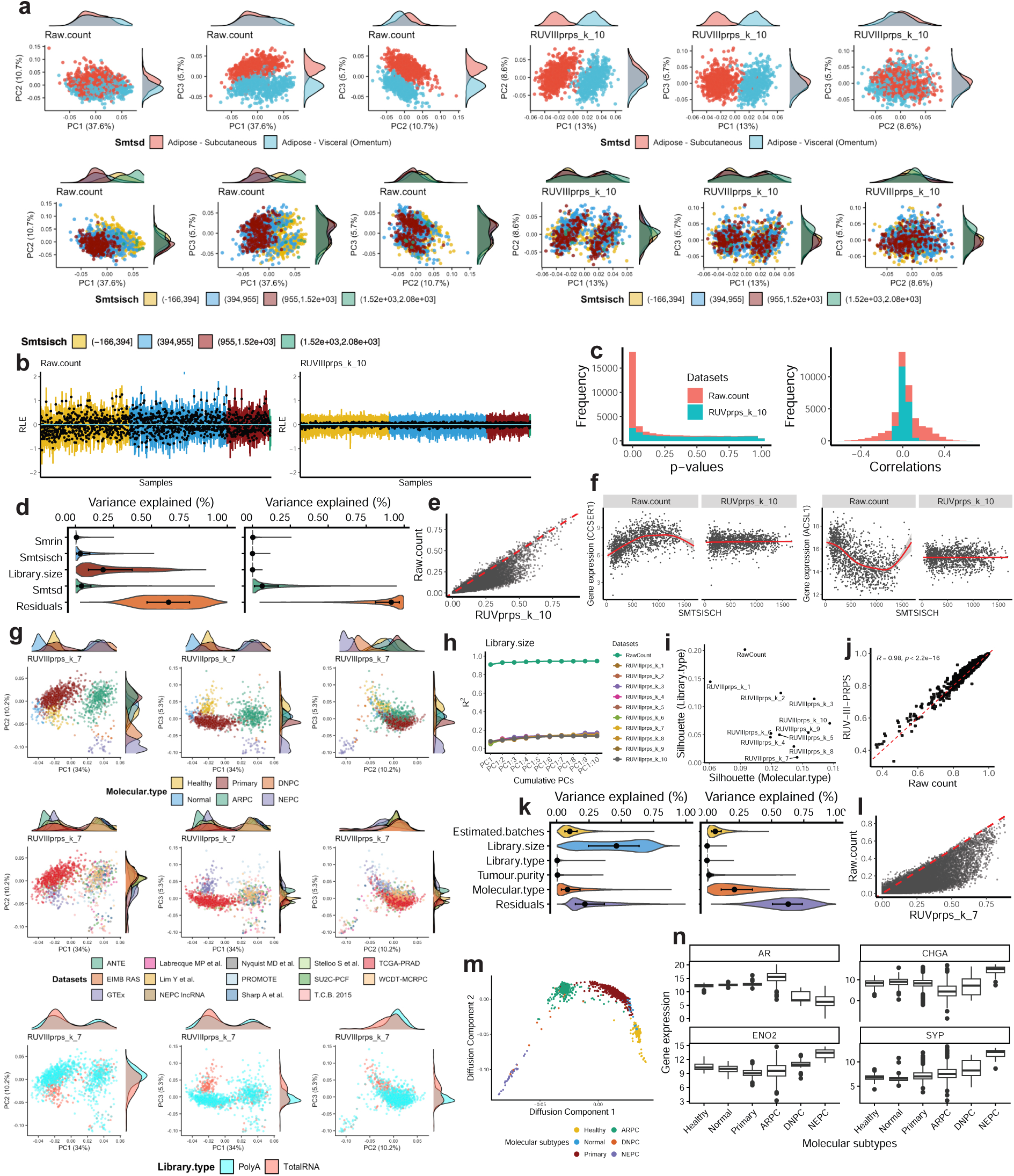
Supervised RUV-III normalization of GTEx and PCCAT RNA-seq data. a) Scatter plots of the first three PCs of GTEx adipose RNA-seq data using raw counts(log_2_ +1) and RUV-III-PRPS normalized data. Samples are colored by tissue type (top row) and by ischemic time (bottom row). b) RLE plots for the raw counts (log_2_ +1) (left) and RUV-III-PRPS–normalized data (right). RLE medians are colored and ordered according to ischemic time intervals. c) Left: Histograms of p-values obtained from differential expression analyses between all pairs of ischemic time intervals using raw counts (log_2_ +1) and RUV-III-PRPS–normalized data. Right: Distributions of correlation coefficients between individual gene expression levels and ischemic time for the raw (log_2_ +1)and RUV-III-PRPS–normalized data. d) Linear mixed-effects model–based variance decomposition showing the contribution of known sources of unwanted variation and biological factors to gene expression variability for the raw counts(log_2_ +1) (first plot) and RUV-III-PRPS normalized data (second plot). e) Scatter plot comparing the proportion of gene expression variance explained by the tissue types in the raw counts (log_2_ +1) to that after RUV-III-PRPS normalization. f) Scatter plots illustrating the expression of two representative genes differentially affected by ischemic time in the raw data (log_2_ +1). Plots show the expression of those genes in both raw data (log_2_ +1) and the RUV-III-PRPS normalized data. g) Scatter plots of the first PCs of the RUV-III-PRPS–normalized PCCAT RNA-seq data. h) Line–dot plots showing the cumulative association between the first ten PCs and library size across different normalized datasets. i) Scatter plots of silhouette coefficients for major molecular subtypes and library preparation protocols. j) Correlation between tumor purity estimates derived from raw count data (log_2_ +1) and that from RUV-III-PRPS–normalized data. k) Linear mixed-effects model–based variance decomposition quantifying the contributions of known sources of unwanted variation and biological factors to gene expression variability. l) Scatter plots showing the proportion of gene expression variance explained by tissue type in the raw log_2_(*x* + 1)-transformed data (left) and after RUV-III-PRPS normalization (right). m) Diffusion map of the RUV-III-PRPS–normalized PCCAT data showing a biologically expected trajectory from healthy prostate tissue to metastatic prostate cancer. n) Expression patterns of established prostate cancer marker genes across the RUV-III-PRPS–normalized PCCAT data samples.

We next applied RUVprps in a fully supervised setting, treating total ischemic time and library size as sources of unwanted variation and adipose tissue subtype as the primary biological variable of interest. PCA demonstrates that RUVprps effectively removes variation associated with ischemic time while preserving the biological separation between adipose tissue subtypes (Fig. 8a). Consistently, both the gene-level and global assessments indicate that RUVprps satisfactorily mitigates unwanted variation (Fig. 8b,c). Linear mixed model analysis of the RUV-III–normalized data shows that gene-level variation explained by sources of unwanted variation is substantially mitigated, while variation associated with adipose tissue subtypes is preserved or enhanced. We refer to the Supplementary File for all details of RUVprps normalization process.

### Prostate cancer cell atlas

Normalization of the prostate cancer cell atlas bulk RNA-seq data is challenging due to substantial confounding between major molecular subtypes and study-specific effects. For example, all healthy prostate tissue samples were obtained from the GTEx RNA-seq dataset. In addition, the cohort includes data generated using two different RNA-seq library preparation protocols, poly(A) selection and total RNA sequencing, further complicating normalization. We applied RUV-III with PRPS in a fully supervised setting, in which the major molecular subtypes or tissues and estimated tumor purity were considered as known biological variables, whereas library size, library preparation protocols and the studies were considered as sources of unwanted variation. Detailed descriptions of data preprocessing and RUV-III normalization are provided in the Supplementary File.

The first three PCs of the RUV-III–normalized data clearly indicate that RUV-III achieves satisfactory normalization (Fig. 8g).These plots clearly show that RUV-III with PRPS preserved the expected biological separation between all molecular subtypes in the data. Linear regression analyses between the first ten PCs and library size show that RUV-III normalization effectively removes the impact of library size from the data (Fig. 8h). Silhouette coefficient analysis demonstrates that RUV-III normalization reduces the effect of library preparation protocols while preserving clustering by molecular subtype (Fig. 8i). Tumor purity estimates derived from the RUV-III–normalized data indicate that this biological variation is preserved (Fig. 8j). Linear mixed model analysis shows that RUV-III-PRPS effectively reduces the contribution of multiple sources of unwanted variation while preserving, and in some cases enhancing, variation explained by to the molecular subtypes. Notably, due to the presence of strong confounding effects in this dataset, near-zero variance attributable to unwanted sources is not expected. Gene expression diffusion analysis reveals a biologically expected transition from healthy prostate tissue to metastatic disease, further confirming preservation of biological signal following normalization (Fig. 8m). Finally, expression patterns of several well-characterized genes associated with molecular subtypes, particularly neuroendocrine prostate cancer—remain biologically consistent after RUV-III-PRPS normalization (Fig. 8n).

## Discussion

Accurate identification of unwanted variation, efficient normalization and comprehensive evaluation of normalization are essential steps for all gene expression data analysis, regardless of the scale of the study. RNA-seq datasets are consistently impacted by one or more sources of unwanted variation, which can result in inaccurate or misleading outcomes if not corrected. This paper introduces RUVprps, a comprehensive R package that offers three major steps in normalization of RNA-seq data. First, it provides a wide range of statistical and computational tools, categorized as global and gene-level approaches, proposed to reveal unwanted variation in RNA-seq data. If sources of unwanted variation are not completely known, RUVprps can identify them using several unsupervised and robust approaches. Second, it offers RUV-III normalization with either genuine control samples, pseudo control samples or a combination of both. Selection of NCG and creation of PRPS can be accomplished using supervised and unsupervised approaches. Third, RUVprps provides a range of global and gene-level tools to visually and numerically assess the performance of normalizations.

In this study, we analysed a number of large-scale RNA-seq datasets spanning cell lines and tissue samples from both healthy and cancer samples to evaluate the performance of RUVprps across a range of real-world scenarios. The results clearly demonstrate that the package can satisfactory deal with RNA-seq data in situations ranging from all sources of unwanted and biological variation being known to when all sources being unknown. In situations when all or some sources of unwanted variation are unknown, RUVprps can robustly identify them. This feature of RUVprps creates a valuable opportunity for the effective use of publicly available datasets, for which complete details of experimental design and sample processing are often unavailable. Furthermore, RUVprps can estimate sources of unwanted variation even when they are strongly correlated with biological variation. We showed that RUVprps achieved satisfactory performance on the PCCAT dataset, which is characterized by different sources of unwanted variation and substantial confounding.

Analysis of 2,400 RNA-seq studies from the Recount3 resources, as well as several large RNA-seq projects, clearly highlights the importance of using both gene-level and global assessments to evaluate unwanted variation in RNA-seq data. Notably, in a large number of studies, the first three PCs did not effectively capture unwanted variation, whereas the gene-level analyses revealed its presence.

The fully unsupervised application of RUV-III-PRPS implemented in RUVprps is particularly valuable, as complete information on all sources of biological variation is often unavailable or cannot be reliably inferred from raw count data alone due to presence of unwanted variation. Across multiple studies, the unsupervised application of RUV-III-PRPS consistently demonstrated strong performance. In more complex scenarios, an unsupervised application can be used for initial normalization to mitigate dominant unwanted variation, followed by estimation of known biological factors and a subsequent supervised normalization step, potentially resulting in improved overall normalization.

By leveraging the *SummarizedExperiment* infrastructure, RUVprps minimizes common user errors, particularly for users who are not familiar with the RUV normalization framework. Furthermore, although RUVprps can be run using only a small number of high-level functions, the availability of modular, stand-alone functions enables advanced users to optimize parameters for their specific study designs and aims.

One of the major advantages of RUVprps is its ability to address any source of variation considered unwanted in a study. By specifying these variables as unwanted, RUVprps can identify true NCGs and generate PRPS data to estimate and remove them from the dataset. Moreover, RUVprps can simultaneously account for multiple sources of unwanted variation. We demonstrated that RUVprps effectively removed the effects of library size, tumor purity, and batch effects in the TCGA BRCA RNA-seq data while preserving known biological variation associated with the PAM50.

Normalization of three large breast cancer RNA-seq datasets and PCCAT data clearly demonstrates that RUVprps can effectively normalize RNA-seq data across different studies. This indicates the promise of RUVprps as a valuable tool for consortium-level data normalization, where datasets from multiple studies are combined.

Finally, many normalization methods have been proposed for RNA-seq data. Regardless of the method chosen, it is essential to assess its performance. Such assessments will include the effectiveness of the method to mitigate or remove the impact of unwanted variation, as well as its ability to retain or enhance known biological variation. RUVprps provides the tools for doing this.

## Methods

### Datasets

TCGA RNA-seq datasets were downloaded using the TCGAbiolinks R package (version 2.3). Batch annotations for the TCGA RNA-seq datasets were obtained from our previous study [3]. Clinical information, including overall survival, was obtained from Liu *et al.* [23]. GTEx gene expression read count data (version v10) and corresponding sample annotations were downloaded from the GTEx Portal (https://gtexportal.org/home/downloads/adult-gtex/bulk_tissue_expression). The Treehouse RNA-seq dataset was downloaded from the UCSC Xena Browser (https://xenabrowser.net/datapages/?hub=https://xena.treehouse.gi.ucsc.edu:443). CCLE gene expression data were downloaded from the CCLE data portal (URL). PCCAT gene expression data and sample annotations were obtained from the Prostate Cancer Cell Atlas website (https://pccat.net). Recount3 datasets were downloaded using the recount3 R/Bioconductor package (version 1.16.0). Two additional non-TCGA RNA-seq datasets were downloaded from the NCBI Gene Expression Omnibus under accession numbers GSE96058 and GSE81538 [21, 22]. For reproducibility, all RNA-seq datasets except those obtained from Recount3 were deposited on Zenodo.

### Overview of the RUV-III model

Before describing the linear model underlying RUV-III, we first introduce the *m* × *m*_1_ mapping matrix *M*, that connects the assays (individual-level measurements) to different samples (biological replicates), which captures the replication pattern in data (Fig. 9). Here, *m* is the number of assays, and *m*_1_ is the number of distinct samples being profiled. *M* (*i, h*) = 1 if assay *i* is on sample *h* and *M* (*i, h*) = 0 otherwise. Each row of *M* sums to 1, and the columns sum to the distinct sample replication numbers, the elements of *M^T^ M*. We also define an *m*_1_ × *p* design matrix *X* to capture the biological factor(s) of interest indexed by sample rather than assay. There are no constraints on *p*; indeed, *X* could be the *m*_1_ × *m*_1_ identity matrix. Our goal here is to remove unwanted variation, not to estimate regression parameters. The linear model we use is:

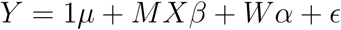

where the data *Y* = (*y_ij_*) and the unobserved errors *ɛ* = (*ɛ_ij_*) are *m* × *n*; the matrices *X* and *M* have just been defined; *µ* is the 1 × *n* row of gene means; *β* is *p* × *n*; the matrix *W* whose columns capture the unwanted variation is *m* × *k*; and *α* is *k* × *n*. 1 = 1*_m_*is the *m* × 1 column vector of 1s. Here *n* = number of genes, *p* is the dimension of the wanted variation *X* i.e. the biological factors of interest and *k* that of the unwanted variation *W*. We assume that *W* ⊥ 1. Also, we suppose that we have a subset of *n_c_*negative control genes whose *m* × *n_c_* submatrix *Y_c_* satisfies *Y_c_* = 1*µ_c_* + *Wα_c_*+ *ɛ_c_*, where we have assumed that *β_c_*= 0—that is, that there is no true association between these genes and the biology of interest. The projection *P_M_* = *M* (*M^T^M*)*^−^*^1^*M* replaces each entry *y_ij_* of *Y* by the simple average of the entries *y*_i′j_ over all *i^′^*for which *M* (*i, h*) = *M* (*i^′^, h*) = 1—that is and over all *i^′^* such that *i^′^* and *i* are label replicate or pseudo-replicate assays of the same unique sample (or pseudo-sample) labeled *h*. Write *R_M_* = *I* − *P_M_* for the corresponding residual projector. This is our source of information on the unwanted variation that we will remove. If the replication is technical at some level, then *R_M_ Y* mainly contains information about unwanted variation in the system after the technical replicates were created. Depending on the study details, technical replicates could be created immediately before the assay is run, in parallel with or immediately after sample is collected or somewhere in between. The earlier the creation of technical replicates, the more unwanted variation will be captured in their differences. The use of pseudo-replicates of suitable pseudo-samples enables us to deal with pre-technical unwanted variation. Write the spectral decomposition of *R_M_Y Y ^T^ R_M_* = *UDU^T^*, where *U* is an *m* × *m* orthogonal matrix and *D* is an *m* × *m* diagonal matrix with entries ordered from largest to smallest eigenvalue. Let *P*_1_ be the orthogonal projection onto 1*_m_*. For a chosen *k*, 1 ≤ *k* ≤ *m* − *m*_1_, define 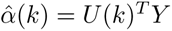, where *U* (*k*) is the first *k* columns of *U*. We estimate *W* by regressing the centered negative controls (*I* − *P*_1_)*Y_c_* on 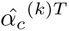. and then form the adjusted/normalized data *Y*, 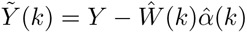

**Figure 9:**
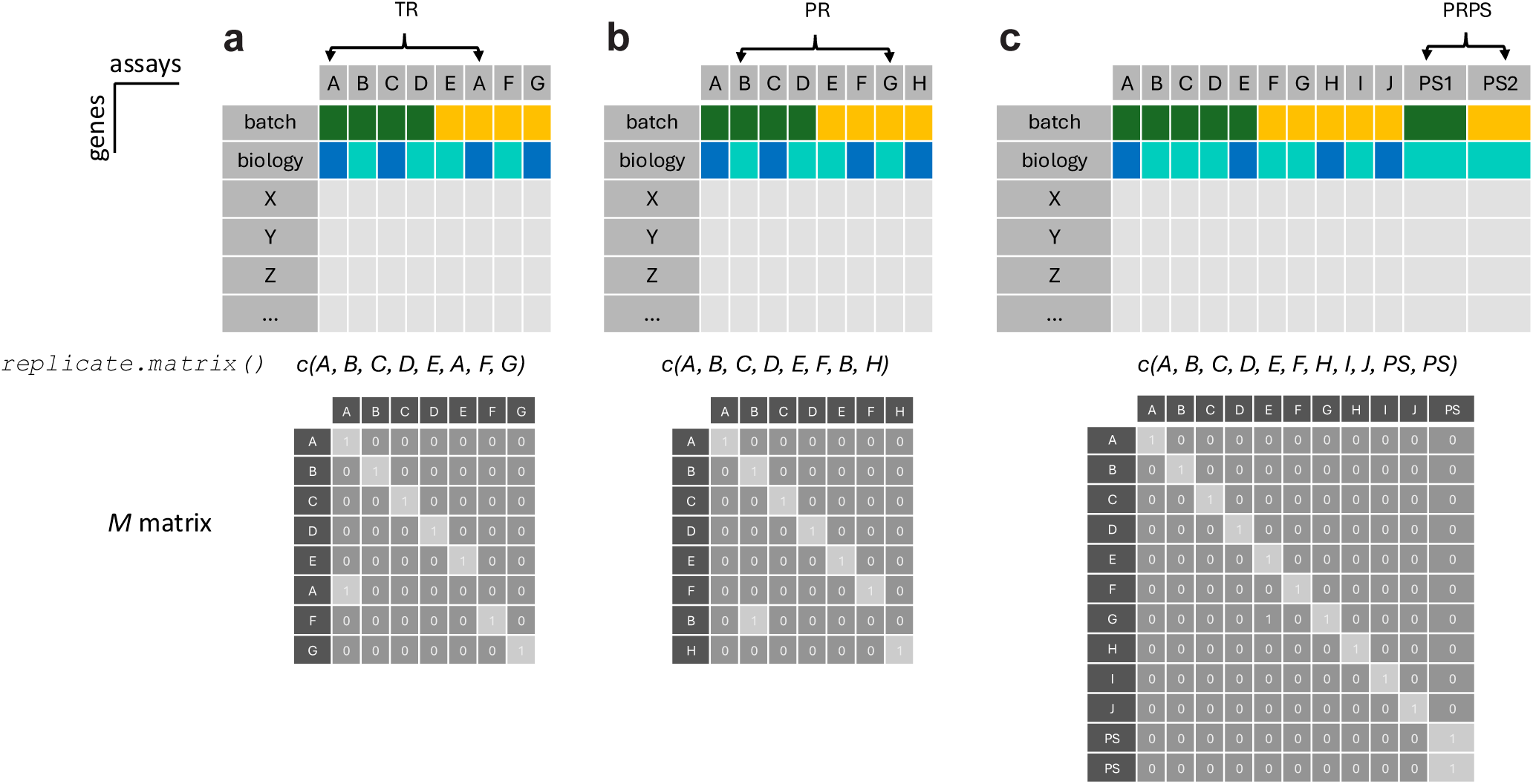
Illustrative examples of M matrix structure for different types of control samples in RUV-III. In these examples, top panels represent the gene expression matrix of a group of assays (columns) and a list of genes (rows). In each matrix, the first row indicates the experimental batches (dark green and orange), and the second row shows the biological groups (blue and cyan). Bottom panels display the replicate matrix M for each example. a) In this example, sample A is repeated across two batches as technical replicates (TR). The function `replicate.matrix()` can be applied to the column names of the gene expression matrix to create the M matrix. All assays are in rows and all unique samples are in the columns of the M matrix. b) In this example, samples B and G are considered as pseudo-replicates (PR). The labels of these two sample must be the same before applying the `replicate.matrix()` function. Here, the label of sample G is changed to B (though it can be done in another way or any unique labels that are different compared to the rest of the labels). c) In this example, samples B, C and D are averaged to create pseudo-sample PS1 and samples F, G and I are averaged to create pseudo-sample PS2. PS1 and PS2 are then added to the initial gene expression matrix as two extras columns representing pseudo-replicates of pseudo-samples (PRPS). The labels of the PS1 and PS2 must be the same before creating the M matrix. Here, those labels are changed to PS. These two samples are considered as a PR set for RUV-III normalization with PRPS. The overview of the RUV-III model has rows and columns representing samples and features, respectively, consistent with previous publications [3, 5]. Then, the RUV-III normalized data will be in the same format. For consistency with R, the RUVprps package transposes the RUV-III normalized data back to the assay format, with rows representing features and columns representing samples for downstream analysis.

### Different types of control samples in RUV-III

RUV-III uses sets of control samples to estimate the gene-wise effects of unwanted variation in high-throughput omics data including RNA-seq. The gene expression differences among a set of control samples indicates the presence of unwanted variation. There are three types of control samples that can be used in RUV-III.

#### Technical replicates (TR)

These are genuine control samples where the same biological sample is measured at least two times across anticipated sources of unwanted variation in a study. For example, if multiple reagent lots will be used for gene expression profiling, including technical replicates across the reagent lots is highly desirable [5, 24].

#### Pseudo-replicates (PR)

Pseudo-replicates are samples that are not true replicates but are used to estimate unwanted variation in the data [5]. These include samples from the same or highly similar biological groups that are measured across batches. The major gene expression differences between samples from a pseudo-replicate set will be unwanted variation. For example, when studying cancer and normal tissues, we can use normal tissues samples as pseudo-replicates if they are well-distributed across sources of unwanted variation [5].

#### Pseudo-replicates of pseudo-samples (PRPS)

Pseudo-replicates of pseudo-samples are a special type of pseudo-replicate set used in RUV-III. They involve creating pseudo-samples by aggregating data from multiple homogeneous biological samples. Pseudo-replicates of these pseudo-samples are then used to estimate unwanted variation. Averaging samples to form pseudo-samples can reduce the genuine biological heterogeneity between the samples so that the major gene expression differences between individual PS in a PR set will be unwanted variation [3]. It is important to note that, when sample size permits, PRPS sets should be homogeneous for each source of unwanted variation to avoid PRPS contamination. This approach is essential for RNAseq studies, where significant biological variation is expected across samples.

#### Key considerations

(1) There are sources of unwanted variation that cannot be corrected by using only genuine technical replicates. For example, if variation in tumor purity is considered an unwanted source of variation, the gene expression differences between any set of technical replicates will not capture that variability. Other examples include unwanted effects due to age or gender, which also cannot be accounted for through technical replicates alone. In such situations, we must use the PR or PRPS approaches. (2) When technical replicates are available but insufficient to capture all sources of unwanted variation, we recommend using them in combination with PR or PRPS. For instance, in a cancer RNA-seq study affected by both library size variation and batch effects, technical replicates may only account for batch-related differences. To remove all unwanted variation, it is still necessary to create PR or PRPS to address variation due to library size, alongside the use of technical replicates. (3) Samples used as technical replicates should have similar gene expression profiles to the other samples in the study. For example, lung cancer tissue samples should not be used as technical replicates in a breast cancer study, as there are breast tissue-specific genes that may not be present in lung cancer tissue, leading to missing information for those genes in the technical replicates. Additionally, samples used as technical replicates should have experienced the same conditions as the other samples in the study. If not, the technical replicates may exhibit variation that is unrelated to that experienced by the rest of the samples, or they may fail to capture certain variation’ that affect the other samples. (4) It is essential to achieve a reasonable balance i.e. to have similar numbers of control samples for each source of unwanted variation. In situations with two sources of unwanted variation, if a larger number of control samples are created for one source compared to the other, it is possible that the other source will not be satisfactorily corrected. (5) We highly recommend assessing the performance of control samples before using them in RUV-III as explained in subsequent sections. Ideally, variation in control assays should have high and low association with unwanted and wanted variation i.e. the biological factors of interest, respectively. (6) All samples or pseudo-samples in an individual replicate or pseudo-replicate set of control samples must have the same unique label before creating the M matrix for RUV-III. For example, if we have an RNA-seq data with two sets of control samples with each set contains three samples. The labels for one set of control samples could be “Rep1”, “Rep1”, “Rep1”, and the labels for a second set could be “Rep2”, “Rep2”, “Rep2”. The remaining samples should each have their own unique labels, such as “sample1”, “sample2”, and so on. (7) A PR set of control samples consists of at least two assays (samples), which are considered “replicates” of each other. The `replicate.matrix()` function treats any samples with the same labels as a PR set of control samples.

### Sources of unwanted variation in gene expression data

We define any source of variation that influences gene expression and compromises the accurate interpretation of downstream analyses as *unwanted* variation. These variations can be divided into two groups: those that influence between-sample analyses and those that impact within-sample analyses. The most apparent sources of unwanted variation between-samples in gene expression data are technical factors, such as differences in library size, sequencing depth, and batch effects. Batch effects can arise when data are generated across different experimental conditions, including variations in platforms, sequencing chemistries, processing times, laboratories, and personnel. These factors can introduce systematic biases that obscure true biological signals or create spurious associations, thereby complicating downstream analyses. Another important source of technical variation is RNA degradation, which can distort transcript abundance estimates, particularly in archived or low-quality samples. In addition to technical sources, biological variation unrelated to the primary focus of a study may also be considered unwanted. For instance, in cancer transcriptomics studies, factors such as tumor purity, patient age, sex, or immune infiltration can introduce variability that complicates the identification of tumor-intrinsic expression patterns. Specifically, tumor purity, the proportion of malignant cells relative to the surrounding stromal, immune, and normal tissue components, can significantly influence gene expression measurements [3]. When the objective is to investigate cancer cell–specific transcriptional programs, variability in tumor purity acts as a confounding factor and should therefore be treated as unwanted variation. To obtain biologically meaningful insights that reflect the intrinsic properties of cancer cells, it is essential to mitigate both technical and non-target biological sources of variation during preprocessing and analysis.

### Creation of a *SummarizedExperiment* object

Tabular data, including gene expression matrices, sample annotations, and gene details, constitute the common data structures obtained after sequencing data quantification. RUVprps offers a function called `prepareSeObj()` that enable users to create a *SummarizedExperiment* object from the tabular data (Fig. 1a). The function can create a sample annotation, gene annotation, or both if they are not available. This allows all sample- or gene-level details to be added to the object throughout the analysis. The function can also remove lowly expressed genes, calculate library size, estimates tumor purity, add several lists of housekeeping genes, include immune and stromal gene signatures [20], and supplement many details about individual genes. The function employs the `filterByExpr()` function from the edgeR R/Bioconductor package to find and remove lowly expressed genes [12]. Further, for cancer RNA-seq data, the function applies both the ESTIMATE [20] and singscore [25] methods to estimate tumor purity. It should be noted that the function can accommodate different data types for different purposes. For example, raw count data are required for calculating library sizes and for removing lowly expressed genes. If such data are not available, the filtering of lowly expressed genes must be performed externally prior to creating the *SummarizedExperiment* object. However, different data types such as CPM or FPKM can be used for the estimation of tumor purity, if they are available. The housekeeping gene sets were obtained from different studies and platforms including bulk and single-cell RNA-seq, microarray gene expression and Nanostring nCounter [26–28]. These genes may be used as a pre-selected NCG for several steps, including the identification of unknown sources of unwanted variation, the RUV-III normalization and normalization assessment. Further, the function can retrieve any given gene-level feature from the biomaRt R/Bioconductor package. The input data for the `prepareSeObj()` function can be also a *SummarizedExperiment* object and benefit from all or some of the mentioned options in the function.

### How to interpret the gene and global-level assessments

#### RLE boxplots

When data is well normalized, the medians of the boxplots in the RLE plot should be centered around zero, and the interquartile ranges should be similar across samples. An RLE plot with medians deviating from zero and varying interquartile ranges indicates the presence of unwanted variation in the data.

#### PCA plot

The scatter plots of the first few PCs after a good normalization should primarily reflect major biological groups, assuming they are distinguishable based on gene expression data, rather than being influenced by sources of unwanted variation. Additionally, the PCA plots of selected sets of true NCG (e.g., housekeeping genes) should not show any distinguishable clusters when the data is well normalized.

#### Line-dot plots of vector correlation and linear regression analysis

Ideal normalizations should result in low correlation values between the first several PCs and unwanted variables, and high correlation values between these quantities and known biological variables. It is recommended to use the average correlation behavior across multiple PCs rather than relying on individual points.

#### Barplot of ARI and silhouette coefficients

A better normalization method should lead to higher ARI and silhouette coefficients for biological groups and lower coefficients for batch groups.

#### Barplot of local inverse Simpson’s index (LISI)

When batch information is provided, a high LISI score indicates better mixing of batches, whereas when biological information is provided, a high LISI score reflects better separation of biological signals.

#### Barplot of k-nearest neighbor batch eflect test (kBET) score

A high kBET score indicates better mixing of batches. *Boxplots of ANOVA and correlations.* An ideal normalization should result in near-zero values for F-statistics and correlation coefficients between unwanted variables and individual gene expression levels. The analysis for biological variation will depend on prior knowledge regarding how genes are affected by these variables. For example,if there is variation in tumor purity and this is considered as biological variation, then we usually expect to see a proportion of genes that have high correlation with tumor purity.

#### Scatter plot from gene-gene partial correlation analysis

Unwanted variation can introduce spurious correlations or obscure true correlations between genes [3]. An ideal normalization method should show similar correlation coefficients between the partial and ordinary correlations for gene pairs.

#### Histograms of p-values from diflerential gene expression (DGE) analysis

In the absence of batch effects, DGE analysis across batches should produce unadjusted p-values that are uniformly distributed (i.e., a flat histogram). Differential gene expression analysis between biological groups expected to differ should yield a reasonable number of differentially expressed genes. If no DE genes are detected, this likely indicates the presence of unwanted variation. *Key considerations:* (1) An ideal RLE plot, where medians are centered around zero and interquartile ranges are similar across samples, does not always indicate well-normalized data. In cases where data is over-corrected (biological variation is removed), the plot may still appear ideal. Therefore, we strongly recommend using multiple tools to assess the performance of normalization. (2) The medians of RLE boxplots can only reveal one source of unwanted variation at a time. Generally, they are highly associated with library size in raw count RNA-seq data. If the impact of library size has been effectively removed but variation in the medians persists, we recommend exploring associations with other variables, such as tumor purity. If a strong association is found and the variable is considered biological variation, we recommend adjusting for it (using either RUV-III or regression analysis) and then examining the association of the RLE medians with other variables. This process should be repeated until the medians of the RLE boxplots are all close to zero. Following this process, all sources of unwanted variation can be identified and normalized more effectively. (3) Correlation between the medians of RLE boxplots and individual gene expression levels should show no or weak correlations (4) If the PCA plots show no visible clusters after a normalization, this may indicate over-correction. We recommend exploring the PCA plots colored by biological factors of interest within homogeneous sample groups considering all sources of unwanted variation. (5) Even if the PCA plots of a pre-selected set of NCG (e.g., housekeeping genes) show no visible clusters after a normalization, this may not be strong evidence of unwanted variation having been removed, especially if the selected genes are not initially affected by it. (6) Partial correlation may be useful when biological variables are provided, in addition to their use for addressing unwanted variables. In this scenario, similar correlation coefficients between partial and ordinary correlations for gene pairs may indicate having removed the specified biological variation.

### Estimation of unknown sources of unwanted variation in RNA-seq data

RUVprps offers different robust approaches to estimated unwanted variation in RNA-seq data.

#### The rle approach

This approach relies on the robust performance of relative log expression (RLE) plot in revealing sources of unwanted variation in transcriptomics data [29]. The underlying assumption of this approach is that in the absence of unwanted variation, the RLE medians should be all zero or near zero, and the RLE IQRs should be similar to each other [3, 5, 29]. To identify unknown sources of unwanted variation, the `identifyUnknownUV()` function first computes the RLE of the specified dataset and extracts the medians and interquartile ranges (IQRs) of the RLE values. The function provides two approaches for identifying potential sources of unwanted variation based on the RLE data. If the samples are chronologically ordered, a changepoint algorithm can be applied to detect any deviations in the RLE medians. Each detected deviation may indicate a potential source of unwanted variation. The second approach involves applying supervised or unsupervised clustering techniques to the RLE medians, IQRs, or both, independently. In the absence of unwanted variation, no clearly distinguishable clusters are expected. However, if unwanted variation is present, distinct clusters may emerge and can be interpreted as technical batches for downstream analyses. The `identifyUnknownUV()` function supports several supervised and unsupervised clustering methods to identify such clusters.

#### Advantages

The key advantage of this approach is that it does not require prior knowledge about biological variation as RLE medians and IQRs are usually robust to them. This can be useful when biological variation are entirely or partially unknown. We previously demonstrated that the RLE plots exhibit high robustness in studies where a small panel of genes (200 - 700) with high biological variability were measured [5]. More importantly, when unwanted variation is largely confounded with biological variation, the RLE plot can reveal the true unwanted variation.

#### Disadvantages

In RNA-seq studies where a very small number of genes are affected by unwanted variation, the RLE medians and IQRs may not reveal them. Further, the variation in RLE medians and IQRs could reflect true biological variation in rare situations where a large number of genes are expected to be differentially expressed between biological conditions of interest. To overcome this issue, we recommend using a set of true NCG (see the selection of NCG section for more details) to generate the RLE data and then obtain the RLE medians and IQRs to estimate possible sources of unwanted variation.

#### The pca approach

In this approach, PCA is first applied to the specified data. Then, a supervised or unsupervised clustering technique is used to group samples based on the first few principal components in order to identify potential unknown sources of unwanted variation (Figure 3). The underlying assumption is that the impact of unwanted variation is larger than biological variation in the selected PCs. It should be noted that the `identifyUnknownUV()` function provides an option to select principal components regardless of the order of their variance. This is particularly useful in situations where the first few principal components capture biological variation [3].

#### Advantages

PCA can be more sensitive to technical variation that affects just a small set of genes, in contrast with the RLE plots. Further, PCA can capture different sources of unwanted variation simultaneously, compared to the RLE medians. For instance, in TCGA raw count RNA-seq data, the first few PCs typically capture variation associated with library size, tumor purity, and batch effects, whereas RLE primarily reflects library size effects.

#### Disadvantages

PCA is sensitive to both biological and unwanted variation. We recommend using a set of true NCG to perform PCA when biological variation is larger than unwanted variation or select some higher principal components for clustering analysis. Further, when the biological and unwanted variation are highly correlated, PCA is limited in its ability to distinguish them. In this situation using a set of true NCG to perform PCA is highly recommended.

#### The sample.scoring approach

This approach relies on gene sets that are capable of capturing unwanted variation while being unrelated to biological variation of interest. These gene sets can be either pre-selected or derived directly from the data [3]. Each sample is scored against each gene set individually, and supervised or unsupervised clustering techniques are then applied to these scores to identify potential unknown sources of unwanted variation (Fig. 3). In the absence of unwanted variation, the scores should not show significant variation, resulting in no distinguishable clusters. A notable example is the use of immune and stromal gene sets from the Yoshihara et al. study [20], which can be leveraged to estimate tumor purity as a source of unwanted variation.

#### Advantages

It can capture and quantify subtle sources of unwanted variation in situations where both RLE and PCA may not be able to effectively find them. For instance, if variation due to hypoxia is considered a source of unwanted variation in cancer RNA-seq data, a gene signature can be used to estimate hypoxia-associated variation in the data. Such variation is typically not captured by RLE or PCA.

#### Disadvantages

Pre-selected gene sets may not either capture or be affected by all sources of unwanted variation.

### Selection of negative control genes

In the RUV framework for RNA-seq data analysis, an ideal NCG is characterized by three key criteria. Firstly, it should exhibit reasonable expression levels: not too high, not too low; secondly, its expression should be influenced by some or all the same unwanted variation experienced by the genes in general; and thirdly, its expression should lack any real association with the biological factors of interest. In general, the success of a set of NCG in RUV-III is a property of the entire set, not each individual member. RUV-III shows robustness to suboptimally selected NCG, and we previously showed in some cases, all genes can be utilized effectively as NCG [5]. However, this is not always the case, and careful selection is sometimes required. It is essential to note that different sources of unwanted variation can require different sets of NCG, so that the final set of NCG may contain different subsets of genes to cover all sources of unwanted variation.

We offer two approaches to obtaining NCG, one which involves using pre-selected genes and the other using data-derived genes. The pre-selected approach relies on a set of available NCG such as housekeeping genes or RNA spike-ins, and the second approach uses the dataset itself to find suitable sets of NCG. Regardless of the approach used, we highly recommend that users assess the performances of selected NCG before using them for RUV-III normalization. We will explain how to perform this assessment in the next section. It should be noted that RNA spike-ins may not be affected by all sources of unwanted variation that endogenous genes are influenced by in RNA-seq study. Furthermore, these control sequences may have their own variation, which is unrelated to the variation of endogenous genes.

There are five major steps involved in selecting NCG in both supervised and unsupervised approaches. The first step involves data pre-processing (Supplementary File) before applying any statistical tests. This mitigates the impact of either unwanted or biological variation, helping to identify genes influenced by each type of variation. This step is not required for all NCG selection functions. The second step is to define homogeneous sample groups with respect to known biological and unwanted variables separately. The third step involves applying statistical tests to quantify the variation of each gene with respect to both biological and unwanted variables. The fourth step is to summarize the statistical results from third step and select the most suitable NCG (Supplementary File). The final step is to evaluate the performance of the selected NCG.

### Supervised selection of negative control genes

The first function, `findNcgSupervisedByTwoWayAnov()`, employs a two-way ANOVA approach to find a set of suitable genes as NCG. In this method, all specified biological and unwanted variables are separately summarized into two categorical variables (see the `createHomogeneousBioGroups()` and `createHomogeneousUvGroups()` functions for more details). A two-way ANOVA is then applied to the expression levels of individual genes, using the summarized biological and unwanted variables as the two groups of factors. Genes are selected as NCG if they exhibit a high F-statistic or effect size for the unwanted variation, while accounting for the biological groups, and a low F-statistic or effect size for the biological variation, while accounting for the unwanted groups. See Supplementary File for available options to summarize the two F-statistics. The effect size is calculated as the proportion of variance explained by each factor, biological or unwanted variation group, relative to the variance remaining after controlling for the other factors.

The second function, `findNcgSupervisedByAnovaCorr()`, combines gene-level ANOVA and correlation analysis to identify genes strongly influenced by the specified categorical and continuous variables, respectively. This analysis is conducted separately for all unwanted and biological variables either across all samples or within groups of samples. The later analysis is performed within sample groups that are homogeneous with respect to either biological or unwanted variation. Genes are selected as NCG if they display appropriately strong associations with the unwanted variables and weak associations with the biological variables, as measured by both ANOVA and correlation results. We refer to Supplementary File for available options and details on how correlation coefficients and F-statistics are summarized for NCG selection.

The third function, `findNcgSupervisedByLinearMixedModel()`, fits a linear mixed model (LMM) to partition the variation (variance) in each gene across biological and unwanted factors [30]. Genes are then selected as NCGs if they exhibit high variation attributable to technical factors and low variation attributable to biological factors. See Supplementary File for available options to summarize the variation calculated for each group of variables.

#### Key Considerations

An implicit assumption of most functions that identify NCGs in a supervised manner is that the biological factors of interest are not strongly associated with unwanted variation. We strongly recommend assessing this association before applying the functions. In cases where biological and unwanted variation are highly correlated, we recommend using the `findNcgSupervisedByAnovaCorr()` function, with the option to perform the analysis within sample groups.

### Unsupervised selection of NCG

Methods to identify unknown sources of unwanted variation are described in the previous sections. The essential part of unsupervised selection of NCG is to identify genes that are highly variable due to biological variation. RUVprps offers two functions, `findNcgsUnSupervisedByAnovaCorr()` and `findNcgUnSupervisedByLinearMixedModel()` to find NCG in unsupervised manners. To streamline the analysis, these two functions are embedded within a major function, `findNcgUnSupervised()`. Similar to supervised selection of NCG, we recommend applying both approaches and comparing the results to identify the most appropriate set of NCGs.

The first function creates the most homogeneous sample groups with respect to all unwanted variation in the data. This reduces the influence of unwanted variation within each group, allowing for the detection of genes whose variability is more likely driven by true biological signals. The function performs an initial adjustment to remove sources of unwanted variation, such as library size or tumor purity, within each group. Then, it uses different statistical summaries e.g. MAD or CV to assess variability of each gene within each sample groups, to identify genes that are highly variable due to biological variation. Subsequently, the function utilizes gene-level correlations and ANOVA to identify genes significantly influenced by continuous and categorical sources of estimated unwanted variation, respectively. Lastly, various methods are used to combine the statistical summaries to identify a subset of genes that are strongly influenced by unwanted variation, while accounting for their biological variation (Supplementary File). The second function uses a LMM from variancePartition R package with several approaches to identify NCG (Supplementary Fig. 3). In all RNA-seq analyses in this study, categorical variables were modeled as random effects, whereas continuous variables were modeled as fixed effects. One of the approaches is to fit an LMM that includes only unwanted sources of variation, thereby selecting genes with the greatest proportion of variance attributable to technical factors as NCG. A second approach is to regress out the sources of unwanted variation from the data, using either a linear model or LMM, followed by performing PCA on the residuals. The first few PCs are then used as major sources of biological variation in the data. Finally, the LMM is fitted that incorporates both the unwanted variation factors and the selected PCs, allowing identification of genes that exhibit high variation explained by unwanted variation but low variation explained by to biological factors. We refer to Supplementary File and Figure 3 for more details.

#### Key Consideration

The main advantage of these approach is that prior knowledge of biological variation is not required. Furthermore, this approach may be more accurate in accounting for all types of biological variation in the data, rather than just the known ones (supervised approach). The main disadvantage is that it may be less effective compared to the supervised approaches, when biological variation is highly associated with unwanted variation. Furthermore, it may be inaccurate to identify truly highly variable genes driven by biological variation if the data preprocessing within sample groups that are homogeneous with respect to unwanted variables is inadequate.

### Assessing and comparing the performance of NCG sets

Each NCG selection function in RUVprps incorporates the `assessNCGs()` function, which evaluates how effectively the selected NCGs are associated with biological (when available) and unwanted variation. In this procedure, PCA is first performed on the data (the one used to find NCG) using only the selected NCGs. Subsequently, linear regression and vector correlation analyses are applied between the cumulative first few PCs and the continuous or categorical variables specified by the user, respectively. For each variable, the corresponding *R*^2^ or vector correlation values are plotted against the cumulative PCs. An optimal set of NCGs should exhibit high correlation with unwanted variables and low correlation with biological variables. Further, RUVprps provides a function called `compareNCGs()` that summarizes the results of linear regression and vector correlation analyses to compare the performance of different NCG sets. It outputs a score from 0 to 1 for each set, reflecting how well the capture unwanted variation while remaining unassociated with biological variables. To compute such scores, the correlations obtained for biological variables are subtracted from 1, and then the correlations for biological and unwanted variables are averaged, separately. The final score for each NCG set is calculated as the sum of half the average scores. A higher score indicates a better set of NCGs.

RUVprps offers the addNCGs() function which can be used to add any pre-selected list of genes such as housekeeping genes to the metadata of the *SummarizedExperiment* object and then their performance can be assessed by the assessNCGs() function and compared to other NCG sets in the the *SummarizedExperiment* object by the compareNCGs() function.

### Filtering selected NCG

RUVprps incorporates a filtration step within both its supervised and unsupervised functions to refine the selection of NCGs. In this step, the data-driven NCGs identified by RUVprps are cross-referenced with one or more publicly available cancer and non-cancer housekeeping genes sets. Genes that are consistently shared across these reference sets, provided the overlap is sufficiently large, are retained as the final NCGs. Importantly, this filtration step is flexible and can be performed using any other publicly available control gene sets, including those not bundled with RUVprps.

### Construction of PRPS

The pseudo-replicates of pseudo-samples (PRPS) essentially play the role of TR in the RUV-III method [3, 5]. The gene-wise expression differences between suitable pseudo-samples (PS) within a pseudo-replicate (PR) set should mainly be unwanted variation. A PRPS set contains at least two PSs created across a minimum of two batches, but may include additional PSs spanning more than two batches. PRPS must be created for all distinct sources of unwanted variation. For example, PRPS sets constructed based on library size will not capture unwanted variation coming from differences in tumor purity. It is essential to maintain a reasonable balance of PRPS sets for each source of unwanted variation. In situations where this criterion is not met, sources of unwanted variation with under-representation of PRPS sets may not be effectively removed. The created PRPS data of individual sources of unwanted variation is by default stored in the metadata of the *SummarizedExperiment* object and they are all combined for the RUV-III normalization. In addition, a sample annotation that contains the sample ids of each PRPS set is generated and stored in the metadata of the object for the PRPS assessment and visualization steps.

### Supervised generation of PRPS

RUVprps offers two functions, `createPrPsSupervisedForCategoricalUV()` and `createPrPsSupervisedForContinuousUV()`, which can be used to create PRPS sets for categorical and continuous unwanted variables, respectively. These functions are combined into a major function, `createPrPsSupervised()`, to facilitate the procedure. These functions first generate all possible homogeneous biological groups with respect to the user-specified biological variables. For example, if a biological subtype factor with five levels and one continuous biological variable are provided, the function will generate 5× the number of user-specified bins for the continuous variable. Then, PRPS data are constructed within each homogeneous biological group that is present across at least two batches. We refer to Supplementary File for more details.

### Unsupervised generation of PRPS

RUVprps offers three functions to create PRPS data in unsupervised way (Fig. 10). The first function, `createPrPsUnsupervisedByKnnMnn(`), uses a combination of *k*-nearest neighbor (KNN) followed by mutual nearest neighbors (MNN) identification. The function first applies a KNN algorithm within each batch to find at least two nearest neighbors (default setting) for each sample. The distances between each sample and its *k* nearest neighbors are computed, averaged, and then ranked to identify the most similar samples for PS construction. The lower the distances, the more similar the samples are. Note that the function can perform different data pre-processing within each batch to remove potential unwanted variation that can have a impact on the KNN analysis. For example, it performs CPM to remove the impact of library size before applying KNN. Then, the function performs MNN to find *m* mutual nearest neighbors across all possible pairs of batches. Finally, for each *m* MNN sets across a pair of batches, the corresponding identified KNN sets will be averaged to create a PS and consequently a set of PRPS. If the number of PRPS sets exceeds the user-specified limit, the function selects those KNN sets with the lowest ranks (smallest distances). The analysis is performed on either gene expression data or PCs using either all genes or a set of highly variable genes with respect to biological variation (highly recommended)(Fig. 10c).

**Figure 10:**
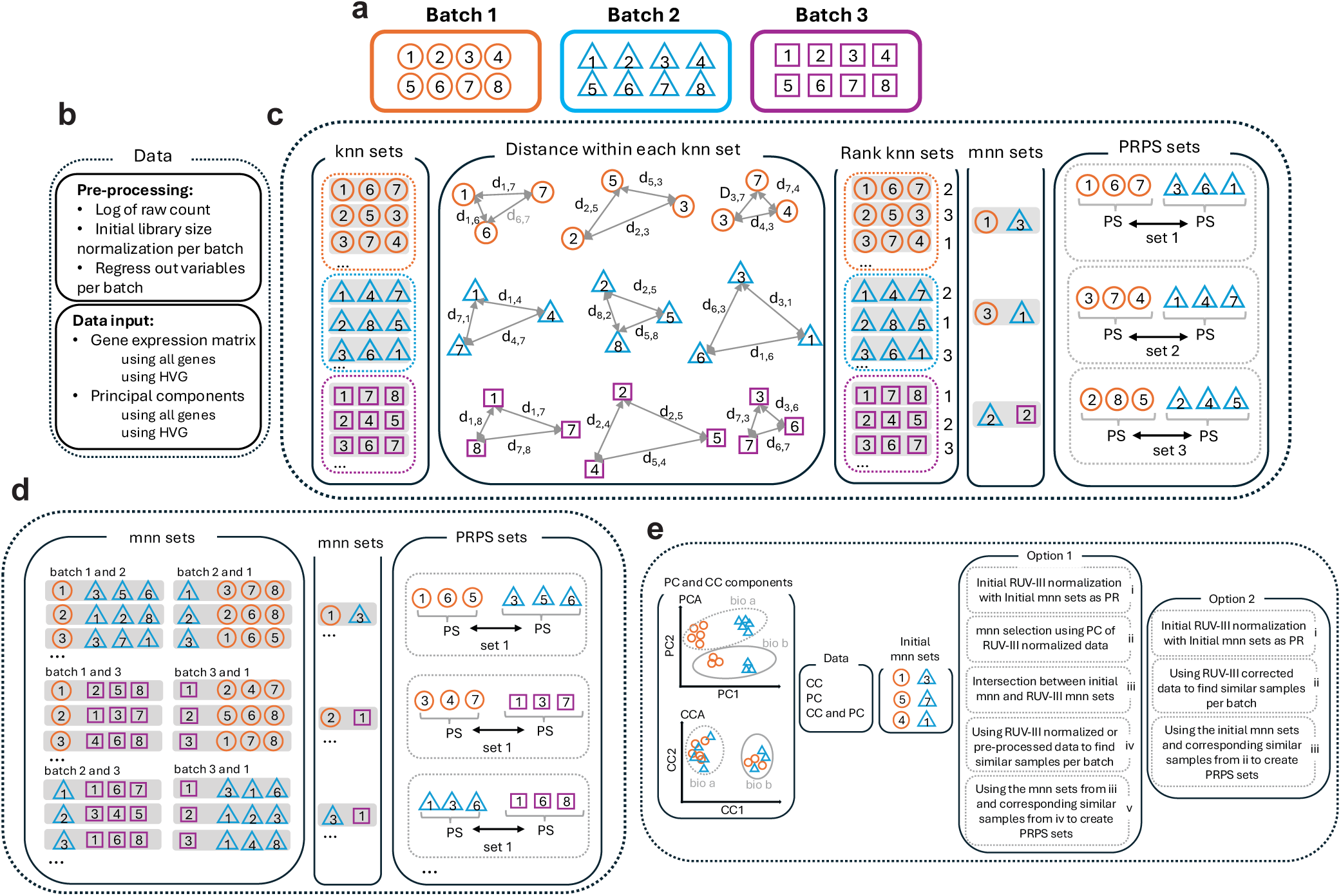
An illustration of the unsupervised approaches to creating PRPS data by the `createPrPsUnSupervised(`) function in RUVprps. a) Shows an RNA-seq study with three batches, each containing 8 samples. We assume that the biological factors of interest are reasonably well-distributed across batches. b) The data pre-processing procedure includes logging of raw counts, library size normalization per batch, and regression of specified variables per batch. The input data for identifying similar samples across batches can be a gene expression matrix containing all genes or a specified set of highly variable genes, along with the first several PCs of either all genes or the selected highly variable genes. c) Describes the steps involved in the `createPrPsUnsupervisedByKnnMnn()` function. Two (default number) KNNs for each individual sample per batch are identified. For example, the two KNN for sample 1 from batch 1 are samples 6 and 7. The KNN sets of the first three samples from each batch are shown here. The distances between each pair of samples in each KNN sets are calculated. The KNN sets are ordered based on their average distances, if filtration is specified by the user. The mutual nearest neighbors (MNNs) across all possible batch pairs are found. Finally, For each MNN sample, the corresponding set of KNNs is averaged to create a PS, and the two PS are considered as a set of PRPS data. d) Describes the steps involved in the `createPrPsUnsupervisedByMnn()` function. For each individual sample per batch, three (default number) MNNs from all other batches are identified. For example, sample 1 from batch 1 has samples 4, 5, and 6 as its MNNs from batch 2. The same sample has samples 2, 5, and 8 as its MNNs from batch 3. I) All possible MNNs between all possible batch pairs are identified. The corresponding MNNs from I are averaged to create PRPS data. e)Shows the steps involved in the `createPrPsUnsupervisedByCca()` function. For each pair of batches, CCA or CCA and PCA are performed to obtain the first few low-dimensional coordinates. The clusters in PCA are mainly driven by batch effects, whereas the clusters in CCA are driven by biological populations. Initial MNNs are then identified using coordinates from PCA or CCA or both separately. Option 1)These initial MNNs can serve as pseudo-replicates for the initial application of RUV-III. PCA is subsequently applied to the RUV-III normalized data, and the resulting PCs are used to identify another set of MNNs, which are then compared to those identified via initial CCA or PCA or both to obtain the common set. The RUV-III normalized data or a pre-processed data can also be used to identify the most similar samples within each batch. Finally, the similarities from each MNN set are used to generate PS and subsequently PRPS. Option 2) MNNs are identified solely based on CCA coordinates. The data are then pre-processed either by RUV-III with PRS or the other pre-processing to identify the most similar samples within each batch. Finally, the similarities from each CCA-derived MNN set are used to generate PS and, subsequently, PRPS. There more options in`createPrPsUnsupervisedByCca()`.

The second function, `createPrPsUnsupervisedByMnn()`, uses only MNN to find at least 3 MNN for each sample across batches (Fig. 4d). Then, each MNN sets will be averaged to create two PS and then they will be matched as a PR set. Similar to the previous function, the analysis cane be performed using either gene expression data or PCs using either all genes or a set of highly variable genes (Fig. 10d).

A potential limitation of the first two approaches is that the gene expression data or PCs used to identify MNNs are influenced by unwanted variation present in the study. This can compromise an accurate identification of MNN. To improve that, RUVprps offers the third function, `createPrPsUnSupervisedByCca()`, that applies canonical correlation analysis (CCA) for dimensionality reduction and uses the CC components to perform MNN analysis for PRPS construction. This function offers different strategies (Fig. 10e). First, for each pair of batches, dimensionality reduction is performed using both CCA and PCA. The first few CC and PC coordinates are then used separately to identify MNNs across the batches. The common MNNs obtained from CCA and PCA can be used as pseudo-replicates to apply an initial RUV-III for data normalization. The normalized data is subsequently subjected to PCA, followed by MNN detection using PCA coordinates. These MNNs are compared to the initial ones to identify the common subset. Next, either the RUV-III–normalized data or a pre-processed dataset is used to find the most similar samples within each batch using different statistical tests. Finally, the most similar samples (the default is three) of each MNN are averaged to generate PS and, ultimately, PRPS. The second approach relies only on CC coordinates to identify MNNs. In this case, a pre-processed dataset is used to find the most similar samples from each MNN set, which are averaged to generate PS and then PRPS.

Note that, both the supervised and unsupervised functions offer the option to either apply the log transformation to the samples before averaging them to create PS, or to average the samples first and then apply the log transformation.

### Contamination, distribution and connectedness of PRPS sets

We highly recommend assessing the contamination of PRPS sets with regard to unwanted variation and their distribution across batches before using them. In situations where multiple sources of unwanted variation are provided, it is possible for PRPS data created for one source to inadvertently capture variation from another. We refer to this as “contamination”. Such contamination may reduce the effectiveness of the PRPS in capturing variation from the intended source. To mitigate this, both the supervised and unsupervised PRPS functions can incorporate all specified unwanted variables when generating PRPS sets for a particular source. In other words, to generate PRPS for a given source, the other sources are taken into account when defining biologically homogeneous groups of samples. It should be noted that considering too many factors when grouping samples into homogeneous subsets may result in groups with too few samples. This can lead to situations where there are not enough samples to create PRPS. In such cases, decisions must be made regarding the relative importance of different sources of unwanted variation, and limited contamination from less important sources may need to be tolerated.

Both the supervised and unsupervised PRPS functions create different plots to show how well the PRPS sets are distributed across sources of unwanted variation. We refer to these plots as PRPS maps. There are helpful to visually explore how PRPS distributed across sources of unwanted variation. These plots are stored in the metadata of the *SummarizedExperiment* object by default.

We say that a replicate or pseudo-replicate set *R* spans two batches if *R* includes samples or pseudo-samples from both batches. When this is the case, RUV-III will remove the batch differences captured by the replicate or pseudo-replicate samples across those two batches. In situations involving multiple batches, it may not be possible to have a single replicate or pseudo-replicate set that spans all of them. In such cases, it is necessary to construct replicate or pseudo-replicate sets with connections that collectively cover all batches. We refer to Supplementary Figure 4 for some examples. The functions that create PRPS data include an argument called `check.prps.connectedness()`. When it is set to `TRUE`, the function checks the connectivity between PRPS sets and displays how they collectively cover all the specified batches.

### Selection of highly variable genes

Using highly variable genes (HVGs) driven by biological variation is helpful for identifying biologically similar samples across batches, which can then be used to create PS and subsequently PRPS in an unsupervised manner. However, identifying such genes prior to a proper data normalization is challenging. RUVprps provides the `findHVG()` function, which implements multiple statistical tests and strategies to identify highly variable genes (HVGs) in RNA-seq data. These include statistical methods based on median absolute deviation (MAD), coefficient of variation (CV), variance, variance-stabilizing transformation (VST), LMM or different combination of them. Most methods are generally applied to homogeneous groups of samples with respect to unwanted variation to find HVGs. The resulting statistical values across groups are then summarized to select a specified number of HVGs. The function offers a variety of data processing options designed to mitigate the influence of unwanted variation and improve the identification of true HVGs within each groups.

The LMM approach in `findHVG()` is analogous to the method used to identify NCGs. However, in this case, the LMM results are used to detect genes whose variability is primarily explained by biological rather than technical variation. It should be noted that The`findHVG()` can be applied in supervised and unsupervised way.

### Selecting maximum value of k for RUV-III normalization

To apply RUV-III with the chosen control samples and NCG set, one needs to specify the number *k* of (linear) dimensions of unwanted variation to estimate and remove from the data. It should be noted that RUV-III is generally robust to overestimating *k*, but not always. There is always a constraint on the maximum value of *k* for any specified set of control samples and NCG. RUVprps offers the `getMaximumK()` function that can be used to find the maximum possible *k* value. To compute this maximum *k* value:

1. Calculate the difference between the columns and rows of the *M* matrix:

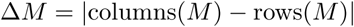
2. Consider the number of NCG:

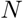
3. The maximum possible *k* value is the minimum of the above two values:

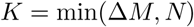

It should be noted that if the user-provided value of *k* exceeds the maximum possible value, the RUVIIIprps() function will automatically calculate the maximum allowed value and use it for normalization.

To address the limited efficiency in optimizing the RUV-III method with respect to the choice of *k*, we designed a novel approach that enables a fast and automated application process. In this approach, the alpha is calculated for the maximum value *K* of *k*, and then corresponding dimensions are the used for other values of *k*. This modification significantly speeds up the implementation of RUV-III with multiple values of *k*. Additionally, the residual operation step in the RUV-III method is replaced with C++ code, further accelerating the calculation. Altogether, the package includes two novel calculations that enable users to run an efficient and automated RUV-III normalization process.

### Assessing the unwanted variation factors estimated by RUV-III

RUVprps provides the `assessW()` function to evaluate the association between biological and unwanted variables with the estimated unwanted variation factors W from RUV-III. The function performs linear regression between each individual continuous variable specified by users and the columns of the W matrix in a cumulative manner, and then computes *R*^2^. For categorical variables, the function applies vector correlation between the variables and the columns of the W matrix. Ideally, the columns of the W matrix should show a strong correlation with unwanted variation and no or a weak correlation with biological variation. To compare the performance of different *k* values, the correlations obtained for biological variables are subtracted from 1, and then the correlations for biological and unwanted variables are averaged separately. The final score for each set of *k* values is sum of the half of the average scores. A higher score indicates a better value for *k*. It is important to note that the assessment of W should be used in combination with all other normalization assessments to select suitable RUV-III normalized data.

### The structure of the metadata in RUVprps

The graphical and numerical outputs of all functions in RUVprps are, by default, stored in the metadata of the *SummarizedExperiment* object (Supplementary Fig. 5). The metadata contains eight major slots, each corresponding to a specific group of outputs. The “Metrics” slot stores all graphical and numerical outputs from individual variation assessment functions for each dataset separately. When multiple datasets are provided and assessed, the resulting plots are combined into a single grid layout and stored in the “Plot” slot. In the RUV-III normalization procedure, all outputs from the NCG selection function are stored in the “NCG” slot, and outputs from the PRPS function are stored in the “PRPS” slot. During normalization, the relevant files from these two slots are retrieved and used. The estimated W factors from RUV-III and their associated diagnostic plots are stored in the “RUVIII.W” slot, while the estimated unknown sources of unwanted variation and exploratory analysis plots are stored in the “UnKnownUV” slot. Finally, the output of the `getAssessmentMetrics()` function, including both plots and assessment tables, is stored in the “AssessmentMetrics” slot. RUVprps provides two functions, `obtainMetric()` and `obtainPlot()`, which facilitate the retrieval of metrics and plots from the metadata, respectively. Most major RUVprps functions also include an option to remove the existing metrics or plots before re-running the function with different parameters.

### Linear mixed effects modeling for variance decomposition

To quantify the relative contributions of biological and technical factors to gene expression variability, RUVprps adopts variance partitioning using a linear mixed-effects modeling framework in variancePartition R package (version 1.36.3) [30]. Specifically, for each gene, expression is modeled as a function of multiple sources of variation, including known biological covariates such as tissue type and molecular subtype and technical or unwanted factors including batch, sequencing platform, and library preparation protocol The proportion of variance explained by each factor is estimated using restricted maximum likelihood, enabling decomposition of total expression variability into biologically meaningful and technical components. These gene-level variance fractions are subsequently aggregated across genes to obtain global estimates of the contribution of each factor. RUVprps use this model to find negative control genes and highly variable genes as well as assess the performance of normalizations.

## Software and data availability

All computational source code required for reproducing all results presented in this paper and the RUVprps R package are available in https://github.com/RMolania/RUVprps. RUVprps will be submitted to Bioconductor.

## Author contributions

R.M conceived and designed the study and the RUVprps package. R.M. developed the functions, the unsupervised approaches for NCG and PRPS, and fast implementation of the RUV-III method in the RUVprps package. R.M. and T.P.S designed the numerical normalization assessment pipeline. M.M and R.M implemented C++ computation. R.M, M.T and M.F performed the data analysis and prepared the vignettes. R.M and T.P.S drafted the manuscript, which was revised and approved by all authors.

## Acknowledgments

R.M. was supported by Prostate Cancer Foundation (PCF) Young Investigated Award and the Lorenzo and Pamela Galli Medical Research Trust. We thank Luke C. Gandolfo, Connie SN Li-Wai-Suen and Adrian Salavaty for their helpful comments on the RUVprps package.

## Competing interests

The authors declare no competing interests

## Notes

### Competing Interest Statement

The authors have declared no competing interest.

