## Supplementary File for "Fast, accurate and scalable normalization for RNA sequencing data with the RUVprps software package"

### Functions designed for assessing variation in RNA-seq data

Here, we describe individual functions in RUVprps that are designed to assess both biological and unwanted variation in RNA-seq data.

*Relative log expression (RLE).* The `computeRLE()` function in the RUVprps package computes the RLE of all specified datasets (assays) in the `SummarizedExperiment` object. The function can return specific properties of the RLE data including the medians, interquartile ranges (IQRs) or both. To visualize the RLE data, the package provides two functions: `plotRLE()` and `plotRleVariable()`. The `plotRLE()` generates boxplots of the RLE distributions for individual samples and colors the medians by a specified variable. We recommend ordering the samples based on known potential sources of unwanted variation to better visualize trends in that variation. The `plotRleVariable()` function plots the RLE medians or IQRs against a categorical or continuous variable and calculates an association metric using either ANOVA and correlation, respectively.

*Principal component analysis (PCA).* The `computePCA()` function calculates principal components (PCs) through the singular vector decomposition (SVD). The default parameter is to center the data without scaling before applying the SVD. The function also offers a fast approach for calculating the SVD for a certain number of singular vectors. It should be noted that when only certain number of left-singular values are calculated, the percentage variation of each component will only be based on that number, not entire possible number of singular vectors in the data. The `plotPCA()` function offers scatter and boxplots to visualize the PCs. The function can generate pairwise scatter plots of the first $n$ PCs colored by a categorical variable or creates boxplots of each PC across the variable. Boxplots of PCs become beneficial when there are many levels of a categorical variable, making visualization through colored scatter plots challenging. If a continuous variable is provided, the function creates a scatter plot of each PC against that variable. ANOVA and Correlation are calculated to assess the association between each PC and a categorical and continuous variable, respectively.

*Vector correlation analysis (VCA).* We utilize the Rozeboom squared vector correlation to quantify the strength of (linear) relationships between two sets of variables, such as the first $k$ PCs ($1 \leq k \leq 10$) and dummy variables representing unwanted or biological variables [1]. Not only does this quantity summarize the full set of canonical correlations, it also reduces to the familiar $R^2$ from multiple regression when one of the variable sets contains just one element. The `computePCVariableCorrelation()` function calculates the vector correlation and the `plotPCVariableCorrelation()` function generates a line-dot plot between the first few PCs, cumulatively, and the vector $R^2$.

*Linear regression analysis (LRA).* $R^2$ values of fitted linear models are used to quantify the strength of the (linear) relationships between a single quantitative source of unwanted variation, such as sample (log) library size or tumor purity, and global sample gene expression summary statistics, such as the first $k$ PCs ($1 \leq k \leq 10$). The `computePCVariableRegression()` function calculates the linear regressions and the `plotPCVariableRegression()`

function generates a line-dot plot between the first few PCs, cumulatively, and the $R^2$ values.

*Adjusted Rand Index (ARI).* The ARI is the corrected-for-chance version of the Rand index. It measures the proportion of matches between two label lists. The `computeARI()` function uses PCs calculated by the `computePCA()` function to compute the ARI for any specified categorical variable for each dataset. This function offers hierarchical clustering through the implementation of the `hclust()` function from the *stats* R package, and model-based clustering based on Gaussian mixture models using the `Mclust()` function of the *mclust* R package. The hclust option is recommended for its faster calculation of ARI. The `plotARI()` function will generate a barplot of ARI values for all specified datasets. If two variables are provided, the function can plot their ARI values against each other. Note that the PCs are not scaled prior to calculation of ARI.

*Silhouette coefficient.* Silhouette coefficient analysis is used to assess the quality of a sample clustering. To compute silhouette coefficient, the first few PCS are used to calculate both the distance between a sample and the other samples in its cluster and the distance between samples in different clusters.. The `computeSilhouette()` function uses the PCs calculated by the `computePCA()` function to compute silhouette coefficients for any specified categorical variable for individual datasets. This function offers several methods to calculate the distances using the `dist()` function in the *stats* R package. The silhouette coefficient of individual samples within each assay are averaged for a bar plot. The `plotSilhouette()` function will generate the barplot of averaged silhouette coefficients for all specified datasets. If two variables are provided, the function can plot their averaged values against each other. It should be noted that the PCs are not scaled prior to calculation of silhouette coefficient.

*Local Inverse Simpson's Index (LISI).* LISI is a metric originally proposed for single-cell RNA-seq analysis to quantify local diversity. RUVprps adopts LISI to evaluate batch mixing, i.e., how well samples from different batches or conditions are integrated. A high LISI score for batch indicates good mixing, meaning samples from different batches are well-integrated locally. Additionally, RUVprps uses LISI to assess biological population separation, i.e., how well distinct cell types remain separated after integration. A low LISI score for sample type indicates good separation, meaning samples of the same type cluster together. RUVprps provides the `computeLISI()` and `plotLISI()` functions to perform LISI analysis and visualize the results.

*k-Nearest Neighbor Batch Effect Test (kBET).* kBET is a statistical test designed to assess batch effects in single-cell data. It evaluates whether samples from different batches are well-mixed locally after normalization. For each cell, kBET examines the batch composition of its k nearest neighbors and tests whether the local batch distribution matches the expected global distribution. A low rejection rate indicates good batch mixing, meaning batch effects have been effectively removed, whereas a high rejection rate indicates poor mixing and residual batch effects. RUVprps provides the `computeKbet()` and `plotKbet()` functions to perform kBET analysis and visualize the results.

*Analysis of variance (ANOVA).* ANOVA enables us to assess the association between a given categorical variable (which we call a factor) on individual gene expression measurements across any set of groups (labeled by the levels of the factor) under study. The `computeGenesVariableAnova()` applies a fast implementation of one-way ANOVA using the `row_oneway_equalvar()` or the `row_oneway_welch()` functions from the *matrixTests* R package. The first function assumes equal variance between the groups while the second function applies the Welch correction for the groups with unequal variances. The `plotGenesVariableAnova()` generates boxplots of $log_2$ of F-statistics of individual assays.

*Correlation analysis.* The RUVprps package provides a function named `computeGenesVariableCorrelation()` to compute Spearman or Pearson correlations between the gene-level expression values of each data and a continuous variable e.g. library size or tumor purity in the data. This function offers both correlations through the implementation of the `correls` function from the *Rfast* R package. The `plotGenesVariableCorrelation()` function generates boxplots of correlation coefficients of individual datasets.

*Partial correlation.* Partial correlation is used to estimate the correlation between two variables while controlling for a third variable. This approach is used to compute gene-gene correlations while controlling for variable such as library size or tumor purity. RUVprps includes a function named `computeGenesPartialCorrelation()` to calculate both ordinary and partial correlation for all possible gene-gene pairs. The formula for the partial correlation $r_{xy.z}$ is given by:

$$r_{xy.z} = \frac{r_{xy} - r_{xz} \cdot r_{yz}}{\sqrt{(1 - r_{xz}^2) \cdot (1 - r_{yz}^2)}}$$

where $r_{xy}$ is correlation between variable $x$ and $y$, $r_{xz}$ is correlation of the third variable $z$ with the variable $x$, and $r_{yz}$ is correlation of the third variable $z$ with the variable $y$. The function `plotGenesPartialCorrelation()` generates histograms, scatter plots, and boxplots of both ordinary and partial correlation coefficients for individual datasets. This step can be computationally expensive when considering all possible gene pairs in RNA-seq data. Therefore, we recommend using a subset of genes that show a high association with the specified variable in the partial correlation analysis.

*Gene set scoring.* Gene set enrichment analysis is adopted to evaluate the overall expression levels of predefined gene sets in individual samples. Gene sets can be either biological or non-biological. RUVprps offers a function named `computeGeneSetScore` by implementing the *singscore* R package [2]. This function calculates sample-level scores for all specified gene sets for individual assays. Then, the function `plotGeneSetScore()` generates either histograms or paired-scatter plots of the scores of each data.

*Differential gene expression (DGE) analysis.* Differential gene expression analyses are performed using either Limma or the Wilcoxon signed-rank test with log-transformed data e.g. raw counts or normalized data. To evaluate the effects of the sources of unwanted variation on the data, DE analyses are performed across all possible pairs of batches. Further, differential gene expression analysis can be performed across the levels of a biological factor of interest. The `computeDGE()` function performs DGE analysis and `plotDGE` function generates histograms of unadjusted P values for individual dataset.

The `plotStudyOutline()` function generates a heatmap of sample-level features such as batches, biological populations, library size, and tumor purity. This plot is helpful for exploring how these factors are distributed across samples and for examining visible unwanted variation in the data. Further, it can visually reveal confounding of factors in the data. The plot can be generated initially using known variables and subsequently updated to incorporate estimated biological and unwanted factors during normalization. We highly recommend generating the plot before performing downstream analyses.

The `getAssessmentMetrics()` function creates a visualization showing all possible metrics that can be calculated for categorical and continuous variables, including both biological and unwanted sources of variation. It assigns a unique barcode to each metric, which can be used to specify which metrics to exclude during the variation assessment steps.

The `assessVariablesAssociation()` function examines the association between all specified biological and unwanted variation variables separately. If two continuous variables are highly correlated, the function keeps the one with the larger variance, and if two categorical variables are highly associated, the function keeps one with the larger number of levels. This assessment is useful for creating PRPS and finding NCGs. If two variables exhibit high correlation, utilizing just one of them may be sufficient for creating PRPS or for identifying NCGs. This step will speed up the computational processes underlining RUV-III normalization. Spearman correlation and Pearson's contingency coefficients are used to assess association between continuous and categorical variables, respectively. The function `ContCoef()` is adopted from the *DescTools* R package for the Pearson contingency coefficient. It should be noted that including two highly correlated variables in PRPS construction and NCGs identification steps will not cause an issue.

### Structure of *SummarizedExperiment* object

The *SummarizedExperiment* object is a data structure used in R for representing and manipulating high-dimensional experimental data. Here are some key features and components of the *SummarizedExperiment* object: The *SummarizedExperiment* object allows for the incorporation of multiple assays (experimental data). Each assay is a separate matrix of data associated with the same features and samples. Rows typically represent features (e.g., genes, transcripts), and columns represent samples or experimental conditions. The assay data can be accessed using the `assay()` function from the the *SummarizedExperiment* R/Bioconductor package.

*Row and Column data:* The *SummarizedExperiment* object includes data associated with both rows and columns. Row data can contain information about the features, such as gene annotations or genomics coordinates. Column data may include sample information, experimental conditions, or other relevant details. The row and column data can be retrieved using the `rowData()` and `colData()` functions from the *SummarizedExperiment* R/Bioconductor package.

*Metadata:* The Metadata of the *SummarizedExperiment* allows for flexibility in terms of data types and structures.

This makes it suitable for saving all outputs of the functions in the RUVprps R package. The metadata can be accessed through SummarizedExperiment@metadata.

**Notes:**(1) The overview of the RUV-III model has rows and columns representing samples and features, respectively, consistent with previous publications [1, 3]. Then, the RUV-III normalized data will be in the same format. For consistency with R, the RUVprps package transposes the RUV-III normalized data back to the assay format, with rows representing features and columns representing samples for downstream analysis.(2) Any changes to a *SummarizedExperiment* object, e.g. removing a gene or sample in an assay, will also be applied to the row or column data, and vice versa. However, these changes will not be reflected in the Metadata.

### General functions in RUVprps

RUVprps provides a suite of general-purpose functions that facilitate manipulation, exploration, and analysis of `SummarizedExperiment` objects.

The `removeAssay()` and `renameAssay()` functions can be used to remove specific datasets and to rename existing datasets within the *SummarizedExperiment* object, respectively.

The `renameVariables()` function can be used to change the column names of specified variables in the sample annotation (colData) of the *SummarizedExperiment* object.

The `createLogAssay()` applies a $log_2$ transformation to the counts from the specified dataset(s). If there are zero counts, a pseudo-count (default is 1) specified by users will be added to the measurements before applying the log transformation. The logged data can be either added as new assay or replace the corresponding assay in the *SummarizedExperiment* object. Each function within RUVprps provides the option to apply a logarithmic transformation to the data.

The `checkSeObj()` function assesses the structure of the `SummarizedExperiment` object and removes any missing values from specified dataset(s), variable(s) or both. Note that the current RUV-III method does not support missing values in the dataset(s). As a result, when multiple datasets are provided, if there are missing values in only one of the datasets, the corresponding rows will be removed in other datasets as well. This function is embedded in most functions of the RUVprps package and can be accessed through the argument named `assess.se.ob`. If this is set to `TRUE`, the `checkSeObj()` function will be applied inside the functions. We recommend applying the `checkSeObj()` once, and then the `assess.se.obj` argument can be set to `FALSE` in all subsequent functions to improve the runtime of the package.

The `orderSeObj()` function can be used to order the samples in a dataset based on specified variables. This can be particularly useful when chronological information is available, allowing users to arrange samples temporally to visualize patterns over time. It can also help in organizing samples by experimental conditions, batch, or other relevant metadata, facilitating downstream analyses and interpretation.

The `groupContinuousVariable()` function bins a continuous variable into discrete groups. This function is usually used in the functions that find NCG and create PRPS data in unsupervised manner.

The `tidyGenes()` function provides several options for organizing feature names (genes) in the *SummarizedExperiment* object. It can remove duplicate gene IDs and retain specific types of genes, such as protein-coding or long non-coding RNAs, if this information is available.

The `createHomogeneousBioGroups()` and `createHomogeneousUVGroups()` functions generate all possible sample groups that are approximately homogeneous with respect to the specified biological and unwanted variables, respectively. If continuous variables are provided, the function splits each into a number of clusters using a clustering method specified by users. Ultimately, all combinations of all clusters are created and each such combination is regarded as a sample group roughly homogeneous with respect to the biological or unwanted variables. For example, if library size and time e.g., years from 2010-2014, are provided as sources of unwanted variation, the *createHomogeneousUVGroups()* will divide the library size into $n$ groups and then the homogeneous sample groups respect to these unwanted variables will be the $5n$ combinations of year and library size groups. These combinations are the sample groups which will be regarded as homogeneous for the known unwanted variation. They are essential for creating PRPS and NCGs, as we will explain in the next section. It should be noted that creating large numbers of sample groups may result in insufficient sample size within each group for creating PRPS or identifying NCGs.

The `obtainMetric()` function can retrieve any specified calculated metric, such as PCA or RLE, for each dataset from the metadata of the *SummarizedExperiment* object.

The `plotMetric()` function can be used to plot any variable, pairs of variables, or heatmap of variables from the *SummarizedExperiment* object. For example, it can be used to visualize the distribution of cancer subtypes across library sizes. The function also calculates appropriate statistics to assess associations between variables.

The `selectGeneSets()` function can retrieve any available gene sets from the gene annotation in the *SummarizedExperiment* object. For example, it can retrieve immune or stromal gene signatures, as well as lists of housekeeping genes.

The `computeSurvival()` function performs survival analysis at either the gene or variable level. For gene-level analyses, expression values of specified genes are stratified into defined groups, and their associations with survival outcomes are assessed. This function generates Kaplan–Meier survival plots.

The `addTcgaClinicInfo()` function provides clinical annotations—including survival outcomes, tumor stage, and related variables for TCGA RNA-seq data.

The `addTcgaBatchInfo()` function adds batch related metadata to individual RNA-seq datasets from TCGA project. The `estimateCMS()` function estimates consensus molecular subtypes (CMS) for colon and rectal adenocarcinoma RNA-seq data. The `estimatePAM50()` function estimates PAM50 molecular subtypes for breast cancer RNA-seq data.

### Gene-level and global assessments of unwanted variation in the Rcount3 database

We used the Rcount3 database to illustrate the importance of assessing unwanted variation at both the gene- and global-levels in RNA-seq data. To quantify library size effects within each dataset, we applied the following procedure: (1) lowly expressed genes were removed; (2) outlier samples based on library size were excluded, these being defined as values exceeding 1.5 $\times$ the interquartile range of library size; (3) counts were normalized using counts per million (CPM) followed by log transformation to mitigate library size variation; (4) principal component analysis (PCA) was performed on the CPM matrix to obtain the first two principal components (PCs); (5) a linear regression model was fitted between the cumulative first two PCs and library size to estimate the corresponding $R^2$ values; and (6) gene-level correlations with library size were computed in the CPM data, with genes showing an absolute correlation $> 0.3$ considered strongly associated. As shown in Supplementary Figure.1.a, in a reasonable number of studies, the first two PCs after CPM normalization fail to effectively capture the influence of library size, whereas gene-level correlation analysis reveals that a substantial number of genes remain highly associated with library size.

To demonstrate the value of gene-level assessment for detecting batch effects, we applied a similar workflow. Steps (1–4) were identical to those described above. (5) Relative log expression (RLE) medians were computed using CPM data, and subjected to unsupervised clustering to estimate batch effects (using the default parameters of the `identifyUnknownUV()` function in RUVprps ); (6) ANOVA was performed for each gene against the inferred batch labels, and genes with an adjusted P value $<$ 0.05 were retained; and (7) vector correlations were computed between the cumulative first two PCs and the inferred batch structure. As shown in Supplementary Figure 1.b, the first two PCs after CPM normalization exhibit limited sensitivity to batch-associated variation, whereas gene-level testing identifies a substantial number of genes that remain strongly associated with estimated batch effects.

Furthermore, the same procedure described above for assessing batch effects was applied using the median of the RLE as a continuous variable rather than as a categorical factor. As shown in Supplementary Figure. 1c, the first two principal components are not strongly associated with the RLE median, whereas a substantial proportion of genes exhibit high correlation with the RLE median across a considerable number of studies.

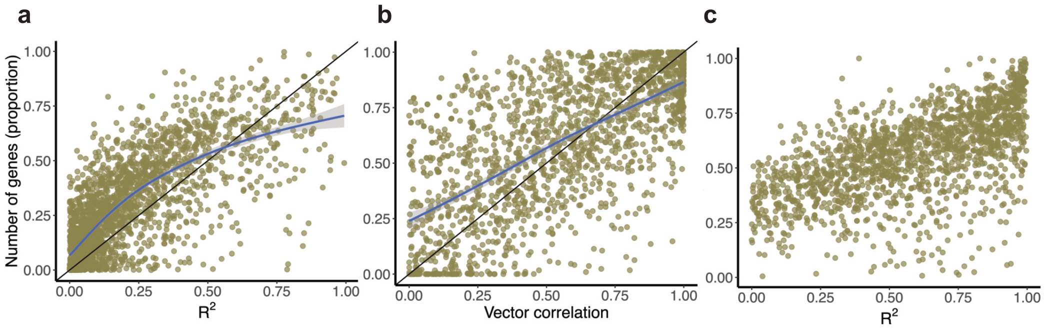


Supplementary Figure 1: Gene-level and global assessments of unwanted variation in Recount3 RNA-seq database. The Recount3 dataset comprises 2,284 RNA-seq studies, each with more than 20 samples. In all scatter plots, each point represents a single study. a) The x-axis shows the $R^2$ from linear regression between the first two PCs of counts-per-million (CPM)–normalized data and library size. The y-axis shows the proportion of genes having Spearman correlation $> 0.3$ with library size in the CPM-normalized data. b) The x-axis shows vector correlations between the first two PCs of CPM-normalized data and estimated batch effects derived using the RLE approach for individual studies. The y-axis shows the proportion of genes having $P < 0.05$ from gene-level ANOVA testing association with the estimated batch effects. c) The x-axis shows the $R^2$ from linear regression between the first two PCs of CPM-normalized data and the median RLE values. The y-axis shows the proportion of genes having Spearman correlation $> 0.3$ with the median RLE in the CPM-normalized data.

### Identification of unwanted variation in large RNA-seq datasets

Before describing how to estimate unwanted variation in large RNA-seq datasets using RUVprps, we first explain the `identifyUnknownUV` function. This function includes an approach argument that can be set to one of three options: `rle`, `pca`, or `sample.scoring`. The `rle` and `pca` approaches can be applied either using all genes or a subset of genes (for example, negative control genes), specified via the `ncg` argument. For the rle approach, either the median or interquartile range (IQR) of the RLE data can be selected for downstream clustering analyses. When gene sets associated with unwanted variation are available, such as gene sets related to tumor purity, housekeeping genes affected by unwanted variation, or cell cycle genes, the `sample.scoring` approach can be used. In cases where known sources of biological variation are available, these variables can be regressed out prior to estimating unwanted variation to improve detection accuracy, using the `regress.out.bio.variables` argument. If the `rle` approach is selected and samples are chronologically ordered (when such information is available), we recommend using change-point detection rather than clustering to estimate unwanted variation. When chronological information is unavailable, or when using other approaches such as `pca` or `sample.scoring`, the `chronological.detection` argument should be set to `FALSE`, and either supervised or unsupervised clustering methods can be applied. The function supports a range of supervised and unsupervised clustering strategies, with multiple tunable parameters to accommodate different study designs and analytical goals. We refer to the RUVprps package documentation for full details of available options and parameter settings.

Based on our previous study, TCGA RNA-seq data are affected by sequencing plate, sequencing time (year), and tissue source site (TSS). Additional sources of unwanted variation may also be present but have not yet been annotated. To identify individual sources of unwanted variation in the TCGA RNA-seq study, we first ordered the data according to each factor and then applied the RLE-based approach with the chronological ordering option to estimate unwanted variation. Cramér's V was used to quantify the association between the estimated unwanted variation and the corresponding technical factors.

For the CCLE RNA-seq dataset, we used the PCA approach implemented in the `identifyUnknownUV()` function to estimate unwanted variation. In this analysis, for individual cancer cell lines, the first three PCs were derived using the top 1,000 stable genes from the *singscore* R package as NCGs. The resulting PCs were clustered in an unsupervised manner using the *mclust* R package. Cramér's V was used to quantify the association between the estimated unwanted variation and known technical factors, including growth medium, collection time (year), strandedness, and DepMap code.

For the GTEx RNA-seq data, we applied approaches similar to those used for TCGA. For each homogeneous tissue, samples were ordered according to potential sources of unwanted variation, including total ischemic time (SMTSISCH), RNA integrity number (RIN; SMRIN), and immune cell heterogeneity. For individual datasets, library size effects were removed using CPM normalization prior to identifying unwanted variation. As with other large RNA-seq projects, additional sources of unwanted variation may exist that are not captured by currently known variables.

For the Treehouse and PCCAT RNA-seq datasets, samples were ordered according to their dataset of origin within each collection, and the RLE-based approach with change-point detection was applied.

To estimate batch effects in the Recount3 database, we used the median RLE values in conjunction with unsupervised clustering. Supplementary Figure 2 summarizes the number of estimated batches across studies.

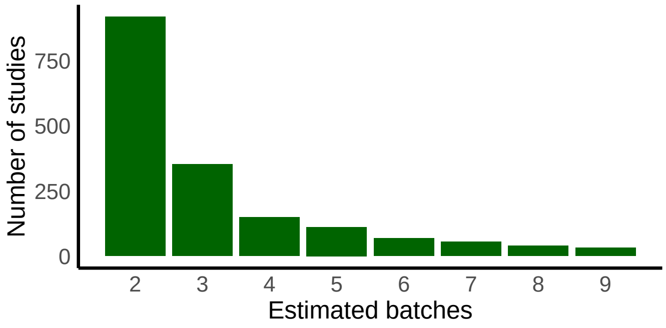


Supplementary Figure 2: Number of batches estimated using RUVprps on the Recount3 studies.

### Data pre-processing for selection of NCG

In data pre-processing for the supervised and unsupervised selection of NCGs, the overarching goal is mitigating unwanted variation to enhance the identification of genes significantly influenced by biological factors, and removing biological variation to find genes impacted by unwanted variation. The input data for the statistical analyses can be one or a combination of the following options:

$log_2$`(raw count data + pseudo-count)`. This is mainly used when raw count data is provided for all NCG selection functions in RUVprps. Notably, this is the only input data required for the `findNcgSupervisedByTwoWayAnova()` function.

*Library size normalization.* Initial library size normalization facilitates the identification of genes significantly affected by biological variation. This can be performed using widely used methods such as CPM as initial normalization, or by regressing out library size with a linear model. Both options are available in all the RUVprps function. Normalization can be applied either within sample groups or across all samples. Note that if CPM or any other library size–normalized data are provided to the functions, the normalization argument must be set to either `FALSE` or `NULL`.

*Regressing out biological and unwanted variation.* Regressing out biological and unwanted variation facilitates the identification of genes that are strongly influenced by unwanted and biological variation, respectively. For example, if tumor purity is considered a biological factor, it can be regressed out to identify genes primarily affected by unwanted variation. This regression analysis can be performed using either a linear model or a linear mixed-effects model options in the functions.

*Regressing out the medians of RLE.* The variation in the medians of RLE data is generally associated with sources of unwanted variation. The NCG functions include an option to calculate and regress out RLE medians from the data prior to identifying genes that are highly variable due to biological factors.

*Key considerations:* (1) Regressing out biological variation that is highly correlated with unwanted variation may inadvertently remove both sources of variation, thereby compromising the identification of genes primarily affected by unwanted variation. The same principle applies when regressing out unwanted variation that is strongly correlated with biological variation. In these scenarios, regression-based adjustment may not be appropriate. (2) To identify biologically affected genes, the order of data processing is as follows: first perform an initial library size normalization, then regress out other sources of unwanted variation, and finally regress out the medians of RLE.

For example, if `apply.log = TRUE, normalization = CPM, regress.out.uv.variables = NULL, regress.out.bio.var` `= NULL`, the preprocessing order is as follows: CPM normalization is applied, followed by a $log_2$ transformation of the CPM-normalized data. This transformed data is then used to identify genes that are strongly influenced by biological variation. In parallel, the $log_2$ transformation of the raw count data (with a pseudo-count) is used to identify genes that are predominantly affected by unwanted variation. Another example: if `apply.log = TRUE, normalization = CPM, regress.out.uv.variables = batch, regress.out.bio.variables = c(subtypes, tumor purity)`, to find genes that are highly affected by biological variation, the function applies CPM normalization followed by $log_2$ transformation first and then regresses out the 'batch' variable from the data. To obtain data to permit finding genes that are highly affected by unwanted variation, the function first applies the $log_2$ transformation and then regresses out both 'subtypes' and 'tumor purity' variables from the data.(3) RUVprps provides the function *regressOutVariables()*, which can be applied to any dataset and the results added to the *SummarizedExperiment* object for use as input in NCG selection. In this case, all other preprocessing options should be set to `NULL` or `FALSE`.

### Identification of NCG in unsupervised way in large RNA-seq datasets

To identify unsupervised NCG for individual TCGA studies, we applied the following procedure. First, for each dataset, lowly expressed genes were removed using the `prepareSeObj()` function, and only protein-coding genes were retained using the `tidyGenes()` function. Second, sources of unwanted variation were estimated using the `rle` approach with change-point detection and unsupervised clustering in the `identifyUnknownUV()` function, as the samples were chronologically ordered. Third, the `findNcgUnSupervised()` function was applied using the `LinearMixedModel` option, fitting the model `~ (1 | Estimated.batches) + Library.size + Tumor.purity` , with `ncg.identification.approach = "LMM.BioUvAdjustment"`. In addition, we evaluated three sets of housekeeping genes: (1) the top 1,000 most stable genes from the *singscore* R package [4]; (2) single-cell housekeeping genes proposed by [5]; and (3) a combined gene set consisting of 1,000 stable genes from *singscore* together with stromal and immune gene signatures from [6]. The latter gene set contains stromal and immune gene signatures to capture variation in tumor purity.

To assess the performance of the NCG gene sets, PCA was performed using only genes in each set separately, and associations between the first 3-5 PCs and multiple sources of unwanted variation including library size, sequencing-plates, time (years) and tumor purity were assessed. In addition, for each individual study, a set of highly variable genes (HVGs) was identified using the `findHVG()` function with the LMM approach. The top 5,000 most stable genes from the singscore R package were also selected. Overlap among the NCGs, HVGs, and stable gene sets was assessed separately for each study. It should be noted that the single-cell housekeeping genes and the stable gene sets derived from the *singscore* R package are not expected to capture variation associated with tumor purity.

The majority of tissues in the GTEx RNA-seq dataset are affected by variation introduced by total ischemic time (SMTSISCH), RNA integrity number (RIN; SMRIN), and immune cell heterogeneity. Variation due to immune cell heterogeneity was estimated using an immune gene signature from [6] and the *singscore* R package. The SMTSISCH and SMRIN variables were categorized into 6 groups, and immune cell scores were grouped into 3 categories using the `cut` function in R. For individual tissues, we applied the following procedure to identify NCGs. First, lowly expressed genes were removed using the `prepareSeObj()` function, and only protein-coding genes were kept using the `tidyGenes()` function. Second, CPM normalization was applied to mitigate library size effects. Third, sources of unwanted variation were estimated using the `rle` approach with change-point detection and unsupervised clustering implemented in the `identifyUnknownUV()` function, with samples ordered according to one of the unwanted variables at a time. Fourth, the `findNcgUnSupervised()` function was applied using the `LinearMixedModel` option, fitting the model `~ (1 | Estimated.batches)` and setting `ncg.identification.approach = "LMM.BioUvAdjustment"`. Finally, the top 5% of genes were selected as NCGs. The same sets of publicly available housekeeping genes used in the TCGA analysis were applied to the GTEx data. Furthermore, the procedures adopted to assess different NCG sets in TCGA were similarly applied to evaluate NCG sets in the GTEx dataset.

### Unsupervised identification of NCG with LMM

RUVprps adopts a linear mixed model from the variancePartition R package [7] to find NCGs in both supervised and unsupervised way by RUVprps. The *variancePartition* framework decomposes gene expression variance into components explained by multiple biological and technical factors by fitting a linear mixed-effects model for each gene. Variance components are estimated genome-wide and summarized to quantify the relative contribution of biological signals and unwanted variation across genes. The variancePartition R package models categorical variables a random effect. (https://www.bioconductor.org/packages/release/bic In this study, we modeled all categorical variables as random effects and all continuous variables as a fixed effects. RUVprps provides different strategies to use the LMM to find genes in unsupervised manners (Supplementary Figure.3). The first strategy fits a LMM using all estimated sources of unwanted variation. Genes for which a large proportion of expression variance is explained by the unwanted factors are selected as NCGs (Supplementary Figure 3a). The second strategy proceeds in four steps. First, all estimated unwanted factors are regressed out from the expression data. Second, PCA is applied to the residuals to obtain the first few PCs. Third, an LMM is fitted including both the estimated unwanted factors and the selected PCs. Finally, genes are selected as NCGs if they collectively exhibit high variance explained by the unwanted factors while showing minimal variance explained by the PCs (Supplementary Figure. 3b). The third strategy comprises a multi-step procedure. First, all estimated sources of unwanted variation are regressed out from the expression data. Second, PCA is applied to the residuals to extract the first few PCs, which are treated as candidate sources of biological variation. Third, a LMM is fitted that includes both the estimated unwanted variation factors and the selected PCs. Fourth, a subset of genes exhibiting high variance explained by unwanted factors and minimal variance explained by the PCs is selected. Fifth, PCA is then performed using this subset of genes to obtain PCs capturing unwanted variation. Sixth, an LMM is fitted incorporating both sets of PCs derived from step 2 (biological PCs) and step 5 (unwanted variation PCs). Finally, a refined subset of genes is selected that collectively shows high variance explained by unwanted variation while displaying minimal variance explained by biological PCs( Supplementary Figure. 3c).

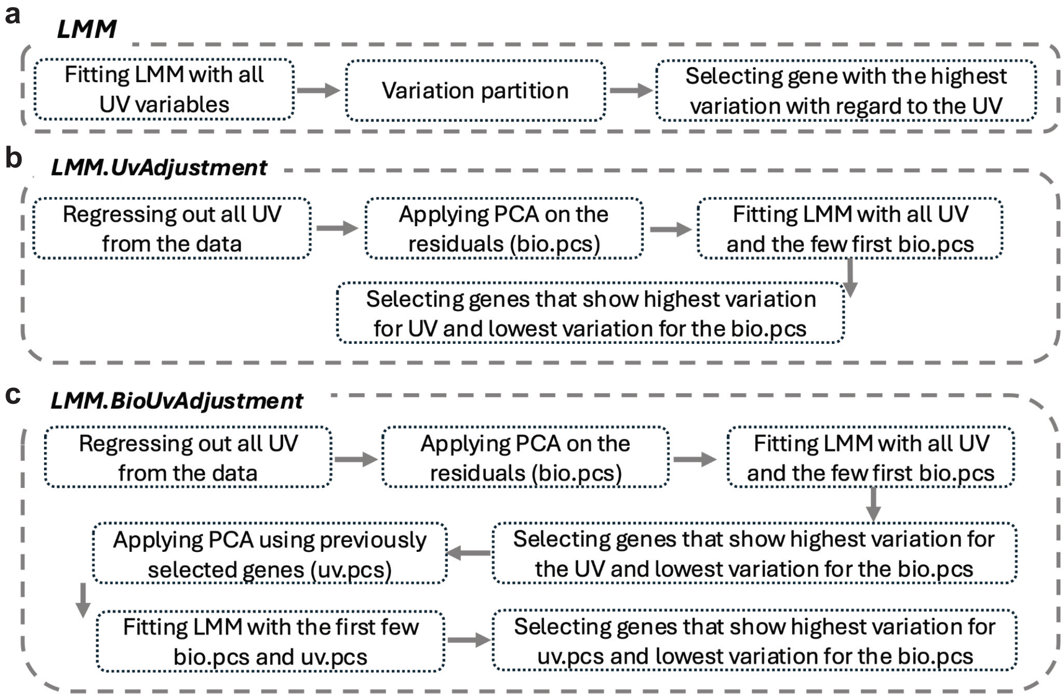


Supplementary Figure 3: Different approaches to find NCGs in unsupervised way using the `findNcgUnSupervisedByLinearMixedModel()` function. The default, all categorical variables are modeled as random effects and continuous variables as fixed effects. a) b) and c)show the major steps involved in the selection of NCGs using the function `findNcgUnSupervisedByLinearMixedModel()` when the approach is set to `LMM`, `LMM.UvAdjustment`, and `LMM.BioUvAdjustment`, respectively.

### Selection procedures for NCG

The supervised and unsupervised functions for identifying NCG generate gene-level statistics, including correlation coefficients, F-statistics, effect size, MAD, ... for individual or a combination of variables. Depending on the function used and the types of variables specified, we propose different strategies to summarize these statistics in order to select the most suitable set of NCG. *Rank-based selection for individual variables:* The options below are based on the ranking of gene-level statistics obtained for each individual variable. To use one of the option below, the `use.rank` argument in the function must be set to `TRUE`. It should be noted for functions that uses ANOVA, there is a option to use either the F-statistic or effect size. “product”, “average” or “sum” of ranks: In this procedure, statistical results obtained for unwanted and biological variable are ranked differently such that a lower rank indicates a stronger influence of unwanted variation on gene expression, while a higher rank reflects a stronger influence of biological variation. These ranks are then combined using either the product, average, or sum of ranks. Finally a number specified by users of genes with the lowest combined ranks are selected as NCG. *none.overlap approach:* First, similar to the approach described above, the statistical results obtained for unwanted and biological variables are ranked separately. Next, the top percentage of genes with the highest ranks for biological variation are selected. In parallel, the top percentage of genes with the lowest ranks for unwanted variation are also selected. Finally, the genes identified in the second step (highly associated with biological variation) are excluded from those identified in the third step (highly associated with unwanted variation). The remaining genes constitute the set of NCG. It should be noted that the final number of NCG cannot be predetermined in this approach as the degree of non-overlap between the two sets may vary.

*"auto" approach*: This approach is similar to the "none.overlap" method described above. However, it incorporates an automated search procedure that adjusts the initial values for selection top highly affected genes by biological, unwanted variation or both to approximate the number of NCG specified by the user. If the initial selection yields more or fewer NCG than requested, the function iteratively modifies the values of these parameters to converge on the target number of NCG. In exact number of NCG can be determined by users. If the selection method is set to the "auto" approach and the initial values of "top.rank.uv.genes" and "top.rank.bio.genes " result in a number of NCG that is either greater or fewer than the user-specified target, the function will initiate an automated search procedure. The examples below illustrate how to specify the function arguments.

*Ratio-based selection:* In this approach, the summarized statistics for biological and unwanted variables are first aggregated separately. To identify genes predominantly influenced by biological variation, the ratio of the aggregated biological values to the aggregated unwanted values is computed. Higher ratios correspond to genes that are primarily affected by biological variation, while the same principle can be applied to detect genes strongly impacted by unwanted variation. Users may specify a threshold percentile of the summarized statistics to be considered prior to ratio calculation, thereby reducing bias from extremely low values. Following ratio computation, users can select the top n genes as negative control genes (NCGs).

### Supervised PRPS functions

RUVprps offers two functions to create different sets of PRPS for categorical and continuous sources of unwanted variation. For categorical variables, such as platform effects, the function `createPrPsSupervisedForCategoricalUV()` first identifies all possible homogeneous biological groups across batches. It then selects biological groups that contain at least three samples in each of two or more batches. Within each batch, the biologically similar samples are averaged, and the resulting PS are subsequently paired across batches to form sets of PRPS.

RUVprps provides the function `createPrPsSupervisedForContinuousUV()` to create PRPS for continuous sources of unwanted variation. The function first identifies all possible homogeneous biological groups in the data. Within each biological group, samples are ordered according to the continuous source of unwanted variation. The top three and the bottom three samples, with the highest and lowest values of the unwanted variation, respectively, are averaged to create two PS, which are then paired to form a PRPS set.

Both functions provide options to categorize continuous sources of biological and unwanted variation. When more than one source of unwanted variation is provided, the function also allow users to account for other unwanted variables while generating PRPS for a specific source of unwanted variation. Furthermore, the number of samples used to create each PS can be specified, with a minimum of two samples. The functions additionally offer the option to either apply a log transformation to the data before creating PS or to create PS first and then apply the log transformation. Finally, both functions can generate a PRPS map to visualize how well the PRPS are distributed and how effectively they capture unwanted variation in the data.

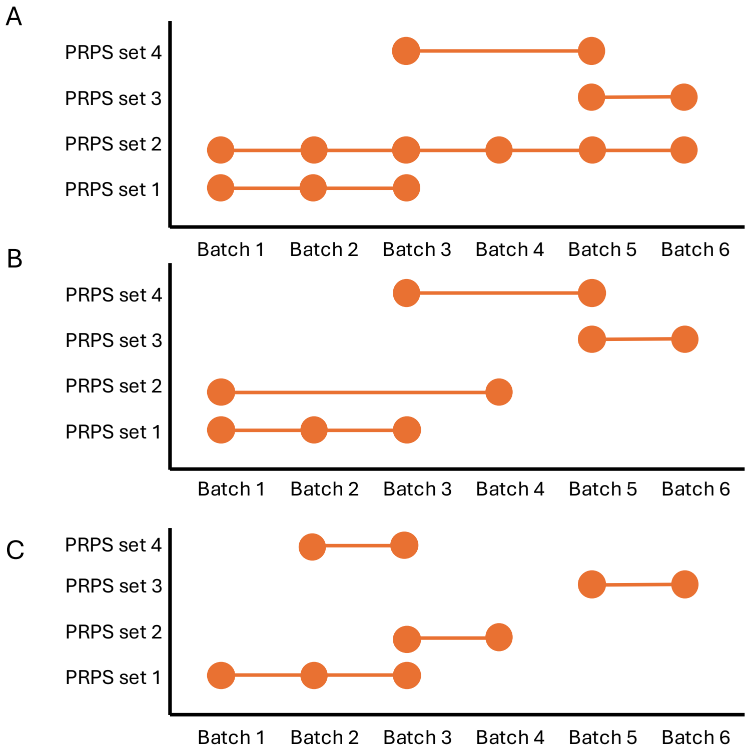


Supplementary Figure 4: Illustrative examples of connectedness between the PRPS sets. In all examples, there are six batches and four PRPS sets. A) The second PRPS set covers all six batches. In this case, there is no need to check for connectedness between the PRPS sets, as full coverage is already achieved. B) The first PRPS set covers batches 1 to 3, the second set covers batches 1 and 4, the third set covers batches 5 and 6, and the fourth set covers batches 3 and 5. Batch 4 is connected to batches 1 to 3 via the second PRPS set. Batches 5 and 6 are connected to batches 1 to 4 through the fourth PRPS set. Therefore, complete connectedness is achieved across all batches. C) Batches 1 to 4 are connected through the first two PRPS sets. However, batches 5 and 6 are not connected to these batches. As a result, there is no overall connectedness among all six batches. The batch gap between 1–4 and 5–6 is not addressed effectively by the PRPS strategy illustrated.

### Definition of control samples sets and how to label them to create M matrix

Individual groups of samples that are considered as control samples should have the same labels regardless of whether they are TR, PR and PRPS. We will refer to each group of control samples as a 'set'. We will explain this in greater detail with examples below.

*Using only TR:* Supposed we have an RNA-seq data with 100 samples and they are labled as sample 1, sample 2, ..., and sample 100. Assume we have two sets of TR as follows: The first set includes samples 10, 50, and 99, while the second set includes samples 30, 40, and 67. We can label all samples from the first set as 'sample.TR.set1' (it can be any name) and all samples from the second set as 'sample.TR.set2'. The vector of sample labels for creating the M matrix should now have 96 unique labels. This results in a replicate matrix M with 100 rows (all samples) and 96 columns (unique samples).

*Using only PRPS:* Consider an RNA-seq dataset with 100 samples. Suppose we create a pseudo-sample (PS) by averaging samples 3, 20, and 22. Then, we create another PS by averaging samples 11, 78, and 61. These two PSs (extra columns in the data) can be labeled as "ps.set1". If we create two additional PSs as part of another PRPS set, their labels can be "ps.set2". The vector of sample labels for creating the M matrix should now have 102 unique labels. Therefore, the dimension of the M matrix will be 104 rows (all 100 genuine samples and 4 PSs) and 102 columns (all genuine samples and PRPS sets).

*Using Both TR and PRPS:* In the examples above, if we use both TR and PRPS, the vector of sample labels for creating the M matrix will have 98 unique labels. Accordingly, the dimension of the M matrix will be 104 rows (assays) and 98 columns (unique samples).

*Note:* To perform a sanity check on the M matrix, the sum of each row should be equal to 1. The sum of the columns should be 1 for unique (non-replicated) samples and greater than 1 for replicated samples. If the sum of any row deviates from 1 or if all column sums equal 1, it indicates that the M matrix has not been generated correctly.

### Details of the *RUVIIIprps()* function

Here, we explain several major arguments of the `RUVIIIprps()` function. We refer to Supplementary Figure 5 for a schematic overview of the metadata structure used to store PRPS and NCG sets identified by both supervised and unsupervised approaches.

`prps.type`. This argument specifies which type of PRPS data, created by PRPS functions, should be used for RUV-III normalization. The available options are `supervised`, `unsupervised`, or `both`. For example, one may create supervised PRPS for one source of unwanted variation and unsupervised PRPS for another, and then apply both during RUV-III normalization. All PRPS data must be stored in the metadata of the *SummarizedExperiment* object in order to be accessed by the `RUVIIIprps()` function

`prps.group.names`. This argument specifies which PRPS group(s) should be used for RUV-III normalization. On can create multiple groups of PRPS data using different parameters and then assess their performance separately or together.

`prps.set.names`. This argument specifies which PRPS set(s) should be used for RUV-III normalization. For example, one may generate PRPS data for library size and tumor purity. This argument allows the user to select one or all of these sets. The default is set to `all`, meaning that all available PRPS sets will be selected.

`technical.replicates`. This option allows users to use TRs if any are available. TRs can be either used along with PRPS data or alone. If one wants to use only TRs, the `prps.type` option must be set to `none`.

`ncg.type`. This argument specifies which type of NCG , created by NCG functions, should be used for RUV-III normalization. The available options are `supervised`, `unsupervised`, or `both`. All NCG must be stored in the metadata of the *SummarizedExperiment* object in order to be accessed by the `RUVIIIprps()` function.

`ncg.group.names`. This argument specifies which NCG group(s) should be used for RUV-III normalization. On can create multiple groups of NCG using different parameters and then assess their performance separately or together.

`ncg.set.names`. This argument specifies which NCG set(s) should be used for RUV-III normalization. The default is set to `all`, meaning that all available NCG sets will be selected.

`K`. This argument specifies the value(s) of $k$, which represents the number of unwanted variation factors to estimate and remove from the data. A separately normalized dataset is generated for each $k$ value and stored in the *SummarizedExperiment* object.

`data.to.log`. This option allows users to apply a log transformation on either the specified data, the control samples, or both. If the raw count data is used to create the PRPS data and `apply.log = TRUE` in the PRPS functions, then PRPS data is already log-transformed, and thus no further log transformation is needed. In this case, `data.to.log` should be set to `assay`. If the PRPS data is not log-transformed when they are created (`apply.log = FALSE` in the PRPS functions), but the specified assay for RUV-III is log-transformed, this option should be set to `data.to.log = prps`.
`return.w`. When it is set to 'TRUE', this returns the $W$ matrix, which contains the sample-wise estimates of unwanted variation by RUV-III. We highly recommend exploring and assessing the $W$ matrix as explained in this study.

### Gene expression data type as an input for RUVprps

The RUVprps package can accommodate various types of transformed and normalized RNA-seq data. While we highly recommend using raw count data as input, this may not always be available. For instance, when using publicly available datasets, the only available format may be CPM or FPKM. Below we will explain what kind of data type can be used for major steps in the RUVprps package.
*Preparation of the SummarizedExperiment object.* The `prepareSeObj()` function requires raw count data without any prior transformation in order to identify lowly expressed genes and calculate library size. If raw count data is unavailable, the removal of lowly expressed genes and library size calculation should be performed outside the RUVprps package. In this case, the processed data should then be provided to the `prepareSeObj()` function and the corresponding arguments, including `remove.lowly.expressed.genes` and `calculate.library.size`, should be set to `FALSE`. However, estimation of tumor purity can be performed using any type of data but it is important to note that lowly expressed genes should be removed previously. Indeed both the ESTIMATE and singscore methods employed to estimating tumor purity, are rank-based approaches for gene set enrichment analysis.
*Identification of unknown sources of unwanted variation:*The `identifyUnknownUV()` function can accommodate any type of data as input. If the data is not logarithmically transformed, the `apply.log` argument should be set to `TRUE`. Note that, there is no need to logarithmically transform the data, if the `approach = sample.scoring` is set in the function.
*Assessment of variation:* The `assessVariation()` function performs all global and gene-level assessments on all specified data, regardless of how they were transformed or normalized. All specified data must have been logarithmically transformed prior to applying the function. If any data is not transformed, the `applyLog()` function can be used to apply a log-transformation, with or without a pseudo-count.
`RUV-III normalization`:The RUV-III normalization steps, including the selection of negative control genes, PRPS creation, and RUV-III normalization, can be performed on raw count data (if available) or normalized data e.g. CPM, FPKM, .... However the data must be logarithmically transformed prior to these steps. Note that, the output of the RUV-III method is log-transformed data.
*Evaluation of normalization:* Similar to the assessment of variation step, the `normalizationAssessment()` function applies all global and gene-level assessments on all specified assays, regardless of how they were transformed or normalized. All specified assays must have been logarithmically transformed before applying the function. If any assay is not transformed, the `applyLog()` function can be used to apply a log-transformation, with or without a pseudo-count.

Examples:

*Example 1:* Consider a situation where the available datasets are raw count and FPKM data followed by upper-quartile normalization (FPKM.UQ) . Neither of the datasets is logarithmically transformed. Suppose that the goal is to normalize the raw count data using the RUV-III method and compare it with FPKM.UQ normalized data. The *SummarizedExperiment* object can be created by including both datasets. Two options are available for applying all downstream functions: (1) The `applyLog` function can be used to create logarithmically transformed data of both the raw count and FPKM.UQ within the object. Then, all downstream functions can be applied to both assays (log-transformed raw counts and log-transformed FPKM.UQ data) with the `applyLog` argument of the functions set to `FALSE`. (2) apply all functions except those involved in the normalization assessment step to the raw count assay and FPKM.UQ assay with the `applyLog` argument set to `TRUE`. After applying RUV-III normalization, both the raw count and FPKM.UQ data should be logarithmically transformed to allow for normalization assessments on the two assays and the RUV-III normalized data.

*Example 2:* Consider a situation where both raw count data and log-transformed CPM data are available. Suppose the aim is to re-normalize the raw count data using RUV-III and compare it with the log-transformed CPM data. The *SummarizedExperiment* object can be created by including both datasets. The `applyLog()` function can then be used to apply a log-transformation to the raw count data within the object. Subsequently, all downstream functions can be applied to both assays, the log-transformed raw counts and the log-transformed CPM data, with the `applyLog()` argument of the functions set to `FALSE`.

*Notes:* Some normalization approaches can remove biological signals from data. In such cases, the RUV-III method might not be able to recover those signals if only the normalized data is provided. (2) One may prefer to use some transformed data such as FPKM over raw count data to estimate tumor purity [8].

### Normalization assessment metrics

*RLE scores:* An ideal RLE plot should have all the medians close to zero and IQRs similar to each other. Two separate scores are calculated for the medians and IQR of the RLE data. To obtain a score for the RLE medians, we first find the largest absolute value $S$ of the RLE medians across samples in all assays:

$$S = \max_{i,j} |R_{ij}|$$

where $R_{ij}$ represents the RLE median for assay $i$ and sample $j$, and $\max_{i,j}$ denotes the maximum value of the absolute RLE medians across all assays $i$ and samples $j$. Then, this value is multiplied by the number of samples $n$ in the data. This is the largest possible deviation of RLE medians that a normalized dataset can have. Finally, the sum of the absolute RLE medians across samples in each normalized dataset is divided by this largest value. The result is subtracted from 1, ensuring that higher scores reflect better performance of the method.

To calculate a score for IQRs, let {IQR} denote the collection of $n$ IQRs. Our goal is to ensure the these IQRs are uniform, i.e. their range to be small. Then, as above, we need to standardize to have no units and subtract from 1. Let $Q_{10}$, $Q_{50}$ and

$Q_{90}$ be the 10th, 50th and 90th deciles of {IQR} (with modification for smaller numbers $n$). We propose:

$$\text{RLE IQR score} = 1 - \frac{Q_{90} - Q_{10}}{Q_{50}}$$

*Association between RLE medians or IQRs and variables:* We have previously demonstrated that the medians of RLE values exhibit a strong correlation with library size in the raw counts of the TCGA RNA-seq dataset [1]. Once the impact of library size was effectively removed, the medians of RLE values showed a high correlation with tumor purity estimates. If tumor purity is regarded as a source of biological variation, this high correlation can serve as evidence of the effectiveness of a normalization method. Then, Spearman and Pearson correlation analyses are used to summarize the relationship between the RLE medians or IQRs and continuous variables. The directionality of correlation is disregarded, and so the absolute correlation coefficients are computed. The coefficients will be subtracted from 1 for unwanted variables. Then, for both unwanted and biological variables, higher values indicate better performance. The Kruskal-Wallis rank sum test is used to assess the association between RLE medians or IQRs and categorical variables. The calculated p-values are subtracted from 1 for biological variables. Then, for both unwanted and biological variables, higher values indicate better performance.

*Vector correlation and regression analysis scores:* To summarize the results of the vector correlation and regression analyses between $n$ PCs and a categorical or continuous variable, we calculate the average of the first $n$ vector correlations or $R^2$ values, respectively. The average values are subtracted from 1 for unwanted variables. Therefore, for both unwanted and biological variables, higher values indicate better performance.

*Silhouette and ARI scores:* The Silhouette coefficients and ARIs range from -1 to 1. To standardize the range between 0 and 1, we add 1 to each coefficient and then divide the result by 2. The values are subtracted from 1 for unwanted variables. For both unwanted and biological variables, higher values indicate better performance.

*kBET scores:* The kBET method is only used to assess the removal of batch effects in RUVprps. The range of kBET values across samples is from 0 to 1. To calculate an overall kBET score, these values are averaged. The higher values indicate better performance of normalization in terms of mixing batches.

*LISI scores:* In RUVprps, the LISI method is used to assess normalization performance in terms of both preserving biological variation and removing unwanted variation. In general, the minimum batch LISI score is 1, indicating that all neighbors originate from the same group or label. The maximum LISI score equals the total number of groups or labels in the data, reflecting perfectly even mixing. To standardize batch LISI scores, min–max normalization is applied:

$$\text{LISI score} = \frac{1}{n}\sum_{i=1}^{n} \frac{\text{LISI}_i - \min(\text{LISI})}{\max(\text{LISI}) - \min(\text{LISI})}$$

This rescales the scores to the range 0–1, where a value of 0 corresponds to the lowest observed LISI (worst mixing) and a value of 1 corresponds to the highest observed LISI (best mixing). The normalized values are then averaged to obtain a final batch LISI score. For biological variation, the final sample-level LISI score is subtracted from 1, so that higher values consistently indicate better performance for both unwanted and biological variables.

*Differential gene expression scores:* In RUVprps, DGE analysis is primarily used to evaluate normalization performance in terms of removing unwanted variation. In this setting, DGE is performed between groups defined by unwanted variables. If normalization is effective, we expect no or very few differentially expressed genes (DEGs) between these groups. To quantify this, two complementary summaries are used; 1) Proportion of DEGs estimated using Storey and Tibshirani's method, as implemented in the q-value R/Bioconductor package (version 2.32.0). A smaller proportion of DEGs indicates better removal of unwanted variation (since no true biological DEGs are expected in this context). This value is subtracted from 1 to be consistent with the interpretation of other metrics, so that higher values indicate better performance. 2) Uniformity of p-values – assessed using the Kolmogorov–Smirnov test via the `ks.test()` function in R. The test compares the empirical cumulative distribution function (ECDF) of the observed p-values to the cumulative distribution function (CDF) of the uniform distribution (punif). The Kolmogorov–Smirnov statistic D measures the maximum absolute difference between the two distributions. A smaller D value indicates that the p-values more closely follow the expected uniform distribution. This value is subtracted from 1 to be consistent with the interpretation of other metrics, so that higher values indicate better performance.

In RUVprps, DGE analysis is also used to assess normalization performance in terms of preserving biological variation. However, interpreting the number of DEGs requires biological knowledge. If few or no DEGs are expected between two biological groups but normalization reveals a large number, this indicates a shortcoming of that normalization. Conversely, if a substantial number of DEGs are expected but normalization reveals few or none, this also indicates a shortcoming of the normalization.

*Gene-level correlation and ANOVA scores:* RUVprps uses three different measurements to summarize the outputs of the gene-level correlation and ANOVA with continuous and categorical variables. First, cutoffs will be specified based on absolute correlation coefficients and F-statistics to select genes with values lower than the specified cutoffs. More genes indicates a better performance for unwanted variables, yet interpreting biological variables' true values proves challenging as they are unknown. Second, we estimate the the proportion of null p-values obtained from gene-level correlation and ANOVA, separately. Higher values indicate the better performance for unwanted variables. Third, the uniformity of p-values $>$ .05 is assessed. This is obtained as explained in DGE section.

*Gene-gene partial correlation scores:* To summarize the gene-gene partial correlation results, we calculate the proportion of gene-gene pairs whose differences between their ordinary and partial correlations are less than a cut off. This proportion will be subtracted from 1 for continuous biological variables. For both unwanted and biological variables, higher values indicate better performance.

*Overall score:* Overall performance scores are computed to rank the effectiveness of normalizations. The metric scores are averaged separately across biological and unwanted variables. Then, the overall score, $Score_{overall,n}$ for each normalization $n$ is calculated by taking the weighted mean of the batch removal score, $Score_{batch,n}$, and the bio-conservation score, $Score_{bio,n}$, following the equation:

$$Score_{\text{overall},n} = 0.6 \times Score_{\text{bio},n} + 0.4 \times Score_{\text{batch},n}$$

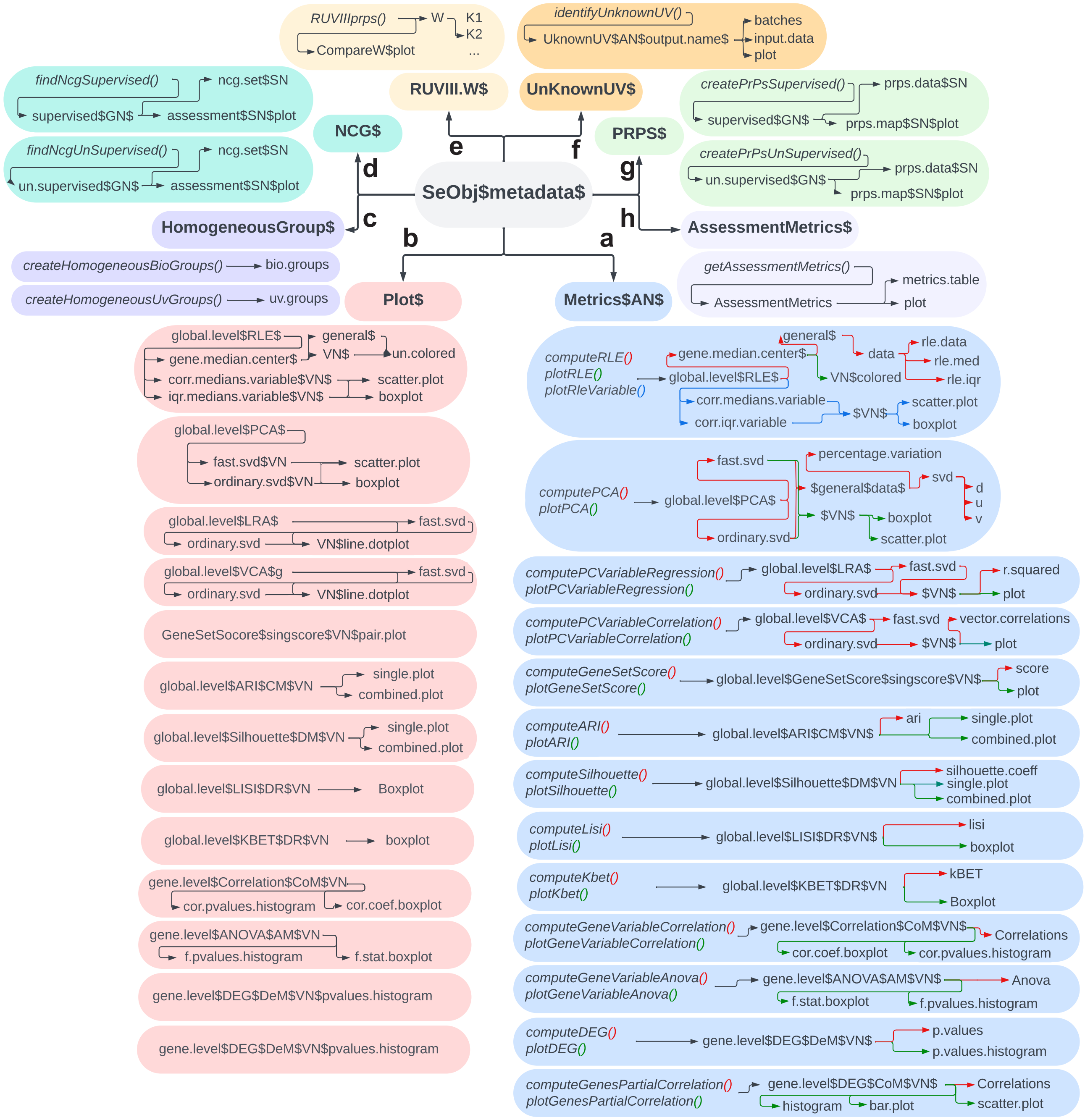


Supplementary Figure 5: An overview of the metadata structure of a *SummarizedExperiment* in RUVprps. a) Illustrates the organization and location of the outputs derived from various assessment functions within the metadata. For each dataset, both numerical results and graphical summaries of all assessments are stored. b ) Indicates that, when multiple data sets (assays) are provided during the assessment steps, the corresponding plot outputs are combined and stored in the metadata. c) Indicates the location of outputs associated with the creation of homogeneous sample groups, considering both unwanted and biological variations. d) Highlights the storage of both supervised and unsupervised identification of NCGS. e) Shows where the estimated unwanted variation factors, W is stored in the RUV-III method. f) Outlines the storage of any estimated sources of unknown unwanted variation within the metadata. g) Analogous to d, shows the storage of outputs related to PRPS data. h) illustrates the storage location for the comprehensive numerical assessment table of normalizations.

### Data pre-processing in RUVprps

The RUVprps package involves three major data pre-processing steps including library size normalization, regression of either biological or unwanted variables, and a combination of both. We will explain the pre-processing steps in several major procedures of the RUVprps package.

*Identification of unknown sources of unwanted variation.* The `identifyUnknownUV()` function has two arguments for preprocessing: `regress.out.bio.variables` and `regress.out.bio.gene.sets` that help find unknown sources of unwanted variation by regressing out known biological variation from the data. The first option regresses out any specified known biological variables from the data and then estimates unwanted variation from the residuals. The second option performs gene-set scoring analysis of any given biological gene signatures. It then regresses out the sample-level scores from the data and finally uses the residuals to identify unknown sources of unwanted variation. We recommend regressing out only variables or gene sets that do not have association with unwanted variation. If there is uncertainty with a variable or gene-set that might have a strong association with unwanted variation, we suggest not to include it.

*Creation of PRPS in an unsupervised manner.* The `createPrPsUnSupervised()` function has three arguments for preprocessing: the `normalization`, `apply.cosine.norm`, and `regress.out.variables` that facilitate the identification of biologically similar samples across batches. The first option applies library size normalizations such as CPM or TMM to mitigate the impact of library size. The cosine normalization helps deal with situations where different batches are normalized differently. To do so, this normalization removes scaling differences between expression values from different batches. The *regress.out.variables* option will regress out any specified variables from the data and use the residuals to find similar samples. In some situations, a combination of these pre-processing steps may be desirable. For example, when cancer RNA-seq data is affected by both library size and tumor purity and these are considered as unwanted variation. One can specify library normalization to remove the library size variation and then regress out tumor purity from the data before identifying biologically similar samples.

*Identification of negative control genes.* Regressing out biological and unwanted variables can help identify negative control genes in either the supervised or unsupervised approaches.

#### Overview of the RUV-III normalization of RNA-seq datasets

To demonstrate the performance of the RUVprps package, we present several different procedures, each with distinct objectives. Each procedure consists of four stages. The first two stages involve data pre-processing and the estimation of biological and unwanted variation. If this information is already available, these stages become unnecessary. The third stage involves assessing variation in the data, and the final stage applies RUV-III normalization and evaluates performance. In RUV-III normalization, we define different scenarios to reflect a range of real-world challenges. Fig.1 shows overall pipeline of RUV-III with PRPS normalization. Some functions have a parallelization option to speed up the underlying calculations. In such cases, the number of cores used for the function is usually set to the number of available cores minus one.

### RUV-III normalization of the TCGA BRCA RNA-seq data

In this procedure our goal is to remove the variation associated with tumor purity alongside other sources of unwanted variations. This variation will then be included as part of the known unwanted variation in the RUV-III with PRPS normalization process. We selected the TCGA breast invasive carcinoma (BRCA) RNA-seq data for this purpose, as it is significantly influenced by purity variation and batches [1]. In this procedure, we will focus on only cancer tissue samples.

##### Required R packages and installations

The R packages below are required to be installed for this procedure.
The TCGAbiolinks R/Bioconductor package (version 2.34) will be used to download the TCGA RNA-seq data from Genomic Data Commons (GDC) data repository.

```
if (!require("BiocManager", quietly = TRUE))
    install.packages("BiocManager")
BiocManager::install("TCGAbiolinks")
```

##### Stage 1: Data preparation and processing

##### Step 1: Downloading the data

The most recent TCGA BRCA RNA-seq data can be downloaded from the GDC data portal using the TCGAbiolinks R/Bioconductor package. The required files will be downloaded to the current home directory.

```
tcga.brca.rnaseq.data <- TCGAbiolinks::GDCquery(
    project = 'TCGA-BRCA',
    data.category = 'Transcriptome Profiling',
    data.type = 'Gene Expression Quantification')
TCGAbiolinks::GDCdownload(tcga.brca.rnaseq.data)
brca.se.obj <- TCGAbiolinks::GDCprepare(query = tcga.brca.rnaseq.data)
brca.se.obj <- brca.se.obj[ , brca.se.obj$tissue_type == 'Tumor']
```

The `brca.se.obj` is a *RangedSummarizedExperiment* object with 60,660 Ensemble transcripts and 1,231 RNA-seq samples. The object contains 6 datasets (assays) including `unstranded`, `stranded_first`, `stranded_second`, `tpm_unstrand`, `fpkm_unstrand` and `fpkm_uq_unstrand`. This object is deposited on the Zenodo website for reproducibility [i will add the link once the MS is finalized]. The `unstranded`, `stranded_first` and `stranded_second` are raw count data without any transformation. We will keep only the `unstranded` assay as raw count data for downstream analysis. The reason for this is that TCGA normalizations, including TPM, FPKM, and FPKM.UQ, were performed on the unstranded data. Note that the normal tissues (113 samples) will be removed from the data before down-stream analysis.

```
# removing some datasets (assays)
brca.se.obj <- removeAssays(
    se.obj = brca.se.obj,
    assays.to.remove = c('stranded_first', 'stranded_second'))
# changing the datasets names
brca.se.obj <- renameAssays(
    se.obj = brca.se.obj,
```

```
    new.names = c('RawCount', 'TPM', 'FPKM', 'FPKM.UQ'))
```

**Step 2: Selecting the gene types of interest (optional)**

The `gene_type` column in the gene annotation (rowData) of the *RangedSummarizedExperiment* object classifies Ensembl transcripts into various gene type categories, such as *protein_ coding*, *lncRNA* (long non-coding RNA), among others. For this analysis, only unique Ensembl transcripts annotated as `protein_coding` will be retained using the `tidyGenes()` function. Additionally, the row names of the *RangedSummarizedExperiment* object will be updated to display gene symbols instead of Ensembl transcript IDs for down-stream analysis.

```
# obtaining gene annottaion
brca.se.obj <- tidyGenes(
    se.obj = brca.se.obj,
    keep.gene.type = 'protein_coding',
    gene.type.col.name = 'gene_type',
    remove.duplicates.ids = TRUE,
    ids.col.name = 'gene_name',
    change.row.names = TRUE,
    new.row.names = 'gene_name')
```

Then a total of 19,934 Ensemble transcripts were retained. It should be noted that any gene type groups of interest can be selected for RUV-III normalization.

**Step 3: Adding batch details**

TCGA RNA-seq samples were collected from various tissue source sites (TSS), shipped to different genomics profiling centers over time, and profiled using 96-sequencing plates. These are known sources of batch information that can impact down-stream analysis [1]. The `addTcgaBatchInfo()` function will be employed to annotate the TCGA BRCA RNA-seq data with all available batch information. This function retrieves batch annotations from the file "41587_2022_1440_MOESM3_ESM.xlsx", available from our previous study [1]. Finally, the `orderSeObj()` function will be applied to order the samples chronologically based on the batch information, in preparation for down-stream analysis.

```
# adding the TCGA RNA-seq batch information
brca.se.obj <- addTcgaBatchInfo(se.obj = brca.se.obj)
brca.se.obj$Years <- as.factor(x = brca.se.obj$Years)
brca.se.obj <- orderSeObj(se.obj = brca.se.obj, factors.to.order = c('Years', 'Plates'))
```

The TCGA BRCA RNA-seq data now comprises 1,083 cancer tissues samples that were collected from 40 tissue sources sites (TSS), distributed across 37 sequencing plates, and profiled over 5 years from 2010 to 2014. The samples collected in 2010 and 2011 were profiled using one flow cell chemistry, and the remaining samples were profiled using a different flow cell chemistry [1]. Note that the TCGA sample barcodes contain several details such as TSS, plates, genomics profiling center, etc . Click here for more details.

**Step 4: Preparing the SummarizedExperiment object**

The `prepareSeObj()` function will be utilized to perform several preprocessing steps, including the identification and removal of lowly expressed genes, calculation of library sizes, estimation of tumor purity, integration of multiple publicly available housekeeping gene lists, and stromal and immune gene signatures. For these tasks, the `RawCount` data will be used for library size calculation, while `FPKM.UQ` will be employed for tumor purity estimation. If a gene annotation file is provided, it must contain at least one column with a name that includes `hgnc_symbol`, `entrezgene_id`, or `ensembl_gene_id` to enable the mapping of gene details and incorporation of housekeeping gene lists. Furthermore, the row names of the *SummarizedExperiment* object must correspond to the identifiers in the specified column. In this analysis, the gene annotation associated with the *brca.se.obj* object includes a column named `gene_name`; therefore, the function will generate a new column labeled `hgnc_symbol` accordingly.

```
brca.se.obj <- prepareSeObj(
    data = brca.se.obj,
    raw.count.assay.name = 'RawCount',
    remove.lowly.expressed.genes = TRUE,
    assay.name.to.estimate.purity = 'FPKM.UQ',
    estimate.tumor.purity = 'both',
    scale.singscore.values = TRUE,
    calculate.library.size = TRUE,
    add.immun.stroma.genes = TRUE,
    add.gene.details = TRUE,
    add.housekeeping.genes = TRUE,
    column.name = 'gene_name',
    gene.group = 'hgnc_symbol')
```

15,304 of 19,934 Ensemble transcripts with expression level of CPM cutoff $\geq 0.18$ in at least 598 samples are kept as highly expressed genes (the default parameters of the function). Tumor purity has been estimated using both the ESTIMATE and singscore methods [2, 6]. The tumor purity estimated by the singscore method will be re-scaled based on the tumor purity estimated by the ESTIMATE method. This adjustment is necessary because the scores from the singscore method exceed 1, whereas the ESTIMATE method provides tumor purity estimates within a more conventional range which is between 0 and 1.

**Step 5: Visualizing the study outline (optional)**

The `plotStudyOutline()` function can be applied to visualize how various variables are distributed across the samples. In addition, this can assess the association between all specified variables. Before creating a study outline plot, the `renameVariables()` function will be used to rename the variables for visualization purposes.

```
# re-naming several columns/variables for visualization
brca.se.obj <- renameVariables(
    se.obj = brca.se.obj,
    current.names = c('tissue_type', 'tumour.purity.estimate', 'library.size'),
    new.names = c('Tissues', 'Tumour.purity', 'Library.size'))
brca.se.obj <- plotStudyOutline(
    se.obj = brca.se.obj,
    variables = c('Years', 'Plates', 'TSS', 'Library.size', 'Tumour.purity', 'Tissues'))
```

The output of the `plotStudyOutline()` function shows that library sizes vary across samples. In particular, samples profiled between 2012 and 2014 exhibit lower library sizes compared with the other samples (Supplementary Figure 6a). We previously found that these differences in library size are mainly explained by the changes in flow cell chemistry. Accordingly, samples are divided into two major flow cell chemistry batches: samples from 2010–2011 are annotated as `Batch1`, and the remaining samples are annotated as `Batch2`. Supplementary Figure 6b shows the tumor purity estimates are highly correlated. Then, the ESTIMATE values will be used for downstream analysis.

```
brca.se.obj$Flow.cells <- factor(
  ifelse(brca.se.obj$Years %in% c(2010, 2011), "Batch1", "Batch2"),
  levels = c("Batch1", "Batch2"))
```

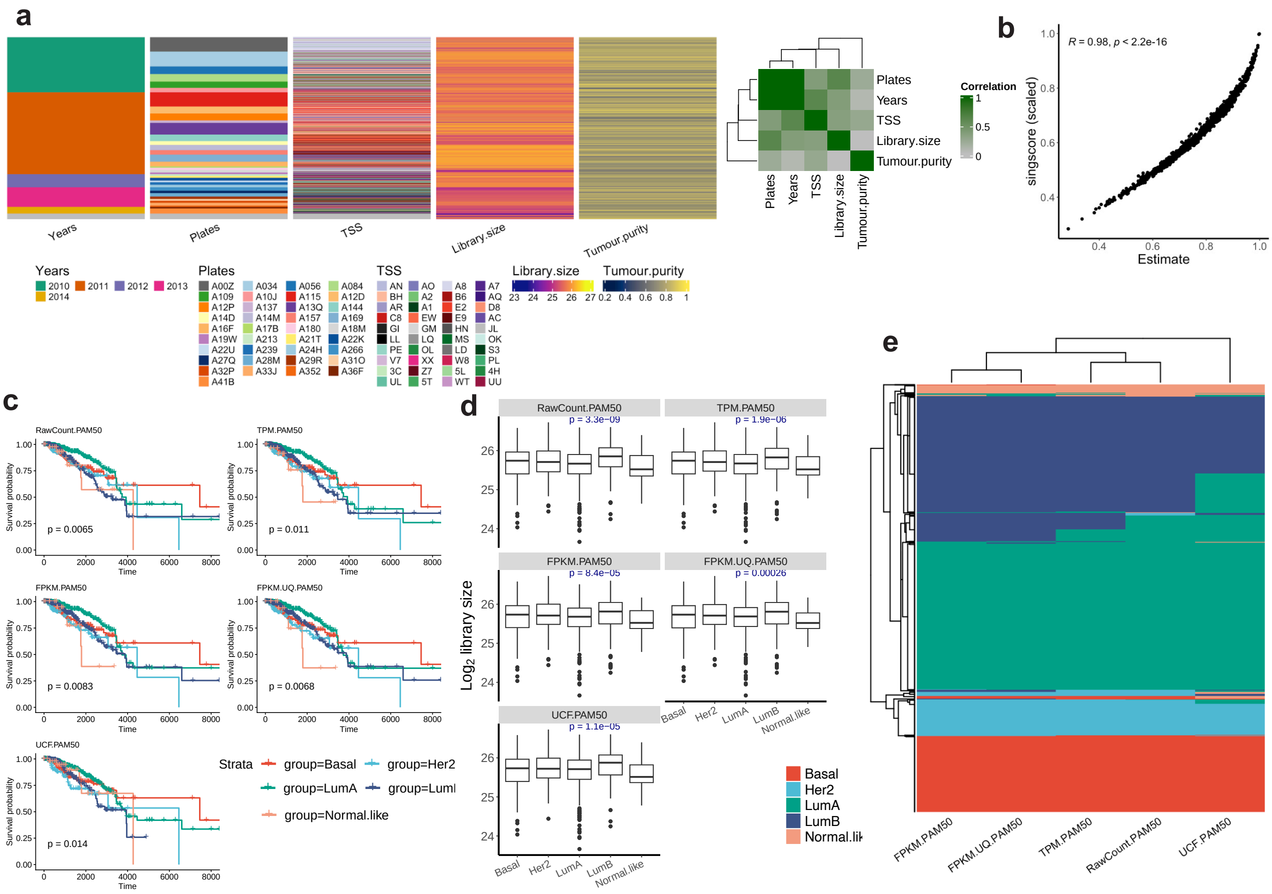

Supplementary Figure 6: Study outline and PAM50 estimates of the TCGA BRCA RNA-seq data. a) The heatmap shows that the data were collected from 40 tissue source sites (TSS) and distributed across 40 sequencing plates for profiling at 5 time points over a span of 5 years. The library sizes were calculated using the raw counts after removing lowly expressed genes. Tumor purity was estimated using the ESTIMATE method on the TCGA FPKM.UQ normalized data. The correlation matrix shows the pairwise correlations among all variables in the dataset. b) This scatter plot shows the correlation between tumor purity obtained using the ESTIMATE and singscore methods. c) Kaplan-Meier plots showing the association between the estimated PAM50 subtypes and the survival outcomes of patients for different datasets. d) Shows the distribution of library size within each estimated PAM50. One-way ANOVA was used to assess the association. e) Heatmap showing the estimated PAM50 subtypes for the differently normalized BRCA samples.

### Stage 2: Identification of major variation

This stage is unnecessary when all sources of unwanted variation and the primary gene expression–based biological variation are already known or have been estimated using appropriate orthogonal platforms or methods. It is important to note that only biological variables that are detectable in the gene expression data can be used effectively in the RUV-III normalization procedure.

##### Step 1: Biological populations

Breast cancers are generally divided into five intrinsic gene-expression-based biological populations including Her2, Basal, Luminal-A, Luminal-B and normal-like. These are obtained using a 50-gene signature (PAM50) that classifies breast cancer into the intrinsic subtypes[9]. It has been shown that PAM50 subtypes are associated with clinical markers including overall survival of patients. Generally, breast cancer patients with Basal and Her2 subtypes are associated with the worst overall survival, while those with Luminal-A are associated with the best overall survival [10]. Several approaches and method are available to estimate the PAM50 subtypes in gene expression data. We will apply the `estimatePAM50` that uses the R package genefu [11] (version 2.32.0) to estimate them in the TCGA BRCA RNA-seq data. Note that the *SummarizedExperiment* object contains the PAM50 annotations from the TCGA breast cancer network. We will label these annotations as `UCF.PAM50`. Any `NA` values in the annotations will be labeled as `Not.classified`.

```
# labeling the TCGA PAM50 subtypes
brca.se.obj$UCF.PAM50 <- brca.se.obj$paper_BRCA_Subtype_PAM50
brca.se.obj$UCF.PAM50[brca.se.obj$UCF.PAM50 == 'Normal'] <- 'Normal.like'
brca.se.obj$UCF.PAM50[is.na(brca.se.obj$UCF.PAM50)] <- 'Not.classified'
brca.se.obj$UCF.PAM50 <- factor(
    x = brca.se.obj$UCF.PAM50,
    levels = c('Basal','Her2','LumA','LumB','Normal.like','Not.classified'))

# applying genefu algorithm
brca.se.obj <- estimatePAM50(
    se.obj = brca.se.obj,
    assay.names = 'all',
    entrez.gene.id = 'entrezgene_id',
    probe = 'gene_name',
    normalization = NULL,
    tissue.type = 'Tissues')
```

*Evaluation of the estimated PAM50 subtypes:* Two analyses will be performed to assess the quality of the estimated PAM50 subtypes for the individual datasets. These analyses will include evaluating the association between the subtypes and the overall survival of patients, as well as examining the relationship with library size. An ideal estimate of the PAM50 should have significant association with survival data and no or weak association with library size variation.

*Survival analysis:* RUVprps provides the function `AddTcgaClinicInfo()`, which enables users to integrate selected TCGA clinical data from Liu et al. [12] into a *SummarizedExperiment* object. The clinical data are available as Supplementary Table 1 ("mmc1.xlsx") from that study. In this analysis, overall survival information for the TCGA BRCA RNA-seq dataset will be extracted and incorporated into the *SummarizedExperiment* object. The function assigns a value of -1 to overall survival times and events for normal tissue samples, facilitating their exclusion from downstream survival analyses. Subsequently, the `computeSurvival()` function will be used to generate Kaplan–Meier survival curves for each PAM50 classification across the differently normalized TCGA datasets. Samples that could not be assigned to any PAM50 subtype will be excluded prior to performing the survival analysis.

```
# adding the TCGA overall survival data
brca.se.obj <- addTcgaClinicalInfo(
    se.obj = brca.se.obj,
    factors = c("OS", "OS.time"),
```

```r
    tissue.type = 'Tissues')

# applying survival analysis
all.survivals <- lapply(
    pam50 <- grep('\\.PAM50$', colnames(colData(brca.se.obj)), value = TRUE),
    function(i) {
        idx <- brca.se.obj$OS != -1 & brca.se.obj[[i]] != 'Not.classified'
        computeSurvival(
            se.obj = brca.se.obj[, idx],
            variable = i,
            survival.time = 'OS.time',
            survival.events = 'OS',
            genes = NULL,
            return.survival.plot = TRUE,
            check.se.obj = FALSE,
            save.se.obj = FALSE)$plot })
```

The survival analysis indicates that the estimated PAM50 for the TCGA FPKM.UQ data are more accurate (lowest p-value) compared to those derived from other datasets (Supplementary Figure 6c). Then, these estimates will be used as major source of known biological variation for downstream analysis.

*Association with library size:* Supplementary Figure 6d shows that the PAM50 subtypes are significantly associated with library size, regardless of the datasets. This indicates that the estimated PAM50 may not be completely accurate. However, the FPKM.UQ shows less association between the PMA50 subtypes and library size compared with the other datasets. Supplementary Figure 6e shows no substantial differences in the PAM50 subtypes estimates across datasets.

```r
# creating boxplots of library size across the estimated PAM50 subtypes
plotVariables(
    se.obj = brca.se.obj[ , brca.se.obj$UCF.PAM50 != 'Not.classified'],
    x.variables = grep('\\.PAM50$', colnames(colData(brca.se.obj)), value = TRUE)[c(2,3,4,5,1)],
    y.variables = 'Library.size',
    y.lab = expression(Log[2]~library~size),
    generate.heatmap = TRUE,
    nb.ncol = 2)
```

**Step 2: Unwanted variation**

Supplementary Figure 6a shows the substantial library size and tumor purity variation across samples. We have previously showed that PC plots of the TCGA BRCA RNAseq data clearly reveal the batch effects introduced by the change in flow cell chemistry [1]. However, there are situations where batch details are not available or are unknown. In these situations batch effects can be estimated from the data using different robust approaches proposed by RUVprps. The `identifyUnknownUV()` function will be applied using the `rle` approach and parameters specified for chronologically ordered samples. Note that biological and known unwanted variables will be used only to assess the estimated unknown unwanted variation and are not required for its estimation.

```r
brca.se.obj <- identifyUnknownUV(
    se.obj = brca.se.obj,
    assay.name = 'RawCount',
```

```
approach = 'rle',
chronological.detection = TRUE,
assess.bio.association = TRUE,
bio.variables = c('FPKM.UQ.PAM50', 'Tumour.purity'),
assess.uv.association = TRUE,
uv.variables = c('Years', 'Plates'),
generate.association.plot = TRUE,
add.to.sample.annotation = TRUE,
col.name = 'Estimated.batches',
output.name = 'brca.estimated.uv')
```

Supplementary Figure 7a and b shows that the RLE medians of the TCGA raw count ($\log_2 +1$) data reflect separation by year and plate effects. These results were further confirmed by confusion matrix between the estimated batches and known sources of unwanted variation in the data (Supplementary Figures 7b and c). Further, the first three PCs of the 500 most strongly affected genes (identified by ANOVA) shows a clear separation of the estimated batches (Supplementary Figures 7d). The first PC1 displays a strong correlation with library size (Supplementary Figure 7e), whereas the first three PCs show no association with tumor purity. These results indicate that the RLE-based approach captures a dominant source of unwanted variation at a time. To estimate sources of unwanted variation such as tumor purity, two strategies can be employed: (i) applying `pca` or `sample.scoring` approaches using validated stromal and immune gene signatures; or (ii) regressing out the initial unwanted variation estimated by the RLE approach and subsequently estimating tumor purity from the residuals using the same RLE framework. Here, we use the estimated tumor purity bu the `prepareSeObj()` function as an additional source of unwanted variation to be removed by RUV-III-PRPS normalization.

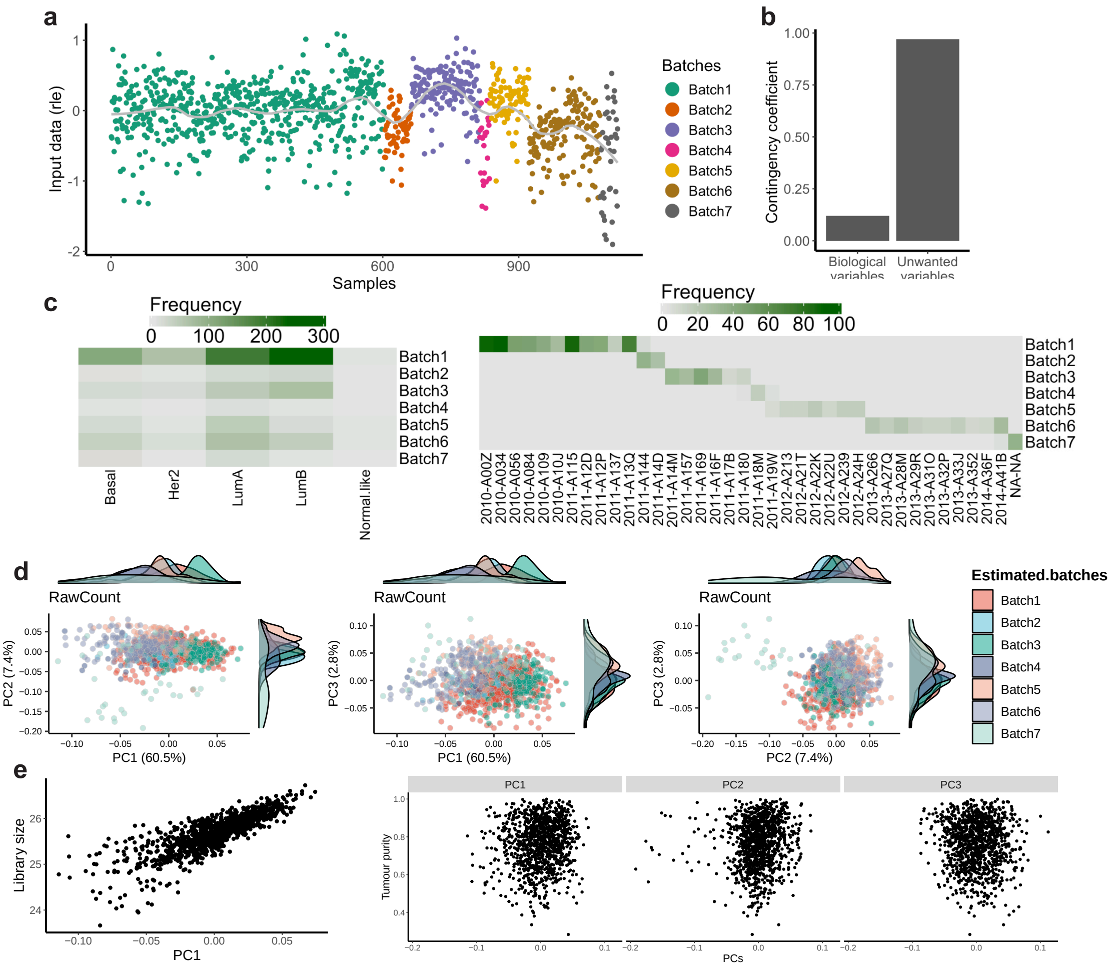


Supplementary Figure 7: Identification of batches in TCGA BRCA RNA-seq data using an unsupervised approach in RUVprps. a) The scatter plot displays the RLE medians calculated from the raw count data ($\log_2 + 1$) using all genes, categorized into potential batches using the changepoint approach in the `identifyUnknownUV()` function. b) Cramer's V contingency coefficient quantifying the association between the estimated batches and biological variable (PAM50 subtypes) as well as unwanted variation (a product of the plates and years variables). c) The left hand side heatmap presents sample counts, illustrating the association between the estimated batches and the PAM50 subtypes. The right hand side heatmap presents sample counts, illustrating the association between the estimated batches and known unwanted sources of variation including the plated and years. d) The scatter plots of first three PCs on TCGA BRCA raw count data ($\log_2 + 1$) using top 500 highly affected genes by the estimated batches, colored by the estimates of batch effects. e) first plot: shows the association between the first PC from d with library size in the data. second plot: displays the association between the firs three PCs from d with tumor purity estimates in the data

#### Stage 3: Variation assessment

Before applying RUV-III normalization, it is essential to evaluate the impact of both known and estimated biological and unwanted sources of variation across all datasets. At this stage, a range of statistical metrics, categorized into gene and global-level measures, are applied to all datasets.

**Step 1: Checking the SummarizedExperiment object**

After selecting both biological and unwanted variables, the `checkSeObj()` function should be applied to remove any `NA` or missing values from both the datasets and the specified variables. Note that the current RUV-III method does not support `NA` or missing values in the data.

```
brca.se.obj$PAM50 <- brca.se.obj$FPKM.UQ.PAM50
brca.se.obj <- checkSeObj(
  se.obj = brca.se.obj,
  assay.names = 'all',
  variables = c('PAM50','Library.size','Tumour.purity','Flow.cells', 'Estimated.batches'),
  remove.na = 'both')
```

The results show that there is no any `NA` or missing values in all the datasets and the specified sample annotation. Note that, the `check.se.obj` argument in all downstream functions can be set to `FALSE`.

**Step 2: Creating all possible assessments**

The `getAssessmentMetrics()` function generates all available plots and assessments for the specified biological and unwanted variables. Any plots or assessments that are not of interest can be excluded in subsequent steps. We recommend generating the full set assessment initially to visually explore the variation, and then selecting which to retain for downstream analyses.

```
brca.se.obj <- getAssessmentMetrics(
    se.obj = brca.se.obj,
    variables = c('PAM50','Library.size','Tumour.purity','Flow.cells', 'Estimated.batches'),
    plot.output = FALSE)
```

Supplementary Figure 8 shows that 66 plots and metrics will be generated to assess the impact of both biological and unwanted variation in individual datasets. Note that the "code" for each assessment can be used to exclude specific assessments from downstream analysis.

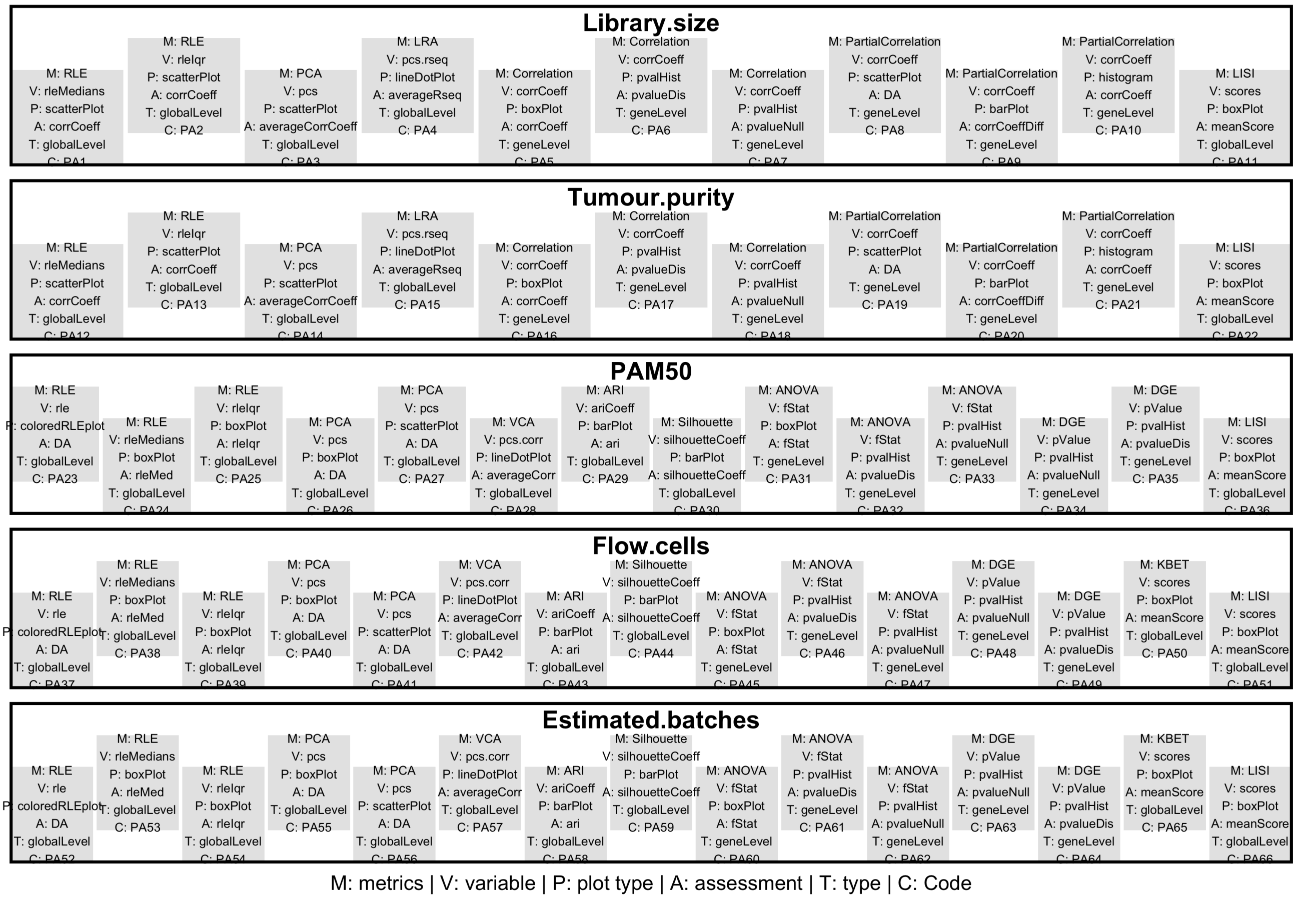


Supplementary Figure 8: Assessment matrices for all the biological and unwanted variation for the TCGA BRCA RNA-seq data. For each individual variable, the specified metrics will be computed and the corresponding plots will be generated. The code of each assessment can be used to exclude the corresponding metrics from the assessment of variation and normalization steps. The 'DA' (Does not Apply) indicates that specific metrics will not be considered for assessing normalization.

#### Step 3: Selecting genes for gene-gene correlation assessment and tumor purity estimation

Forming partial gene-gene correlations is one of the tools in the variation assessment process. If this is selected, it is desirable to provide a list of genes whose expression is highly associated with the corresponding variable; otherwise, all genes will be used, which is computationally expensive. Further, gene set enrichment analysis, particularly for estimation of tumor purity, is one the metrics in the variation assessment process. To estimate tumor purity for individual datasets, the immune and stromal genes sets will be used [6].

The `selectGenesForPPcorr()` function will be applied to find genes that have certain correlation with library size and tumor purity, separately. We specify the `Flow.cells` variable so that correlation analyses are performed within each major batch, thereby mitigating batch effects. The results of each batch will be considered together to select genes.

The `selectGeneSets()` function retrieves stromal and immune gene signatures from the gene annotations provided in the

`SummarizedExperiment` object. These gene sets were added during preparation of the `SummarizedExperiment` object.

```
ppcorr.gene.sets <- selectGenesForPPcorr(
    se.obj = brca.se.obj,
    assay.names = c('RawCount', 'FPKM.UQ'),
    variables = c('Library.size', 'Tumour.purity'),
    cor.cutoff = 0.6,
    groups = 'Flow.cells')

tumour.purity.genes.sets <- selectGenesSets(
    se.obj = brca.se.obj,
    gene.set = "immune.stromal")
```

The 3,327 and 700 genes that have an absolute correlation of more than 0.6 (arbitrary cutoff) with library size and purity were selected for partial gene-gene correlation analysis. The total number of 273 genes were retrieved to estimate the variation of tumor purity in the data. Note that, any other true gene signatures can be used for estimation of tumor purity.

**Step 4: Assessing variation**

The `assessVariation()` function applies all statistical metrics to the specified variables, and stores the resulting summaries and plots within the *SummarizedExperiment* object. The `plotAssessVariation()` function generates a PDF file containing all plots, which is saved to the current home directory.

```
# assessing variation in both biological and unwanted variables
brca.se.obj <- assessVariation(
    se.obj = brca.se.obj,
    assay.names = 'all',
    bio.variables = c('PAM50'),
    uv.variables = c('Library.size','Tumour.purity','Flow.cells'),
    pcorr.genes = ppcorr.gene.sets,
    gene.set.score.list = tumour.purity.genes.sets,
    override.check = FALSE,
    check.se.obj = FALSE)

plotAssessVariation(
    se.obj = brca.se.obj,
    assay.names = 'all',
    variables = c('PAM50','Library.size','Tumour.purity','Flow.cells'),
    output.file.name = paste0(figPathSc0, 'RUVprps_TCGA_BRCA_InitialVariationAssessments'))
```

Supplementary File "RUVprps_TCGA_BRCA_InitialVariationAssessments.pdf" shows all the plots of the result of gene- and global-level metrics that are produced by the `assessVariation()` function. Briefly, the first three PCs of the TCGA normalized datasets clearly show that the effect of the library size, tumor purity and flow cell chemistries in these datasets. This conclusion is further supported by all the gene- and global-level assessments. Collectively, these results indicate that TCGA normalization exhibits shortcomings in effectively removing sources of unwanted variation. We therefore apply RUV-III with PRPS to remove the effects of library size, tumor purity, and flow-cell chemistry from the raw count data while preserving biologically meaningful variation associated with PAM50 subtypes.

**Stage 4: RUV-III normalization**

RUV-III with PRPS will be employed under 4 different scenarios spanning from fully supervised to fully unsupervised applications. The ultimate aim in each scenario is to preserve known biological variation and remove or mitigate the impact of different sources of unwanted variation. The general process within each scenario comprises of identification of a set of NCGs, PRPS sets and RUV-III normalizations with different values for $k$.

**Scenario 1: Both biological and unwanted variation are known**

In this scenario, only known sources of biological and unwanted variation will be considered in the RUV-III normalization. Library size, tumor purity and flow cell chemistries will be used as the known sources of unwanted variation, while the PAM50 subtypes will be recognized as the major biological variation. RUV-III normalization will be conducted in a fully supervised manner.

**Step 1: Selecting negative control genes**

A set of NCGs will be identified by the implementation of the `findNcgSupervised()` function. All the available approaches will be applied and then the `compareNCG()` function will be used to select the most suitable set of NCG. The evaluation of their performance assesses their ability to effectively capture sources of unwanted variation while exhibiting minimal association with biological variation .

```
ncg.options <- c(
    'AnovaCorr.AcrossAllSamples',
    'AnovaCorr.PerBatchPerBiology',
    'LinearMixedModel',
    'TwoWayAnova')
for(i in 1:4){
    brca.se.obj <- findNcgSupervised(
        se.obj = brca.se.obj,
        assay.name = 'RawCount',
        bio.variables = c('PAM50'),
        uv.variables = c('Library.size', 'Flow.cells', 'Tumour.purity'),
        approach = ncg.options[i],
        form =  ~ (1|Flow.cells) + Library.size + (1|PAM50) + Tumour.purity,
        bio.percentile = NULL,
        uv.percentile = 0.8,
        use.rank = FALSE,
        ncg.selection.method = 'quantile',
        nb.bio.clusters = 3,
        pseudo.count = 0.5,
        adjust.data = TRUE,
        adjustment.variables = "uv",
        nb.ncg = 0.05,
        ncg.group.name = 'brca.scenario1',
        ncg.set.name = ncg.options[i],
        nb.cores = 15,
        check.se.obj = FALSE,
        assess.ncg = TRUE)}

brca.se.obj <- compareNCGs(
```

```
    se.obj = brca.se.obj,
    assay.name = 'RawCount',
    ncg.set.names = "all",
    bio.variables = c('PAM50'),
    uv.variables =  c('Library.size','Tumour.purity','Flow.cells'),
    nb.pcs = 10,
    ncg.type = 'supervised',
    ncg.group.name = 'brca.scenario1',
    check.se.obj = FALSE)
```

Supplementary Figure 9a and b show the satisfactory performance of all four NCG selection approaches. The 765 NCG identified by the `LinearMixedModel` approach exhibits slightly better performance compared to other approaches. Then this set of NCG will be used for RUV-III normalization.

**Step 2: Creating the PRPS data**

The PRPS data will be generated for individual sources of unwanted variation using the `createPrPsSupervised()` function. In this process, to avoid possible PRPS contamination, all other sources of unwanted variation will be taken into account while constructing the PRPS for a given source (`apply.other.uv.variables = TRUE`).

```
brca.se.obj <- createPrPsSupervised(
    se.obj = brca.se.obj,
    assay.name = 'RawCount',
    bio.variables = 'PAM50',
    uv.variables = c('Library.size','Tumour.purity','Flow.cells'),
    nb.other.uv.clusters = 4,
    apply.other.uv.variables = TRUE,
    apply.log = FALSE,
    prps.group.name = 'brca.scenario1',
    verbose = TRUE)
```

In total, 28, 28, and 27 PR sets, corresponding to 56, 56, and 54 PS respectively, were generated for the removal of library size, flow cell, and tumor purity effects (Supplementary Figure 9c). Note that the `apply.log` argument was set to `FALSE`, indicating that PRPS values were computed directly from raw count data without log transformation. Therefore, we will ensure that the `RUVIIIprps()` function applies a log transformation to the PRPS data before use. Conversely, if `apply.log` is set to `TRUE`, the function first applies a log transformation to the raw count data before generating PRPS values, in which case `RUVIIIprps()` should not apply a log transformation to the PRPS data.

**Step 3: Performing RUV-III normalization with multiple $k$ values**

First, the `getMaximumK()` function will be applied to determine the maximum possible value of $k$. For the current NCGs and PRPS data, this maximum is 86. A range of $k$ values from 1 to 20 will then be tested. The RUV-III-PRPS normalization will be performed on the `RawCount` ($\log_2 + 0.5$) data. All RUV-III normalized data will be stored as new datasets in the *SummarizedExperiment* object.

```
brca.se.obj <- RUVIIIprps(
    se.obj = brca.se.obj,
    assay.name = 'RawCount',
    control.sample.types = 'prps',
```

```
    prps.type = 'supervised',
    prps.group  = 'brca.scenario1',
    prps.set.names = 'all',
    ncg.type = 'supervised',
    ncg.group.names = 'brca.scenario1',
    ncg.set.names = 'LinearMixedModel',
    k = c(1:20),
    apply.log = TRUE,
    data.to.log = 'both',
    pseudo.count = 0.5,
    check.se.obj = FALSE)
```

Note that the `data.to.log` argument was set to `both`, indicating that both the `RawCount` and PRPS data have not been log-transformed. The `RUVIIIprps` function therefore applied a log transformation with a pseudo-count to both prior to performing RUV-III normalization.

**Step 4: Assessing the W matrix**

Before assessing the performance of normalizations, we will examine the association between the biological and unwanted variables with the unwanted factors W estimated by RUV-III. Ideally, the columns of the estimated W should have strong and weak associations with the unwanted and biological variables, respectively.

```
brca.se.obj <- assessW(
    se.obj = brca.se.obj,
    compare.w = TRUE,
    variables = NULL,
    bio.variables = 'PAM50',
    uv.variables = c('Library.size','Tumour.purity','Flow.cells'))
```

Supplementary Figure 9d indicates that the unwanted factors W estimated by RUV-II are highly associated with all sources of unwanted variation, while they show reasonably low association with the biological variable. This analysis will be considered alongside with other metrics to select optimal $k$ value for RUV-III normalization.

**Step 5: Evaluating normalizations**

To assess the performance of RUV-III normalization and compare it with the TCGA normalizations, we will first apply the `assessVariation()` function, followed by the `assessNormalization()` function, across all RUV-III normalized data in the *SummarizedExperiment* object. Prior to variation assessment, the PAM50 subtypes will be estimated, and their association with patient survival will be evaluated for individual RUV-III normalized datasets.

```
brca.se.obj <- estimatePAM50(
    se.obj = brca.se.obj,
    assay.names = names(SummarizedExperiment::assays(brca.se.obj))[-c(1:4)],
    entrez.gene.id = 'entrezgene_id',
    probe = 'gene_name',
    normalization = NULL,
    apply.log = FALSE,
    tissue.type = 'Tissues')
```

```
all.survivals <- lapply(
    pam50 <- grep('\\.PAM50$', colnames(colData(brca.se.obj)), value = TRUE),
    function(i) {
        idx <- brca.se.obj$OS != -1
        computeSurvival(
            se.obj = brca.se.obj[, idx],
            variable = i,
            survival.time = 'OS.time',
            survival.events = 'OS',
            genes = NULL,
            return.survival.plot = TRUE,
            check.se.obj = FALSE,
            save.se.obj = FALSE)$plot })
```

The results of the survival analysis can be used a metric to select a suitable value of *K*. We will assess the performance of RUV-III normalized datasets and then compare them with TCGA normalized datasets. Note that there is no need to re-calculate the metrics for the TCGA datasets, as they have already been computed and stored in the *SummarizedExperiment* object. By setting `override.check = TRUE`, the function will not recompute metrics already available for each dataset. However, if one wishes to re-calculate the metrics for the TCGA datasets, the data must first be log-transformed, after which the `assessVariation()` function can be applied with `apply.log = FALSE`, since all RUV-III normalized data are already in log scale.

```
brca.se.obj$PAM50 <-  brca.se.obj$FPKM.UQ.PAM50
brca.se.obj <- assessVariation(
    se.obj = brca.se.obj,
    assay.names = 'all',
    assessment.level = 'L2',
    bio.variables = c("PAM50"),
    uv.variables = c('Library.size','Tumour.purity','Flow.cells'),
    pcorr.genes = ppcorr.gene.sets,
    gene.set.score.list = tumour.purity.genes.sets,
    pcorr.filter.genes = FALSE,
    general.points.size = 3,
    override.check = TRUE,
    apply.log = FALSE,
    check.se.obj = FALSE)

brca.se.obj <- assessNormalization(
    se.obj = brca.se.obj,
    assay.names = 'all',
    assessment.level = 'L2',
    select.top.ruv = FALSE,
    bio.variables = c("PAM50"),
    uv.variables = c('Library.size','Tumour.purity','Flow.cells'),
    bio.weight = 0.6,
    uv.weight = 0.4,
    corr.cutoff = list(Tumour.purity = .5, Library.size = .3),
    output.name = 'brca.read.assessNormalization.scenario1')

RUVprps:::plotAssessVariation(
    se.obj = brca.se.obj,
```

```
assay.names = 'all',
variables = c("Tumour.purity", "PAM50", "Library.size", "Flow.cells"),
output.file.name = "RUVprps_TCGA_BRCA_VariationAssessment_Scenario1))
```

Supplementary Figure 9e, which presents the level 2 normalization assessment, clearly demonstrates that RUV-III with PRPS effectively removes sources of unwanted variation while preserving known biological variation, in contrast to the TCGA-normalized data. These analyses indicate that RUV-III-PRPS normalization with $k = 20$ achieves the highest total performance score. However, differences between $k = 20$ and other values of k>5 are not significant. If using a high value of is concerning for downstream analyses, RUV-III-PRPS normalization with $k = 6$ may also be considered.

Supplementary Figures 9f further demonstrate satisfactory separation of the PAM50 molecular subtypes and effective mixing of flow-cell batches in the first three principal components. Supplementary Figures 9g show that purity-associated variation is effectively removed by RUV-III-PRPS when using $k = 20$.

The Supplementary File "RUVprps_TCGA_BRCA_VariationAssessment_Scenario1.pdf" exhibits all the plots summarized in the Supplementary Figure 9e. Supplementary Figure 10 shows the association between PAM50 subtypes and patient survival. These results indicate that PAM50 subtypes derived from RUV-III-PRPS normalization with $k = 20$ are significantly associated with survival.

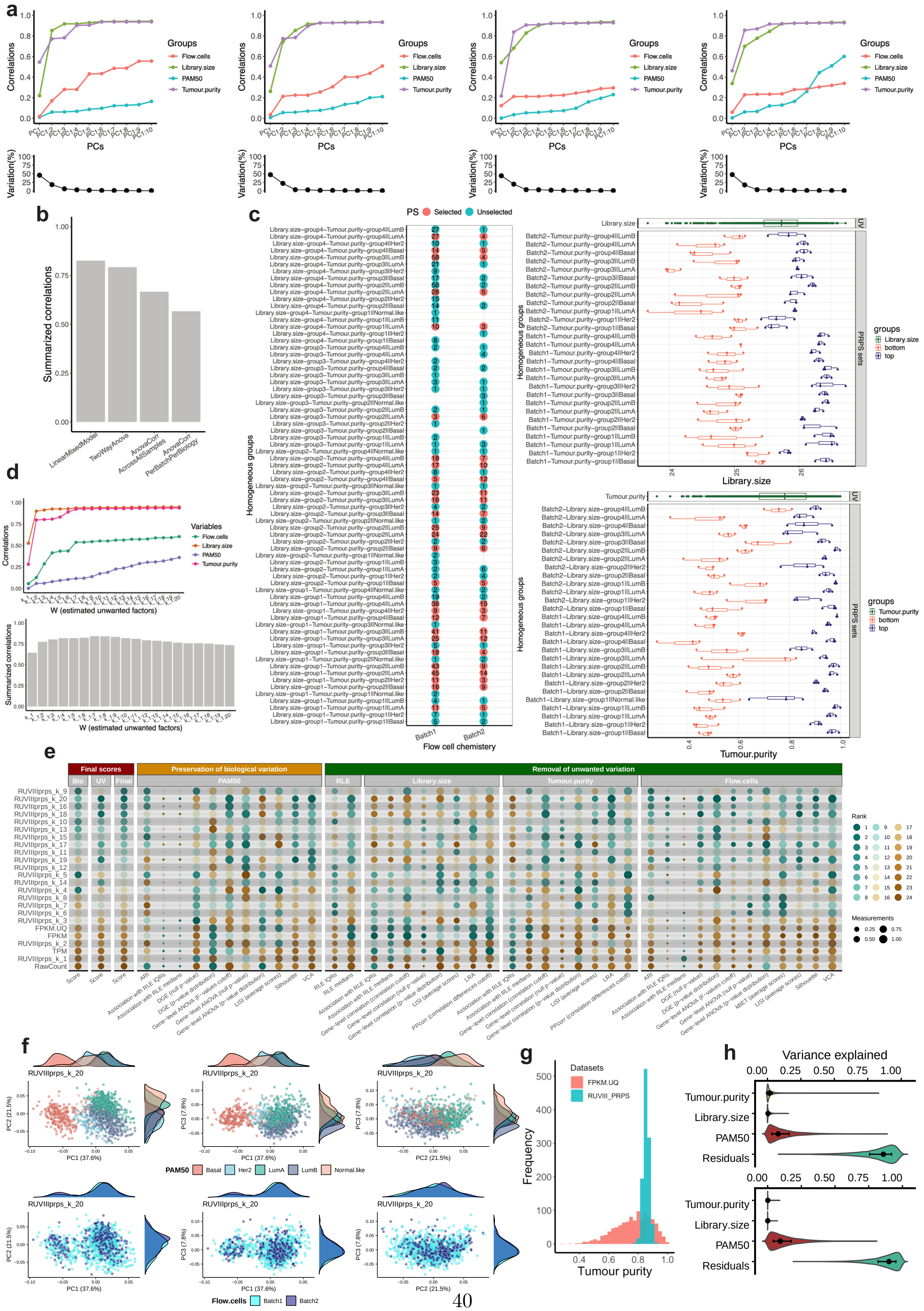

Supplementary Figure 9: RUV-III normalization of the TCGA BRCA RNA-seq data for scenario 1. a) The line-dot plots show the performance of the selected normalization methods using different approaches including 'LinearMixedModel' (first plot),'TwoWayAnova' (second plot), 'AnovaCorr.AcrossAllSamples' (third plot), and 'AnovaCorr.PerBatchPerBiology' (fourth plot), on the TCGA BRCA raw count ($\log_2 + 0.5$)RNA-seq data. b) The barplot summarizes the performance of the selected NCG sets. The higher the correlation, the better the performance. c) The PRPS maps are shown for the batches of flow cell chemistry (first plot), library size (second plot) and tumor purity variables (third plot). For each PRPS map of a unwanted variable, the Y-axis displays all possible homogeneous groups with respect to the known biological variation and other unwanted variables. The X-axes represent the distributions of samples across the flow cell chemistry (batch 1 and batch 2), library size and tumor purity ranges, respectively. In the flow cell chemistry PRPS map, red indicates groups of samples containing more than two samples. Each horizontal line connecting two red points defines a PRPS set across the batches of flow cell chemistry. In the library size PRPS map, each horizontal line corresponds to six samples representing the highest and lowest library sizes within a homogeneous sample group. Every three samples are averaged to create a PS. The X-axis of the library size PRPS map shows the $\log_2 n$ of the library size in the data and highlights the samples selected for PS construction. In the tumor purity PRPS map, each horizontal line corresponds to six samples representing the highest and lowest tumor purity within a homogeneous sample group. Every three samples are averaged to create a PS. The X-axis shows the tumor purity estimate in the data and highlights the samples selected for PS construction. d) The line-dot plot shows the associations between the columns of the estimated W matrix and both biological and unwanted variables. The bar plot summarizes these correlations. A higher correlation the better performance. e) Numerical assessments of all metrics for both biological and unwanted variables are presented. Each point represents the performance of a given normalization method, where higher values indicate better performance. Normalized datasets are ranked according to their effectiveness in removing unwanted variation while preserving biological signals. f) The first three PC plots of RUV-III with $K = 20$ are shown, colored by PAM50 subtypes (top row) and the batches of flow cell chemistries (bottom row). The PCA were performed on a PAM50 subtypes gene signatures proposed by genefu R package. g) The plot shows the tumor purity estimated by the ESTIMATE method using the TCGA FPKM.UQ and the RUV-III-PRPS normalized data with $K = 20$. h) Shows the results of a linear mixed model analysis followed by gene-level variation analysis across all the variables for the TCGA FPKM.UQ (top plot) and the RUV-III-PRPS normalized data with $K = 20$ (bottom plot).

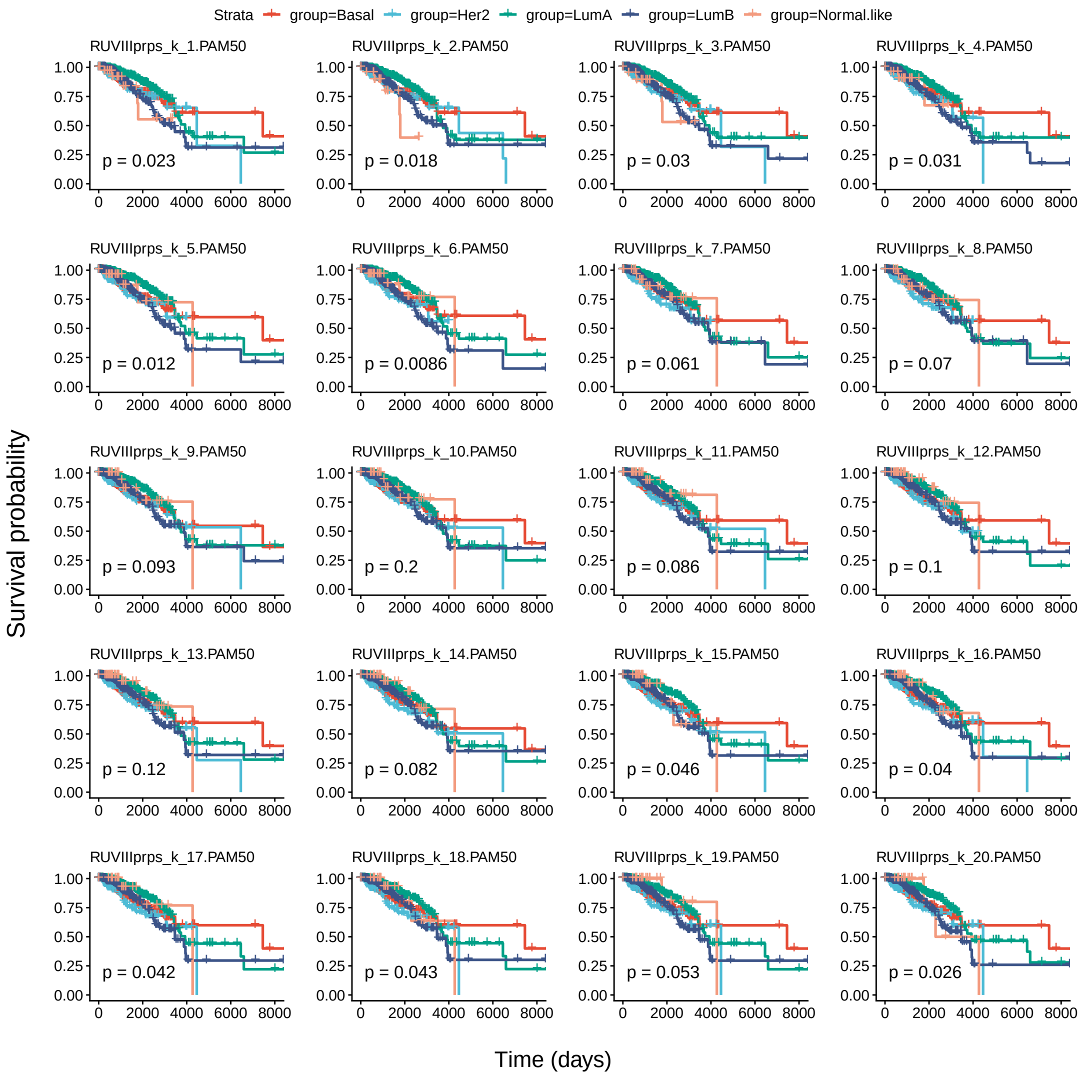


Supplementary Figure 10: Kaplan–Meier analysis of PAM50 subtypes derived from each RUV-III–normalized dataset in Scenario 1, showing their association with patient survival in the TCGA BRCA RNA-seq data

#### Scenario 2: Unknown unwanted and known biological variation

In second scenario, it is assumed that the batches in the data are unknown. Then, the estimated batches (refer to step 2 of stage 2) along with library size and tumor purity will be used as sources of unwanted variation. In this scenario, the PAM50 will be used as known biological variation.

#### Step 1: Selecting negative control genes

A set of NCGs is obtained using the `findNcgsSupervised()` function. In this framework, PAM50 subtypes are considered as representing known biological variation, whereas estimated batches, library size, and tumor purity are treated as sources of unwanted variation. All approaches available within the `findNcgsSupervised()` function will be applied and evaluated to identify a suitable set of NCGs.

```
ncg.options <- c(
    'AnovaCorr.AcrossAllSamples',
    'AnovaCorr.PerBatchPerBiology',
    'LinearMixedModel',
    'TwoWayAnova')
for(i in 1:4){
    brca.se.obj <- findNcgSupervised(
        se.obj = brca.se.obj,
        assay.name = 'RawCount',
        bio.variables = c('PAM50'),
        uv.variables = c('Library.size', 'Estimated.batches', 'Tumour.purity'),
        approach = ncg.options[i],
        form =  ~ (1|Estimated.batches) + Library.size + (1|PAM50) + Tumour.purity,
        bio.percentile = NULL,
        uv.percentile = 0.8,
        use.rank = FALSE,
        ncg.selection.method = 'quantile',
        nb.bio.clusters = 3,
        pseudo.count = 0.5,
        adjust.data = TRUE,
        adjustment.variables = "uv",
        nb.ncg = 0.05,
        ncg.group.name = 'brca.scenario2',
        ncg.set.name = ncg.options[i],
        nb.cores = 15,
        check.se.obj = FALSE,
        assess.ncg = TRUE)}

brca.se.obj <- compareNCGs(
    se.obj = brca.se.obj,
    assay.name = 'RawCount',
    ncg.set.names = "all",
    bio.variables = c('PAM50'),
    uv.variables =  c('Library.size','Tumour.purity','Estimated.batches'),
    nb.pcs = 10,
    ncg.type = 'supervised',
    ncg.group.name = 'brca.scenario2',
    check.se.obj = FALSE)
```

Supplementary Figures 11a and b show that all four NCG selection approaches perform satisfactorily. Among them, the 765 NCGs identified by the `LinearMixedModel` approach show slightly superior performance relative to the others. This set of NCGs will be therefore selected for subsequent RUV-III normalization.

##### Step 2: Creating the PRPS data

Different sets of PRPS will be created for library size, the estimated batches and tumor purity using the the function `createPrPsSupervised()` function. In this process, all other sources of unwanted variation will be taken into account while constructing the PRPS for a given source. This will avoid possible PRPS contamination.

```
brca.se.obj <- createPrPsSupervised(
```

```
    se.obj = brca.se.obj,
    assay.name = 'RawCount',
    bio.variables = 'PAM50',
    uv.variables = c('Library.size','Tumour.purity','Estimated.batches'),
    nb.other.uv.clusters = 2,
    apply.other.uv.variables = TRUE,
    apply.log = FALSE,
    prps.group.name = 'brca.scenario2',
    verbose = TRUE)
```

In total, 36, 16, and 32 PR sets, corresponding to 72, 65, and 64 PS respectively, were generated for the removal of library size, estimated batches, and tumor purity effects (Supplementary Figure 11c). Similar to scenario 1, the PRPS data were created using the raw count data without any log transformation.

##### Step 3: Performing RUV-III normalization with multiple $k$ values

The `getMaximumK()` function will first be applied to determine the maximum feasible value of $k$, which for the current NCG and PRPS data is 84. Subsequently, $k$ values ranging from 1 to 20 will be tested. RUV-III-PRPS normalization will be applied to the `RawCount` data ($\log_2 +1$), and all resulting RUV-III normalized datasets will be stored as new datasets in a *SummarizedExperiment* object.

```
brca.se.obj <- RUVIIIprps(
    se.obj = brca.se.obj,
    assay.name = 'RawCount',
    control.sample.types = 'prps',
    prps.type = 'supervised',
    prps.group  = 'brca.scenario2',
    prps.set.names = 'all',
    ncg.type = 'supervised',
    ncg.group.names = 'brca.scenario2',
    ncg.set.names = 'LinearMixedModel',
    k = c(1:20),
    apply.log = TRUE,
    data.to.log = 'both',
    pseudo.count = 0.5,
    check.se.obj = FALSE)
```

Note that the `RUVIIIprps()` function performed log transformation on both the `RawCount` and PRPS data (`data.to.log = 'both'`).

##### Step 4: Assessing the W matrix

We will examine the association between the biological and unwanted variables with the unwanted factors W estimated by RUV-III.

```
brca.se.obj <- assessW(
    se.obj = brca.se.obj,
    compare.w = TRUE,
    variables = NULL,
    bio.variables = 'PAM50',
    uv.variables = c('Library.size','Tumour.purity','Estimated.batches'))
```

Supplementary Figure (Figure 11d) indicates that the unwanted factors W estimated by RUV-II are highly associated with all sources of unwanted variation, while they show reasonably low association with the biological variable. This result will be considered to select optimal $k$ value for RUV-III normalization. Note that any biological and unwanted variables not used for normalization can be provided to the functions to assess their association with W.

**Step 5: Evaluating normalizations**

Similar to scenario 1, for individual RUV-III normalized data, the PAM50 subtypes will be estimated, and their association with patient survival will be examined by the `estimatePAM50()` and `computeSurvival()` functions, respectively.

```
brca.se.obj <- estimatePAM50(
    se.obj = brca.se.obj,
    assay.names = names(SummarizedExperiment::assays(brca.se.obj))[-c(1:4)],
    entrez.gene.id = 'entrezgene_id',
    probe = 'gene_name',
    normalization = NULL,
    apply.log = FALSE,
    tissue.type = 'Tissues')

all.survivals <- lapply(
    pam50 <- grep('\\.PAM50$', colnames(colData(brca.se.obj)), value = TRUE),
    function(i) {
        idx <- brca.se.obj$OS != -1
        computeSurvival(
            se.obj = brca.se.obj[, idx],
            variable = i,
            survival.time = 'OS.time',
            survival.events = 'OS',
            genes = NULL,
            return.survival.plot = TRUE,
            check.se.obj = FALSE,
            save.se.obj = FALSE)$plot })
```

The results of the PAM50 subtype survival analysis can be used to identify an optimal value of $K$.

The `assessVariation()` function will be applied on all RUV-III normalized data and then the `assessNormalization()` function will be applied across datasets in the *SummarizedExperiment* object.

```
brca.se.obj$PAM50 <-  brca.se.obj$FPKM.UQ.PAM50
brca.se.obj <- assessVariation(
    se.obj = brca.se.obj,
    assay.names = 'all',
    assessment.level = 'L2',
    bio.variables = c("PAM50"),
    uv.variables = c('Library.size','Tumour.purity','Estimated.batches'),
    pcorr.genes = ppcorr.gene.sets,
    gene.set.score.list = tumour.purity.genes.sets,
    pcorr.filter.genes = FALSE,
    general.points.size = 3,
    override.check = TRUE,
    apply.log = FALSE,
    check.se.obj = FALSE)
```

```
brca.se.obj <- assessNormalization(
    se.obj = brca.se.obj,
    assay.names = 'all',
    assessment.level = 'L2',
    select.top.ruv = FALSE,
    bio.variables = c("PAM50"),
    uv.variables = c('Library.size','Tumour.purity','Estimated.batches'),
    bio.weight = 0.6,
    uv.weight = 0.4,
    corr.cutoff = list(Tumour.purity = .5, Library.size = .3),
    output.name = 'brca.read.assessNormalization.scenario2')

RUVprps:::plotAssessVariation(
    se.obj = brca.se.obj,
    assay.names = 'all',
    variables = c("Tumour.purity", "PAM50", "Library.size", "Flow.cells"),
    output.file.name = "RUVprps_TCGA_BRCA_VariationAssessment_Scenario2")
```

Supplementary Figure 11e presents the level 2 normalization assessment, demonstrates that RUV-III with PRPS effectively removes sources of unwanted variation while preserving known biological signals, in contrast to the TCGA-normalized data. We selected the RUV-III-PRPS normalization with $k = 17$ based on assessment of the W matrix, survival analysis of the PAM50 subtypes, and quantitative evaluation of normalization performance. Supplementary Figures 11f show clear separation of the PAM50 subtypes and reasonable mixing of the batches of flow cells in the first three PCs of the RUV-III-PRPS normalized data. Further, the variation in tumor purity was significantly reduced by RUV-III-PRPS normalization (Supplementary Figures 11g).Results from the linear mixed-effects model analysis, followed by gene-level variance decomposition across all variables, showed that RUV-III-PRPS normalization effectively removes variation explained by unwanted sources while preserving variation associated with the PAM50 subtypes. The Supplementary File "TCGA_BRCA_variationAssessment_Scenario2.pdf" provides all plots summarized in Supplementary Figure 11e. Results of the PAM50 survival analysis showed that RUV-III-PRPS normalization with $k = 17$ was significantly associated with survival (Supplementary Figure 12).

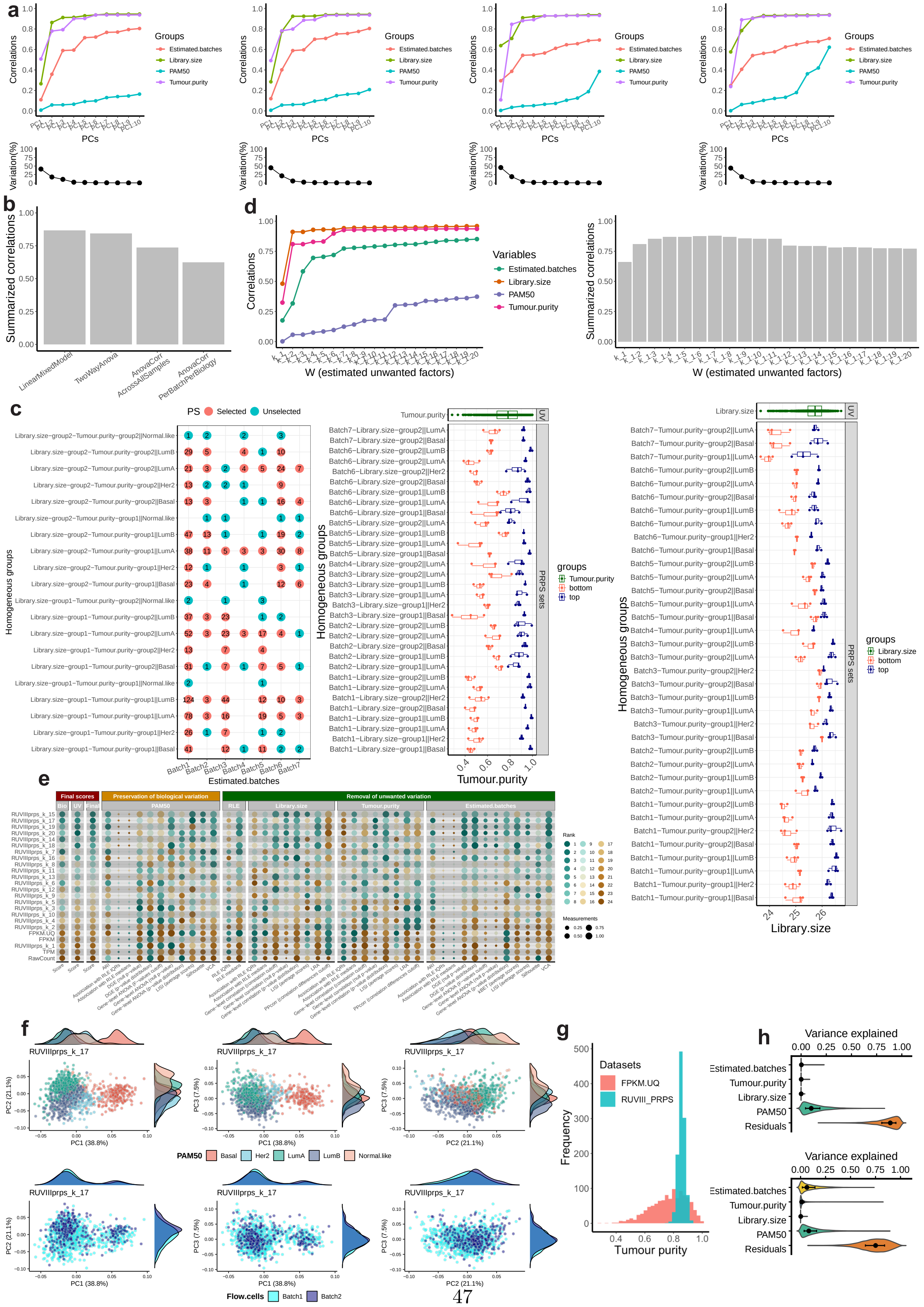
a
Correlations
PCs
Variation(%)
Groups
Estimated.batches
Library.size
PAM50
Tumour.purity
b
Summarized correlations
LinearMixedModel
TwoWayAnova
AnovaCorr AcrossAllSamples
AnovaCorr PerBatchPerBiology
d
Correlations
Variables
Estimated.batches
Library.size
PAM50
Tumour.purity
W (estimated unwanted factors)
Summarized correlations
W (estimated unwanted factors)
c
PS
Selected
Unselected
Homogeneous groups
Estimated.batches
Tumour.purity
Library.size
PRPS sets
UV
groups
Tumour.purity
Library.size
bottom
top
e
Final scores
Preservation of biological variation
Removal of unwanted variation
PAM50
RLE
Library.size
Tumour.purity
Estimated.batches
Rank
Measurements
f
RUVIIIprps_k_17
PC1 (38.8%)
PC2 (21.1%)
PC3 (7.5%)
PAM50
Basal
Her2
LumA
LumB
Normal.like
Flow.cells
Batch1
Batch2
g
Datasets
FPKM.UQ
RUVIII_PRPS
Frequency
Tumour purity
h
Variance explained
Estimated.batches
Tumour.purity
Library.size
PAM50
Residuals

Supplementary Figure 11: RUV-III normalization of the TCGA BRCA RNA-seq data for scenario 2. a) The line-dot plots show the performance of the selected normalization methods using different approaches including 'LinearMixedModel' (first plot),'TwoWayAnova' (second plot), 'AnovaCorr.AcrossAllSamples' (third plot), and 'AnovaCorr.PerBatchPerBiology' (fourth plot), on the TCGA BRCA raw count ($\log_2 + 0.5$)RNA-seq data. b) The barplot summarizes the performance of the selected NCG sets. The higher the correlation, the better the performance. c) The PRPS maps are shown for the batches of flow cell chemistry (first plot), library size (second plot) and tumor purity variables (third plot). For each PRPS map of a unwanted variable, the Y-axis displays all possible homogeneous groups with respect to the known biological variation and other unwanted variables. The X-axes represent the distributions of samples across the flow cell chemistry (batch 1 and batch 2), library size and tumor purity ranges, respectively. In the flow cell chemistry PRPS map, red indicates groups of samples containing more than two samples. Each horizontal line connecting two red points defines a PRPS set across the batches of flow cell chemistry. In the library size PRPS map, each horizontal line corresponds to six samples representing the highest and lowest library sizes within a homogeneous sample group. Every three samples are averaged to create a PS. The X-axis of the library size PRPS map shows the $\log_2 n$ of the library size in the data and highlights the samples selected for PS construction. In the tumor purity PRPS map, each horizontal line corresponds to six samples representing the highest and lowest tumor purity within a homogeneous sample group. Every three samples are averaged to create a PS. The X-axis shows the tumor purity estimate in the data and highlights the samples selected for PS construction. d) The line-dot plot shows the associations between the columns of the estimated W matrix and both biological and unwanted variables. The bar plot summarizes these correlations. A higher correlation the better performance. e) Numerical assessments of all metrics for both biological and unwanted variables are presented. Each point represents the performance of a given normalization method, where higher values indicate better performance. Normalized datasets are ranked according to their effectiveness in removing unwanted variation while preserving biological signals. f) The first three PC plots of RUV-III with $K = 17$ are shown, colored by PAM50 subtypes (top row) and the batches of flow cell chemistries (bottom row). The PCA were performed on a PAM50 subtypes gene signatures proposed by genefu R package. g) The plot shows the tumor purity estimated by the ESTIMATE method using the TCGA FPKM.UQ and the RUV-III-PRPS normalized data with $k = 17$. h) Shows the results of a linear mixed model analysis followed by gene-level variation analysis across all the variables for the TCGA FPKM.UQ (top plot) and the RUV-III-PRPS normalized data with $k = 17$ (bottom plot).

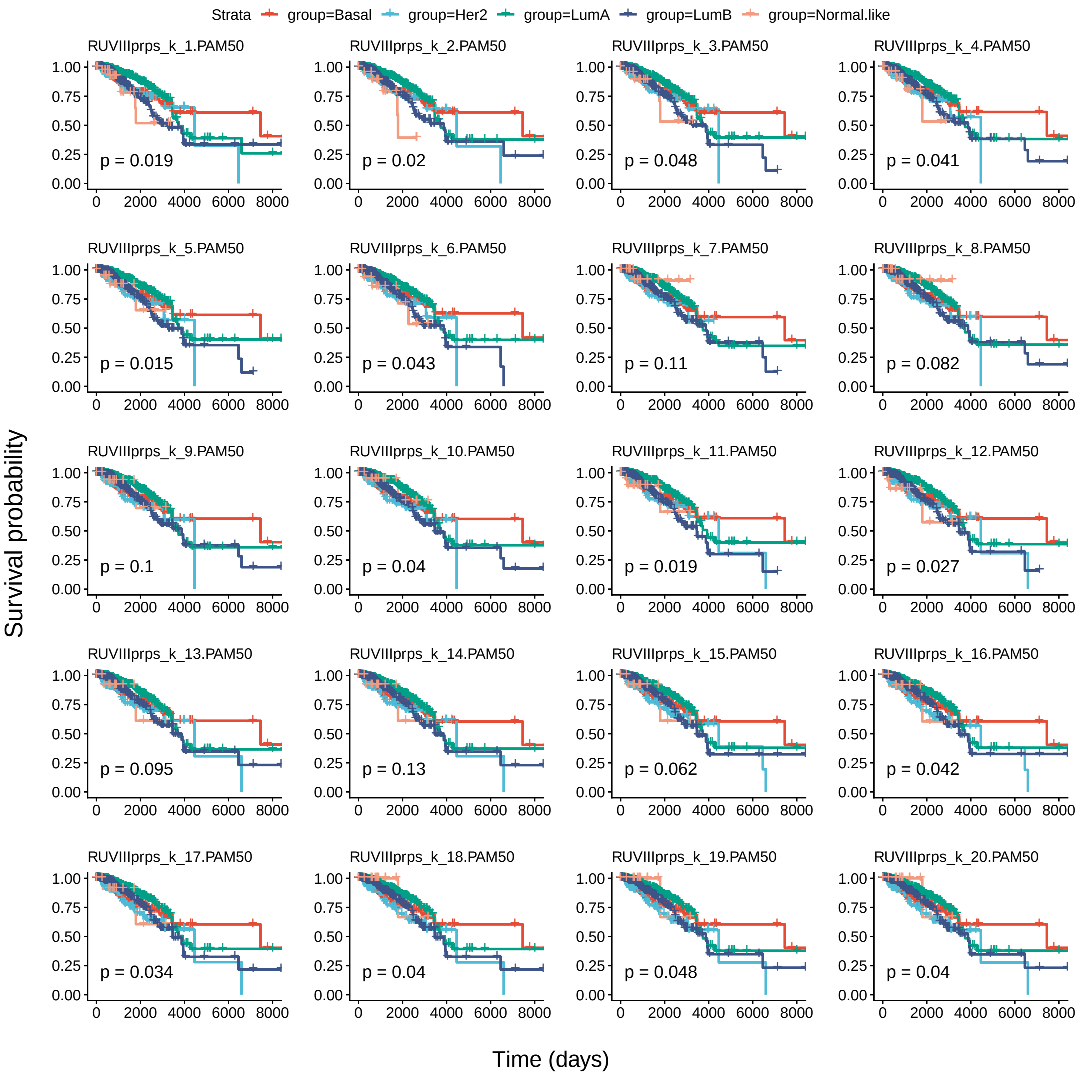


Supplementary Figure 12: Kaplan–Meier analysis of PAM50 estimated from each RUV-III–normalized dataset in scenario 2 and patient survival in the TCGA BRCA RNA-seq data.

#### Scenario 3: Known unwanted and unknown biological variation

In third scenario, it is assumed that the biological variation in the data is wholly unknown. Then, NCG and PRPS will then be identified and created in the unsupervised manner.

#### Step 1: Selecting negative control genes

A set of NCGs will be identified using the `findNcgsUnSupervised()` function. Library size, tumor purity and flow cell chemistries will be regarded as known sources of unwanted variation. Note that the PAM50 subtypes will be used solely to assess the performance of the NCG sets and will not be involved in the selection procedure.

```
for(i in c( 'AnovaCorr', 'LinearMixedModel')){
```

```
    brca.se.obj <- findNcgUnSupervised(
        se.obj = brca.se.obj,
        assay.name = 'RawCount',
        approach = i,
        use.rank = FALSE,
        ncg.selection.method = 'quantile',
        uv.variables = c('Flow.cells', 'Library.size', 'Tumour.purity'),
        form =  ~ (1 | Flow.cells) + Library.size + Tumour.purity,
        bio.percentile = NULL,
        uv.percentile = .8,
        nb.ncg = 0.05,
        variables.to.assess.ncg = c('Tumour.purity', 'PAM50', 'Flow.cells', 'Library.size'),
        ncg.group.name = 'brca.scenario3',
        ncg.set.name = i,
        nb.cores = 15,
        check.se.obj = FALSE)}
```

Supplementary Figures 13 a and b show that the selected genes by `LinearMixedModel` approach exhibits reasonable high correlation with the unwanted variation, and low correlation with known biological variation. This approach performed slightly better than `AnovaCorr` approach for this analysis.

**Step 2: Creating the PRPS data**

PRPS data will be created for individual sources of unwanted variation including library size, tumor purity and flow cell chemistries using the CCA approach in the `createPrPsUnSupervised()` function. First, a set of highly variable genes will be identified for CC-based dimensionality reduction to improve the identification of biologically similar samples across batches. The `findHVG()` function with the `lmm` approach will be used fo find HVG in the data.

```
hvg <- findHVG(
    se.obj = brca.se.obj,
    assay.name = 'RawCount',
    approach = 'lmm',
    uv.variables = c('Tumour.purity', 'Flow.cells', 'Library.size'),
    form = ~ (1 | Flow.cells) + Library.size + Tumour.purity,
    nb.hvg = .2,
    nb.cores = 15,
    save.se.obj = FALSE)
```

PCA analysis of the selected HVG colored by PAM50 subtypes showed a reasonable performance of the `findHVG()` function (Supplementary Figures 13d). We will set the maximum PRPS set argument to 10 to achieve a reasonable balance in the number of PRPS sets across sources of unwanted variation.

```
uv.variables <- c("Library.size", "Tumour.purity", "Flow.cells")
max.prps.sets <- c(10, 10, 10)
for(i in c(1:3)){
    brca.se.obj <- createPrPsUnSupervised(
        se.obj = brca.se.obj,
        assay.name = 'RawCount',
        uv.variables = uv.variables[i],
        other.uv.variables = NULL,
        select.extreme.groups = TRUE,
```

```
        approach = 'cca',
        nb.cca = 40,
        nb.pcs = 5,
        nb.clusters = 3,
        coordinates.to.use = 'cca',
        max.prps.sets = max.prps.sets[i],
        filter.prps.sets = TRUE,
        hvg = hvg,
        apply.log.for.prps = FALSE,
        create.prps.map = TRUE,
        prps.group.name = 'brca.scenario3')
```

Note that the `apply.log.for.prps()` argument was set to `FALSE`, indicating that the function used raw count data without log transformation to construct the PRPS data. In total, 34, 30, and 10 PR sets, corresponding to 68, 60, and 20 PS respectively, were generated for the removal of library size, tumor purity and flow cell chemistries effects (Supplementary Figure 13c).

##### Step 3: Performing RUV-III normalization with multiple $k$ values

The `getMaximumK()` function will first be applied to determine the maximum feasible value of $k$, which for the current NCG and PRPS data is 30. Subsequently, $k$ values ranging from 1 to 20 will be tested. RUV-III-PRPS normalization will be applied to the RawCount ($\log_2 + 0.5$) data, and all resulting RUV-III normalized datasets will be stored as new datasets in the *SummarizedExperiment* object.

```
brca.se.obj <- RUVIIIprps(
    se.obj = brca.se.obj,
    assay.name = 'RawCount',
    control.sample.types = 'prps',
    prps.type = 'un.supervised',
    prps.group  = 'brca.scenario3',
    prps.set.names = 'all',
    ncg.type = 'un.supervised',
    ncg.group.names = 'brca.scenario3',
    ncg.set.names = 'LinearMixedModel',
    k = c(1:20),
    apply.log = TRUE,
    data.to.log = 'both',
    pseudo.count = 0.5,
    return.wa = TRUE,
    check.se.obj = FALSE)
```

##### Step 4: Assessing the W matrix

The association between the estimated unwanted factors W from RUV-III and both biological and unwanted variables will be assessed by `assessW` function. Ideally, the columns of the estimated W should show strong associations with unwanted variables and no or weak associations with biological variables.

```
brca.se.obj <- assessW(
    se.obj = brca.se.obj,
    compare.w = TRUE,
```

```
    variables = NULL,
    bio.variables = 'PAM50',
    uv.variables = c('Library.size','Tumour.purity','Flow.cells'))
```

Supplementary Figure 13e shows that the unwanted factors W estimated by RUV-III are strongly associated with all sources of unwanted variation, while exhibiting relatively low association with the biological variable, particularly, the first 10 estimate factors . This analysis will be used to guide the selection of the optimal k value for RUV-III normalization.

**Step 5: Evaluating normalizations**

To assess the performance of RUV-III normalization and compare it with the TCGA normalizations, we will first apply the `assessVariation()` function, followed by the `assessNormalization()` function, across all RUV-III normalized data in the *SummarizedExperiment* object. Prior to variation assessment, the PAM50 subtypes will be estimated, and their association with patient survival will be evaluated using the RUV-III normalized data.

```
brca.se.obj <- estimatePAM50(
    se.obj = brca.se.obj,
    assay.names = names(SummarizedExperiment::assays(brca.se.obj))[-c(1:4)],
    entrez.gene.id = 'entrezgene_id',
    probe = 'gene_name',
    normalization = NULL,
    apply.log = FALSE,
    tissue.type = 'Tissues')

all.survivals <- lapply(
    pam50 <- grep('\\.PAM50$', colnames(colData(brca.se.obj)), value = TRUE),
    function(i) {
        idx <- brca.se.obj$OS != -1
        computeSurvival(
            se.obj = brca.se.obj[, idx],
            variable = i,
            survival.time = 'OS.time',
            survival.events = 'OS',
            genes = NULL,
            return.survival.plot = TRUE,
            check.se.obj = FALSE,
            save.se.obj = FALSE)$plot })
```

Supplementary Figure 15 shows the result of the PAM50 survival analysis for all the RUV-III-PRPS normalized data.

```
brca.se.obj$PAM50 <- brca.se.obj$FPKM.UQ.PAM50
brca.se.obj <- assessVariation(
    se.obj = brca.se.obj,
    assay.names = 'all',
    assessment.level = 'L2',
    bio.variables = c("PAM50"),
    uv.variables = c('Library.size','Tumour.purity','Flow.cells'),
    pcorr.genes = ppcorr.gene.sets,
    gene.set.score.list = tumour.purity.genes.sets,
    pcorr.filter.genes = FALSE,
    general.points.size = 3,
    override.check = TRUE,
```

```
    apply.log = FALSE,
    check.se.obj = FALSE)

brca.se.obj <- assessNormalization(
    se.obj = brca.se.obj,
    assay.names = 'all',
    assessment.level = 'L2',
    select.top.ruv = FALSE,
    bio.variables = c("PAM50"),
    uv.variables = c('Library.size','Tumour.purity','Flow.cells'),
    bio.weight = 0.6,
    uv.weight = 0.4,
    corr.cutoff = list(Tumour.purity = .5, Library.size = .3),
    output.name = 'brca.read.assessNormalization.scenario3')

RUVprps:::plotAssessVariation(
    se.obj = brca.se.obj,
    assay.names = 'all',
    variables = c("Tumour.purity", "PAM50", "Library.size", "Flow.cells"),
    output.file.name = "RUVprps_TCGA_BRCA_VariationAssessment_Scenario3")
```

Supplementary Figure 13f shows that RUV-III with PRPS achieved effective normalization of the TCGA BRCA RNA-seq data. Quantitative evaluation of normalization performance indicated that RUV-III-PRPS with $k = 9$ yielded the best overall numerical performance. However, considering the results of the PAM50 survival analysis (Supplementary Figure 15), RUV-III-PRPS with $k = 15$ was selected as the most appropriate normalized dataset. The first three PCs of the RUV-III-PRPS–normalized data with $k = 15$ show biologically expected separation of the PAM50 subtypes while effectively removing the impact of flow cell chemistry (Supplementary Figure 13g). The PCA was performed using a PAM50 gene signature provided by the genefu R package. Supplementary Figure 13h shows that variation associated with tumor purity is reduced following RUV-III-PRPS normalization. Results from linear mixed-effects model analysis, followed by gene-level variance decomposition across all variables, further demonstrated that RUV-III-PRPS normalization removes variation attributable to unwanted sources while preserving variation associated with the PAM50 subtypes. The Supplementary File "TCGA_BRCA_variationAssessment_Scenario3.pdf" provides all plots summarized in Supplementary Figure 13f. Finally, results of the PAM50 survival analysis showed that RUV-III-PRPS normalization with $k = 15$ was significantly associated with survival (Supplementary Figure 15).

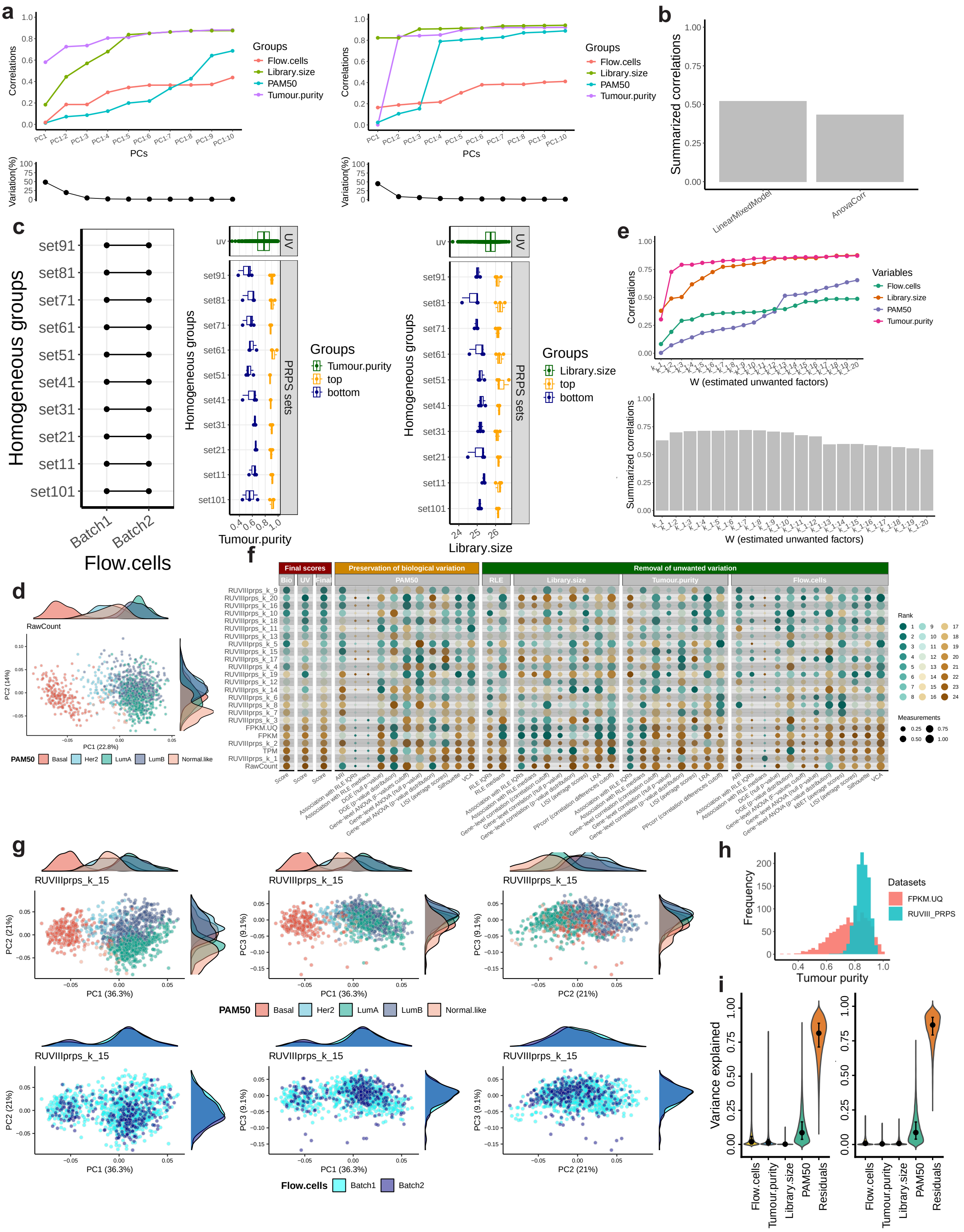
a
Correlations
PCs
Groups
Flow.cells
Library.size
PAM50
Tumour.purity
Variation(%)
b
Summarized correlations
LinearMixedModel
AnovaCorr
c
Homogeneous groups
Batch1
Batch2
Flow.cells
uv
UV
PRPS sets
Tumour.purity
top
bottom
Library.size
d
RawCount
PC1 (22.8%)
PC2 (14%)
PAM50
Basal
Her2
LumA
LumB
Normal.like
e
W (estimated unwanted factors)
Variables
f
Final scores
Preservation of biological variation
Removal of unwanted variation
Rank
Measurements
g
RUVIIIprps_k_15
PC1 (36.3%)
PC2 (21%)
PC3 (9.1%)
Batch1
Batch2
h
Frequency
Tumour purity
Datasets
FPKM.UQ
RUVIII_PRPS
i
Variance explained
Residuals

Supplementary Figure 13: RUV-III normalization of the TCGA BRCA RNA-seq data for scenario 3. a) The line-dot plots show the performance of the selected NCGs using different approaches including `LinearMixedModel` (first plot), and `AnovaCorr` (second plot), on the TCGA BRCA raw count ($\log_2 + 0.5$)RNA-seq data. b) The barplot summarizes the performance of the selected NCG sets. The higher the correlation, the better the performance. c) The PRPS maps are shown for the batches of flow cell chemistry (first plot), library size (second plot) and tumor purity variables (third plot). For each PRPS map of a unwanted variable, the Y-axis displays all possible homogeneous groups found in CCA-based unsupervised manner. The X-axes represent the distributions of samples across the flow cell chemistry (batch 1 and batch 2), library size and tumor purity ranges, respectively. In the flow cell chemistry PRPS map, each horizontal line connecting two black points defines a PRPS set across the batches of flow cell chemistry. In the library size PRPS map, each horizontal line corresponds to six samples representing the highest and lowest library sizes within a homogeneous sample group. Every three samples are averaged to create a PS. The X-axis of the library size PRPS map shows the $\log_2 n$ of the library size in the data and highlights the samples selected for PS construction. In the tumor purity PRPS map, each horizontal line corresponds to six samples representing the highest and lowest tumor purity within a homogeneous sample group. Every three samples are averaged to create a PS. The X-axis shows the tumor purity estimate in the data and highlights the samples selected for PS construction. d) The firs two PCs of the TCGA BRCA raw count data ($\log_2 + 0.5$) using the selected HVGs, colored by the PAM50 subtypes. e) The line-dot plot shows the associations between the columns of the estimated W matrix and both biological and unwanted variables. The bar plot summarizes these correlations. A higher correlation the better performance. f) Numerical assessments of all metrics for both biological and unwanted variables are presented. Each point represents the performance of a given normalization method, where higher values indicate better performance. Normalized datasets are ranked according to their effectiveness in removing unwanted variation while preserving biological signals. g) The first three PC plots of RUV-III with $K = 15$ are shown, colored by PAM50 subtypes (top row) and the batches of flow cell chemistries (bottom row). The PCA were performed on a PAM50 subtypes gene signatures proposed by the genefu R package. h) The plot shows the tumor purity estimated by the ESTIMATE method using the TCGA FPKM.UQ and the RUV-III-PRPS normalized data with $k = 15$. h) Shows results from linear mixed-effects model analysis followed by gene-level variance decomposition across all variables for the TCGA FPKM-UQ data (top) and the RUV-III-PRPS–normalized data with $k = 15$ (bottom).

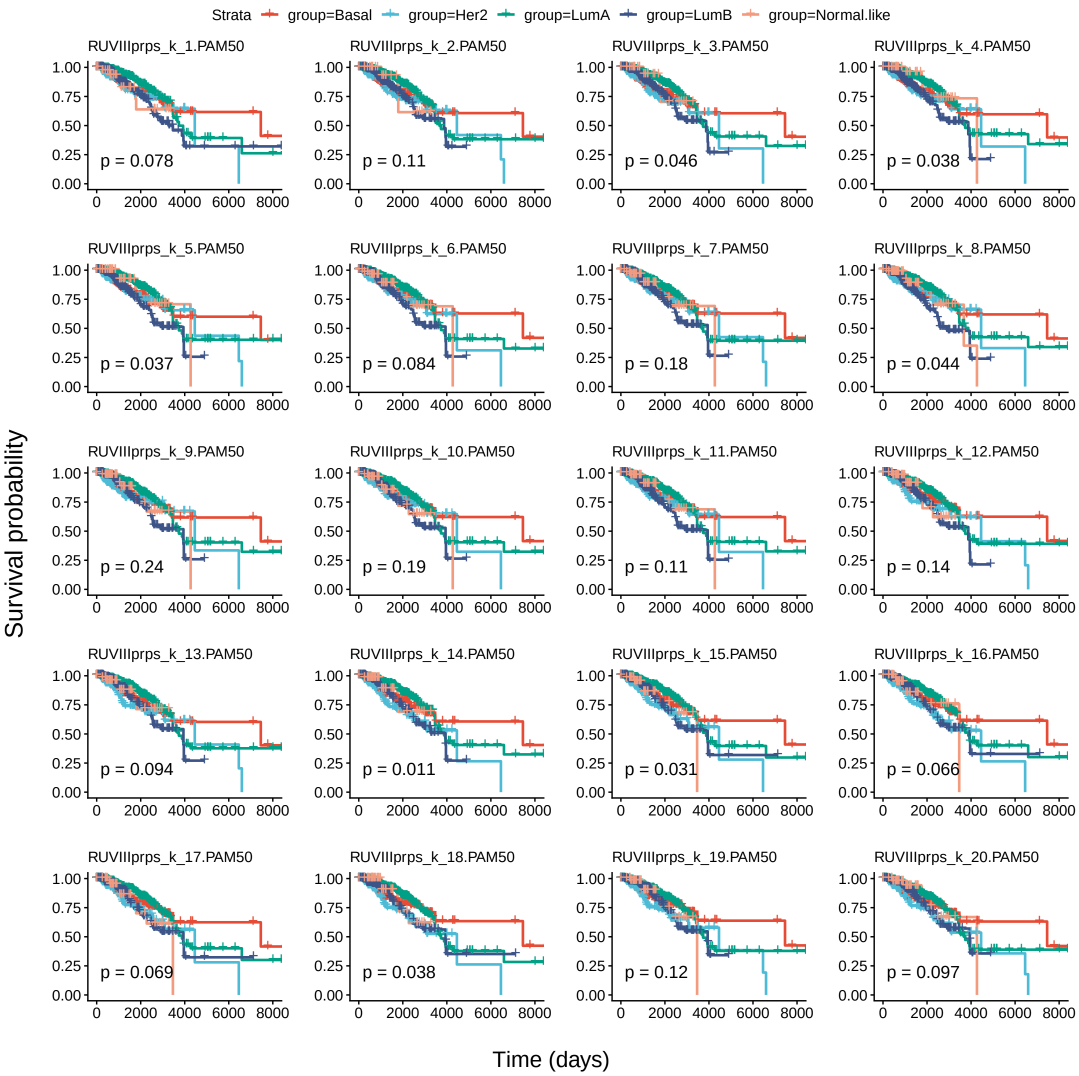


Supplementary Figure 14: Kaplan–Meier analysis of PAM50 estimated from each RUV-III–normalized dataset in scenario 3 and patient survival in the TCGA BRCA RNA-seq data.

#### Scenario 4: Both the unwanted and biological variation unknown

In this scenario, it is assumed that both biological and unwanted variation in the data are wholly unknown. Both NCG and PRPS will be identified in the unsupervised manner.

#### Step 1: Selecting negative control genes

A set of NCGs will be identified using the `findNcgsUnSupervised()` function. Both unsupervised approaches will be tested and the results will be compared to select a true set NCG. For this analysis, library size, tumor purity and estimated batches will be considered as sources of unwanted variation in the data. As we explain before, the PAM50 subtypes will be used solely to assess the performance of the different sets of NCG.

```
for(i in c( 'AnovaCorr', 'LinearMixedModel')){
    brca.se.obj <- findNcgUnSupervised(
        se.obj = brca.se.obj,
        assay.name = 'RawCount',
        approach = i,
        use.rank = FALSE,
        ncg.selection.method = 'quantile',
        uv.variables = c('Estimated.batches', 'Library.size', 'Tumour.purity'),
        form =  ~ (1 | Estimated.batches) + Library.size + Tumour.purity,
        bio.percentile = NULL,
        uv.percentile = .8,
        nb.ncg = 0.05,
        variables.to.assess.ncg = c('Tumour.purity', 'PAM50', 'Estimated.batches', 'Library.size'),
        ncg.group.name = 'brca.scenario4',
        ncg.set.name = i,
        nb.cores = 15,
        check.se.obj = FALSE)}

brca.se.obj <- compareNCGs(
    se.obj = brca.se.obj,
    assay.name = 'RawCount',
    ncg.set.names = "all",
    bio.variables = c('PAM50'),
    uv.variables =  c('Library.size','Tumour.purity','Estimated.batches'),
    nb.pcs = 10,
    ncg.type = 'un.supervised',
    ncg.group.name = 'brca.scenario4',
    check.se.obj = FALSE)
```

Figure 5 a and b show that the selected genes by `LinearMixedModel` approach exhibits reasonable high correlation with the unwanted variation, and low correlation with known biological variation. Further, this approach is slightly better than the other approach for this analysis.

**Step 2: Creating the PRPS data**

PRPS data will be created for individual sores of of unwanted variation including library size, tumor purity and estimated batches using the `createPrPsUnSupervised()` function. First,a set of highly variable genes will be identified using the `findHVG` function. This will improve the identification biological similar samples across batches by CCA. Note that different values will be used for the maximum PRPS set size for individual sources in order to maintain a reasonable balance of PRPS across sources of unwanted variation.

```
# finding the highly variable genes
hvg <- findHVG(
    se.obj = brca.se.obj,
    assay.name = 'RawCount',
    approach = 'lmm',
    uv.variables = c('Tumour.purity', 'Estimated.batches', 'Library.size'),
    form = ~ (1 | Estimated.batches) + Library.size + Tumour.purity,
    nb.hvg = .2,
    nb.cores = 15,
```

```
    save.se.obj = FALSE)

uv.variables <- c("Library.size", "Tumour.purity", "Estimated.batches")
max.prps.sets <- c(10, 10, 1)
for(i in c(1:3)){
    brca.se.obj <- createPrPsUnSupervised(
        se.obj = brca.se.obj,
        assay.name = 'RawCount',
        uv.variables = uv.variables[i],
        other.uv.variables = NULL,
        select.extreme.groups = TRUE,
        approach = 'cca',
        nb.cca = 4,
        nb.pcs = 5,
        clustering.method = 'quantile',
        nb.clusters = 4,
        coordinates.to.use = 'cca',
        max.prps.sets = max.prps.sets[i],
        filter.prps.sets = TRUE,
        hvg = hvg,
        apply.log.for.prps = FALSE,
        create.prps.map = TRUE,
        prps.group.name = 'brca.scenario4')}
```

In total, 10, 10, and 21 PR sets, corresponding to 20, 20, and 42 PS respectively, were generated for the removal of library size, tumor purity and estimated batches (Figure 1d).

**Step 3: Performing RUV-III normalization with multiple $k$ values**

For the current NCG and PRPS data, the possible maximum value for $K$ is 41. We will apply RUV-III-PRPS with $K$ values ranging from 1 to 20. RUV-III-PRPS normalization will be applied to the RawCount ($\log_2$ +.5) data, and all RUV-III normalized datasets will be stored as new datasest in the *SummarizedExperiment* object.

```
brca.se.obj <- RUVIIIprps(
    se.obj = brca.se.obj,
    assay.name = 'RawCount',
    control.sample.types = 'prps',
    prps.type = 'un.supervised',
    prps.group  = 'brca.scenario4',
    prps.set.names = 'all',
    ncg.type = 'un.supervised',
    ncg.group.names = 'brca.scenario4',
    ncg.set.names = 'LinearMixedModel',
    k = c(1:20),
    apply.log = TRUE,
    data.to.log = 'both',
    pseudo.count = 0.5,
    return.wa = TRUE,
    check.se.obj = FALSE)
```

**Step 4: Assessing the W matrix**

Similar to other scenarios, we will assess how the estimated unwanted factors W from RUV-III relate to both biological and unwanted variables.).

```
brca.se.obj <- assessW(
    se.obj = brca.se.obj,
    compare.w = TRUE,
    variables = NULL,
    bio.variables = 'PAM50',
    uv.variables = c('Library.size','Tumour.purity','Estimated.batches'))
```

Figure 1f demonstrates strong performance of the unsupervised RUV-III-PRPS normalization. The unwanted factors W estimated by RUV-III are strongly associated with multiple sources of unwanted variation while exhibiting minimal association with the biological variable. This analysis will be used to guide the selection of an optimal value of k for RUV-III normalization.

**Step 5: Evaluating normalizations**

For individual RUV-III-PRPS normalized data, the PAM50 subtypes will be estimated, and their association with patient survival will be evaluated.

```
brca.se.obj <- estimatePAM50(
    se.obj = brca.se.obj,
    assay.names = names(SummarizedExperiment::assays(brca.se.obj))[-c(1:4)],
    entrez.gene.id = 'entrezgene_id',
    probe = 'gene_name',
    normalization = NULL,
    apply.log = FALSE,
    tissue.type = 'Tissues')

all.survivals <- lapply(
    pam50 <- grep('\\.PAM50$', colnames(colData(brca.se.obj)), value = TRUE),
    function(i) {
        idx <- brca.se.obj$OS != -1
        computeSurvival(
            se.obj = brca.se.obj[, idx],
            variable = i,
            survival.time = 'OS.time',
            survival.events = 'OS',
            genes = NULL,
            return.survival.plot = TRUE,
            check.se.obj = FALSE,
            save.se.obj = FALSE)$plot })
```

We next assessed variation associated with all biological and unwanted variables in the RUV-III-PRPS–normalized data. Their performance will be then evaluated and compared to the TCGA normalized data. Note that the PAM50 survival analysis showed that RUV-III-PRPS normalization with $k = 20$ performed better than the TCGA FPKM-UQ normalization. Therefore, for this analysis, we will use PAM50 subtype calls derived from the RUV-III-PRPS–normalized data to assess variation and normalizations.

```
brca.se.obj$PAM50 <- brca.se.obj$RUVIIIprps_k_20.PAM50
```

```
brca.se.obj <- assessVariation(
    se.obj = brca.se.obj,
    assay.names = 'all',
    assessment.level = 'L2',
    bio.variables = c("PAM50"),
    uv.variables = c('Library.size','Tumour.purity','Estimated.batches'),
    pcorr.genes = ppcorr.gene.sets,
    gene.set.score.list = tumour.purity.genes.sets,
    pcorr.filter.genes = FALSE,
    general.points.size = 3,
    override.check = TRUE,
    apply.log = FALSE,
    check.se.obj = FALSE)

brca.se.obj <- assessNormalization(
    se.obj = brca.se.obj,
    assay.names = 'all',
    assessment.level = 'L1',
    select.top.ruv = FALSE,
    bio.variables = c("PAM50"),
    uv.variables = c('Library.size','Tumour.purity','Estimated.batches'),
    bio.weight = 0.6,
    uv.weight = 0.4,
    corr.cutoff = list(Tumour.purity = .5, Library.size = .3),
    output.name = 'brca.read.assessNormalization.scenario4'
    )
RUVprps:::plotAssessVariation(
    se.obj = brca.se.obj,
    assay.names = 'all',
    variables = c("Tumour.purity", "PAM50", "Library.size", "Estimated.batches"),
    output.file.name = "RUVprps_TCGA_BRCA_VariationAssessment_Scenario4")
```

Comprehensive details of the normalization performance assessment are provided in the main text and Fig. 5. The Supplementary File "TCGA_BRCA_variationAssessment_Scenario4.pdf" contains all plots from the normalization performance assessment.

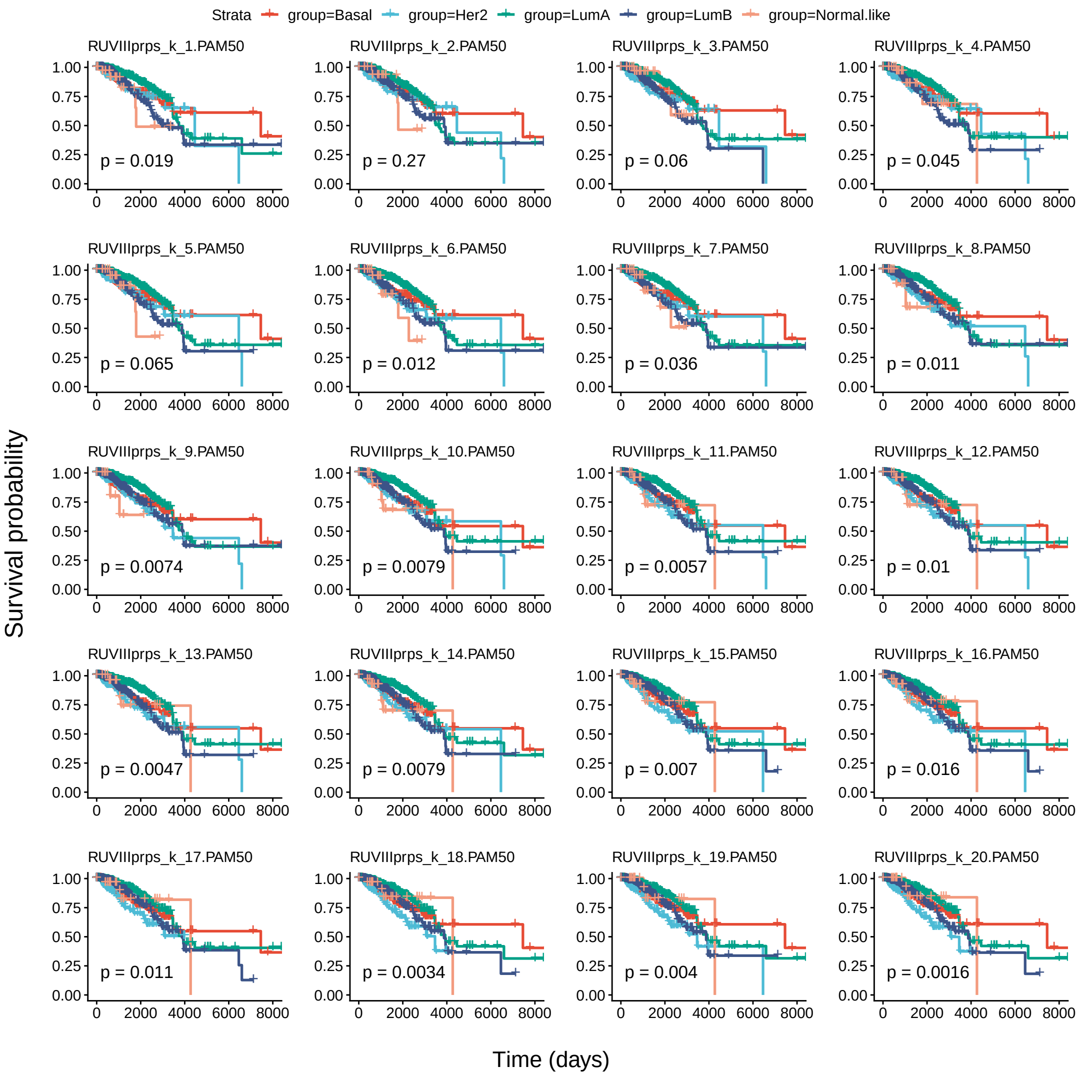


Supplementary Figure 15: Kaplan–Meier analysis of PAM50 estimated from each RUV-III–normalized dataset in scenario 4 and patient survival in the TCGA BRCA RNA-seq data.

### RUV-III normalization of the TCGA READ RNA-seq data

In this procedure, our goal is to preserve the variation associated with tumor purity alongside other known biological variation. We selected the TCGA rectum adenocarcinoma (READ) RNA-seq data for this purpose, as it is significantly influenced by library size and batch effects [1]. It should be noted that there is complete confounding between normal tissues and the batches in the data, which will be explained in the subsequent sections. Normalizing RNA-seq data with such confounding factors proves challenging.

**Required R packages and installations**

The R package below is required to be installed for this procedure.

The TCGAbiolinks R/Bioconductor package (version 2.34.0) will be used to download the TCGA RNA-seq data from the Genomic Data Commons (GDC) data repository.

```
if (!require("BiocManager", quietly = TRUE))
    install.packages("BiocManager")
BiocManager::install("TCGAbiolinks")
```

**Stage 1: Data preparation and processing**

Data preparation and processing procedures may vary across individual studies. In this stage, these steps are specifically tailored for RNA-seq data obtained from TCGA.

**Step 1: Downloading the data**

The most recent TCGA READ RNA-seq data will be obtained from the GDC data portal using the TCGAbiolinks R/Bioconductor package. The required files will be downloaded to the current home directory.

```
tcga.read.rnaseq.data <- TCGAbiolinks::GDCquery(
    project = 'TCGA-READ',
    data.category = 'Transcriptome Profiling',
    data.type = 'Gene Expression Quantification'
    )
TCGAbiolinks::GDCdownload(tcga.read.rnaseq.data)
read.se.obj <- TCGAbiolinks::GDCprepare(query = tcga.read.rnaseq.data)
```

The `read.se.obj` is a *RangedSummarizedExperiment* object with 60,660 Ensemble transcripts and 177 RNA-seq samples. This object is deposited on the Zenodo website for reproducibility [i will add a link once the MS is finalized]. The object contains 6 datasets (assays) including `unstranded`, `stranded_first`, `stranded_second`, `tpm_unstrand`, `fpkm_unstrand` and `fpkm_uq_unstrand`. The `unstranded`, `stranded_first` and `stranded_second` are raw count data without any transformation. The `removeAssays()` function will be used to exclude the `stranded_first`, `stranded_second` and the `unstranded` data will be used as raw count data for RUV-III normalization. The reason for this is that TCGA normalizations, including TPM, FPKM, and FPKM.UQ, were performed on the unstranded data. Note that the names of the datasets in the object will be replaced with `RawCount`, `TPM`, `FPKM` and `FPKM.UQ`.

```
read.se.obj <- RUVprps:::removeAssays(
    se.obj = read.se.obj,
    assays.to.remove = c('stranded_first', 'stranded_second'))
# changing the names of the datasets
read.se.obj <- RUVprps:::renameAssays(
    se.obj = read.se.obj,
    new.names = c('RawCount', 'TPM', 'FPKM', 'FPKM.UQ'))
```

**Step 2: Selecting the gene types of interest (optional)**

The `gene_type` column in the gene annotation (rowData) of the *RangedSummarizedExperiment* object classifies Ensembl transcripts into various gene type categories, such as `protein_coding`, `lncRNA` (long non-coding RNA), among others. For this analysis, only unique Ensembl transcripts annotated as "protein_coding" will be retained using the `tidyGenes()` function. Additionally, the row names of the *RangedSummarizedExperiment* object will be updated to display gene symbols instead of Ensembl transcript IDs for down-stream analysis.

```
read.se.obj <- tidyGenes(
    se.obj = read.se.obj,
    keep.gene.type = 'protein_coding',
    gene.type.col.name = 'gene_type',
    remove.duplicates.ids = TRUE,
    ids.col.name = 'gene_name',
    change.row.names = TRUE,
    new.row.names = 'gene_name')
```

Then a total of 19,934 genes were retained for subsequent steps. It should be noted that any gene type groups of interest can be kept and used for RUV-III normalization.

**Step 3: Adding batch details**

TCGA RNA-seq samples were collected from various tissue source sites (TSS), shipped to different genomics profiling centers over time, and profiled using 96-sequencing plates. These are known sources of batch information that can impact down-stream analysis [1]. The `addTcgaBatchInfo()` function will be employed to annotate the TCGA READ RNA-seq data with all available batch information. This function retrieves batch annotations from the file "41587_2022_1440_MOESM3_ESM.xlsx", available from our previous study [1]. Finally, the `orderSeObj()` function will be applied to order the samples chronologically based on the batch information, in preparation for down-stream analysis.

```
# adding TCGA RNA-seq batch information
read.se.obj <- addTcgaBatchInfo(se.obj = read.se.obj)

# re-ordering the object by variables
read.se.obj <- orderSeObj(
    se.obj = read.se.obj,
    factors.to.order = c('Years','Months','Days','Plates','TSS','Center'))
read.se.obj$Years <- as.factor(x = read.se.obj$Years)
```

All available batch information were added to the sample annotation of the *RangedSummarizedExperiment* object. The TCGA READ RNA-seq data involved 10 normal (adjacent tissue) and 167 cancer tissue samples that were collected from 13 TSS, distributed across 15 sequencing plates, and profiled over the 4 years 2010, 2012, 2013 and 2014 (Supplementary Figure 16a).

**Step 4: Preparing the SummarizedExperiment object**

The `prepareSeObj()` function will be utilized to perform several preprocessing steps, including the identification and removal of lowly expressed genes, calculation of library sizes, estimation of tumor purity, integration of multiple publicly available housekeeping gene lists, and stromal and immune gene signatures. For these tasks, the `RawCount` data will be used for library size calculation, while FPKM.UQ will be employed for tumor purity estimation. If a gene annotation file is provided, it must contain at least one column with a name that includes hgnc_symbol, entrezgene_id, or ensembl_gene_id to enable the mapping of gene details and incorporation of housekeeping gene lists. Furthermore, the row names of the *SummarizedExperiment* object must correspond to the identifiers in the specified column. In this analysis, the gene annotation associated with the read.se.obj object includes a column named gene_name; therefore, the function will generate a new column labeled hgnc_symbol accordingly.

```
read.se.obj <- prepareSeObj(
    data = read.se.obj,
    raw.count.assay.name = 'RawCount',
    remove.lowly.expressed.genes = TRUE,
    assay.name.to.estimate.purity = 'FPKM.UQ',
    estimate.tumor.purity = 'both',
    scale.singscore.values = TRUE,
    calculate.library.size = TRUE,
    add.immun.stroma.genes = TRUE,
    add.gene.details = FALSE,
    add.housekeeping.genes = TRUE,
    column.name = 'gene_name',
    gene.group = 'hgnc_symbol')
```

14,799 of 19,934 genes were kept as highly expressed genes. Tumor purity was estimated using both the ESTIMATE and singscore methods [2, 6]. These estimates are highly correlated (Supplementary Figure 16b), and the ESTIMATE values will be used for down-stream analyses. The tumor purity estimated by the singscore method was re-scaled based on the tumor purity estimated by the ESTIMATE method. This adjustment is necessary because the scores from the singscore method exceed 1, whereas the ESTIMATE method provides tumor purity estimates within a more conventional range which is between 0 and 1.

##### Step 5: Visualizing the study outline (optional)

The plotStudyOutline() function can be applied to visualize how various variables are distributed across the samples. In addition, this can assess the association between all specified variables. This plot can be updated with more variables, such as cancer subtypes once they are calculated throughout the procedure. Before creating a study outline plot, the renameVariables() function will be used to rename the variables for visualization purposes.

```
# re-naming the columns just for visualization purposes
read.se.obj <- renameVariables(
    se.obj = read.se.obj,
    current.names = c('tissue_type', 'tumour.purity.estimate', 'library.size'),
    new.names = c('Tissues', 'Tumour.purity', 'Library.size'))

# generating study outline plot
read.se.obj <- plotStudyOutline(
    se.obj = read.se.obj,
    variables = c('Years', 'Plates', 'TSS', 'Library.size', 'Tumour.purity', 'Tissues'))
```

The output of the plotStudyOutline() function shows that the sample library sizes vary greatly between samples profiled in 2010 and the other samples (Supplementary Figure 16a). The experimental conditions that may explain this variation are not publicly available. All adjacent normal tissues were profiled in 2013-2014, indicating the presence of substantial confounding in the data. Supplementary Figure 16c shows the Plates, Years and library size are highly correlated with each other. Furthermore, the tumor purity estimates show no association with library size or known batch effects.

The study outline plot reveals substantial variation in library size across different years. To account for this, the years will be grouped into two intervals, 2010 and 2011–2014, for downstream analysis. These intervals will be named as `Batch1` and `Batch2`, respectively.

```
read.se.obj[['Time.interval']] <- factor(
  ifelse(read.se.obj$Years == 2010, 'Batch1', 'Batch2'),
  levels = c('Batch1', 'Batch2'))
```

##### Stage 2: Identification of major variation

In situations where all sources of unwanted variation and the main gene expression-based biological variation are already known or estimated using suitable orthogonal platforms or methods, this stage becomes unnecessary. It is crucial to emphasize that any significant biological variables should be discernible in the gene expression data. Variables not evident in the data are generally not applicable to the RUV-III normalization procedure. For instance, if there is no true association between gene expression and gender, including gender as a factor will not aid in normalization. However, these variables can be useful for normalization assessments. If genes show associations with those variables, this might be an indication of the presence of unwanted variation in the data.

Identification of major biological populations is not the primary focus of the RUVprps. Users may apply any appropriate tools or approaches to estimate biological variation according to their study objectives.

##### Step 1: Biological populations

To implement RUV-III with PRPS in a supervised manner and assess the performance of normalizations, it is essential to identify all or major gene expression-based biological populations within the data. This step is particularly challenging for the TCGA READ RNA-seq data due to the substantial impact of library size and batches on both the raw and TCGA normalized datasets [1]. To address this challenge, we suggest two strategies and evaluate both to ascertain the most reliable estimates of the biological populations in the data.

Colorectal cancers can be classified into four major gene expression–based biological subtypes, known as consensus molecular subtypes (CMS) [10]. These subtypes have been shown to correlate with clinical outcomes, including overall patient survival. Specifically, patients classified as CMS4 exhibit the poorest overall survival, whereas those with the CMS2 subtype are associated with more favorable survival outcomes. [10]. RUVprps provides a wrapper function called `estimateCMS()` that leverages the CMScaller R package [13] to assign CMS classifications to samples. The following section outlines our approach for CMS identification and assessment.

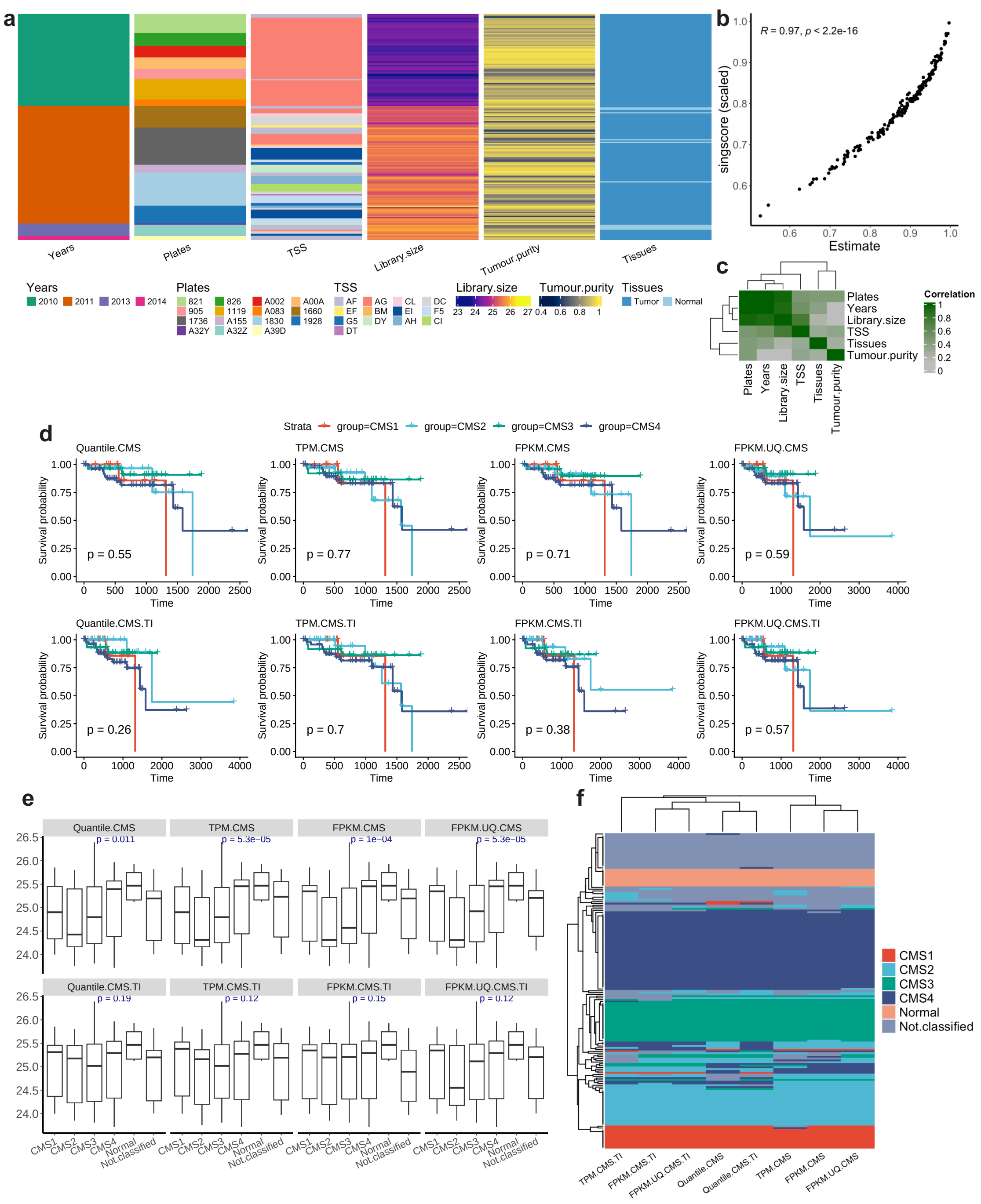


Supplementary Figure 16: Study outline and consensus molecular subtype (CMS) estimates of the TCGA READ RNA-seq data. a) The heatmap shows 176 rectum adenocarcinoma and adjacent normal tissues that were collected from 13 tissue source sites (TSS) and distributed across 15 sequencing plates for profiling at 4 time points over a span of 4 years. The library sizes were calculated using the raw counts after removing lowly expressed genes. Tumor purity was estimated using the ESTIMATE method on the TCGA FPKM.UQ normalized data. b) The scatter plot shows the correlation between estimated tumor purity using the ESTIMATE and singscore methods. c) The heatmap shows the canonical correlation between all the variables in the data. d) Kaplan-Meier plots show the association between the estimated CMSs and the survival outcomes of patients for individual TCGA datasets using different strategies. The first row shows the CMS estimates obtained from strategy 1, and the second row shows the estimates from strategy 2. e) Shows the distribution of library size within each estimated CMS. One-way ANOVA was used to assess the association. f) The heatmap shows the estimated CMS for each TCGA READ RNA-seq dataset using different strategies.

*Strategy 1.* The CMS classifier is applied across all the cancer samples using the TCGA raw count and normalized data. The classifier applies quantile normalization to the raw count data before estimating the CMS. All the estimated CMS will be added to the sample annotation of the *SummarizedExperiment* object for down-stream analysis. Note that, the normal tissues will be removed before applying the classifier.

```
read.se.obj <- estimateCMS(
    se.obj = read.se.obj,
    assay.names = 'all',
    raw.count.assay.name = 'RawCount',
    subtype = 'CMS',
    tissue.type =  'Tissues',
    groups = NULL,
    templates = CMScaller::templates.CMS,
    seed = 122334,
    apply.log = TRUE,
    nb.cores = 1)
```

*Strategy 2.* The CMS classifier will be applied within the major time intervals (2010 or Batch1 and 2011-2014 or Batch2). This adjustment aims to mitigate possible influence of library size and batches on the CMS classifier.

```
read.se.obj <- estimateCMS(
    se.obj = read.se.obj,
    assay.names = 'all',
    raw.count.assay.name = 'RawCount',
    subtype = 'CMS',
    templates = CMScaller::templates.CMS,
    tissue.type =  'Tissues',
    groups = 'Time.interval',
    out.put.name = 'TI',
    apply.log = TRUE,
    seed = 12244,
    nb.cores = 1)
```

*Evaluating of the estimated CMS.* Two analyses will be conducted to evaluate the quality of the estimated CMS for the individual datasets. First, the association between the derived subtypes and patient overall survival will be assessed. Second, the relationship between the estimated subtypes and library size will be examined. An ideal estimate of CMS should have significant association with survival data and no or weak association with library size.

RUVprps provides a function, `AddTcgaClinicInfo()`, which allows users to incorporate specified TCGA clinical data from Liu et al. [12] into a *SummarizedExperiment* object. The clinical data, available as Supplementary Table 1 ("mmc1.xlsx") from that study. In this analysis, overall survival data for the TCGA READ RNA-seq dataset will be extracted and incorporated to the *SummarizedExperiment* object. The function assigns a value of -1 to the overall survival time and events for normal tissue samples to facilitate their exclusion from downstream survival analysis. Subsequently, the `computeSurvival()` function will be employed to generate Kaplan–Meier survival curves for each CMS classification across the differently normalized TCGA datasets. Note that normal samples and those that could not be classified into any CMS subtype will be excluded prior to performing the survival analysis.

```
# adding overall survival details
read.se.obj <- addTcgaClinicalInfo(
    se.obj = read.se.obj,
```

```
    factors = c("OS", "OS.time"),
    tissue.type = 'Tissues')

# obtaining the Kaplan-Meier plots
all.survivals <- lapply(
    cms <- grep('CMS', colnames(colData(read.se.obj)), value = TRUE),
    function(i) {
        idx <- read.se.obj$OS != -1 & read.se.obj[[i]] != 'Not.classified'
        computeSurvival(
            se.obj = read.se.obj[, idx],
            variable = i,
            survival.time = 'OS.time',
            survival.events = 'OS',
            genes = NULL,
            return.survival.plot = TRUE,
            check.se.obj = FALSE,
            save.se.obj = FALSE)$plot})
```

Supplementary Figure 16d indicates that the CMS derived from the Quantile normalized data within the time intervals are more associated with patients' overall survival compared to the other normalized datasets.

The `plotVariables()` function will be used to assess the association between the CMS estimates and library size. Further, the function will generate a heatmap of all the estimated CMS to explore the differences between different datasets and strategies.

```
# generating a boxplot between the library size and the estimated CMS
plotVariables(
    se.obj = read.se.obj,
    x.variables = grep('CMS', colnames(colData(read.se.obj)), value = TRUE),
    y.variables = 'Library.size',
    generate.heatmap = TRUE)
```

Supplementary Figure 16e shows that the CMS subtypes obtained in the strategy 2 are not significantly associated with library size. As a result, the `Quantile.TI` (TI stands for time interval) will be used as reasonable initial estimation of known major biological populations for down-stream analyses. Further, Supplementary Figure 16f shows overall high agreement of estimated CMS between different normalized datasets and strategies.

**Step 2: Unwanted variation**

Supplementary Figure 16a shows substantial library size variation and differences between the major time intervals. However, there are situations where batch details are not available or are unknown. In these situations batches can be estimated from the data using different robust approaches proposed by RUVprps. We now assume that the time interval details of the TCGA READ RNA-seq data are unknown and proceed to estimate them.

The `identifyUnknownUV()` function will be applied using parameters specified for chronologically ordered samples. We will specify known biological and unwanted variables in order to perform an association analysis between the estimated batches and those variables. This analysis is recommended if such information is available. The estimated batches will be added to the sample annotation file of the *SummarizedExperiment* object for down-stream analysis.

```r
# identifying unknown sources of unwanted variation
read.se.obj <- identifyUnknownUV(
    se.obj = read.se.obj,
    assay.name = 'RawCount',
    approach = 'rle',
    chronological.detection = TRUE,
    assess.bio.association = TRUE,
    bio.variables = c('Quantile.CMS.TI', 'Tumour.purity'),
    assess.uv.association = TRUE,
    uv.variables = c('Years', 'Plates'),
    generate.association.plot = TRUE,
    add.to.sample.annotation = TRUE,
    col.name = 'Estimated.batches',
    output.name = 'iteration1')
```

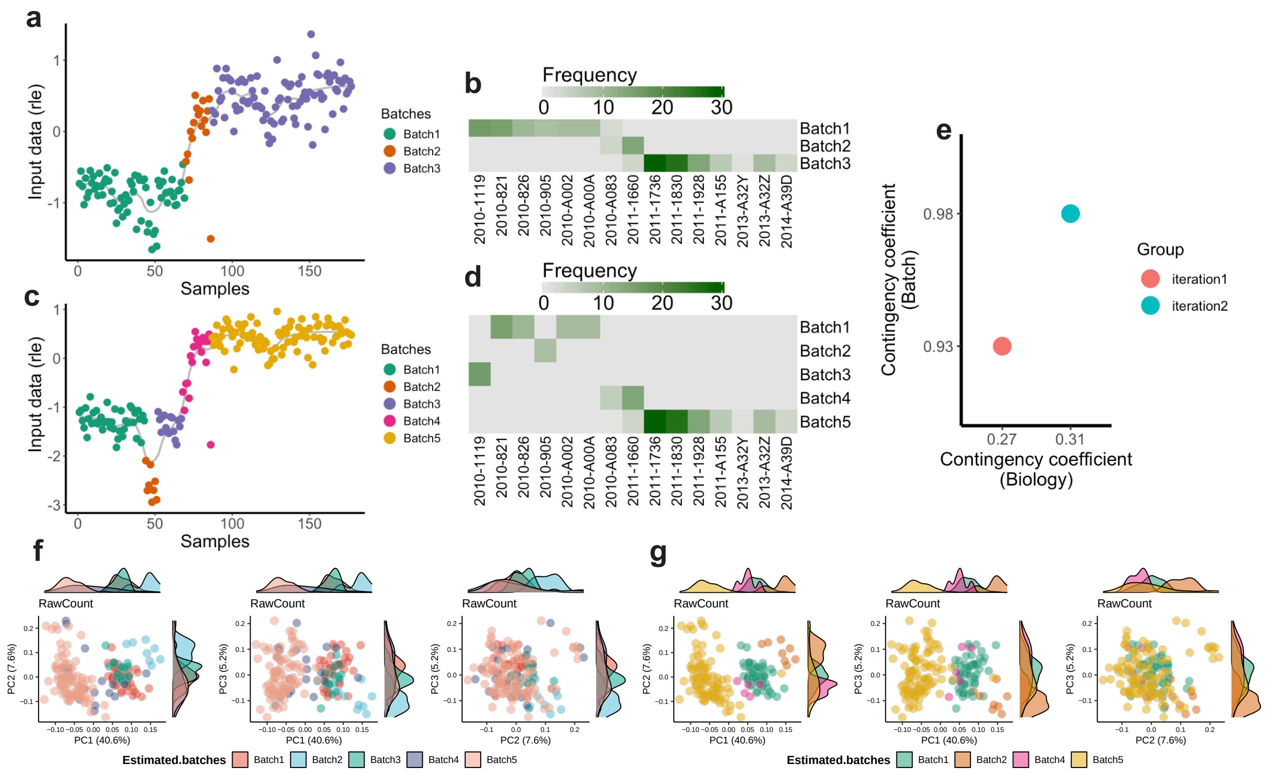


Supplementary Figure 17: Identification of batches in TCGA READ RNA-seq data using an unsupervised approach. a) The scatter plot displays the relative log expression (RLE) medians calculated from the raw count data using all genes, categorized into potential batches using the changepoint approach in the `identifyUnknownUV()` function. The heatmap presents sample counts, illustrating the association between the estimated batches (inferred from RLE medians) and known unwanted sources of variation including the plated and years. b) Similar to panel a, but the RLE medians are computed using the top 500 genes most affected by the estimated batch effects identified in the first iteration (a). c) The scatter plot shows Cramér's V contingency coefficients, quantifying the association between sample distributions in the estimated batches and those in known technical batches (Y-axis), as well as known biological subgroups (X-axis). d) The scatter plots of first three PCs on TCGA READ raw count data colored by the initial estimates of batch effects. e) The scatter plots of first three PCs on TCGA READ raw count data colored by the aggregation of batches obtained from the the iteration in b.

Supplementary Figure 17a shows that the RLE medians of the TCGA raw count data ($log_2 + 1$) can clearly separate the major time intervals in the data. Furthermore, Supplementary Figure 17b indicates that the analysis also captures certain plate effects present in the data. The smooth line across the RLE medians suggests that there might be more extra batch effects in the data. Then, we recommend iterating the step above using a set of genes that are highly affected by the initially estimated batches. These genes will be identified using the `computeGenesVariableAnova()` function, which applies ANOVA between individual genes and the estimated batches as a factor. The top 500 genes (arbitrary cut off) with the highest F-statistic will be used to re-run the `identifyUnknownUV()` function.

```
# Finding highly affected genes by the estimated batches
read.se.obj <- computeGenesVariableAnova(
    se.obj = read.se.obj,
    assay.names = 'RawCount',
```

```
    variable = 'Estimated.batches')

# Selecting the top 500 genes with the highest F-statistic
hag <- obtainMetric(
    se.obj = read.se.obj,
    slot = 'Metrics',
    metric.group = 'gene.level',
    assay.name = 'RawCount',
    metric.name = 'ANOVA',
    test.name = 'aov',
    variable = 'Estimated.batches')
hag <- hag[order(hag$statistic, decreasing = TRUE) , ]

# Estimating batches using only the top 500 genes
read.se.obj <- identifyUnknownUV(
    se.obj = read.se.obj,
    assay.name = 'RawCount',
    approach = 'rle',
    ncg = row.names(hag)[1:500],
    chronological.detection = TRUE,
    assess.bio.association = TRUE,
    bio.variables = c('Quantile.CMS.TI', 'Tumour.purity'),
    assess.uv.association = TRUE,
    uv.variables = c('Years', 'Plates'),
    generate.association.plot = TRUE,
    add.to.sample.annotation = TRUE,
    col.name = 'Estimated.batches',
    output.name = 'iteration2')
read.se.obj[['Estimated.batches']][read.se.obj[['Estimated.batches']] == 'Batch3'] <- 'Batch1'
read.se.obj[['Estimated.batches']][read.se.obj[['Estimated.batches']] == 'Batch4' & read.se.obj[['Time.
       interval']] == 'Batch2'] <- 'Batch5'
```

Supplementary Figure 17c and d show that the RLE medians computed using the selected genes more effectively capture the plate effects present in the data. Supplementary Figure 17c presents Cramér's V contingency coefficients, comparing the estimated batch assignments with known technical and biological groupings. Specifically, the Y-axis reflects agreement with known batch groupings (defined as all combinations of time points and plates), while the X-axis shows agreement with known biological populations (defined by combinations of tumor purity, categorized into three intervals, and CMS subtypes). This analysis shows that the iteration step captures more unwanted variation in the data. However, Supplementary Figure 17c and f suggest that certain batches with similar RLE medians can be aggregated for downstream analysis. For example, Batch 1 and Batch 3 can be combined into a single batch. Similarly, the samples from Batch 4, which originate from the Batch 2 time interval, can be aggregated with Batch 5. Finally, Supplementary Figure 17g shows that the final estimated batches are well separated across first PC of the raw count ($log_2 + 1$) data.

**Stage 3: Variation assessment**

Before proceeding with RUV-III normalization or any other normalization, it is crucial to assess the impact of both biological and unwanted variation on all data sets. At this stage a range of statistical metrics that are categorized into global and gene level will be applied on all the assays.

#### Step 1: Checking the SummarizedExperiment object

After selecting both biological and unwanted variables, applying the `checkSeObj()` function is recommended to remove any `NA` or missing values in both the assays and the variables. Recall that the current RUV-III method does not support `NA` or missing values in the data.

```
# using the 'Quantile.CMS.TI' for down-stream analysis
read.se.obj[[CMS]] <- read.se.obj$Quantile.CMS.TI
read.se.obj <- checkSeObj(
  se.obj = read.se.obj,
  assay.names = 'all',
  variables = c('CMS','Library.size','Tumour.purity','Time.interval', 'Estimated.batches'))
```

The result shows that there are no `NA` or missing values in either the data or the variables. Note that the `check.se.obj` argument of all the function used in subsequent analysis can be set to `FALSE`.

#### Step 2: Creating all possible assessments (optional)

The `getAssessmentMetrics()` function can be used to find all possible plots and assessments for all specified biological and unwanted variables. Plots or assessments that are not of interest can be excluded for the later steps. We recommend generating all the plots to visually explore the variation and then deciding which ones to exclude for down-stream analyses.

```
# Visualizing all possible assessment metrics for the variables
read.se.obj <- getAssessmentMetrics(
    se.obj = read.se.obj,
    bio.variables = c('CMS', 'Tumour.purity'),
    uv.variables = c('Library.size','Time.interval', 'Estimated.batches'),
    plot.output = TRUE)
```

Supplementary Figure 18 shows that 67 plots and metrics will be generated to assess the impact of both biological and unwanted variation in the data. Note that the code for each assessment can be used to exclude specific assessments from downstream analysis.

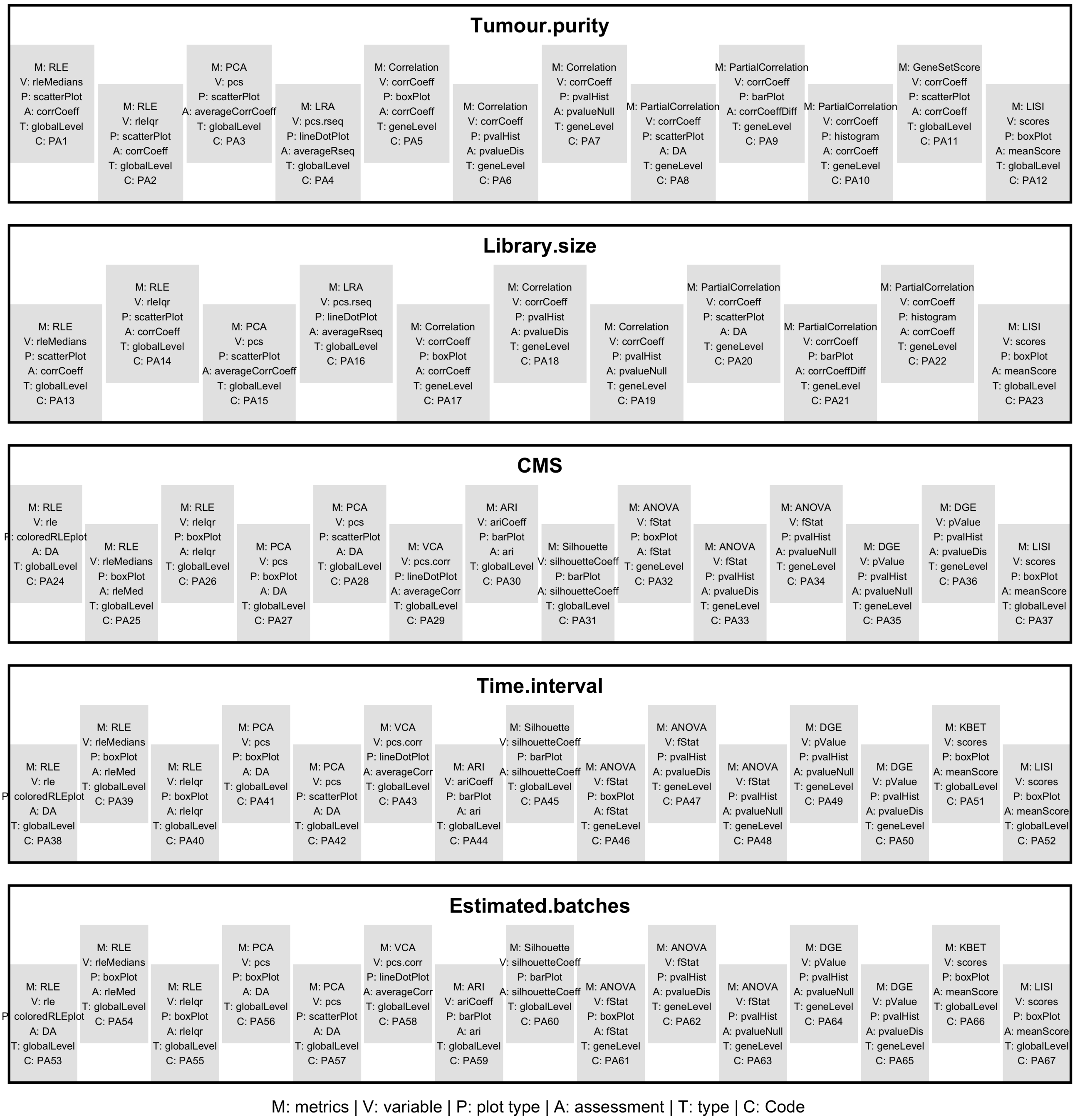


Supplementary Figure 18: Assessment matrices for all the biological and unwanted variation for the TCGA READ RNA-seq data. For each individual variable, the specified metrics will be computed and the corresponding plots will be generated. The code for each assessment can be used to exclude the corresponding metrics from the assessment of variation and normalization steps. The 'DA' (Does not Apply) indicates that specific metrics will not be considered for assessing normalization.

#### Step 3: Selecting genes for gene-gene correlation assessment and tumor purity estimation

Forming partial gene-gene correlations is one of the metrics used in the variation assessment process. If this metric is selected, it is preferable to provide a list of genes whose expression is highly associated with the corresponding variable; otherwise, all genes will be used, which can be computationally expensive. The `selectGenesForPPcorr()` function will be used to identify gene sets for library size and purity variation. Spearman correlation analysis will be performed within each major time interval to account for the large batch effects in the data.
Preserving the variation in tumor purity is one of the aims of this procedure. Gene set scoring analysis will be performed to estimate tumor purity in data sets, serving as one of the metrics in the variation assessment process. To estimate tumor purity for individual samples, predefined immune and stromal gene sets will be used. The `selectGeneSets()` will be used to retrieve an immune and stromal gene signature in the *SummarizedExperiment* object. Note that users may also provide appropriate alternative gene signatures for estimating tumor purity.

```
# selecting genes for gene-gene correlation
ppcorr.gene.sets <- selectGenesForPPcorr(
    se.obj = read.se.obj,
    assay.names = c('RawCount', 'FPKM.UQ'),
    variables = c('Library.size', 'Tumour.purity'),
    cor.cutoff = 0.6,
    groups = 'Time.interval')

#retrieving immune and stromal gene set
tumour.purity.genes.sets <- selectGenesSets(
    se.obj = read.se.obj,
    gene.set = "immune.stromal")
```

The 1,535 and 870 genes that have an absolute correlation of more than 0.6 with purity and library size are identified, respectively. A total of 273 genes are selected to estimate tumor purity in the data.

#### Step 4: Assessing variation

The `assessVariation()` function will apply all the statistical metrics for all specified variables. The results including statistical summaries and plots of individual metrics will be stored in the *SummarizedExperiment* object. The `plotAssessVariation()` function will create a PDF file that contains all the plots. This file will be stored in the current home directory.

```
read.se.obj <- assessVariation(
    se.obj = read.se.obj,
    assay.names = 'all',
    assessment.level = 'L2',
    bio.variables =  c('CMS', 'Tumour.purity'),
    uv.variables =  c('Library.size','Time.interval', 'Estimated.batches'),
    pcorr.genes = ppcorr.gene.sets,
    gene.set.score.list = tumour.purity.genes.sets,
    general.points.size = 3,
    override.check = FALSE,
    check.se.obj = FALSE)
```

```
RUVprps:::plotAssessVariation(
    se.obj = read.se.obj,
    assay.names = 'all',
    variables =  c('CMS','Library.size','Tumour.purity','Time.interval'),
    output.file.name = "RUVprps_TCGA_READ_InitialVariationAssessments")
```

Supplementary File "TCGA_READ_InitialVariationAssessments.pdf" displays all the plots of individual metrics applied by the `assessVariation()` function. Briefly, the first two PCs of the TCGA normalized data sets clearly show that the effect of the major time intervals persists in these datasets. This conclusion is further supported by the other gene-level and global-level assessments. We will now apply RUV-III with PRPS to remove the impact of library size and the major time intervals from the raw count data while preserving variation in two major known biological variables including tumor purity and CMS. Ideal normalization should preserve all biological variation of interest in the data. The relatively high association between and known biological variation is attributed to the presence of confounding in the data.

#### Stage 4: RUV-III normalization

RUV-III with PRPS will be applied in four different scenarios. The general process for each scenario involves identifying a suitable set of NCG, creating PRPS data for each source of unwanted variation, performing RUV-III normalization with multiple values for $k$, and finally assessing the normalization performance.

##### Scenario 1: Both biological and unwanted variation are known

In this scenario, only known sources of biological and unwanted variation will be considered in the RUV-III normalization. Library size and the major time intervals will then be used as the known sources of unwanted variation, while CMS and tumor purity will be recognized as the major biological variation. RUV-III normalization will be conducted in a fully supervised manner.

##### Step 1: Selecting negative control genes

A set of NCG will be identified through implementation of the `findNcgSupervised()` function. All the available approaches will be tested and then the `compareNCGs()` function will be applied to select the most suitable set of genes as NCG. All NCG sets will be stored in the *SummarizedExperiment* object. It should be noted that the results of the `comapreNCGs()` function are suggestive rather than definitive. Users may test all NCG sets and evaluate their performance accordingly.

```
ncg.options <- c(
    'AnovaCorr.AcrossAllSamples',
    'AnovaCorr.PerBatchPerBiology',
    'LinearMixedModel',
    'TwoWayAnova'
    )

for(i in 1:4){
    read.se.obj <- findNcgSupervised(
        se.obj = read.se.obj,
        assay.name = 'RawCount',
        bio.variables = c('Tumour.purity', 'CMS'),
        uv.variables = c('Library.size', 'Time.interval'),
        samples.to.use = 'all',
```

```
        approach = ncg.options[i],
        form = ~ (1|Time.interval) + Library.size + (1|CMS) + Tumour.purity,
        use.rank = FALSE,
        bio.percentile = NULL,
        uv.percentile = 0.8,
        pseudo.count = 1,
        adjust.data = TRUE,
        adjustment.variables = "uv",
        ncg.selection.method = 'quantile',
        nb.ncg = 0.05,
        ncg.group.name = 'read.scenario1',
        ncg.set.name = ncg.options[i],
        nb.cores = 15,
        check.se.obj = FALSE,
        assess.ncg = TRUE)}

countNCG(
    se.obj = read.se.obj,
    ncg.selection = 'supervised',
    ncg.group.name = 'read.scenario1',
    create.venn.diagram = FALSE
    )
```

Supplementary Figure 19a and b show the satisfactory performance of all four NCG selection approaches. The 740 NCG identified by the `LinearMixedModel` approach show slightly better performance compared to other approaches. Then this set of NCG will be used for RUV-III normalization.

**Step 2: Creating the PRPS data**

The PRPS data will be generated for individual sources of unwanted variation using the `createPrPsSupervised()` function. The parameter `apply.other.uv.variables` will be set to `FALSE` due to insufficient sample size. In addition, both normal tissues and samples annotated as "Not.classified" for CMS will be excluded. Normal tissues will be omitted because they were all profiled within the second batch of the time interval and therefore are not informative for PRPS construction to remove the batch effects. Samples annotated as "Not.classified" for CMS will be excluded because their subtypes were undefined, which could lead to unsatisfactory PRPS data. All PRPS data will be stored in the *SummarizedExperiment* object. Note that tumor purity will be divided into four categorical groups using k-means clustering (the default option) to find homogeneous biological groups for creating the PRPS data. Increasing the number of categories will result in smaller homogeneous biological groups and may lead to insufficient samples to create PRPS data across batches. The function will print messages if the number of groups is too small or too large. PRPS data will be created using the RawCount ($log_2 + 1$) data (`apply.log = TRUE`).

```
read.se.obj <- createPrPsSupervised(
    se.obj = read.se.obj,
    assay.name = 'RawCount',
    bio.variables = c('Tumour.purity', 'CMS'),
    samples.to.use = !read.se.obj$CMS %in%  c('Normal', 'Not.classified'),
    uv.variables = c('Library.size', 'Time.interval'),
    bio.clustering.method = "kmeans",
    nb.bio.clusters = 4,
    apply.log = TRUE,
    pseudo.count = 1,
```

```
    check.prps.connectedness = FALSE,
    apply.other.uv.variables = FALSE,
    prps.group.name = 'read.scenario1',
    verbose = TRUE)
```

In total, 8 PR sets with a total of 16 PS were created to capture library size effects, and 8 PR sets from 18 PS were created for the major time interval batch difference. Supplementary Figure 19C shows the distribution of PRPS sets (PRPS map) across each source of unwanted variation.

**Step 3: Performing RUV-III normalization with multiple $k$ values**

First, the `getMaximumK()` function will be applied to find the possible maximum value, $K$, for the $k$. This shows that the maximum value for the current NCG and PRPS data is $K = 16$. If any values bigger than 16 are provided, the `RUVIIIprps()` function will ignore them. Here, all ranges of values for $k$ will be then tested and the performance for each $k$ will be evaluated. All the RUV-III normalized data will be stored as assays in the *SummarizedExperiment* object. The RUV-III-PRPS normalization will be applied on the RawCount ($log_2 + 1$) data.

```
read.se.obj <- RUVIIIprps(
    se.obj = read.se.obj,
    assay.name = 'RawCount',
    control.sample.types = 'prps',
    prps.type = 'supervised',
    prps.group  = 'read.scenario1',
    prps.set.names = 'all',
    ncg.type = 'supervised',
    ncg.group.names = 'read.scenario1',
    ncg.set.names = 'LinearMixedModel',
    k = c(1, seq(3, 16, 2)),
    apply.log = TRUE,
    data.to.log = 'assay',
    pseudo.count = 1,
    return.wa = TRUE,
    check.se.obj = FALSE)
```

Note that the `data.to.log` parameter was set to `assay`, indicating that RUV-III-PRPS applied log transformation exclusively to the `RawCount` data, whereas the PRPS data were not transformed because they had already been log-transformed during their construction.

**Step 4: Assessing the W matrix (optional)**

Prior to evaluating normalization performance, the `assessW()` function can be applied to assess the associations between the biological and unwanted variables and the unwanted factors estimated by RUV-III, represented by the columns of the estimated matrix $W$. Ideally these factors should have strong and no or weak associations with the unwanted and biological variables, respectively.

```
read.se.obj <- assessW(
    se.obj = read.se.obj,
    variables = NULL,
    compare.w = TRUE,
    bio.variables = c('Tumour.purity', 'CMS'),
```

```
    uv.variables = c('Library.size', 'Time.interval'))
```

Supplementary Figures 19d show that the estimated unwanted factors have a strong association with library size and the major time intervals, but a relatively low association with CMS and tumor purity. This plot will be useful for selecting the most appropriate $k$ value for RUV-III normalization. Overestimating the value of $k$ might result in removing variation between the CMS. Note that in RUV-III with PRPS normalization, we prefer to use lower values for $k$ as long as they reasonably capture the unwanted variation.

#### Step 5: Evaluating normalizations

To assess the performance of RUV-III and compare it with the TCGA normalizations, we will first estimate CMS for the RUV-III normalized datasets. Then, the association between the CMS and overall survival will be assessed to select the most suitable CMS for down-stream analysis. All the CMS estimates will be added to the sample annotation in the *SummarizedExperiment* object. Note that if the CMS estimates derived from the TCGA-normalized data are considered reliable, this step becomes unnecessary.

```
read.se.obj <- estimateCMS(
    se.obj = read.se.obj,
    assay.names = names(SummarizedExperiment::assays(read.se.obj))[-c(1:4)],
    raw.count.assay.name = NULL,
    tissue.type = 'Tissues',
    groups = NULL,
    remove.na = 'none',
    apply.log = FALSE,
    seed = 122334,
    nb.cores = 14)
```

The association of the estimated CMS with patient survival will be assessed. This analysis will determine which estimated CMS should be used for evaluating variation and normalization. Furthermore, this analysis can inform the selection of an appropriate value of $K$. Note that, in the next analysis, both CMS and survival analysis for the TCGA datasets will not be re-calculated.

```
all.survivals <- lapply(
    cms <- grep('RUVIIIprps', colnames(colData(read.se.obj)), value = TRUE),
    function(i) {
        idx <- read.se.obj$OS != -1 & read.se.obj[[i]] != 'Not.classified'
        computeSurvival(
            se.obj = read.se.obj[, idx],
            variable = i,
            survival.time = 'OS.time',
            survival.events = 'OS',
            genes = NULL,
            return.survival.plot = TRUE,
            check.se.obj = FALSE,
            save.se.obj = FALSE)$plot})
```

Supplementary Figure 20 shows that the CMS estimated by RUV-III with $k = 15$ yields the lowest p-value compared to the `Quantile.CMS.TI` method (Supplementary Figure 16d). These estimates will be used for variation assessment and evaluation of normalization performance.

Next, the `assessVariation()` function, followed by `assessNormalization()`, will be applied to all RUV-III normalized data in the *SummarizedExperiment* object. Note that there is no need to re-calculate the metrics for the TCGA datasets, as they have already been computed and stored in the *SummarizedExperiment* object. By setting `override.check = TRUE`, the function will not recompute metrics already available for each dataset. However, if one wishes to re-calculate the metrics for the TCGA datasets, the data must first be log-transformed, after which the `assessVariation()` function can be applied with `apply.log = FALSE`, since all RUV-III normalized data are already in log scale.

```
read.se.obj$CMS <- read.se.obj$RUVIIIprps_k_15.CMS
read.se.obj <- assessVariation(
    se.obj = read.se.obj,
    assay.names = 'all',
    assessment.level = 'L2',
    bio.variables = c("Tumour.purity", "CMS"),
    uv.variables = c("Library.size", "Time.interval"),
    pcorr.genes = ppcorr.gene.sets,
    gene.set.score.list = tumour.purity.genes.sets,
    pcorr.filter.genes = FALSE,
    general.points.size = 3,
    override.check = TRUE,
    apply.log = FALSE,
    check.se.obj = FALSE)

read.se.obj <- assessNormalization(
    se.obj = read.se.obj,
    assay.names = 'all',
    assessment.level = 'L1',
    select.top.ruv = FALSE,
    bio.variables = c("Tumour.purity", "CMS"),
    uv.variables = c("Library.size", "Time.interval"),
    bio.weight = 0.6,
    uv.weight = 0.4,
    corr.cutoff = list(Tumour.purity = .5, Library.size = .3),
    output.name = 'tcga.read.assessNormalization.scenario1')

RUVprps:::plotAssessVariation(
    se.obj = read.se.obj,
    assay.names = 'all',
    variables =  c('CMS','Library.size','Tumour.purity','Time.interval'),
    output.file.name = "RUVprps_TCGA_READ_VariationAssessment_Secnario1")
```

Supplementary Figure 19e demonstrates, level 2 normalization assessment, that RUV-III-PRPS more effectively removes unwanted variation while preserving known biological variation compared to the TCGA normalization methods. The Supplementary File "TCGA_READ_variationAssessment_Scenario1.pdf" contains all the plots summarized in the Supplementary Figure 19e. Across all evaluation metrics, RUV-III-PRPS with $k = 11$ exhibits the strongest overall performance among the tested normalization approaches, however, the differences between the RUV-III-PRPS normalized data are not significant. It should be noted that the choice of k may be tailored to the specific objectives of downstream analyses. For instance, while $k = 11$ yields the most robust results across the full set of metrics, a higher value such as $k = 15$ provides better association between the CMS and survival. Thus, the flexibility of RUV-III-PRPS allows users to select an adjustment level that balances noise reduction and signal preservation in a manner most appropriate for their study.

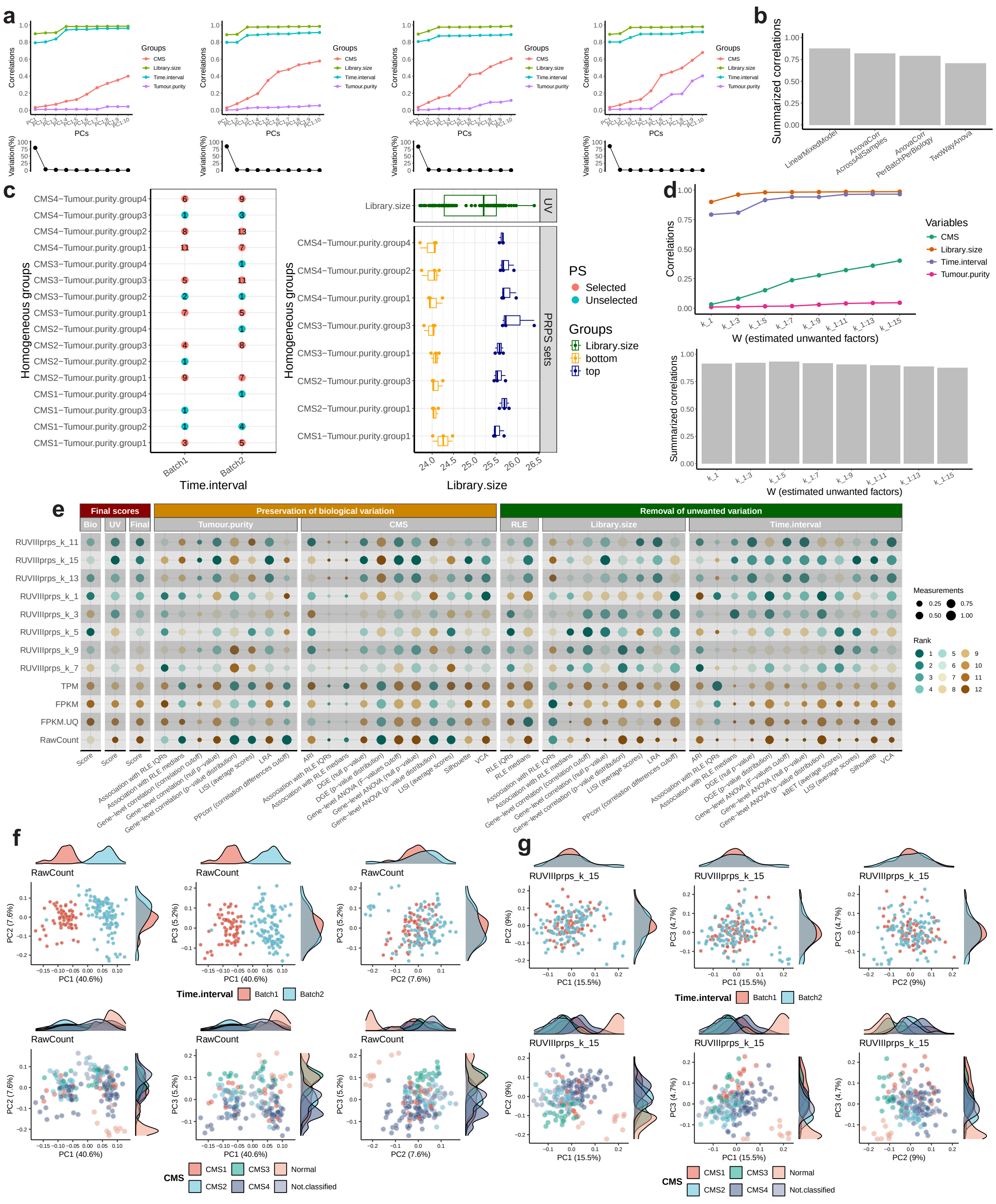

Supplementary Figure 19: RUV-III normalization performance on the TCGA READ RNA-seq data for scenario 1. a) Line-dot plots illustrate the performance of the NCG selection methods, including `LinearMixedModel` (first plot), `AnovaCorr.AcrossAllSamples` (second plot), `AnovaCorr.PerBatchPerBiology` (third plot), and `TwoWayAnova` (fourth plot), applied to the raw count data $log2_{+}1$ from the TCGA READ RNA-seq dataset. b) The bar plot summarizes the performance of the selected NCG sets, where a higher correlation indicates better performance. c) PRPS maps are presented for the time interval (first plot) and library size (second plot) variables. The Y-axis shows all possible homogeneous groups with respect to known biological variation. The X-axes represent the distributions of samples across time intervals (batch 1 and batch 2) and library size ranges, respectively. In the time interval PRPS map, red points indicate groups containing more than two samples, and each horizontal line connecting two red points defines a PRPS set across the time interval. In the library size PRPS map, each red line corresponds to six samples representing the highest and lowest library sizes within a partially homogeneous biological sample group. Every three samples are averaged to create a PS. The X-axis of the library size PRPS map displays the $\log_2(n)$ of library size and highlights the samples selected for PS construction. d) The line-dot plot shows the associations between the columns of the estimated W matrix and both biological and unwanted variables, with the accompanying bar plot summarizing these correlations. A higher correlation the better performance. e) Numerical assessments of all metrics for both biological and unwanted variables are presented. Each point represents the performance of a given normalization method, with higher values indicating better performance. Normalized datasets are ranked according to their effectiveness in removing unwanted variation while retaining biological signals. f) The first three PC plots of the TCGA READ raw count data $log_2 + 1$ with $K = 15$, colored by the major time intervals (top row) and the CMS (bottom row). g) similar to f, for the RUV-III-PRPS normalization with $K = 15$

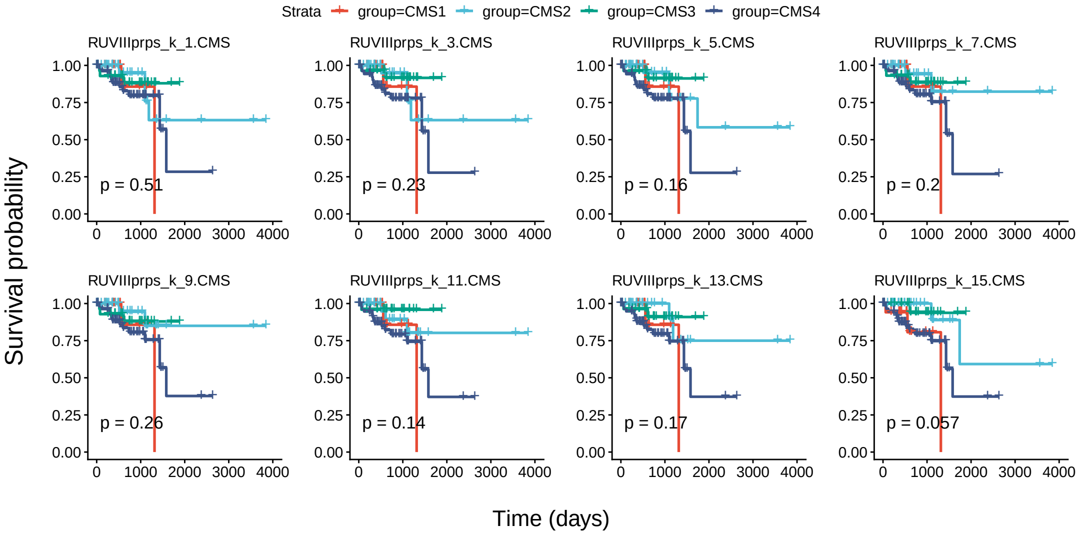


Supplementary Figure 20: Kaplan–Meier analysis of CMS estimated from each RUV-III–normalized dataset in scenario 1 and patient survival in the TCGA READ RNA-seq data.

#### Scenario 2: Unknown unwanted and known biological variation

In this second scenario it is assumed that the batches in the data are unknown. Then, the estimated batches (see Stage 2 above) along with library size will be used as sources of unwanted variation. In this scenario, similar to the scenario 1, the CMS and tumor purity will be considered as the known biological variation.

#### Step 1: Selecting negative control genes

A set of NCG will be obtained using the `findNcgsSupervised()` function. Similar to scenario 1, all the four options will be tested and then their performance will be assessed.

```
ncg.options <- c(
    'AnovaCorr.AcrossAllSamples',
    'AnovaCorr.PerBatchPerBiology',
    'LinearMixedModel',
    'TwoWayAnova'
    )
for(i in ncg.options){
    read.se.obj <- findNcgSupervised(
        se.obj = read.se.obj,
        assay.name = 'RawCount',
        samples.to.use = 'all',
        bio.variables = c('Tumour.purity', 'CMS'),
        uv.variables = c('Estimated.batches', 'Library.size'),
        form = ~ (1|Estimated.batches) + Library.size + (1|CMS) + Tumour.purity,
        approach = i,
        bio.percentile = NULL,
        uv.percentile = .8,
        use.rank = FALSE,
        ncg.selection.method = 'quantile',
        adjust.data = TRUE,
        adjustment.variables = 'uv',
        nb.ncg = 0.05,
        ncg.group.name = 'read.scenario2',
        nb.cores = 15,
        ncg.set.name = i,
        check.se.obj = FALSE,
        assess.ncg = TRUE)}

countNCG(
    se.obj = read.se.obj,
    ncg.selection = 'supervised',
    ncg.group.name = 'read.scenario2',
    create.venn.diagram = FALSE
    )ead.scenario2',
    nb.pcs = 10)
```

Supplementary Figure 21 a and b show that the 760 NCG selected by the `LinearMixedModel` option exhibits the highest and lowest correlations with the unwanted and biological variation, respectively. This set of genes will be used for RUV-III normalization.

#### Step 2: Creating the PRPS data

PRPS data will be created for library size and the estimated batches using the the function `createPrPsSupervised()` function. Similar to scenario 1, the `apply.other.uv.variables` parameter will be set to `FALSE` due to insufficient sample size.

```
read.se.obj <- createPrPsSupervised(
```

```
    se.obj = read.se.obj,
    assay.name = 'RawCount',
    bio.variables = c('Tumour.purity', 'CMS'),
    samples.to.use = !read.se.obj$CMS %in%  c('Normal', 'Not.classified'),
    uv.variables = c('Estimated.batches', 'Library.size'),
    nb.bio.clusters = 4,
    apply.log = TRUE,
    bio.clustering.method = 'quantile',
    apply.other.uv.variables = FALSE,
    check.prps.connectedness = FALSE,
    prps.group = 'read.scenario2',
    verbose = TRUE)
```

In total, 8 PR sets with a total of 16 PS are created for library size effects, and 6 PR sets from 13 PS are created for the estimated batch effects. Supplementary Figure 21c shows the distribution of PRPS sets across each source of unwanted variation.

##### Step 3: Performing RUV-III normalization with multiple $k$ values

The *getMaximumK()* function shows that the maximum $k$ for the scenario 2 is $K = 14$. RUV-III with with all possible values of $k$ will then be applied using the selected set of NCG and all the PRPS data generated above.

```
read.se.obj <- RUVIIIprps(
    se.obj = read.se.obj,
    assay.name = 'RawCount',
    prps.type = 'supervised',
    prps.group  = 'read.scenario2',
    ncg.type = 'supervised',
    ncg.group = 'read.scenario2',
    ncg.set.names = 'LinearMixedModel',
    k = c(1:15),
    apply.log = TRUE,
    data.to.log = 'assay',
    pseudo.count = 0.5,
    return.wa = TRUE,
    check.se.obj = FALSE)
```

##### Step 4: Assessing the W matrix

The function `assessW()` will be applied to examine the association between the biological and unwanted variables with the columns of unwanted factors matrix W estimated by RUV-III.

```
read.se.obj <- assessW(
    se.obj = read.se.obj,
    compare.w = TRUE,
    bio.variables = c('Tumour.purity', 'CMS'),
    uv.variables = c('Estimated.batches', 'Library.size'))
```

Figure 21d shows that the estimated unwanted factors have significant association with the unwanted variables and reasonably weak association with the biological variables.

#### Step 5: Evaluating normalizations

As with scenario 1, CMS will be estimated using the RUV-III normalized datasets and then the `assessVariation()` function will be applied on only RUV-III normalized data sets. Finally, the `assessNormalization()` function will be performed on all available assays in the *SummarizedExperiment* object.

```
read.se.obj <- estimateCMS(
    se.obj = read.se.obj,
    assay.names = names(SummarizedExperiment::assays(read.se.obj))[-c(1:4)],
    raw.count.assay.name = NULL,
    tissue.type = 'Tissues',
    remove.na = 'none',
    apply.log = FALSE,
    seed = 122335,
    nb.cores = 10)

all.survivals <- lapply(
    cms <- grep('RUVIIIprps', colnames(colData(read.se.obj)), value = TRUE),
    function(i) {
        idx <- read.se.obj$OS != -1 & read.se.obj[[i]] != 'Not.classified'
        computeSurvival(
            se.obj = read.se.obj[, idx],
            variable = i,
            survival.time = 'OS.time',
            survival.events = 'OS',
            genes = NULL,
            return.survival.plot = TRUE,
            check.se.obj = FALSE,
            save.se.obj = FALSE)$plot})
```

Supplementary Figure 22 shows the survival analyses of the CMS estimated using the RUV-III normalized datasets. The RUV-III data with $k = 10$ shows greater significance compared to the 'Quantile.CMS.TI' method (Supplementary Figure 16d). The newly estimated CMS will then be used for assessing both the variation and normalization steps. Note that the `override.check` parameter of the `assessVariation()` function will be set to `TRUE`. This setting will prevent the need to recompute all the metrics from the raw count and TCGA normalized data in Stage 3.

```
read.se.obj$CMS <- read.se.obj$RUVIIIprps_k_10.CMS
read.se.obj <- assessVariation(
    se.obj = read.se.obj,
    assay.names = 'all',
    assessment.level = 'L2',
    bio.variables = c("Tumour.purity", "CMS"),
    uv.variables = c("Library.size", "Estimated.batches"),
    pcorr.genes = ppcorr.gene.sets,
    gene.set.score.list = tumour.purity.genes.sets,
    pcorr.filter.genes = FALSE,
    general.points.size = 3,
    override.check = TRUE,
    apply.log = FALSE)

read.se.obj <- assessNormalization(
    se.obj = read.se.obj,
```

```
    assay.names = 'all',
    bio.variables = c('CMS', 'Tumour.purity'),
    uv.variables = c('Library.size', 'Estimated.batches'),
    bio.weight = 0.6,
    uv.weight = 0.4,
    assessment.level = 'L2',
    select.top.ruv = F,
    corr.cutoff = list(Tumour.purity = .5, Library.size = .3),
    output.name = 'tcga.read.assessNormalization.scenario2')

RUVprps:::plotAssessVariation(
    se.obj = read.se.obj,
    assay.names = 'all',
    variables =  c('CMS','Library.size','Tumour.purity','Estimated.batches'),
    output.file.name = "RUVprps_TCGA_READ_VariationAssessment_Secnario2")
```

Supplementary Figure 21e presents the level 2 normalization assessment, illustrating that RUV-III-PRPS more effectively removes unwanted variation while preserving known biological signals compared to the TCGA normalization methods. All plots summarized in Figure 21e are provided in the Supplementary File "TCGA_READ_variationAssessment_Scenario2.pdf". Across all evaluation metrics, RUV-III-PRPS with $k = 10$ demonstrates the strongest overall performance among the normalization approaches tested.

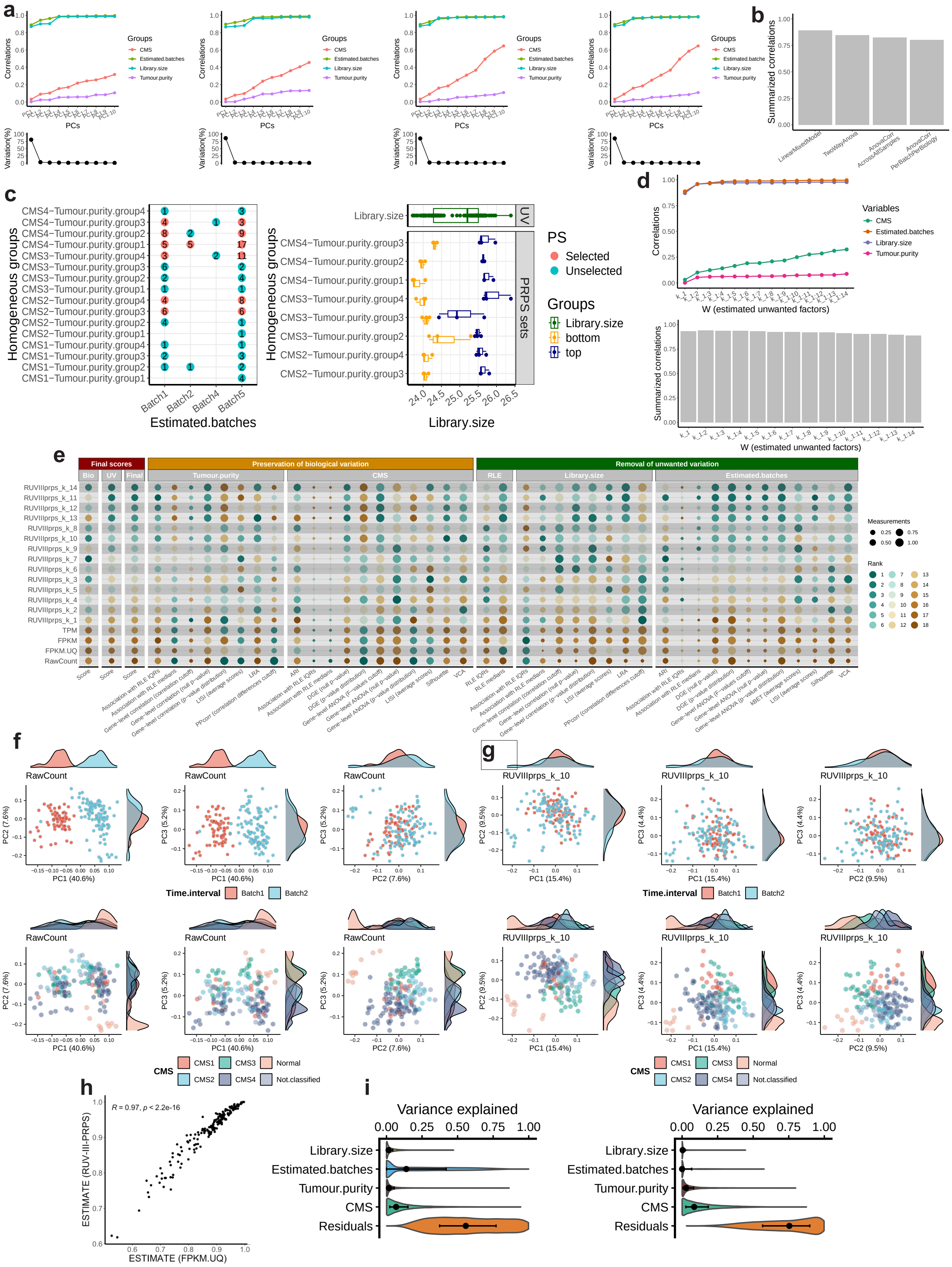

Supplementary Figure 21: RUV-III normalization performance on TCGA READ RNA-seq data for scenario 2. a) Line–dot plots compare the performance of negative control gene (NCG) selection methods applied to the TCGA READ raw count data ($\log_2(x + 1)$), including `LinearMixedModel` (first panel), `TwoWayAnova` (second panel), `AnovaCorr.AcrossAllSamples` (third panel) and `AnovaCorr.PerBatchPerBiology` (fourth panel). b) Bar plots summarize the performance of the resulting NCG sets, with higher correlations indicating superior performance. c) PRPS maps are shown for the estimated batches (first panel) and library size (second panel). The y-axis enumerates all homogeneous groups with respect to known biological variation, whereas the x-axes represent sample distributions across estimated batches and library size ranges, respectively. In the estimated batches PRPS map, red points denote groups containing more than two samples, and each horizontal line connecting two red points defines a PRPS set across time intervals. In the library-size PRPS map, each red line corresponds to six samples representing the highest and lowest library sizes within a partially homogeneous biological group. Every three samples are averaged to form a pseudo-sample (PS). The x-axis displays $\log_2(n)$ of the library size and highlights the samples selected for PS construction. d) Line–dot plots depict the associations between columns of the estimated $W$ matrix and both biological and unwanted covariates, with the accompanying bar plots summarizing these correlations; higher values indicate improved performance. e) Quantitative summaries of all evaluation metrics for both biological and unwanted variables are shown. Each point represents a normalization method, with higher values reflecting more effective removal of unwanted variation while preserving biological signal. Normalized datasets are ranked accordingly. f) Principal component plots of the first three components of the TCGA READ raw count data ($\log_2(x + 1)$) and RUV-III-PRPS normalized data with $K = 15$, colored by estimated batches (top row) and CMS (bottom row). g) As in f, but for data normalized using RUV-III-PRPS with $K = 10$. h) The tumor purity estimates for the TCGA FPKM.UQ data and RUV-III-PRPS normalized data. i) Shows the results of a linear mixed model analysis followed by gene-level variation analysis across all the variables for the TCGA FPKM.UQ (first plot) and the RUV-III-PRPS normalized data with $k = 10$ (second plot).

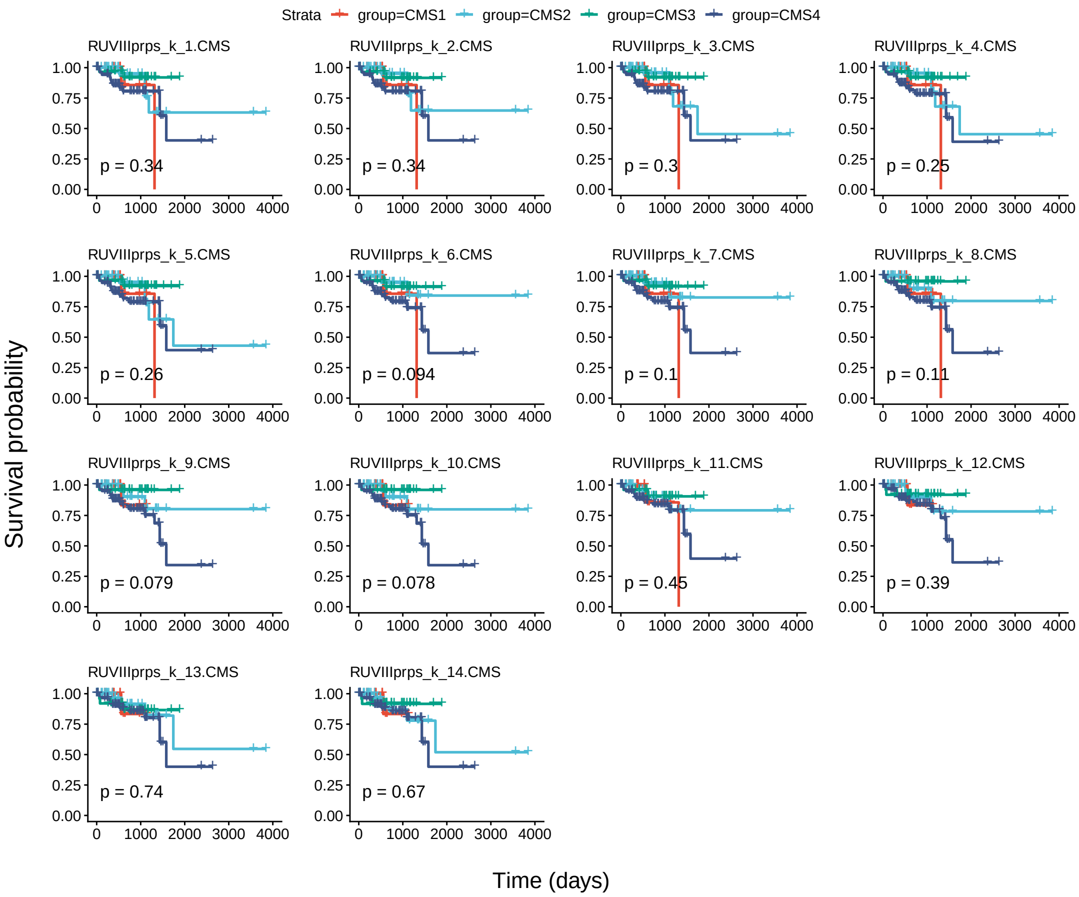


Supplementary Figure 22: Kaplan–Meier analysis of CMS estimated from each RUV-III–normalized dataset in scenario 2 and patient survival in the TCGA READ RNA-seq data.

#### Scenario 3: Known unwanted and unknown biological variation

In third scenario, it is assumed that the biological variation in the data is wholly unknown. The NCG and PRPS will then be identified in the unsupervised manner. In this scenario, library size and the major time intervals will be considered as known sources of unwanted variation.

#### Step 1: Selecting negative control genes

A set of NCG will be obtained using the `findNcgUnSupervised()` function. Both available approaches will be applied, and the performance of selected NCG sets will be assessed. It should be noted that known biological variables including CMS and tumor purity will only be used to assess the performance of the NCG sets.

```
for(i in c( 'AnovaCorr', 'LinearMixedModel')){
    read.se.obj <- findNcgUnSupervised(
        se.obj = read.se.obj,
        assay.name = 'RawCount',
        approach = i,
```

```
        use.rank = FALSE,
        ncg.selection.method = 'quantile',
        uv.variables = c('Time.interval', 'Library.size'),
        form =  ~ (1 | Time.interval) + Library.size,
        bio.percentile = NULL,
        uv.percentile = .8,
        nb.ncg = 0.05,
        variables.to.assess.ncg = c('Tumour.purity', 'CMS', 'Time.interval', 'Library.size'),
        ncg.group.name = 'read.scenario3',
        ncg.set.name = i,
        nb.cores = 15,
        check.se.obj = FALSE)}

countNCG(
    se.obj = read.se.obj,
    ncg.selection = 'un.supervised',
    ncg.group.name = 'read.scenario3')
```

Supplementary Figure 24A and B show that the performance of NCGs selected by the `LinearMixedModel` approach is superior to that of `AnovaCorr`. The first plot indicates a very strong association with unwanted variation and a moderate association with known biological variation.

It is important to emphasize that this scenario assumes biological variation is fully unknown, although we used CMS and tumor purity variation to assess the NCG selection approaches. If biological variation were indeed fully unknown, we would recommend following approach to estimate major biological variation in the data. First, a set of NCG that captures all sources of unwanted variation should be identified in the unsupervised manner. Subsequently, PCA could be performed on the NCGs, followed by regressing out a small number of PCs from the data to obtain residuals. Performing PCA on these residuals would then yield the first few PCs, which could serve as reasonable proxies for the major biological variation, and thus be useful for assessing NCG performance and other downstream steps.

#### Step 2: Creating the PRPS data

PRPS data will be generated for library size and major time intervals using the `createPrPsUnSupervised()` function. The top 20% of highly variable genes will be selected using the `findHVG()` function to improve the performance of CCA and MNN in creating PRPS.

```
hvg <- findHVG(
    se.obj = read.se.obj,
    assay.name = 'RawCount',
    approach = 'lmm',
    uv.variables = c('Time.interval', 'Library.size'),
    form = ~ (1 | Time.interval) + Library.size,
    nb.hvg = .2,
    nb.cores = 14,
    save.se.obj = FALSE)

uv.variables <- c("Library.size", "Time.interval")
max.prps.sets <- c(5, 5)
for(i in c(1:2)){
    read.se.obj <- createPrPsUnSupervised(
```

```
        se.obj = read.se.obj,
        assay.name = 'RawCount',
        uv.variables = uv.variables[i],
        other.uv.variables = NULL,
        select.extreme.groups = TRUE,
        approach = 'cca',
        nb.cca = 4,
        nb.pcs = 5,
        nb.clusters = 3,
        clustering.method = 'quantile',
        coordinates.to.use = 'cca',
        max.prps.sets = max.prps.sets[i],
        filter.prps.sets = TRUE,
        hvg = hvg,
        apply.log.for.prps = TRUE,
        create.prps.map = TRUE,
        prps.group.name = 'read.scenario3')}
```

In total, 5 PR sets comprising 19 PS samples were generated to remove time interval effects, and 5 PR sets comprising 10 PS samples were constructed to account for library size variation. Figure 24c shows the distribution of the PRPS data across these sources of unwanted variation. Note that the parameter `select.extreme.groups = TRUE` was used, indicating that the extreme groups of samples with the highest and lowest library sizes were selected for PRPS construction.

**Step 3: Performing RUV-III normalization with multiple $k$ values**

The `getMaximumK()` function shows that the maximum $k$ for the scenario 3 is 10. Then RUV-III with all possible values of $k$ will be applied using the selected set of NCG and all the PRPS data.

```
read.se.obj <- RUVIIIprps(
    se.obj = read.se.obj,
    assay.name = 'RawCount',
    prps.type = 'un.supervised',
    prps.group.names = 'read.scenario3',
    prps.set.names = "all",
    ncg.type = 'un.supervised',
    ncg.group.names = 'read.scenario3',
    ncg.set.names = 'LinearMixedModel',
    technical.replicates = NULL,
    k = c(1:10),
    apply.log = TRUE,
    data.to.log = 'assay',
    return.wa = TRUE,
    check.se.obj = FALSE)
```

**Step 4: Assessing the W matrix**

The function `assessW()` will be applied to examine the association between the biological and unwanted variables with the unwanted factors estimated by RUV-III.

```
read.se.obj <- assessW(
    se.obj = read.se.obj,
```

```
    compare.w = TRUE,
    variables = NULL,
    bio.variables = c("Tumour.purity", "CMS"),
    uv.variables = c("Time.interval", "Library.size"))
```

Supplementary Figure 24d shows that the columns of the W matrix exhibit very high association with unwanted variation and low association with known biological variation. This indicates that the unsupervised selection of NCGs using the PRPS data failed to adequately capture the known biological variation present in the dataset.

**Step 5: Evaluating the normalizations**

As in scenarios 1 and 2, CMS will first be estimated using the RUV-III normalized datasets, after which the `assessVariation()` function will be applied exclusively to the RUV-III normalized data. Finally, the `assessNormalization()` function will be performed across all available assays contained in the *SummarizedExperiment* object.

```
read.se.obj <- estimateCMS(
    se.obj = read.se.obj,
    assay.names = names(SummarizedExperiment::assays(read.se.obj))[-c(1:4)],
    raw.count.assay.name = NULL,
    apply.log = FALSE,
    tissue.type = 'Tissues',
    remove.na = 'none',
    seed = 122330,
    nb.cores = 10)

all.survivals <- lapply(
    cms <- grep('RUVIIIprps', colnames(colData(read.se.obj)), value = TRUE),
    function(i) {
        idx <- read.se.obj$OS != -1 & read.se.obj[[i]] != 'Not.classified'
        computeSurvival(
            se.obj = read.se.obj[, idx],
            variable = i,
            survival.time = 'OS.time',
            survival.events = 'OS',
            genes = NULL,
            return.survival.plot = TRUE,
            check.se.obj = FALSE,
            save.se.obj = FALSE)$plot
})
```

Supplementary Figure 24 presents the survival analyses of CMS estimated from the RUV-III normalized datasets. RUV-III with $k = 9$ shows a reasonable association between CMS and survival, and therefore will be selected for variation and normalization assessment.

```
read.se.obj$CMS <- read.se.obj$RUVIIIprps_k_9.CMS
read.se.obj <- assessVariation(
    se.obj = read.se.obj,
    assay.names = 'all',
    assessment.level = 'L2',
    bio.variables = c("Tumour.purity", "CMS"),
```

```
    uv.variables = c("Library.size", "Time.interval"),
    pcorr.genes = ppcorr.gene.sets,
    gene.set.score.list = tumour.purity.genes.sets,
    pcorr.filter.genes = FALSE,
    general.points.size = 3,
    override.check = TRUE,
    apply.log = FALSE)

read.se.obj <- assessNormalization(
    se.obj = read.se.obj,
    assay.names = 'all',
    bio.variables = c('CMS', 'Tumour.purity'),
    uv.variables = c('Library.size', 'Time.interval'),
    bio.weight = 0.6,
    uv.weight = 0.4,
    assessment.level = 'L1',
    select.top.ruv = F,
    corr.cutoff = list(Tumour.purity = .5, Library.size = .3),
    output.name = 'tcga.read.assessNormalization.scenario3')

plotAssessVariation(
    se.obj = read.se.obj,
    assay.names = 'all',
    variables =  c('CMS','Library.size','Tumour.purity','Time.interval'),
    output.file.name = "RUVprps_TCGA_READ_VariationAssessment_Secnario3")
```

Supplementary Figure 21e shows the level-2 normalization assessment, demonstrating that RUV-III-PRPS more effectively removes unwanted variation while better preserving known biological signals than the TCGA normalization methods. All corresponding diagnostic plots are provided in Supplementary File TCGA_READ_variationAssessment_Scenario2.pdf. Across all evaluation metrics, RUV-III-PRPS with $k = 7$ exhibits the strongest overall performance among the normalization strategies evaluated.

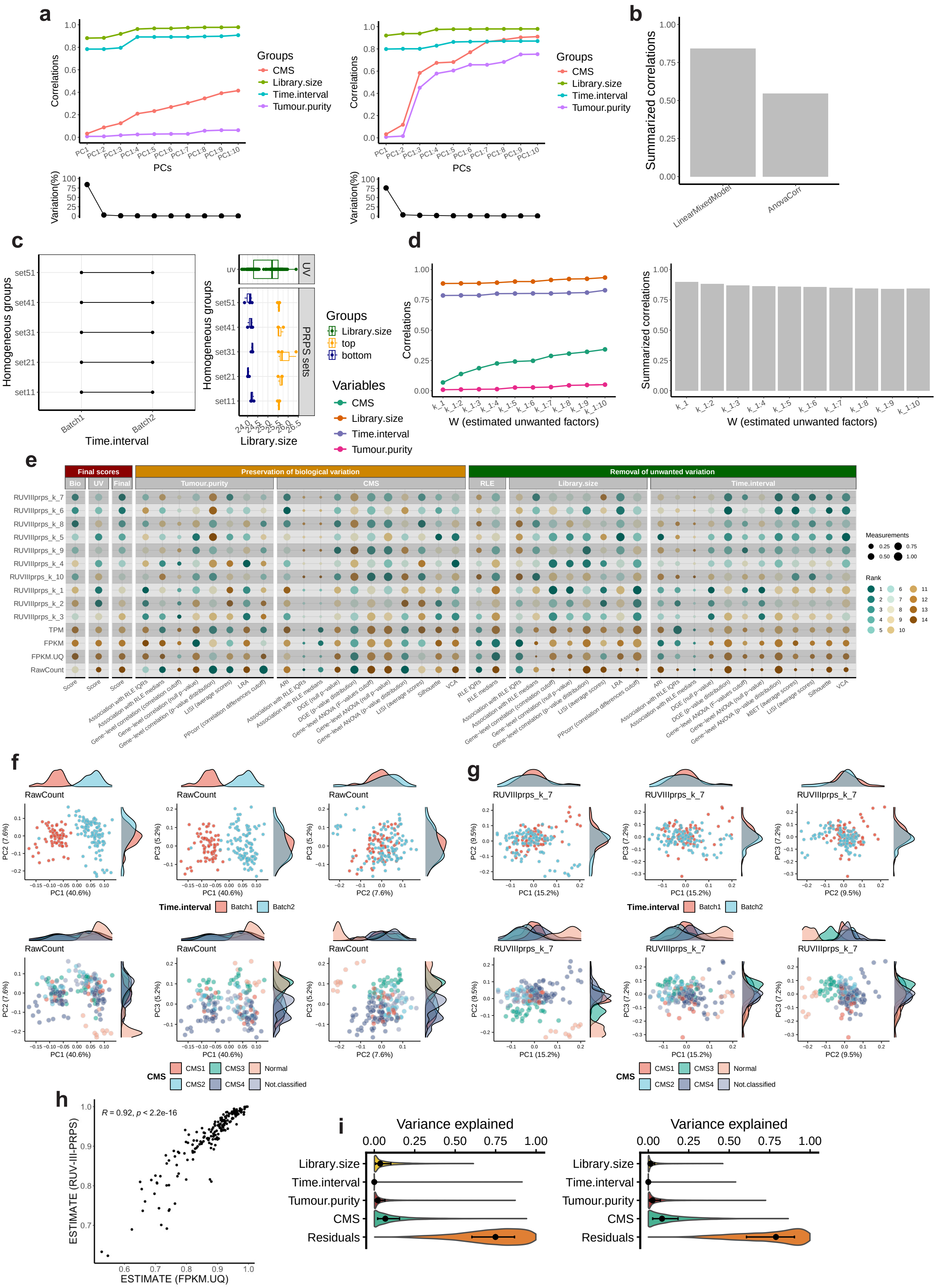
a
Groups
CMS
Library.size
Time.interval
Tumour.purity
Correlations
PCs
Variation(%)
b
Summarized correlations
LinearMixedModel
AnovaCorr
c
Homogeneous groups
Time.interval
Library.size
UV
PRPS sets
Groups
Library.size
top
bottom
d
Variables
CMS
Library.size
Time.interval
Tumour.purity
W (estimated unwanted factors)
e
Final scores
Preservation of biological variation
Removal of unwanted variation
Measurements
Rank
f
RawCount
Time.interval
Batch1
Batch2
CMS
CMS1
CMS2
CMS3
CMS4
Normal
Not.classified
g
RUVIIIprps_k_7
h
R = 0.92, p < 2.2e-16
ESTIMATE (RUV-III-PRPS)
ESTIMATE (FPKM.UQ)
i
Variance explained
Library.size
Time.interval
Tumour.purity
CMS
Residuals

Supplementary Figure 23: RUV-III normalization performance on TCGA READ RNA-seq data for scenario 3. a) Line–dot plots compare the performance of negative control gene (NCG) selection methods applied to the TCGA READ raw count data ($\log_2(x+1)$), including `LinearMixedModel` (first panel), `TwoWayAnova` (second panel), `AnovaCorr.AcrossAllSamples` (third panel) and `AnovaCorr.PerBatchPerBiology` (fourth panel). b) Bar plots summarize the performance of the resulting NCG sets, with higher correlations indicating superior performance. c) PRPS maps are shown for the estimated batches (first panel) and library size (second panel). The y-axis enumerates all homogeneous groups with respect to known biological variation, whereas the x-axes represent sample distributions across estimated batches and library size ranges, respectively. In the estimated batches PRPS map, red points denote groups containing more than two samples, and each horizontal line connecting two red points defines a PRPS set across time intervals. In the library-size PRPS map, each red line corresponds to six samples representing the highest and lowest library sizes within a partially homogeneous biological group. Every three samples are averaged to form a pseudo-sample (PS). The x-axis displays $\log_2(n)$ of the library size and highlights the samples selected for PS construction. d) Line–dot plots depict the associations between columns of the estimated $W$ matrix and both biological and unwanted covariates, with the accompanying bar plots summarizing these correlations; higher values indicate improved performance. e) Quantitative summaries of all evaluation metrics for both biological and unwanted variables are shown. Each point represents a normalization method, with higher values reflecting more effective removal of unwanted variation while preserving biological signal. Normalized datasets are ranked accordingly. f) Principal component plots of the first three components of the TCGA READ raw count data ($\log_2(x+1)$) and RUV-III-PRPS normalized data with $K=15$, colored by estimated batches (top row) and CMS (bottom row). g) As in f, but for data normalized using RUV-III-PRPS with $K=10$. h) The tumor purity estimates for the TCGA FPKM.UQ data and RUV-III-PRPS normalized data. i) Shows the results of a linear mixed model analysis followed by gene-level variation analysis across all the variables for the TCGA FPKM.UQ (first plot) and the RUV-III-PRPS normalized data with $k=10$ (second plot).

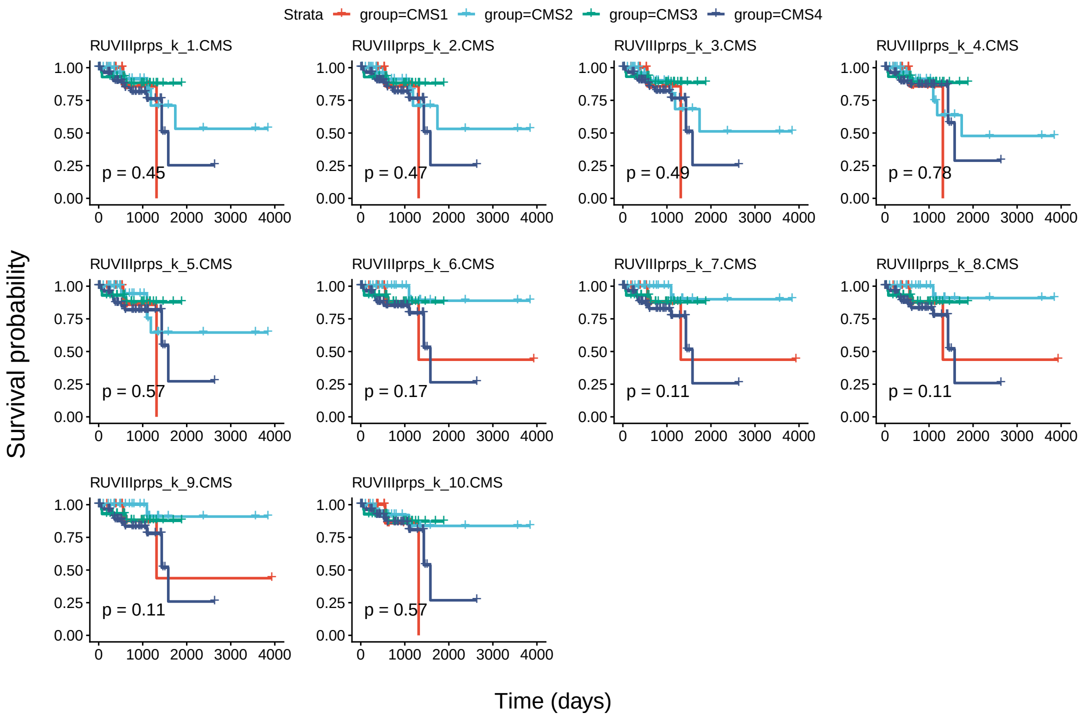


Supplementary Figure 24: Kaplan–Meier analysis of CMS estimated from each RUV-III–normalized dataset in scenario 3 and patient survival in the TCGA READ RNA-seq data.

#### Scenario 4: Both unwanted and biological variation unknown

In the final scenario, it is assumed that both the biological and unwanted variation in the data are completely unknown. Both NCG and PRPS will be identified in an unsupervised manner.

#### Step 1: Selecting negative control genes

A set of NCG will be obtained using both approaches in the `findNcgUnSupervised()` function. The estimated batches and library size will be regarded as sources of unwanted variation.

```
for(i in c( 'AnovaCorr', 'LinearMixedModel')){
    read.se.obj <- findNcgUnSupervised(
        se.obj = read.se.obj,
        assay.name = 'RawCount',
        approach = i,
        nb.ncg = 0.05,
        use.rank = FALSE,
        ncg.selection.method = 'quantile',
        uv.variables = c('Estimated.batches', 'Library.size'),
        form =  ~ (1 | Time.interval) + Library.size,
        bio.percentile = NULL,
        uv.percentile = .8,
        variables.to.assess.ncg = c('Tumour.purity', 'CMS', 'Estimated.batches', 'Library.size'),
        ncg.group.name = 'read.scenario4',
        ncg.set.name = i,
        nb.cores = 15)

read.se.obj <- compareNCGs(
    se.obj = read.se.obj,
    assay.name = 'RawCount',
    apply.log = TRUE,
    variables = NULL,
    bio.variables = c("Tumour.purity", "CMS"),
    uv.variables = c("Estimated.batches", "Library.size"),
    ncg.type = 'un.supervised',
    ncg.group = 'read.scenario4',
    nb.pcs = 10)
```

Supplementary Figure 26a and b exhibit that the performance of NCGs selected by the `LinearMixedModel` approach is significantly better than a set that identified by the `AnovaCorr` approach. This set will be used as an NCG set for RUV-III normalization.

#### Step 2: Creating the PRPS data

Different sets of PRPS will be created for library size and the estimated batches using the the function `createPrPsUnSupervised()` function. Similar to scenario 3, we will first select a set highly variable genes to improved the procedure.

```
# finding a set of highly variable genes
hvg <- findHVG(
    se.obj = read.se.obj,
```

```
    assay.name = 'RawCount',
    approach = 'lmm',
    uv.variables = c('Estimated.batches', 'Library.size'),
    form = ~ (1 | Time.interval) + Library.size,
    nb.hvg = .2,
    nb.cores = 14,
    save.se.obj = FALSE)

uv.variables <- c("Library.size", "Estimated.batches")
max.prps.sets <- c(5, 2)
for(i in c(1:2)){
    read.se.obj <- createPrPsUnSupervised(
        se.obj = read.se.obj,
        assay.name = 'RawCount',
        uv.variables = uv.variables[i],
        other.uv.variables = NULL,
        select.extreme.groups = TRUE,
        approach = 'cca',
        nb.cca = 4,
        nb.pcs = 5,
        nb.clusters = 3,
        clustering.method = 'quantile',
        coordinates.to.use = 'cca',
        max.prps.sets = max.prps.sets[i],
        filter.prps.sets = TRUE,
        hvg = hvg,
        apply.log.for.prps = TRUE,
        create.prps.map = TRUE,
        prps.group.name = 'read.scenario4')}
```

In total, 12 PR sets comprising 24 PS samples were generated to remove the effects of estimated batches, and 5 PR sets comprising 10 PS samples were constructed to account for library size variation. Supplementary Figure 26c illustrates the distribution of the PRPS data across these sources of unwanted variation.

##### Step 3: Performing RUV-III normalization with multiple $k$ values

The `getMaximumK()` function displays that the maximum $k$ for the scenario 4 is 17. Then RUV-III with all possible values of $k$ will be applied using the selected set of NCG and all PRPS data.

```
read.se.obj <- RUVIIIprps(
    se.obj = read.se.obj,
    assay.name = 'RawCount',
    prps.type = 'un.supervised',
    prps.group.names = 'read.scenario4',
    prps.set.names = "all",
    ncg.type = 'un.supervised',
    ncg.group.names = 'read.scenario4',
    ncg.set.names = 'LinearMixedModel',
    technical.replicates = NULL,
    k = c(1:27),
    apply.log = TRUE,
    data.to.log = 'assay',
```

```
    return.wa = TRUE,
    check.se.obj = FALSE)
```

#### Step 4: Assessing the W matrix

The function `assessW()` will be applied to examine the association between the biological and unwanted variables with the unwanted factors estimated by RUV-III.

```
read.se.obj <- assessW(
    se.obj = read.se.obj,
    compare.w = TRUE,
    variables = NULL,
    bio.variables = c("Tumour.purity", "CMS"),
    uv.variables = c("Estimated.batches", "Library.size"))
```

Supplementary Figure 26d shows that the columns of the W matrix exhibit very high association with unwanted variation and low association with known biological variation. This indicates that the unsupervised selection of NCGs using the PRPS data failed to adequately capture the known biological variation present in the dataset.

#### Step 5: Evaluating Normalizations

As in other scenarios, CMS will first be estimated using the RUV-III normalized datasets, after which the `assessVariation()` function will be applied exclusively to the RUV-III normalized data. Finally, the `assessNormalization()` function will be performed across all available assays contained in the *SummarizedExperiment* object.

```
read.se.obj <- estimateCMS(
    se.obj = read.se.obj,
    assay.names = names(SummarizedExperiment::assays(read.se.obj))[-c(1:4)],
    raw.count.assay.name = NULL,
    apply.log = FALSE,
    tissue.type = 'Tissues',
    remove.na = 'none',
    seed = 122334,
    nb.cores = 10)

all.survivals <- lapply(
    cms <- grep('RUVIIIprps', colnames(colData(read.se.obj)), value = TRUE),
    function(i) {
        idx <- read.se.obj$OS != -1 & read.se.obj[[i]] != 'Not.classified'
        computeSurvival(
            se.obj = read.se.obj[, idx],
            variable = i,
            survival.time = 'OS.time',
            survival.events = 'OS',
            genes = NULL,
            return.survival.plot = TRUE,
            check.se.obj = FALSE,
            save.se.obj = FALSE)$plot})
```

Supplementary Figure 26 presents the survival analyses of CMS estimated from the RUV-III normalized datasets. RUV-III with $k = 10$ shows a reasonable association between CMS and survival, and therefore will be selected for variation and normalization assessment.

```
read.se.obj$CMS <- read.se.obj$RUVIIIprps_k_10.CMS
t1 <- Sys.time()
read.se.obj <- assessVariation(
    se.obj = read.se.obj,
    assay.names = 'all',
    assessment.level = 'L2',
    bio.variables = c("Tumour.purity", "CMS"),
    uv.variables = c("Library.size", "Estimated.batches"),
    pcorr.genes = ppcorr.gene.sets,
    gene.set.score.list = tumour.purity.genes.sets,
    pcorr.filter.genes = FALSE,
    general.points.size = 3,
    override.check = TRUE,
    apply.log = FALSE
    )
t2 <- Sys.time()
t2 - t1

read.se.obj <- assessNormalization(
    se.obj = read.se.obj,
    assay.names = 'all',
    bio.variables = c('CMS', 'Tumour.purity'),
    uv.variables = c('Library.size', 'Estimated.batches'),
    bio.weight = 0.6,
    uv.weight = 0.4,
    assessment.level = 'L2',
    select.top.ruv = F,
    corr.cutoff = list(Tumour.purity = .5, Library.size = .3),
    output.name = 'tcga.read.assessNormalization.scenario4'
    )

plotAssessVariation(
    se.obj = read.se.obj,
    assay.names = 'all',
    variables =  c('CMS','Library.size','Tumour.purity','Time.interval'),
    output.file.name = paste0(figPathSc4, "RUVprps_TCGA_READ_VariationAssessment_Secnario4"))
```

Supplementary Figure 26e shows the level 2 normalization assessment, demonstrating that RUV-III-PRPS more effectively mitigates unwanted variation while preserving known biological signals compared with the TCGA normalization methods. The complete set of plots summarized in Figure 26e is provided in the Supplementary File "TCGA_READ_variationAssessment_Scenario4.pdf." Across all evaluation metrics, RUV-III-PRPS with $k = 10$ exhibits the strongest overall performance among the normalization methods evaluated.

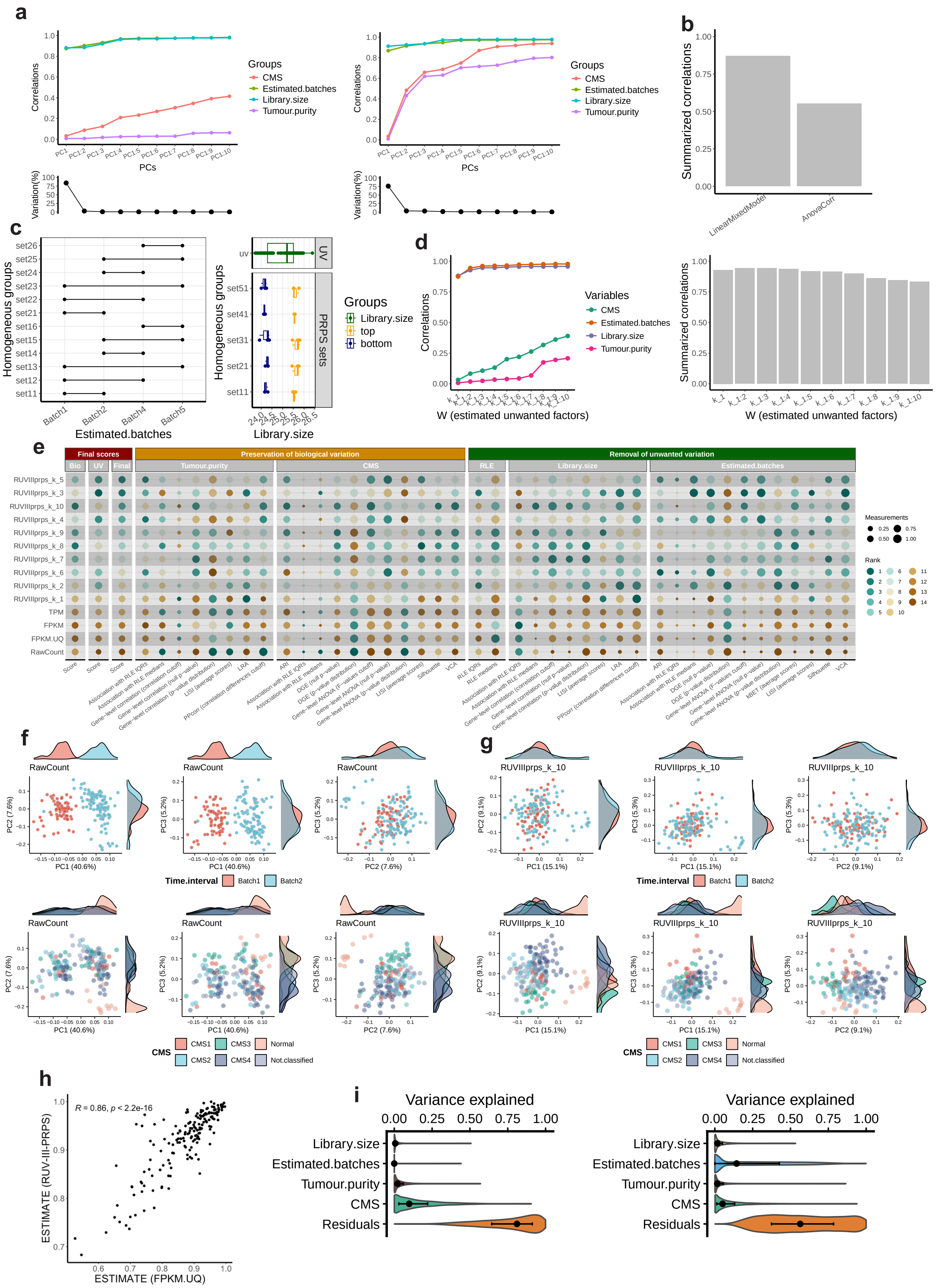
a
b
c
d
e
f
g
h
i
Groups
CMS
Estimated.batches
Library.size
Tumour.purity
Correlations
PCs
Variation(%)
Summarized correlations
LinearMixedModel
AnovaCorr
Homogeneous groups
Estimated.batches
Library.size
top
bottom
UV
PRPS sets
Variables
W (estimated unwanted factors)
Final scores
Preservation of biological variation
Removal of unwanted variation
Bio
UV
Final
Tumour.purity
CMS
RLE
Library.size
Estimated.batches
RUVIIIprps_k_5
RUVIIIprps_k_3
RUVIIIprps_k_10
RUVIIIprps_k_4
RUVIIIprps_k_9
RUVIIIprps_k_8
RUVIIIprps_k_7
RUVIIIprps_k_6
RUVIIIprps_k_2
RUVIIIprps_k_1
TPM
FPKM
FPKM.UQ
RawCount
Measurements
Rank
RawCount
RUVIIIprps_k_10
Time.interval
Batch1
Batch2
CMS1
CMS2
CMS3
CMS4
Normal
Not.classified
PC1 (40.6%)
PC2 (7.6%)
PC3 (5.2%)
PC1 (15.1%)
PC2 (9.1%)
PC3 (5.3%)
R = 0.86, p < 2.2e-16
ESTIMATE (RUV-III-PRPS)
ESTIMATE (FPKM.UQ)
Variance explained
Library.size
Estimated.batches
Tumour.purity
CMS
Residuals

Supplementary Figure 25: RUV-III normalization performance on TCGA READ RNA-seq data for scenario 4. a) Line–dot plots compare the performance of negative control gene (NCG) selection methods applied to the TCGA READ raw count data ($\log_2(x+1)$), including `LinearMixedModel` (first panel), `TwoWayAnova` (second panel), `AnovaCorr.AcrossAllSamples` (third panel) and `AnovaCorr.PerBatchPerBiology` (fourth panel). b) Bar plots summarize the performance of the resulting NCG sets, with higher correlations indicating superior performance. c) PRPS maps are shown for the estimated batches (first panel) and library size (second panel). The y-axis enumerates all homogeneous groups with respect to known biological variation, whereas the x-axes represent sample distributions across estimated batches and library size ranges, respectively. In the estimated batches PRPS map, red points denote groups containing more than two samples, and each horizontal line connecting two red points defines a PRPS set across time intervals. In the library-size PRPS map, each red line corresponds to six samples representing the highest and lowest library sizes within a partially homogeneous biological group. Every three samples are averaged to form a pseudo-sample (PS). The x-axis displays $\log_2(n)$ of the library size and highlights the samples selected for PS construction. d) Line–dot plots depict the associations between columns of the estimated $W$ matrix and both biological and unwanted covariates, with the accompanying bar plots summarizing these correlations; higher values indicate improved performance. e) Quantitative summaries of all evaluation metrics for both biological and unwanted variables are shown. Each point represents a normalization method, with higher values reflecting more effective removal of unwanted variation while preserving biological signal. Normalized datasets are ranked accordingly. f) Principal component plots of the first three components of the TCGA READ raw count data ($\log_2(x+1)$) and RUV-III-PRPS normalized data with $K = 15$, colored by estimated batches (top row) and CMS (bottom row). g) As in f, but for data normalized using RUV-III-PRPS with $K = 10$. h) The tumor purity estimates for the TCGA FPKM.UQ data and RUV-III-PRPS normalized data. i) Shows the results of a linear mixed model analysis followed by gene-level variation analysis across all the variables for the TCGA FPKM.UQ (first plot) and the RUV-III-PRPS normalized data with $k = 10$ (second plot).

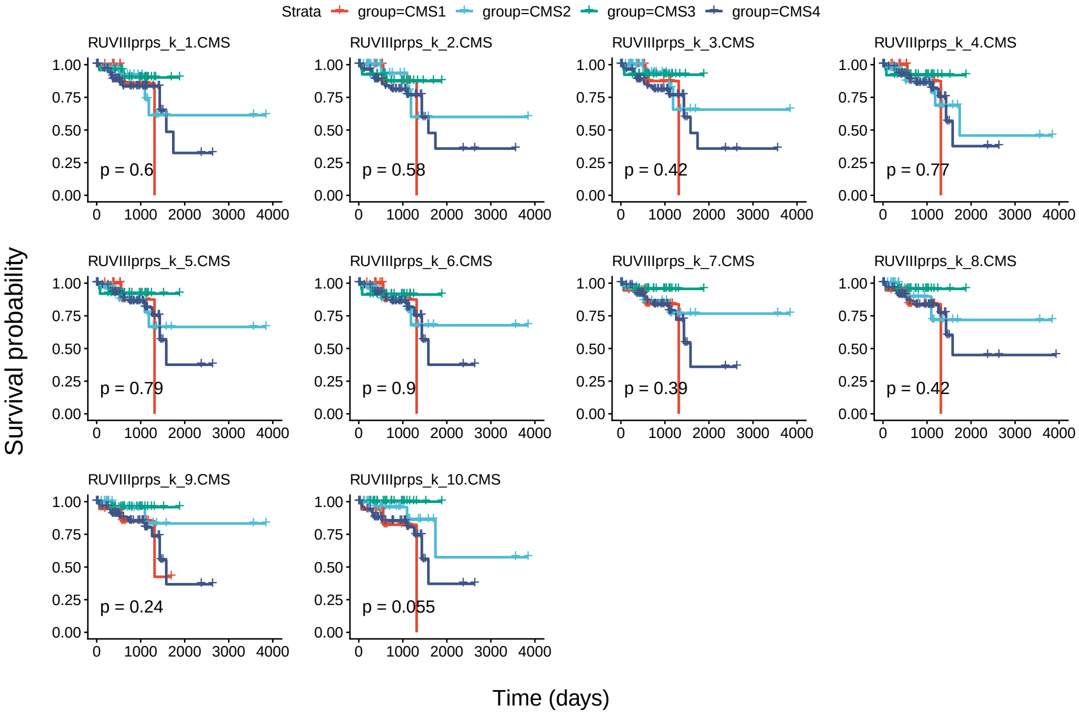


Supplementary Figure 26: Kaplan–Meier analysis of CMS estimated from each RUV-III–normalized dataset in scenario 4 and patient survival in the TCGA READ RNA-seq data.

#### RUV-III normalization of three large breast cancer RNA-seq studies

In this procedure, our goal is to show how to use RUVprps to normalize multiple RNA-seq studies. We will use three large breast cancer RNA-seq studies from TCGA and two studies by Brueffer et al. [14, 15]. Our aim is to remove unwanted variation within and between studies, while preserving known biological variation, including the PAM50 subtypes and tumor purity.

##### Required R packages and installations

The R packages below are required to be installed for this procedure.
The TCGAbiolinks R/Bioconductor package (version 2.28.4) will be used to download the TCGA RNA-seq data from Genomic Data Commons (GDC) data repository.

```
if (!require("BiocManager", quietly = TRUE))
    install.packages("BiocManager")
BiocManager::install("TCGAbiolinks")
```

The GEOquery R package (version 2.74.0) will be employed to download RNA-seq data from the GEO database.

```
if (!require("BiocManager", quietly = TRUE))
    install.packages("BiocManager")
BiocManager::install("GEOquery")
library('GEOquery')
```

##### Stage 1: Data preparation and processing

##### Dataset 1: GSE81538 cohort

##### Step 1: Downloading the data

The log of the FPKM count data of the GSE81538 GEO repository will be downloaded from the GEO data portal using the GEOquery R/Bioconductor package. The required files will be downloaded to the current home directory.

```
# downloading gene expression data
GEOquery::getGEOSuppFiles(GEO = "GSE81538")
data.gse81538 <- data.table::fread(
  input = paste0(
    data.path,
    'GSE81538/',
    'GSE81538_gene_expression_405_transformed.csv.gz')
  ) %>%
  data.frame()
row.names(data.gse81538) <- data.gse81538$V1
data.gse81538 <- data.gse81538[, -1]
data.gse81538 <- as.matrix(data.gse81538)

# downloading sample annotation
Sys.setenv("VROOM_CONNECTION_SIZE" = 7048576 * 2)
geo.obj.gse81538 <- GEOquery::getGEO(GEO = "GSE81538")
sample.annot.gse81538 <- Biobase::pData(object = geo.obj.gse81538[[1]])
indentical(colnames(data.gse81538), sample.annot.gse81538$title)
```

```
row.names(sample.annot.gse81538) <- sample.annot.gse81538$title
```

The `data.gse81538` includes 18,802 genes and 405 samples. Both gene expression and sample annotation data were deposited here [i will add a Zenodo link] for reproducibility. The `sample.annot.gse81538` contains several sample level details including the PAM50 subtypes.

##### Step 2: Removal of lowly expressed genes

The lowly expressed genes will be identified and removed from the data. Please note that, the `prepareSeObj()` function cannot be used for this purpose as the raw count data is not available. A sample annotation containing the PAM50 classification, sample IDs, and study name will be created for downstream analyses.

```
# removing lowly expressed genes
keep.genes <- rowSums(data.gse81538 > -1) >= 22
data.gse81538 <- data.gse81538[keep.genes , ]
colnames(data.gse81538) <- sample.annot.gse81538$geo_accession
row.names(sample.annot.gse81538) <- sample.annot.gse81538$geo_accession

# creating sample annotation
sample.annot.gse81538 <- data.frame(
    sample.ids = row.names(sample.annot.gse81538),
    PAM50 = sample.annot.gse81538$`pam50 subtype:ch1`,
    Studies = rep('GSE81538', nrow(sample.annot.gse81538))
    )
```

Out of 18,802 genes, 15,709 with an expression level of log FPKM $\geq -0.1$ in at least 22 samples, the smallest PAM50 populations, are retained as highly expressed genes in the data. The current gene expression data contains 15,709 genes and 405 samples.

##### Dataset 2: GSE96058 cohort

##### Step 1: Downloading the data

Similar to GSE81538 cohort, the log of the FPKM count data and sample annotation of the GSE96058 cohort will be downloaded from the GEO data portal using the GEOquery R/Bioconductor package. Please note that, the sample annotation contains more samples than the gene expression expression data. Then, the common samples between the two data will be identified and kept for down-stream analysis.

```
# downloading gene expression data
options(timeout = 10^7)
GEOquery::getGEOSuppFiles(GEO = "GSE96058")
data.gse96058 <- data.table::fread(
  input = paste0(
    'GSE96058/',
    "GSE96058_gene_expression_3273_samples_and_136_replicates_transformed.csv.gz")
  ) %>%
  data.frame()
row.names(data.gse96058) <- data.gse96058$V1
data.gse96058 <- data.gse96058[, -1]
```

```
# downloading sample annotation
SSys.setenv("VROOM_CONNECTION_SIZE" = 14048576 * 2)
geo.obj.gse96058 <-  GEOquery::getGEO(GEO = "GSE96058")
sample.annot.gse96058 <- Biobase::pData(object = geo.obj.gse96058[[1]])
row.names(sample.annot.gse96058) <- sample.annot.gse96058$title

# finding common samples between the expression and sample annotation data
common.samples <- intersect(colnames(data.gse96058), sample.annot.gse96058$title)
sample.annot.gse96058 <- sample.annot.gse96058[common.samples , ]
data.gse96058 <- data.gse96058[ , common.samples]
data.gse96058 <- as.matrix(data.gse96058)
colnames(data.gse96058) <- sample.annot.gse96058$geo_accession
row.names(sample.annot.gse96058) <- sample.annot.gse96058$geo_accession
```

The `data.gse96058` includes 30,865 genes and 3409 samples. The sample annotation and gene expression data were deposited on the Zenodo website for reproducibility [i will add the link]. The `sample.annot.gse96058` contains several sample level details including the PAM50 subtypes. There are 3,069 common samples between the sample annotation and gene expression data.

**Step 2: Removal of lowly expressed genes**

A similar gene expression cut-off, log FPKM $\geq -0.1$, will be applied to identify and remove lowly expressed genes from the data. The smallest PAM50 population in this study consists of 202 samples. Furthermore, as described above, a sample annotation containing the PAM50 classification, sample IDs, and study name will be created for downstream analyses.

```
# removing lowly expressed genes
keep.genes <- rowSums(data.gse96058 > -1) >= 202
data.gse96058 <- data.gse96058[keep.genes , ]
colnames(data.gse96058) <- sample.annot.gse96058$geo_accession
row.names(sample.annot.gse96058) <- sample.annot.gse96058$geo_accession

# creating a sample annotation
sample.annot.gse96058 <- data.frame(
    sample.ids = rownames(sample.annot.gse96058),
    PAM50 = sample.annot.gse96058$`pam50 subtype:ch1`,
    Studies = rep('GSE96058', nrow(sample.annot.gse96058 )))
```

Out of 30,865 genes, 18,844 with an expression level of log FPKM $\geq -0.1$ in at least 202 samples, the smallest PAM50 populations, are retained as highly expressed genes. The current gene expression data contains 18844 genes and 3069 samples.

**Dataset 3: TCGA BRCA RNA-seq**

**Step 1: Downloading the data**

We will use the TCGA BRCA *SummarizedExperiment* object that was downloaded and used for normalization of the data.

```
# download gene expression data
tcga.se.obj <- qs::qread('TCGA_BrcaRnaSeq_StarCounts38_TCGAbiolinks.021424.qs')
```

This object includes 60,660 Ensemble gene ids and 1231 samples.

#### Step 2: Adding TCGA batch details

Before adding the TCGA batch information and filtering out lowly expressed genes, the adjacent normal samples will be removed from the TCGA RNA-seq dataset, as the other two datasets do not include this sample type. Furthermore, the TCGA FPKM data will be retained to ensure consistency with the other two datasets, which are also in FPKM format.

```
# keeping the tumour samples
brca.se.obj <- brca.se.obj[ , brca.se.obj$tissue_type == 'Tumor']

# keeping FPKM data
brca.se.obj <- removeAssays(
    se.obj = brca.se.obj,
    assays.to.remove = c('unstranded', 'stranded_first', 'stranded_second', 'tpm_unstrand', 'fpkm_uq_
           unstrand'))

# changing the assay names
brca.se.obj <- renameAssays(se.obj = brca.se.obj, new.names = 'FPKM')

# tidying up the genes ids
brca.se.obj <- tidyGenes(
    se.obj = brca.se.obj,
    keep.gene.type = 'protein_coding',
    gene.type.col.name = 'gene_type',
    remove.duplicates.ids = TRUE,
    ids.col.name = 'gene_name',
    change.row.names = TRUE,
    new.row.names = 'gene_name')
brca.se.obj <- addTcgaBatchInfo(se.obj = brca.se.obj)
brca.se.obj$Years <- as.factor(x = brca.se.obj$Years)
brca.se.obj <- orderSeObj(se.obj = brca.se.obj, factors.to.order = c('Years', 'Plates'))
```

Similar gene expression cut-off, log FPKM $\geq -0.1$, will be applied on the log of the 'fpkm_unstrand' data. The sample size of the smallest PAM50 population is 38 in the data. Further, the duplicated gene ides will be remove as well. A sample annotation containing the PAM50 classification available in *SummarizedExperiment* object., sample IDs, and study name will be created for downstream analyses.

```
keep.genes <- rowSums(log2(assay(x = brca.se.obj, i = 'FPKM') + .1) > -1) >= 38 # 25843
brca.se.obj <- brca.se.obj[keep.genes , ]
brca.sample.annot <- as.data.frame(colData(brca.se.obj))
brca.sample.annot <- brca.sample.annot[ , c('barcode', 'Years', 'paper_BRCA_Subtype_PAM50')]
colnames(brca.sample.annot) <- c('sample.ids', 'sample.ids.numric', 'pam50')
brca.sample.annot$studies <- 'TCGA'
tcag.data <- log2(assay(x = brca.se.obj, i = 'FPKM') + .1)
```

### Stage 2: Merging the data

#### Step 1: Merging the gene expression and sample annotation files

We first need to find common genes across all the studies. Then, we will merge the datasets for down-stream analyses.

```
common.genes <- Reduce(
  intersect,
```

```
  list(
    a = row.names(data.tcga),
    b = row.names(data.gse96058),
    c = row.names(data.gse81538)))
```

A total of 13,665 genes are common between all the three studies. Further, the individual sample annotation will be merged as well.

```
merged.expr.data <- cbind(
    data.tcga[common.genes , ],
    data.gse81538[common.genes , ],
    data.gse96058[common.genes , ])

merged.sample.annot <- rbind(
    sample.annot.tcga,
    sample.annot.gse81538,
    sample.annot.gse96058)
row.names(merged.sample.annot) <- merged.sample.annot$sample.ids
```

Now, the function `prepareSeObj()` will be applied on the merged expression data and sample annotation to create a *SummarizedExperiment* object. Tumor purity will be estimated using both the ESTIMATE and singscore methods [2, 6].

```
ms.se.obj <- prepareSeObj(
    data = list(FPKM = merged.expr.data),
    sample.annotation = merged.sample.annot,
    create.gene.annotation = TRUE,
    add.gene.details = FALSE,
    gene.group = 'hgnc_symbol',
    scale.singscore.values = TRUE,
    estimate.tumor.purity = 'both',
    assay.name.to.estimate.purity = 'FPKM',
    add.housekeeping.genes = TRUE,
    add.immun.stroma.genes = TRUE)
```

**Step 2: Visualization of study outline (optional)**

The `plotStudyOutline()` function will be used to explore the distribution of the biological and unwanted variables across the samples and studies.

```
# renaming variables for visualization
ms.se.obj <- renameVariables(
    se.obj = ms.se.obj,
    current.names = c('pam50', 'tumour.purity.estimate', 'studies'),
    new.names = c('PAM50', 'Tumour.purity', 'Studies'))

ms.se.obj <- plotStudyOutline(
    se.obj = ms.se.obj,
    variables = c('Studies','PAM50', 'Tumour.purity'))
```

Supplementary Figure 27a show the PAM50 subtypes are well distributed across samples. Supplementary Figure 27b show that the tumor purity estimates are highly correlated. Then the ESTIMATE values will be used for down-stream analyses. It should be noted that singscore tumor purity estimations were re-scaled base on the ESTIMATE score.

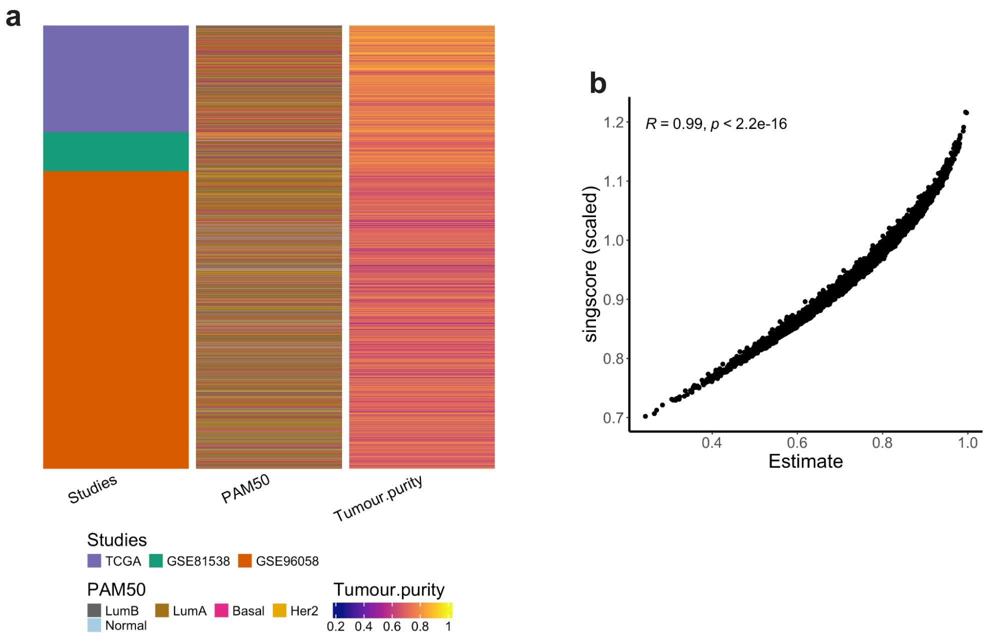


Supplementary Figure 27: Overview of the three breast cancer RNA-seq datasets. a) Heatmap showing 4,538 breast cancer tissue samples collected from three studies. Tumor purity was estimated using the ESTIMATE method on the FPKM expression data. b) Scatter plot showing the correlation between tumor purity estimates obtained using two different methods, ESTIMATE and singscore. The singscore estimates were rescaled based on the ESTIMATE values.

#### Stage 3: Identification of major variation

#### Step 1: Biological populations

In this procedure, the PAM50 subtypes and tumor purity are considered as major sources of biological variation. For the PAM50 subtypes, we will use those provided in each study.

#### Step 2: Unwanted variation

Aside from the evident inter-study batch effects (Fig. 7a), we applied the `identifyUnknownUV()` function to estimate additional, potentially unannotated sources of unwanted variation within each study.

```
ms.se.obj <- identifyUnknownUV(
    se.obj = ms.se.obj,
    assay.name = 'FPKM',
    approach = 'rle',
    chronological.detection = TRUE,
    assess.bio.association = TRUE,
    bio.variables = c('PAM50', 'Tumour.purity'),
    assess.uv.association = TRUE,
    uv.variables = c('Studies'),
    generate.association.plot = TRUE,
    add.to.sample.annotation = TRUE,
```

```
    apply.log = FALSE,
    col.name = 'Estimated.batches',
    output.name = 'ms.estimated.uv')
```

Figure 7c shows that the `identifyUnknownUV()` function found differences between studies and three batches in TCGA data which are mainly time effects in the data.

**Stage 3: Variation assessment**

Here, the impact of both biological and unwanted variation will be assessed on the FPKM data. This step will help to explore the impact of the biological and unwanted variation on the data before applying RUV-III.

**Step 1: Checking the SummarizedExperiment object**

The `checkSeObj()` function will be applied to remove any `NA` or missing values in both the assays and the variables. Recall that the current RUV-III method does not support `NA` or missing values in the data.

```
ms.se.obj <- checkSeObj(
  se.obj = ms.se.obj,
  assay.names = 'all',
  variables = c('PAM50','Tumour purity','Studies', 'Estimated batches'),
  remove.na = 'both',
  verbose = TRUE)
```

**Step 2: Creating all possible assessments**

The `getAssessmentMetrics()` function can be used to find all possible plots and assessments that RUVprps will perform for each selected variables. Any plots and assessments that are not of interest can be excluded for the later steps. We recommend generating all the plots to visually explore the variation and then deciding which ones to exclude for RUV-III normalization.

```
bio.variables <- c('PAM50', 'Tumour purity')
uv.variables <- c('Studies', 'Estimated batches')

# Visualize all possible assessment metrics for the variables
ms.se.obj <- getAssessmentMetrics(
    se.obj = ms.se.obj,
    variables = c(bio.variables, uv.variables),
    plot.output = FALSE
    )
```

Supplementary Figure 28 summarizes the full set of assessment metrics that will be utilized by RUVprps to evaluate the influence of both biological and unwanted sources of variation.

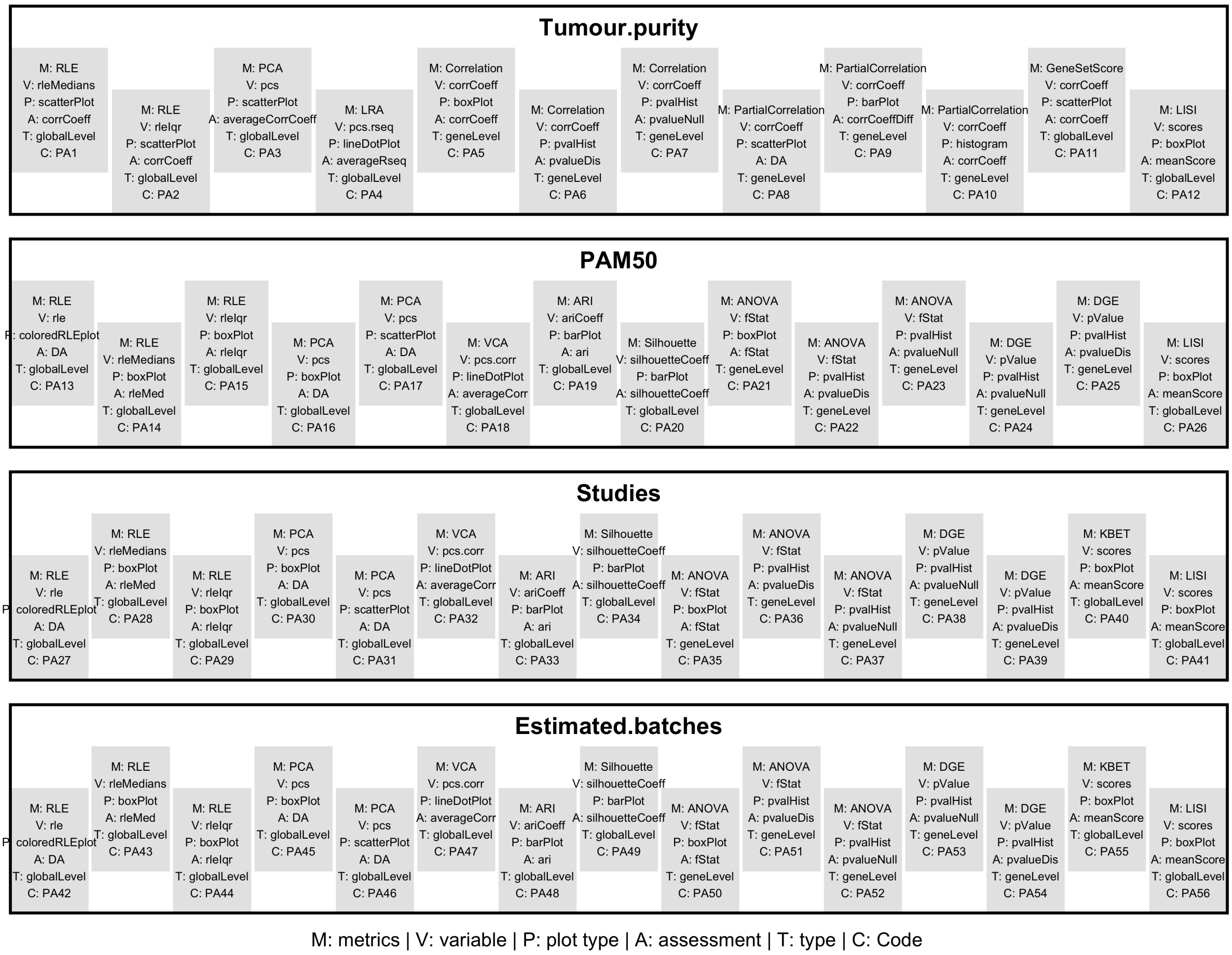


Supplementary Figure 28: Assessment matrices for all the biological and unwanted variation for the normalization of the large breast cancer studies For each individual variable, the specified metrics will be computed and the corresponding plots will be generated. The code of each assessment can be used to exclude the corresponding metrics from the assessment of variation and normalization steps. The 'DA' (Does not Apply) indicates that specific metrics will not be considered for assessing normalization.

#### Step 3: Assessing variation

The `assessVariation()` function applies all the statistical metrics for each specified variable. The individual metrics are stored in the *SummarizedExperiment* object. Then, the `plotAssessVariation()` function creates a PDF file that contains all the plots for visual investigation.

**Step 4: Selecting genes for gene-gene correlation assessment and tumor purity estimate**

Forming partial gene-gene correlations is one of the tools in the variation assessment process. If this is selected, it is desirable to provide a list of genes whose expression is highly associated with the corresponding variable; otherwise, all genes will be used, which is computationally expensive.

Here, a list involving a gene set for the purity variation will be identified.

```
ms.se.obj$Studies <- factor(ms.se.obj$Studies)
ppcorr.gene.sets <- selectGenesForPPcorr(
    se.obj = ms.se.obj,
    assay.names = 'FPKM',
    variables = 'Tumour.purity',
    cor.cutoff = 0.5,
    groups = 'Studies',
    abs.cor = TRUE,
    apply.log = FALSE,
    check.se.object = FALSE)

tumour.purity.genes.sets <- selectGenesSets(
    se.obj = ms.se.obj,
    gene.set = "immune.stromal")
```

The 1,110 genes that have an absolute correlation more than 0.5 with the tumor purity variation are selected. Now, the `assessVariation()` function will be applied.

```
ms.se.obj <- assessVariation(
    se.obj = ms.se.obj,
    assay.names = 'all',
    assessment.level = 'L2',
    bio.variables = c('PAM50','Tumour.purity'),
    uv.variables = c('Studies', 'Estimated.batches'),
    pcorr.genes = ppcorr.gene.sets,
    gene.set.score.list = tumour.purity.genes.sets,
    apply.log = FALSE,
    override.check = FALSE,
    check.se.obj = FALSE
    )
RUVprps:::plotAssessVariation(
    se.obj = ms.se.obj,
    assay.names = 'all',
    variables = c('PAM50','Tumour.purity', 'Estimated.batches', 'Studies'),
    output.file.name = 'RUVprps_MS_InitialVariationAssessments')
```

Supplementary File "RUVprps_MS_InitialVariationAssessments.pdf" shows all the plots that are produced by the `assessVariation()` function. Notably, the RLE and PC plots of the FPKM data clearly show a substantial variation between the studies. These indicate FPKM was inadequate for normalization of the datasets.

**Stage 4: RUV-III normalization**

RUV-III with PRPS will be employed under two different scenarios. The ultimate aim in each scenario is to preserve known biological variation and remove or mitigate the impact of different sources of unwanted variation. The general process within each scenario comprises of identification of a set of NCGs, PRPS sets and RUV-III normalizations with different values for k.

#### Scenario 1: Both biological and unwanted variation are known

In this scenario, only known sources of biological and unwanted variation will be considered in the RUV-III normalization. Then, studies will be used as a major known source of unwanted variation, while the PAM50 and tumor purity will be recognized as the major biological variation. RUV-III normalization will be conducted in a fully supervised manner.

#### Step 1: Selecting negative control genes

A set of NCGs will be identified through implementation of the `findNcgSupervised()` function. All the available approaches will be applied and then the `compareNCGs()` function will be used to select the most suitable set of NCG.

```
ncg.options <- c(
    'AnovaCorr.AcrossAllSamples',
    'AnovaCorr.PerBatchPerBiology',
    'LinearMixedModel',
    'TwoWayAnova')
for(i in 1:4){
    ms.se.obj <- findNcgSupervised(
        se.obj = ms.se.obj,
        assay.name = 'FPKM',
        bio.variables = c('Tumour.purity', 'PAM50'),
        uv.variables = c('Studies'),
        samples.to.use = 'all',
        approach = ncg.options[i],
        form = ~ (1|Studies) + (1|PAM50) + Tumour.purity,
        use.rank = FALSE,
        bio.percentile = NULL,
        uv.percentile = 0.8,
        pseudo.count = 0.5,
        adjust.data = TRUE,
        adjustment.variables = "uv",
        ncg.selection.method = 'quantile',
        nb.ncg = 0.05,
        ncg.group.name = 'ms.scenario1',
        ncg.set.name = ncg.options[i],
        nb.cores = 15,
        normalization = NULL,
        apply.log = FALSE,
        check.se.obj = FALSE,
        assess.ncg = TRUE)}
```

Supplementary Figure 29a and b show the performance of NCG identification methods. The results indicate that the NCG sets identified by the `TwoWayAnova` approach exhibited better performance compared to the other approaches.

#### Step 2: Creating the PRPS data

The PRPS data will be produced for between the studies using the `createPrPsSupervised()` function.

```
ms.se.obj <- createPrPsSupervised(
    se.obj = ms.se.obj,
```

```
    assay.name = 'FPKM',
    bio.variables = bio.variables,
    uv.variables = uv.variables,
    nb.bio.clusters = 20,
    nb.other.uv.clusters = 3,
    apply.log = FALSE,
    apply.other.uv.variables = FALSE,
    prps.group = 'ms.scenario1',
    verbose = TRUE)
```

In total, 75 PR sets with a total of 200 pseudo-samples were created for removing the batch effects between the studies effects (Supplementary Figure 29c).

**Step 3: Performing RUV-III normalization with multiple $k$ values**

The `getMaximumK()` function is applied to find the possible maximum value for the $k$. This shows that the maximum value for the current NCGs and PRPS data is 75. A range of values for $k$ is then selected and the performance for each $k$ will be evaluated. It should be noted the RUV-III-PRPS normalization is applied on the FPKM assay which is the log of the FPKM count. All the RUV-III normalized data are stored in the SummarizedExperiment object.

```
ms.se.obj <- RUVIIIprps(
    se.obj = ms.se.obj,
    assay.name = 'FPKM',
    prps.type = 'supervised',
    prps.group  = 'ms.scenario1',
    ncg.type = 'supervised',
    ncg.group = 'ms.scenario1',
    ncg.set.names = 'TwoWayAnova',
    k = c(1:10),
    residop.fun = "r2",
    apply.log = FALSE,
    return.wa = TRUE,
    check.se.obj = FALSE)
```

**Step 4: Assessing the W matrix**

Before assessing the performance of normalizations, we examine the association between both biological and unwanted variables with the unwanted factors W estimated by RUV-III.

```
ms.se.obj <- assessW(
    se.obj = ms.se.obj,
    compare.w = TRUE,
    variables = NULL,
    bio.variables = c('PAM50', 'Tumour.purity'),
    uv.variables = 'Studies')
```

Figure 7e shows that the W matrix is highly associated with batch effects but exhibits low association with PAM50 subtypes and tumor purity.

#### Step 5: Evaluating normalizations

To assess the performance of RUV-III normalization, we first apply the `assessVariation()` function, followed by `assessNormalization()`, on all available assays in the *SummarizedExperiment* object.

```
ms.se.obj <- assessVariation(
    se.obj = ms.se.obj,
    assay.names = 'all',
    assessment.level = 'L2',
    bio.variables = c("Tumour.purity", "PAM50"),
    uv.variables = c("Studies"),
    pcorr.genes = ppcorr.gene.sets,
    gene.set.score.list = tumour.purity.genes.sets,
    pcorr.filter.genes = FALSE,
    general.points.size = 2,
    override.check = TRUE,
    apply.log = FALSE,
    check.se.obj = FALSE)

ms.se.obj <- assessNormalization(
    se.obj = ms.se.obj,
    assay.names = 'all',
    assessment.level = 'L2',
    select.top.ruv = FALSE,
    bio.variables = c("Tumour.purity", "PAM50"),
    uv.variables = c("Studies"),
    bio.weight = 0.6,
    uv.weight = 0.4,
    corr.cutoff = list(Tumour.purity = .5),
    output.name = 'ruvprps.ms.assessNormalization.scenario1')

RUVprps:::plotAssessVariation(
    se.obj = ms.se.obj,
    assay.names = 'all',
    variables = c("Tumour.purity", "PAM50", "Studies"),
    output.file.name = figPathSc1, "RUVprps_MS_Scenario1")
```

We refer to the main text and Figure 7 for great details of RUV-III-PRPS normalization performance.

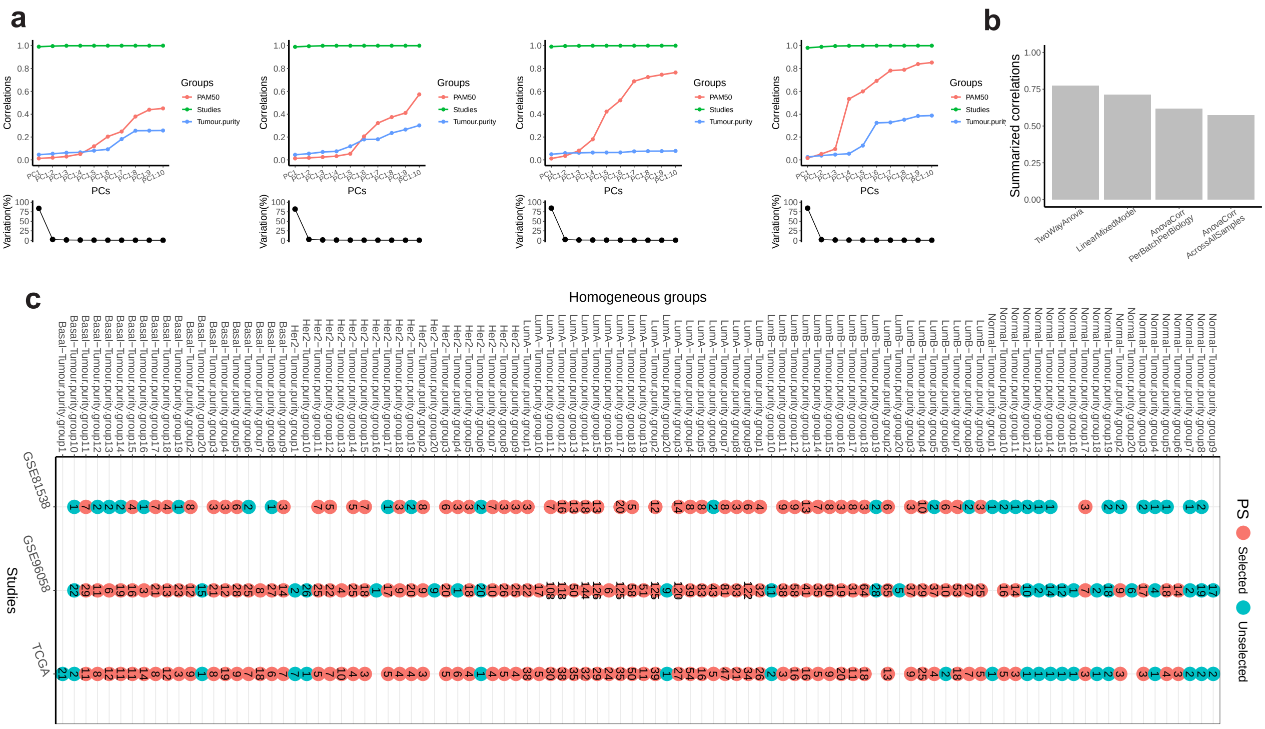


Supplementary Figure 29: NCG and PRPS results for scenario 1 of RUVprps normalization applied to multiple breast cancer RNA-seq datasets. a)Line–dot plots show the performance of NCG selection methods applied to the multiple breast cancer FPKM ($log_2 + 1$)RNA-seq dataset, including `TwoWayAnova` (first panel), `LinearMixedModel` (second panel), `AnovaCorr.PerBatchPerBiology` (third panel) and `AnovaCorr.AcrossAllSamples` (fourth panel). b) Bar plots summarize the performance of the resulting NCG sets, with higher correlations indicating superior performance. c) PRPS maps are shown for the studies. The x-axis enumerates all homogeneous groups with respect to known biological variation, whereas the x-axes represent sample distributions across the studies.

### Scenario 2: Both biological and unwanted variation are unknown

In this scenario, we assume that both biological and unwanted variation are unknown. Then, we will apply RUV-III-PRPS normalization in fully unsupervised manner.

### Step 1: Selecting negative control genes

A set of NCGs is identified through implementation of the `findNcgsUnSupervised()` function. Both approaches in the function will be applied.

```
for(i in c('AnovaCorr', 'LinearMixedModel')){
    ms.se.obj <- findNcgUnSupervised(
        se.obj = ms.se.obj,
        assay.name = 'FPKM',
        approach = i,
        nb.ncg = 0.05,
        use.rank = FALSE,
        ncg.selection.method = 'quantile',
```

```
        uv.variables = c('Estimated.batches'),
        form =  ~ (1 | Estimated.batches),
        bio.percentile = NULL,
        uv.percentile = .9,
        variables.to.assess.ncg = c('Estimated.batches', 'Studies', 'PAM50', 'Tumour.purity'),
        ncg.group.name = 'ms.scenario2',
        ncg.set.name = i,
        apply.log = FALSE,
        normalization = NULL)}

ms.se.obj <- compareNCGs(
    se.obj = ms.se.obj,
    assay.name = 'FPKM',
    ncg.set.names = "all",
    bio.variables = c('PAM50', 'Tumour.purity'),
    uv.variables =  c('Estimated.batches'),
    nb.pcs = 5,
    ncg.type = 'un.supervised',
    apply.log = FALSE,
    ncg.group.name = 'ms.scenario2',
    check.se.obj = FALSE)
```

Supplementary Figure 30a and b show the performance of NCG identification methods. The results indicate that the NCG sets identified by the `LinearMixedModel` method exhibited better performance compared to the other approache.

##### Step 2: Creating the PRPS data

The PRPS data will be produced for between the estimated batches using the `createPrPsUnSupervised()` function. We first find a set of highly variable genes in the data before crating PRPS data.

```
hvg <- findHVG(
    se.obj = ms.se.obj,
    assay.name = 'FPKM',
    approach = 'lmm',
    uv.variables = c('Estimated.batches'),
    form = ~ (1 | Estimated.batches),
    normalization = NULL,
    apply.log = FALSE,
    nb.hvg = .2,
    nb.cores = 14,
    save.se.obj = FALSE)

ms.se.obj <- createPrPsUnSupervised(
    se.obj = ms.se.obj,
    assay.name = 'FPKM',
    uv.variables = 'Estimated.batches',
    other.uv.variables = NULL,
    select.extreme.groups = FALSE,
    approach = 'cca',
    nb.cca = 5,
    nb.pcs = 5,
    nb.clusters = 3,
    coordinates.to.use = 'cca',
```

```
    max.prps.sets = 3,
    normalization = NULL,
    filter.prps.sets = F,
    hvg = hvg,
    apply.log = FALSE,
    apply.log.for.prps = FALSE,
    create.prps.map = TRUE,
    prps.group.name = 'brca.scenario2')
```

In total, 143 PR sets with a total of 286 pseudo-samples were created for removing the batch effects between the studies effects (Supplementary Figure 30c).

**Step 3: Performing RUV-III normalization with multiple $k$ values**

The `getMaximumK()` shows that the maximum value for the current NCGs and PRPS data is 143. A range of values for $k$ is then selected and the performance for each $k$ will be evaluated.

```
ms.se.obj <- RUVIIIprps(
    se.obj = ms.se.obj,
    assay.name = 'FPKM',
    prps.type = 'un.supervised',
    prps.group  = 'brca.scenario2',
    ncg.type = 'un.supervised',
    ncg.group = 'ms.scenario2',
    ncg.set.names = 'LinearMixedModel',
    k = c(1:10),
    apply.log = FALSE,
    return.wa = FALSE,
    check.se.obj = FALSE)
```

**Step 4: Assessing the W matrix**

The association between both biological and unwanted variables with the unwanted factors W estimated by RUV-III will be examined.

```
ms.se.obj <- assessW(
    se.obj = ms.se.obj,
    compare.w = TRUE,
    variables = NULL,
    bio.variables = c('PAM50', 'Tumour.purity'),
    uv.variables = 'Estimated.batches')
```

Figure **??**D shows a high association of the W matrix with the estimated batches and a low association with the biological variables.

**Step 5: Evaluating normalizations**

To assess the performance of RUV-III normalization, we first apply the `assessVariation()` function, followed by `assessNormalization()`, on all available assays in the *SummarizedExperiment* object.

```
ms.se.obj <- assessVariation(
```

```
    se.obj = ms.se.obj,
    assay.names = 'all',
    assessment.level = 'L3',
    bio.variables = c("Tumour.purity", "PAM50"),
    uv.variables = c("Estimated.batches"),
    pcorr.genes = ppcorr.gene.sets,
    gene.set.score.list = tumour.purity.genes.sets,
    pcorr.filter.genes = FALSE,
    general.points.size = 2,
    override.check = TRUE,
    apply.log = FALSE,
    check.se.obj = FALSE,)

ms.se.obj <- assessNormalization(
    se.obj = ms.se.obj,
    assay.names = 'all',
    assessment.level = 'L3',
    select.top.ruv = FALSE,
    bio.variables = c("Tumour.purity", "PAM50"),
    uv.variables = c("Estimated.batches"),
    bio.weight = 0.6,
    uv.weight = 0.4,
    corr.cutoff = list(Tumour.purity = .5),
    output.name = 'ruvprps.ms.assessNormalization.scenario2')

plotAssessVariation(
    se.obj = ms.se.obj,
    assay.names = 'all',
    variables = c("Tumour.purity", "PAM50", "Studies"),
    output.file.name = paste0('../MS/Main_Figures/MutipleStudies_RNAseq/Scenario2/', "RUVprps_MS_
           AssessVariation_Scenario2"))
```

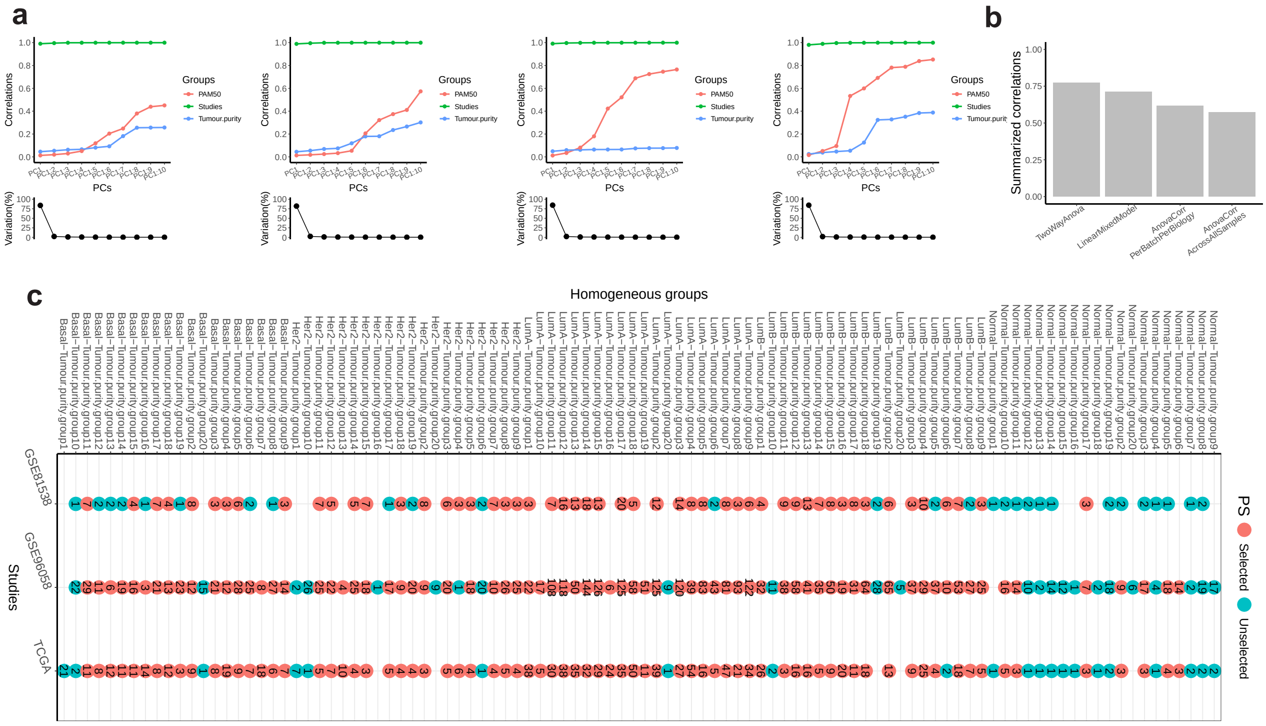

Supplementary Figure 30: NCG and PRPS results for scenario 2 of RUVprps normalization applied to multiple breast cancer RNA-seq datasets. a)Line–dot plots show the performance of NCG selection methods applied to the multiple breast cancer FPKM ($log_2 + 1$)RNA-seq dataset, including `TwoWayAnova` (first panel), `LinearMixedModel` (second panel), `AnovaCorr.PerBatchPerBiology` (third panel) and `AnovaCorr.AcrossAllSamples` (fourth panel). b) Bar plots summarize the performance of the resulting NCG sets, with higher correlations indicating superior performance. c) PRPS maps are shown for the studies. The x-axis enumerates all homogeneous groups with respect to known biological variation, whereas the x-axes represent sample distributions across the studies.
